# The Enzymes that beyond Non-Oxidative Glycolysis

**DOI:** 10.1101/362269

**Authors:** Zixiang Xu, Qiaqing Wu, Dunming Zhu

## Abstract

High yield is an important objective of cell factory. One or several genes cloned into the bacterial may make the synthetic pathway much more optimal, so can increase the yield. But the global benefit enzymes are rare, which can increase the yields of many chemical products for a cell factory such as *E.coli*. Two of these kinds of global benefit enzymes are the famous enzymes, D-fructose-6-phosphate D-erythrose-4-phosphate-lyase and D-Xylulose 5-phosphate D-glyceraldehyde-3-phosphate-lyase, of non-oxidative glycolysis (NOG) published in *Nature*, which can improve the utilization ratio of carbon. We expect to find other global benefit enzymes. We use an integrated model, which integrated *in silico* model of *E.coli* and KEGG. By computation, we analyze the effect of adding each reaction from KEGG on the theoretical yields of several products with *E.coli* and find 83 enzymes that may be potentially global benefit enzymes. By comparison, we find about 30 of the 83 enzymes are better in improving the theoretical yields than the two enzymes of NOG.

In order to compare the global benefit enzymes with NOG, as an example, we select “Glycerol:NADP+ oxidoreductase” (GNO) which can increase the supply of NADPH in *E.coli*. But To increase the supply of reducing power, such as NADPH will probably increase the yield of chemicals in a cell factory. We use flux balance analysis method to testify our assumption. By comparing the maximum yields of 80 products produced by *E.coli* with respectively using GNO and NOG, we find GNO has better performance in the product production of *E.coli*. So GNO is a global benefit enzyme which can increase the yields of many chemical products in *E.coli*.

## Introduction

Synthetic biology has developed rapidly in recent years and the construction of cell factory is one of the main tasks of synthetic biology. For microorganisms to produce a variety of chemicals, cell factories have greatly improved industrial bio-economy. The construction of synthetic pathway is the key aspect for the development of cell factories. If a bacterial can’t produce a chemical natively, some genes should be cloned into the bacterial and make it can produce the chemical. High yield is an important objective of cell factories. Gene knockout and overexpression are two ways to increase the yields of cell factories. Sometimes, we may clone one or several genes into the bacterial and this will make the synthetic pathway much more optimal, so can increase the yield as well. But the global benefit enzymes are rare, which can increase the yields of many chemical products for a cell factory such as *E.coli*. The famous global benefit enzyme is the non-oxidative glycolysis (termed NOG) that was found by James Liao group in the year 2013 and published in Nature [1], which can improve the utilization ratio of carbon. One enzyme name of NOG is D-fructose-6-phosphate D-erythrose-4-phosphate-lyase, its reaction formula is “D-Fructose 6-phosphate + Orthophosphate <=> Acetyl phosphate + D-Erythrose 4-phosphate + H2O” and its KEGG id is R00761; another enzyme name of NOG is “D-Xylulose 5-phosphate D-glyceraldehyde-3-phosphate-lyase”, its reaction formula is “D-Xylulose 5-phosphate + Orthophosphate <=> Acetyl phosphate + D-Glyceraldehyde 3-phosphate + H2O” and its KEGG id is R01621. KEGG (Kyoto Encyclopedia of Genes and Genomes) is a database resource for understanding high-level functions and utilities of the biological system [2]. NOG can increase the yields of many chemical products produced by *E.coli*.

We expect to find other global benefit enzymes and we utilize the computational method. Computation biology has been used as testification method widely. We use an integrated model, which integrated *in silico* model of *E.coli* and KEGG [3]. By computation, we analyze the effect of adding each reaction from KEGG on the theoretical yields of several products and find 83 enzymes that are potentially global benefit enzymes. By comparison, we find about 30 from the 83 enzymes are better in improving the theoretical yields than NOG.

As we know, to increase the supply of, such as NADPH, will probably increase the yields of chemicals. In order to compare the global benefit enzymes with NOG, as an example, we select the enzyme, “Glycerol:NADP+ oxidoreductase” (termed GNO) from the 83 enzymes, its KEGG id is R01041, its reaction formula is “Glycerol + NADP^+^ <=> D-Glyceraldehyde + NADPH + H^+^”, and we make our test in this study. We have compared the maximum yields of 80 products produced by E.coli respectively using GNO and NOG.

## Methods

### About 80 enzymes that are potentially global benefit enzymes

An integrated model [3] was used, which integrated *in silico* model of *E.coli* and KEGG. The reactions from KEGG that added to *E.coli* model were balance checked and reversibility checked. There were 4733 reactions that were added to the *E.coli* model. When we used one reaction from KEGG, the up/low flux bounds were not constrained, while the flux bounds of other reactions were set to zero. So it is equivalent to add one reaction to *E.coli* every time.

*E.coli* can produce many chemicals, and here we did not calculate all the yields of them. We choose Acetate (Ace), L-Glutamate (Glu), L-Lysine (Lys), L-Threonine (Thr), Succinate (Suc), Formate (For), L-Alanine (Ala), L-Tryptophan (Try), L-Phenylalanine (Phe), L-Malate (Mal) as the representatives.

Firstly, we calculated the growth rate (Grw) and the theoretical yields of the above 10 chemicals with wild-type *E.coli*. The following constraints were applied, when we did the calculation: glucose consumption rate was −10 mmol g^−1^(Dw)h^−1^; there was no constraint for oxygen consumption rate. The FBA (flux balance analysis) calculation was carried out by COBRA Toolbox [4] with a loopless function which eliminates all steady-state flux solutions that are incompatible with the loop-law [5]. The optimization solver is Gurobi. The growth rate and the theoretical yields were calculated by setting the corresponding flux as objectives of FBA model.

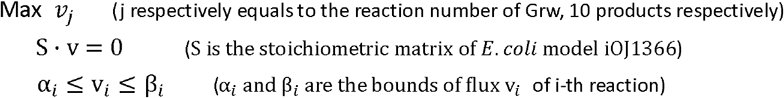

Secondly, we calculated the growth rate and the theoretical yields of above 10 chemicals with *E.coli* but added one reaction every time. That is to say, when we used one reaction from KEGG, the up/low flux bounds were not constrained, while the flux bounds of other reactions were set to zero. The computational condition was the same as above.

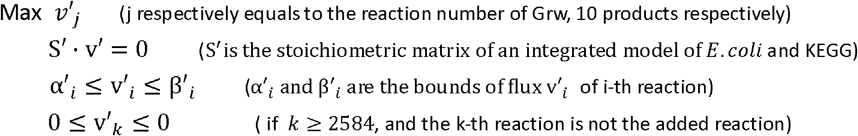

Thirdly, we calculated the growth rate and the theoretical yields of above 10 chemicals with *E.coli* by adding one reaction every time. But for this time, considering the enzyme activity, when we used one reaction from KEGG, up/low flux bounds were constrained to not more than 100 in abstract value. At the same time, we calculated the smallest balanced pathway for every chemical by the tool FIND_tfSBP [8].

### The example of GNO (Glycerol:NADP+ oxidoreductase)

In order to compare the impact of GNO and NOG for the products of *E.coli*, we add both reactions to the model iAF_1260 of *E.coli* [6]. In iAF_1260, there are about 300 exchange reactions, we calculate the maximum yield of each product with FBA (flux balance analysis) method, which has been described in Ref [7]. Here we use the exchange reaction as the objective but not the biomass (growth). The following constraints were applied, when we do the calculation: glucose consumption rate is 10 mmol g^−1^ (Dw)h^−1^; there is no constraint for oxygen consumption rate. The FBA calculation was carried out with COBRA Toolbox [4]. Many of the exchange reactions take no flux, while about 80 exchange reactions have fluxes, i.e. *E.coli* may produce about 80 products actually according to the model of iAF_1260.

We first add NOG (R00761) into the iAF_1260 model and calculate, one by one, the maximum yields of all the 80 products which *E.coli* may produce actually. Then, we add GNO into the iAF_1260 model and calculate the maximum yields of all the 80 products with the same method. We compare GNO and NOG, and hope to find out which will get a larger number with higher yields. We classify the result into three groups, i.e. “NOG is better” (product yields are higher when adding NOG), “GNO is better” (product yields are higher when adding GNO) and “No significant difference between NOG and GNO” (product yields are nearer when adding NOG or GNO).

## Results and Discussion

### 83 enzymes that are potentially global benefit enzymes

The growth rate and the theoretical yields of 10 chemicals, i.e. Acetate (Ace), L-Glutamate (Glu), L-Lysine (Lys), L-Threonine (Thr), Succinate (Suc), Formate (For), L-Alanine (Ala), L-Tryptophan (Try), L-Phenylalanine (Phe), L-Malate (Mal), were obtained with wild-type *E.coli* and with *E.coli* but added one reaction every time (4733 cases in total). In order to find which reactions added would make the theoretical yields of 10 chemicals improved, if any one of the 10 theoretical yields of *E.coli* (added one reaction every time) is larger than (1.01 times) the corresponding value of wild-type *E.coli*, we select out the enzyme (reaction), and we get 83 enzymes (reactions) in total. We list them in **Table 1** with descending order in Ace and Glu yield values. The whole names and equations of these reactions are illustrated in **Supplementary Table S1**. These 83 enzymes are enzymes that have potential ability to improve the theoretical yields of some chemical products in *E.coli*.

**Table 1.** Reactions that have potential ability to improve the theoretical yields of 10 chemical products in *E.coli*.

| No. | KEGG id | Rev | Growth | Ace | Glu | Lys | Thr | Suc | For | Ala | Try | Phe | Mal |
| --- | --- | --- | --- | --- | --- | --- | --- | --- | --- | --- | --- | --- | --- |
|  | Wild_Type |  | 0.982 | 29.093 | 11.937 | 7.833 | 12.853 | 17.096 | 103.740 | 20.000 | 4.622 | 5.507 | 18.412 |
| 1 | R07409 | rev | 6.664 | 207.919 | 80.332 | 60.942 | 98.184 | 132.953 | 1000.000 | 164.401 | 30.603 | 3.222 | 131.302 |
| 2 | R00790 | rev | 6.677 | 195.829 | 76.028 | 56.440 | 93.812 | 118.034 | 1000.000 | 144.410 | 29.143 | 35.589 | 122.509 |
| 3 | R00659 | rev | 1.384 | 30.000 | 13.333 | 8.571 | 15.000 | 17.143 | 120.000 | 20.000 | 5.217 | 6.000 | 20.000 |
| 4 | R00206 | rev | 1.384 | 30.000 | 13.333 | 8.571 | 15.000 | 17.143 | 120.000 | 20.000 | 5.217 | 6.000 | 20.000 |
| 5 | R02537 | rev | 1.384 | 30.000 | 13.333 | 8.571 | 15.000 | 17.143 | 120.000 | 20.000 | 5.217 | 6.000 | 20.000 |
| 6 | R01050 | rev | 1.384 | 30.000 | 13.333 | 8.571 | 15.000 | 17.143 | 120.000 | 20.000 | 5.217 | 6.000 | 20.000 |
| 7 | R00709 | rev | 1.384 | 30.000 | 13.333 | 8.571 | 15.000 | 17.143 | 120.000 | 20.000 | 5.217 | 6.000 | 20.000 |
| 8 | R00149 | rev | 1.384 | 30.000 | 13.333 | 8.571 | 15.000 | 17.143 | 120.000 | 20.000 | 5.217 | 6.000 | 20.000 |
| 9 | R00224 | rev | 1.384 | 30.000 | 13.333 | 8.571 | 15.000 | 17.143 | 120.000 | 20.000 | 5.217 | 6.000 | 20.000 |
| 10 | R08515 | rev | 1.384 | 30.000 | 13.333 | 8.571 | 15.000 | 17.143 | 120.000 | 20.000 | 5.217 | 6.000 | 20.000 |
| 11 | R01224 | rev | 1.384 | 30.000 | 13.333 | 8.571 | 15.000 | 17.143 | 120.000 | 20.000 | 5.217 | 6.000 | 20.000 |
| 12 | R00572 | rev | 1.384 | 30.000 | 13.333 | 8.571 | 15.000 | 17.143 | 120.000 | 20.000 | 5.217 | 6.000 | 20.000 |
| 13 | R03004 | rev | 1.384 | 30.000 | 13.333 | 8.571 | 15.000 | 17.143 | 120.000 | 20.000 | 5.217 | 6.000 | 20.000 |
| 14 | R02301 | rev | 1.384 | 30.000 | 13.333 | 8.571 | 15.000 | 17.143 | 120.000 | 20.000 | 5.217 | 6.000 | 20.000 |
| 15 | R02145 | rev | 1.384 | 30.000 | 13.333 | 8.571 | 15.000 | 17.143 | 120.000 | 20.000 | 5.217 | 6.000 | 20.000 |
| 16 | R07159 | rev | 1.384 | 30.000 | 13.333 | 8.571 | 15.000 | 17.143 | 120.000 | 20.000 | 5.217 | 6.000 | 20.000 |
| 17 | R01138 | rev | 1.384 | 30.000 | 13.333 | 8.571 | 15.000 | 17.143 | 120.000 | 20.000 | 5.217 | 6.000 | 20.000 |
| 18 | R00397 | rev | 1.384 | 30.000 | 13.333 | 8.571 | 15.000 | 17.143 | 120.000 | 20.000 | 5.217 | 6.000 | 20.000 |
| 19 | R07758 | rev | 1.384 | 30.000 | 13.333 | 8.571 | 15.000 | 17.143 | 120.000 | 20.000 | 5.217 | 6.000 | 20.000 |
| 20 | R00724 | rev | 1.384 | 30.000 | 13.333 | 8.571 | 15.000 | 17.143 | 120.000 | 20.000 | 5.217 | 6.000 | 20.000 |
| 21 | R06846 | rev | 1.384 | 30.000 | 13.333 | 8.571 | 15.000 | 17.143 | 120.000 | 20.000 | 5.217 | 6.000 | 20.000 |
| 22 | R00430 | rev | 1.384 | 30.000 | 13.333 | 8.571 | 15.000 | 17.143 | 120.000 | 20.000 | 5.217 | 6.000 | 20.000 |
| 23 | R00885 | rev | 1.384 | 30.000 | 13.333 | 8.571 | 15.000 | 17.143 | 120.000 | 20.000 | 5.217 | 6.000 | 20.000 |
| 24 | R00522 | rev | 1.384 | 30.000 | 13.333 | 8.571 | 15.000 | 17.143 | 120.000 | 20.000 | 5.217 | 6.000 | 20.000 |
| 25 | R01858 | rev | 1.384 | 30.000 | 13.333 | 8.571 | 15.000 | 17.143 | 120.000 | 20.000 | 5.217 | 6.000 | 20.000 |
| 26 | R00126 | rev | 1.384 | 30.000 | 13.333 | 8.571 | 15.000 | 17.143 | 120.000 | 20.000 | 5.217 | 6.000 | 20.000 |
| 27 | R00502 | rev | 1.384 | 30.000 | 13.333 | 8.571 | 15.000 | 17.143 | 120.000 | 20.000 | 5.217 | 6.000 | 20.000 |
| 28 | R01217 | rev | 1.384 | 30.000 | 13.333 | 8.571 | 15.000 | 17.143 | 120.000 | 20.000 | 5.217 | 6.000 | 20.000 |
| 29 | R01197 | rev | 0.990 | 30.000 | 13.189 | 7.833 | 12.895 | 17.143 | 104.740 | 20.000 | 4.626 | 5.507 | 18.403 |
| 30 | R00265 | rev | 0.982 | 30.000 | 13.073 | 7.833 | 12.856 | 17.096 | 104.740 | 20.000 | 4.399 | 5.507 | 18.412 |
| 31 | R01621 | irr | 0.982 | 30.000 | 12.433 | 7.833 | 12.853 | 17.096 | 103.740 | 20.000 | 4.622 | 5.507 | 18.412 |
| 32 | R00761 | irr | 0.982 | 30.000 | 12.433 | 7.833 | 12.853 | 17.096 | 103.740 | 20.000 | 4.622 | 5.507 | 18.412 |
| 33 | R01817 | rev | 0.990 | 30.000 | 12.208 | 8.135 | 13.441 | 17.096 | 109.800 | 20.000 | 4.641 | 5.580 | 18.412 |
| 34 | R00834 | rev | 0.990 | 30.000 | 12.208 | 8.135 | 13.441 | 17.096 | 109.800 | 20.000 | 4.641 | 5.580 | 18.412 |
| 35 | R07164 | rev | 0.990 | 30.000 | 12.208 | 8.135 | 13.441 | 17.096 | 109.800 | 20.000 | 4.641 | 5.580 | 18.412 |
| 36 | R01063 | rev | 0.990 | 30.000 | 12.208 | 8.135 | 0.000 | 17.096 | 109.800 | 20.000 | 4.641 | 5.580 | 18.412 |
| 37 | R09281 | rev | 0.990 | 30.000 | 12.208 | 8.135 | 13.441 | 17.096 | 109.800 | 20.000 | 4.641 | 5.580 | 18.412 |
| 38 | R01434 | rev | 0.990 | 30.000 | 12.208 | 8.135 | 13.441 | 17.096 | 109.800 | 20.000 | 4.641 | 5.580 | 18.412 |
| 39 | R07140 | rev | 0.990 | 30.000 | 12.208 | 8.135 | 13.441 | 17.096 | 109.800 | 20.000 | 4.641 | 5.580 | 18.412 |
| 40 | R01976 | rev | 0.990 | 30.000 | 12.208 | 8.135 | 13.441 | 17.096 | 109.800 | 20.000 | 4.641 | 5.580 | 18.412 |
| 41 | R00978 | rev | 0.990 | 30.000 | 12.208 | 8.135 | 13.441 | 17.096 | 109.800 | 20.000 | 4.641 | 5.580 | 18.412 |
| 42 | R01218 | rev | 0.990 | 30.000 | 12.208 | 8.135 | 13.441 | 17.096 | 109.800 | 20.000 | 4.641 | 5.580 | 18.412 |
| 43 | R00396 | rev | 0.990 | 30.000 | 12.208 | 8.135 | 13.441 | 17.096 | 109.800 | 20.000 | 4.641 | 5.580 | 18.412 |
| 44 | R00094 | rev | 0.990 | 30.000 | 12.208 | 8.135 | 13.441 | 17.096 | 109.800 | 20.000 | 4.641 | 5.580 | 18.412 |
| 45 | R00688 | rev | 0.990 | 30.000 | 12.208 | 8.135 | 13.441 | 17.096 | 109.800 | 20.000 | 4.641 | 5.580 | 18.412 |
| 46 | R01738 | rev | 0.990 | 30.000 | 12.208 | 8.135 | 13.441 | 17.096 | 109.800 | 20.000 | 4.641 | 5.580 | 18.412 |
| 47 | R00343 | rev | 0.990 | 30.000 | 12.208 | 8.135 | 13.441 | 17.096 | 106.243 | 20.000 | 4.641 | 5.580 | 18.412 |
| 48 | R01747 | rev | 0.990 | 30.000 | 12.208 | 8.135 | 13.441 | 17.096 | 109.800 | 20.000 | 4.641 | 5.580 | 18.412 |
| 49 | R00746 | rev | 0.990 | 30.000 | 12.208 | 8.135 | 13.441 | 17.096 | 109.800 | 20.000 | 4.641 | 5.580 | 18.412 |
| 50 | R01773 | rev | 0.990 | 30.000 | 12.208 | 8.135 | 13.441 | 17.096 | 109.800 | 20.000 | 4.641 | 5.580 | 18.412 |
| 51 | R02196 | rev | 0.990 | 30.000 | 12.208 | 8.135 | 13.441 | 17.096 | 109.800 | 20.000 | 4.641 | 5.580 | 18.412 |
| 52 | R00936 | rev | 0.990 | 30.000 | 12.208 | 8.135 | 13.441 | 17.096 | 109.800 | 20.000 | 4.641 | 5.580 | 18.412 |
| 53 | R00243 | rev | 0.990 | 30.000 | 12.208 | 8.135 | 13.441 | 17.096 | 109.800 | 20.000 | 4.641 | 5.580 | 18.412 |
| 54 | R02259 | rev | 0.990 | 30.000 | 12.208 | 8.135 | 13.441 | 17.096 | 109.800 | 20.000 | 4.641 | 5.346 | 18.412 |
| 55 | R00842 | rev | 0.990 | 30.000 | 12.208 | 8.135 | 13.225 | 17.096 | 109.800 | 20.000 | 4.641 | 5.580 | 18.412 |
| 56 | R06847 | rev | 0.990 | 30.000 | 12.208 | 8.135 | 13.441 | 17.096 | 109.800 | 20.000 | 4.641 | 5.580 | 18.412 |
| 57 | R02577 | rev | 0.990 | 30.000 | 12.208 | 8.135 | 13.441 | 17.096 | 109.800 | 20.000 | 4.641 | 5.580 | 18.412 |
| 58 | R07759 | rev | 0.990 | 30.000 | 12.208 | 8.135 | 13.441 | 17.096 | 109.800 | 20.000 | 4.641 | 5.580 | 18.412 |
| 59 | R01041 | rev | 0.990 | 30.000 | 12.208 | 8.135 | 13.441 | 17.096 | 109.800 | 20.000 | 4.641 | 5.580 | 18.412 |
| 60 | R00848 | rev | 0.990 | 30.000 | 12.208 | 8.135 | 13.441 | 17.096 | 109.800 | 20.000 | 4.641 | 5.580 | 18.412 |
| 61 | R02165 | rev | 1.040 | 30.000 | 12.118 | 7.960 | 13.227 | 17.143 | 118.425 | 20.000 | 4.727 | 5.592 | 18.668 |
| 62 | R02965 | rev | 1.040 | 30.000 | 12.118 | 7.960 | 13.227 | 17.143 | 100.436 | 20.000 | 4.727 | 5.592 | 18.668 |
| 63 | R01058 | rev | 1.384 | 30.000 | 10.000 | 8.571 | 15.000 | 17.143 | 120.000 | 20.000 | 5.217 | 6.000 | 20.000 |
| 64 | R02639 | rev | 0.990 | 30.000 | 0.000 | 8.135 | 13.441 | 17.096 | 109.800 | 20.000 | 4.641 | 5.580 | 18.412 |
| 65 | R00726 | rev | 0.997 | 29.943 | 12.522 | 8.134 | 13.556 | 17.143 | 104.740 | 20.000 | 4.623 | 5.507 | 19.945 |
| 66 | R00346 | rev | 0.997 | 29.943 | 12.522 | 8.134 | 13.556 | 17.143 | 104.740 | 20.000 | 4.623 | 5.507 | 19.945 |
| 67 | R00431 | rev | 0.997 | 29.943 | 12.522 | 8.134 | 13.556 | 17.143 | 104.740 | 20.000 | 4.623 | 5.507 | 19.148 |
| 68 | R01967 | rev | 0.982 | 29.753 | 12.102 | 7.833 | 12.029 | 17.096 | 103.740 | 20.000 | 4.473 | 5.507 | 18.412 |
| 69 | R02089 | rev | 0.982 | 29.753 | 12.102 | 7.833 | 12.853 | 17.096 | 103.740 | 20.000 | 4.622 | 5.507 | 18.412 |
| 70 | R01010 | rev | 1.014 | 29.737 | 12.247 | 8.011 | 13.044 | 17.143 | 104.309 | 20.000 | 4.710 | 5.652 | 19.181 |
| 71 | R01059 | rev | 0.982 | 29.714 | 12.049 | 7.895 | 12.938 | 17.096 | 106.740 | 20.000 | 4.625 | 5.523 | 18.412 |
| 72 | R07417 | rev | 0.982 | 29.714 | 12.049 | 7.895 | 12.938 | 17.096 | 106.740 | 20.000 | 4.625 | 5.523 | 18.412 |
| 73 | R00585 | rev | 0.982 | 29.406 | 12.012 | 7.833 | 12.853 | 17.096 | 103.740 | 20.000 | 4.622 | 5.507 | 18.412 |
| 74 | R00941 | rev | 0.983 | 29.391 | 11.991 | 7.833 | 12.853 | 17.096 | 103.740 | 20.000 | 4.622 | 5.507 | 18.412 |
| 75 | R00519 | rev | 0.982 | 29.279 | 11.971 | 7.833 | 12.853 | 17.096 | 117.671 | 20.000 | 4.622 | 5.507 | 18.412 |
| 76 | R01199 | rev | 0.986 | 29.225 | 11.961 | 7.833 | 12.853 | 17.096 | 114.309 | 20.000 | 4.622 | 5.507 | 18.412 |
| 77 | R00210 | rev | 0.982 | 29.131 | 11.946 | 7.833 | 12.856 | 17.096 | 113.740 | 20.000 | 4.623 | 5.507 | 18.412 |
| 78 | R09280 | rev | 0.987 | 29.093 | 11.937 | 7.833 | 12.870 | 17.096 | 105.740 | 20.000 | 4.628 | 5.507 | 18.412 |
| 79 | R00352 | rev | 1.001 | 29.093 | 12.345 | 7.833 | 12.895 | 17.143 | 107.945 | 20.000 | 4.638 | 5.507 | 18.525 |
| 80 | R04198 | rev | 0.983 | 29.093 | 11.937 | 7.945 | 12.853 | 17.096 | 103.740 | 20.000 | 4.622 | 5.507 | 18.412 |
| 81 | R07613 | rev | 0.984 | 29.093 | 11.937 | 8.011 | 12.853 | 17.096 | 103.740 | 20.000 | 4.622 | 5.507 | 18.412 |
| 82 | R06975 | rev | 0.000 | 29.093 | 11.937 | 7.833 | 12.531 | 17.096 | 109.740 | 20.000 | 4.622 | 5.507 | 18.412 |
| 83 | R00859 | rev | 0.983 | 29.093 | 11.937 | 7.639 | 12.853 | 17.096 | 110.381 | 20.000 | 4.622 | 5.507 | 18.412 |

\*: All the rate unit is $\text{mmol g}^{-1}(\text{Dw})\text{h}^{-1}$ ; glucose consumption rate is $-10 \text{ mmol g}^{-1}(\text{Dw})\text{h}^{-1}$ ; there is no constraint for oxygen consumption rate;
\*: Rev: Reversibility; Acetate(Ace), L-Glutamate(Glu), L-Lysine(Lys), L-Threonine(Thr), Succinate(Suc), Formate(For) , L-Alanine (Ala), L-Tryptophan (Try), L-Phenylalanine (Phe), L-Malate (Mal).
\*: Descending order in Ace and Glu values.

Among these 83 enzymes, many can get Acetate yield to 3.0 (wild-type *E.coli* is 2.9), L-Glutamate yield to 1.33 (wild-type *E.coli* is 1.19), L-Lysine yield to 0.86 (wild-type *E.coli* is 0.78), L-Threonine yield to 1.5 (wild-type *E.coli* is 1.28), and Formate yield to 12 (wild-type *E.coli* is 10.3), but the yield of Succinate was not improved significantly. Many can improve the growth of the cell as well and the growth rate can get to 1.38, while the growth rate of wild-type *E.coli* is 0.98.

Two reactions of NOG, R00761 and R01621, are on the list of **Table 1**. But they are not the best in improving the yields of products and they are just at the average level of improving the yields of L-Glutamate, L-Lysine, L-Threonine, and Formate. Among these 83 enzymes, about 30 enzymes have better performance in improving the product yields with *E.coli* than the two reactions of NOG.

But the first two reactions, R07409 and R00790, have seemingly abnormal values in yields improved. We checked the reason and it may lie in the reversibility of the two reactions. We list all the fluxes through the added reactions when calculating the yields of the 10 chemicals, in **Supplementary Table S2**, following on the column of the yield of the corresponding chemicals. By comparing the direction of every added reaction, the flux through the reaction and the property of the reaction, we marked all the possible error of reversibility of added reactions in the column **Rev** with red color in **Supplementary Table S2**. The property of the reactions we mention here refer to CO_2_ and H_2_O as metabolites in the reactions, and it is not easy for CO_2_ and H_2_O to be decomposed in vivo. Although the reaction directions of these added reactions were checked in Ref [3], the reversibility quality of marked reaction may result in wrong calculations for improving yields.

At the same time, considering the enzyme activity, when we calculated the growth rate and the theoretical yields of above 10 chemicals with *E.coli* by adding one reaction every time, we constrained the up/low flux bounds to not more than 100 in abstract value. We list all the yields of the 10 chemicals in this condition in **Supplementary Table S3**. Of course, they were lower than the corresponding yield value if without constraint. We also calculated the smallest balanced pathway for every chemical by the tool FIND_tfSBP [8] and provided the pathways of Excel table in **Supplementary Material**. At the same time, we drew the pathways for every chemical by an improved Paint4Net [9] in **Supplementary Pictures**, where the highly correlated nodes, such as the co-factors of ATP, ADP, NADP, NADPH, H, H2O and Pi, are splited with differrent labels, in order to reduce the network complexity. Metabolic flux is proportional to line thickness.

Global benefit enzymes which can increase the theoretical yields of many chemical products for cell factories are what we want. NOG is one of them. In this study, by computation method, we find many other global benefit enzymes, which can increase the theoretical yields of some chemical products in *E.coli*. Especially, some of these global benefit enzymes are better than NOG in many products from the comparison result above. For the number of these global benefit enzymes we find is large, it is not possible to give testification with experiments. Computation method has been used as testification method widely. We think these global benefit enzymes we find will be potentially effective.

### The example of GNO (Glycerol:NADP+ oxidoreductase)

We add NOG (R00761) into the iAF_1260 model and calculate, one by one, the maximum yields of all the 80 products which *E.coli* may produce actually. Then we add GNO into the iAF_1260 model and calculate the maximum yields of all the 80 products with the same method. The maximum yields of *E.coli* for wild-type were listed as a comparison. From the yields of 80 products, we find that all of them are higher than wild-type when adding with GNO, so GNO is also a global benefit enzyme. We classify the result into three groups, i.e. “NOG is better”, “GNO is better” and “No significant difference between NOG and GNO”. “NOG is better” of 30 products was shown in **Table 2**, “GNO is better” of 43 products was shown in **Table 3**, and “No significant difference between NOG and GNO” of 7 products was shown in **Table 4**. GNO gets 43 products better than NOG, while NOG gets 30 better than GNO, so GNO has better performance in the product production of *E.coli*.

**Table 2.**
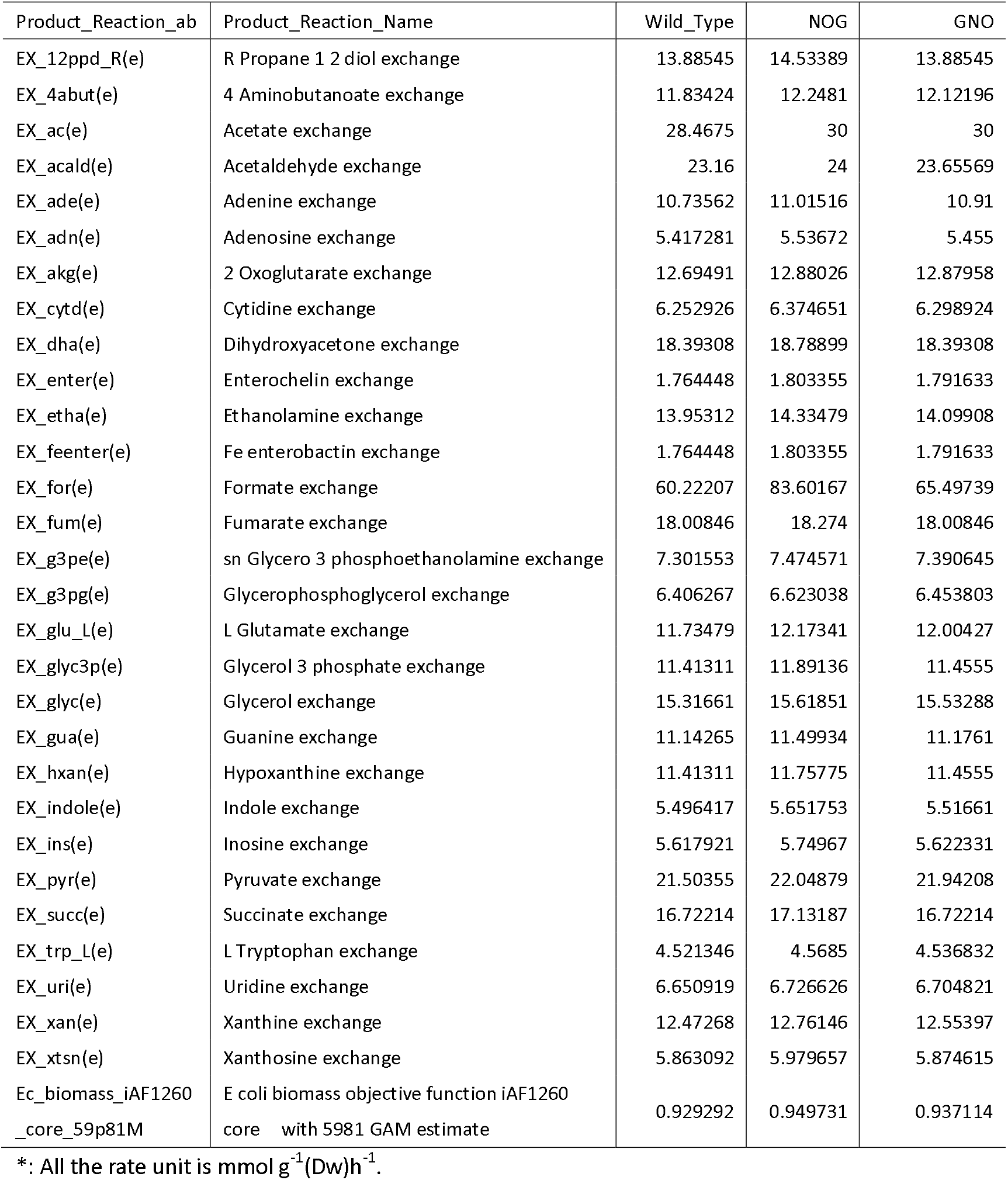
NOG is better (30 products)

**Table 3.**
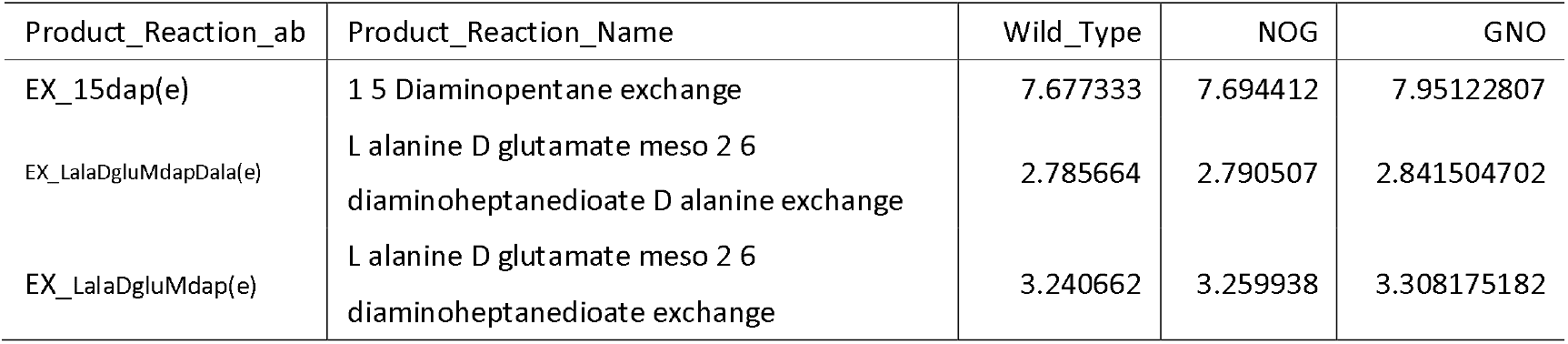
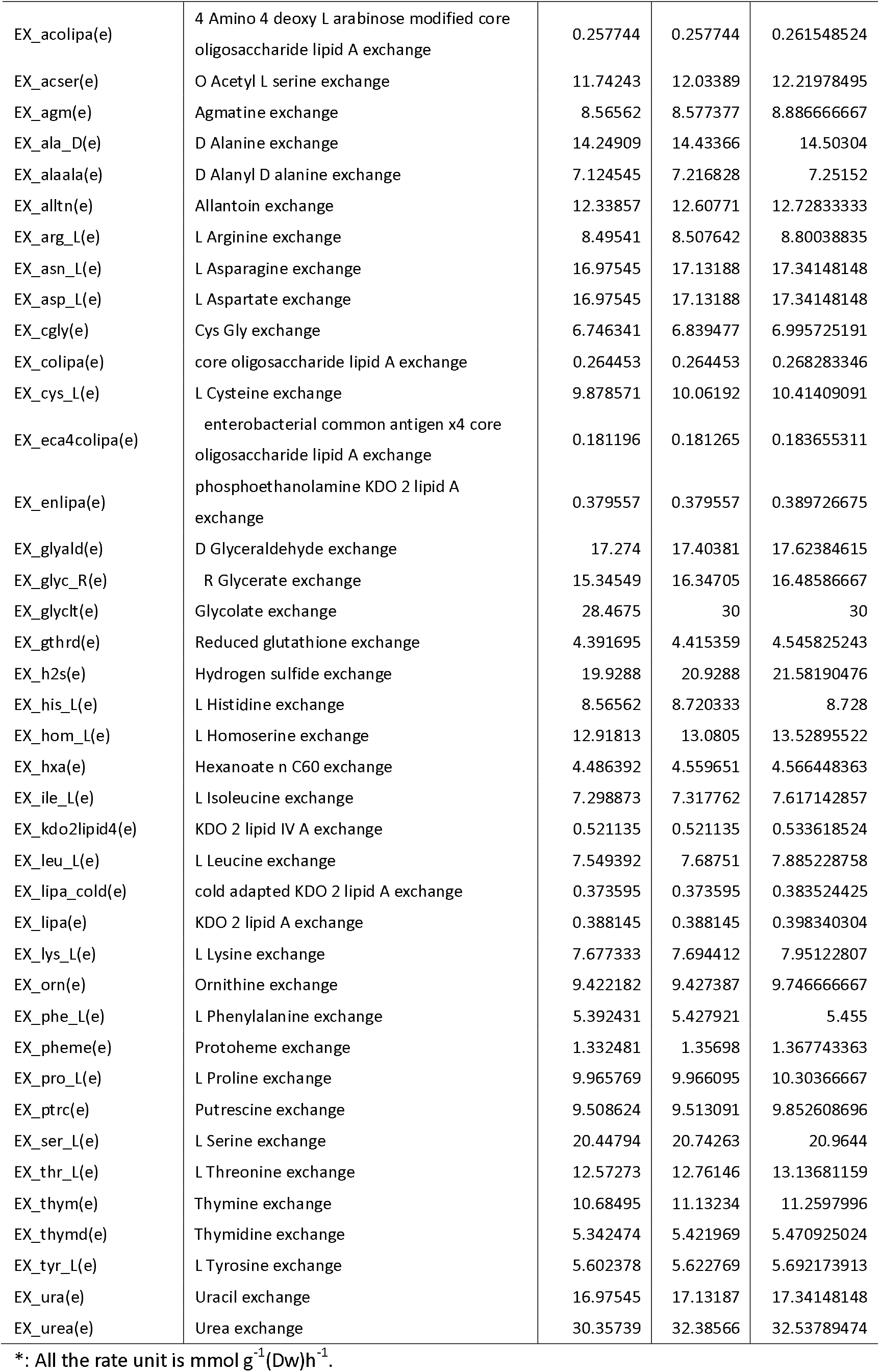
GNO is better (43 products)

**Table 4.**
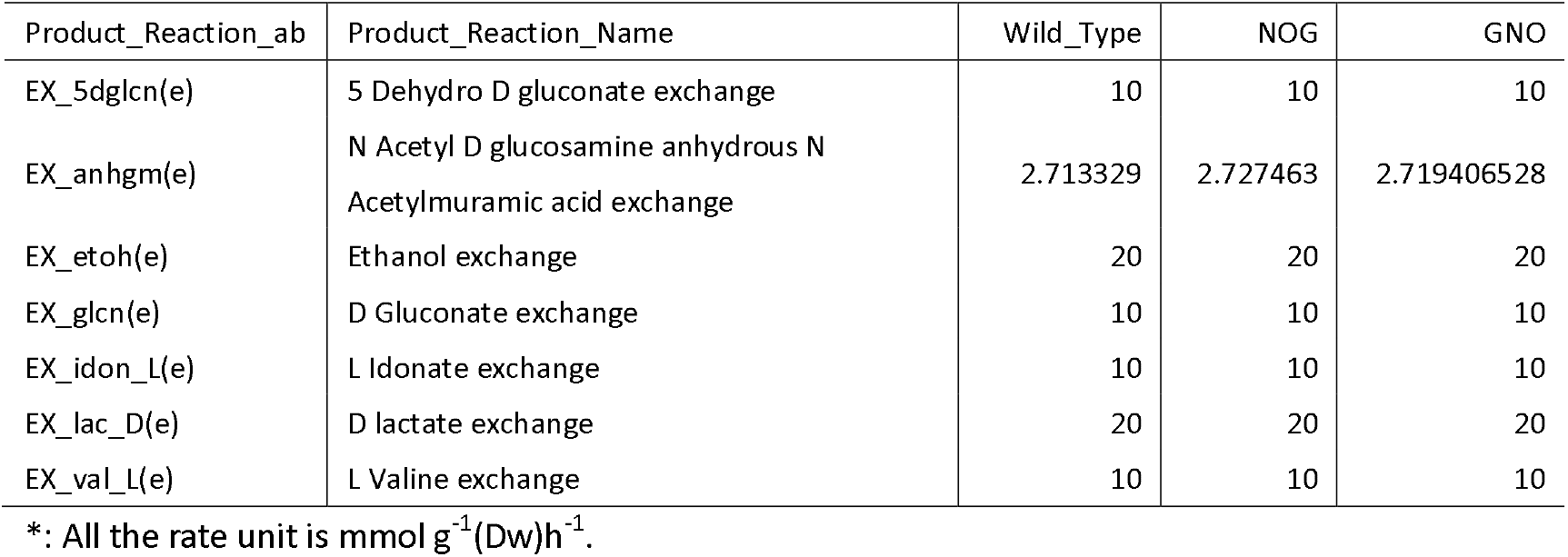
No significant difference between NOG and GNO (7 products)

Global benefit enzymes which can increase the theoretical yields of many chemical products for cell factories are our hope. NOG is one of them. In this study, we find another global benefit enzyme, GNO, which can increase the theoretical yields of 80 chemical products in *E.coli*. Especially, GNO is better than NOG in many products from the comparison result above. GNO is an enzyme related to NADPH production. NADPH is an important cofactor in the construction of cell factory. The cofactor optimization will increase the supply of reducing power, so increase the theoretical yields.

## Supporting information

supplementary pictures

## Acknowledgments

Support for this work was provided by “National Natural Science Foundation of China (31370829)’’, “Tianjin Research Program of Application Foundation and Advanced Technology (15JCYBJC23600)”. The funders had no role in study design, data collection, and analysis, decision to publish, or preparation of the manuscript.

## Additional Information

The authors declare no competing financial interests.

## Author contributions

Conceived and designed the experiments: ZX. Performed the experiments: ZX. Analyzed the data: ZX QW DZ. Contributed reagents/materials/analysis tools: QW DZ. Wrote the paper: ZX.

