## Supplementary figures and images for "The Enzymes that beyond Non-Oxidative Glycolysis"

### R00352_Ace.png

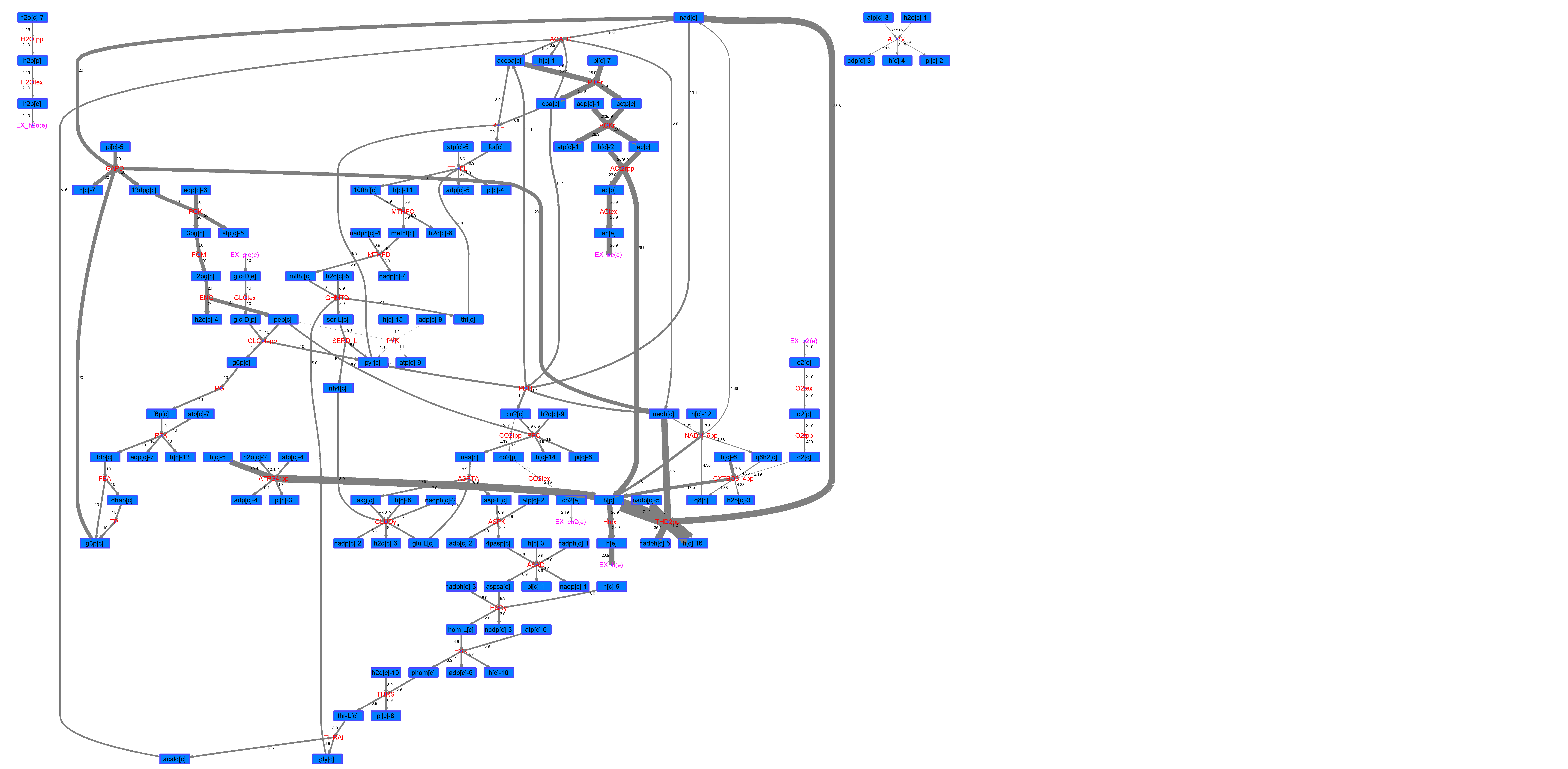

### R00352_For.png

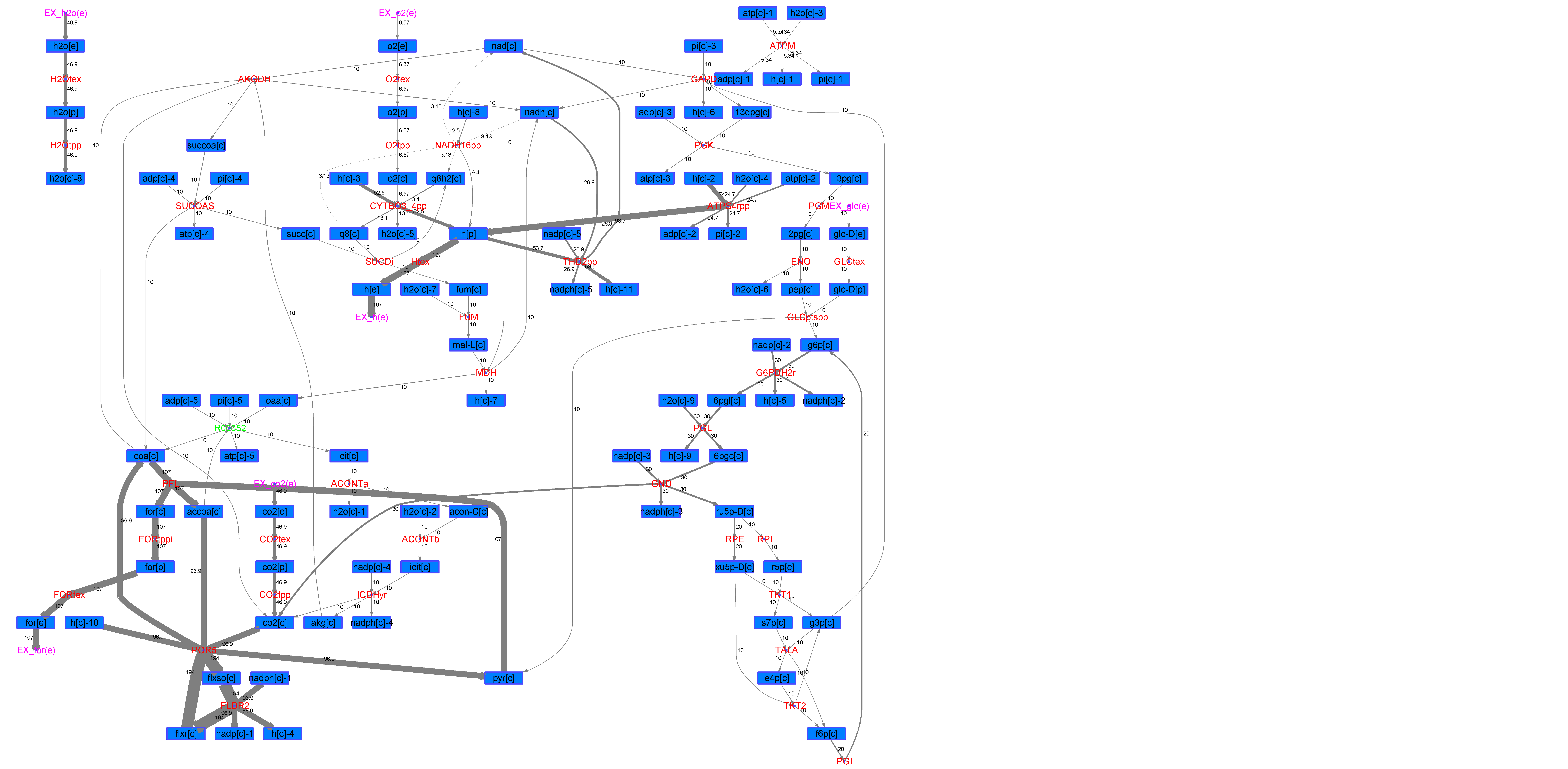

### R00352_Glu.png

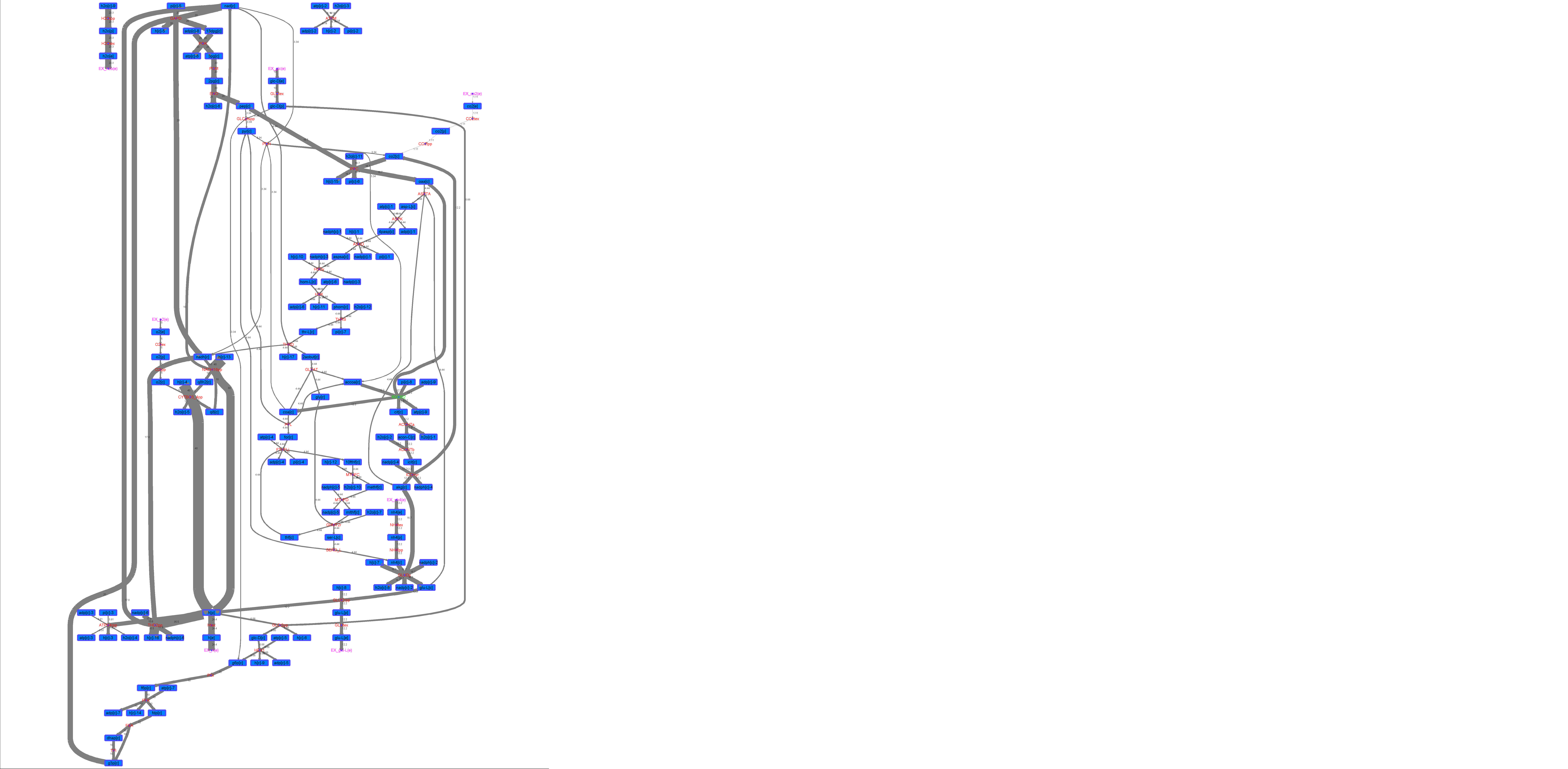

### R00352_L-ala.png

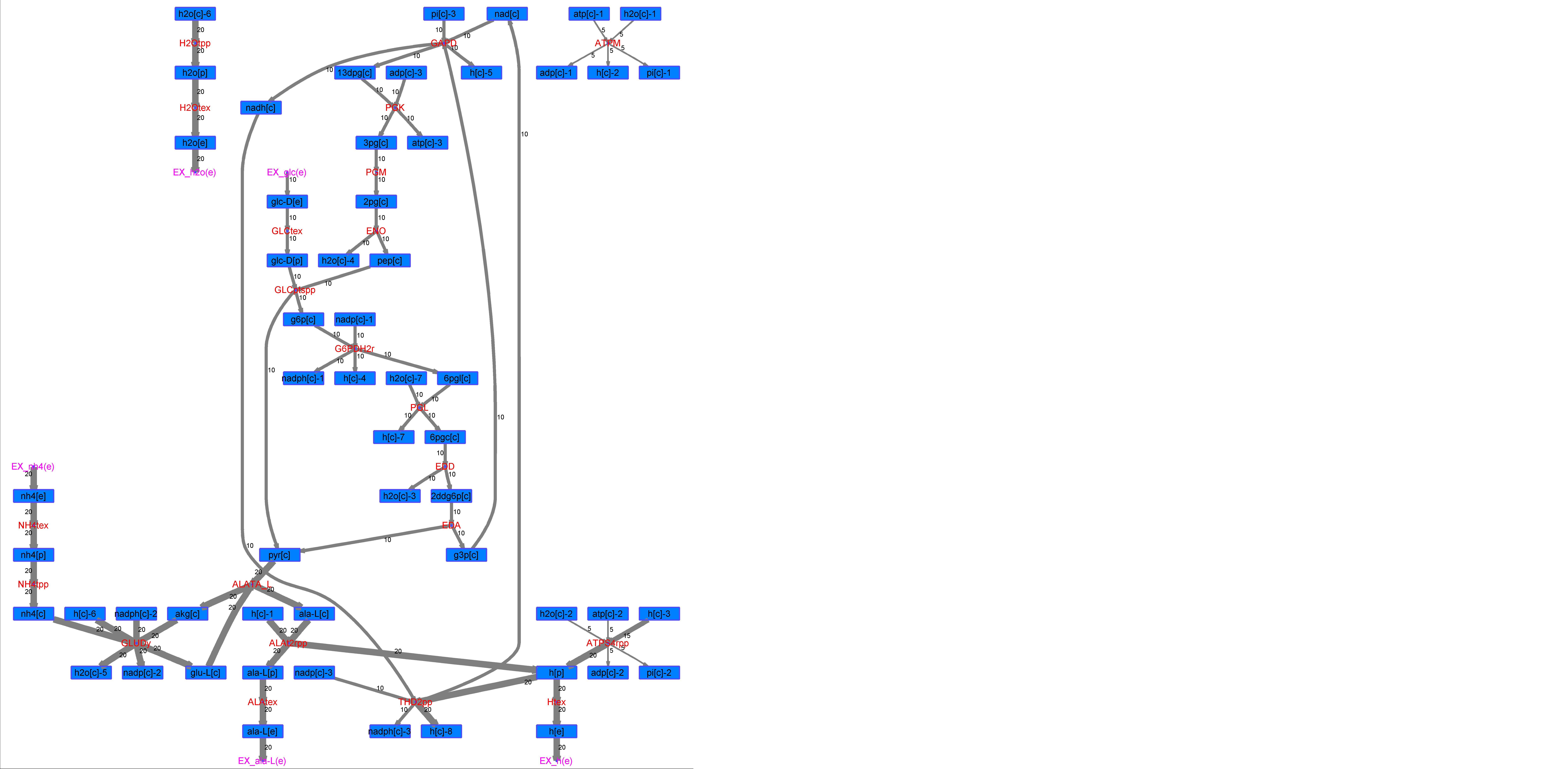

### R00352_L-Mal.png

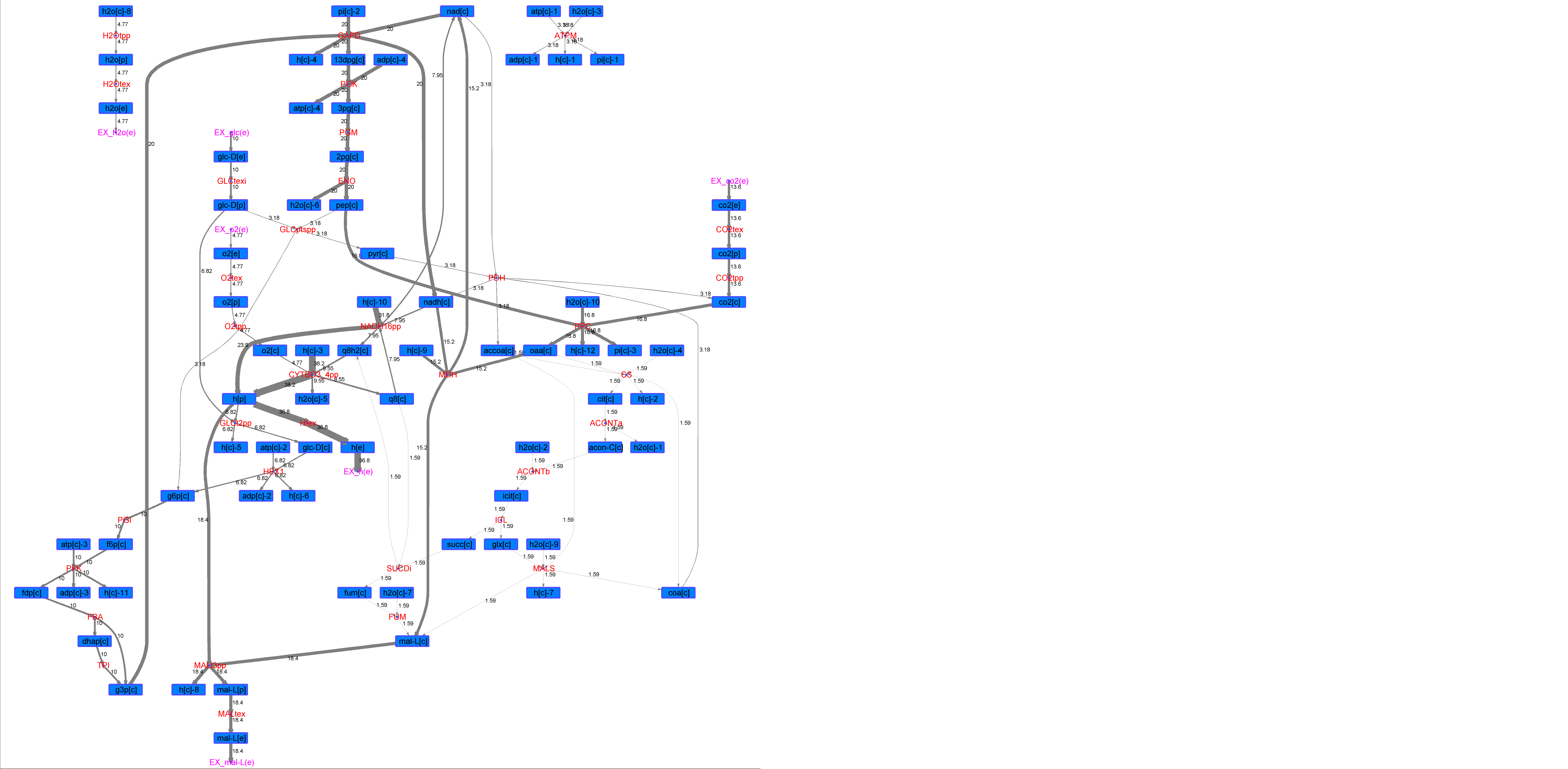

### R00352_L-Phe.png

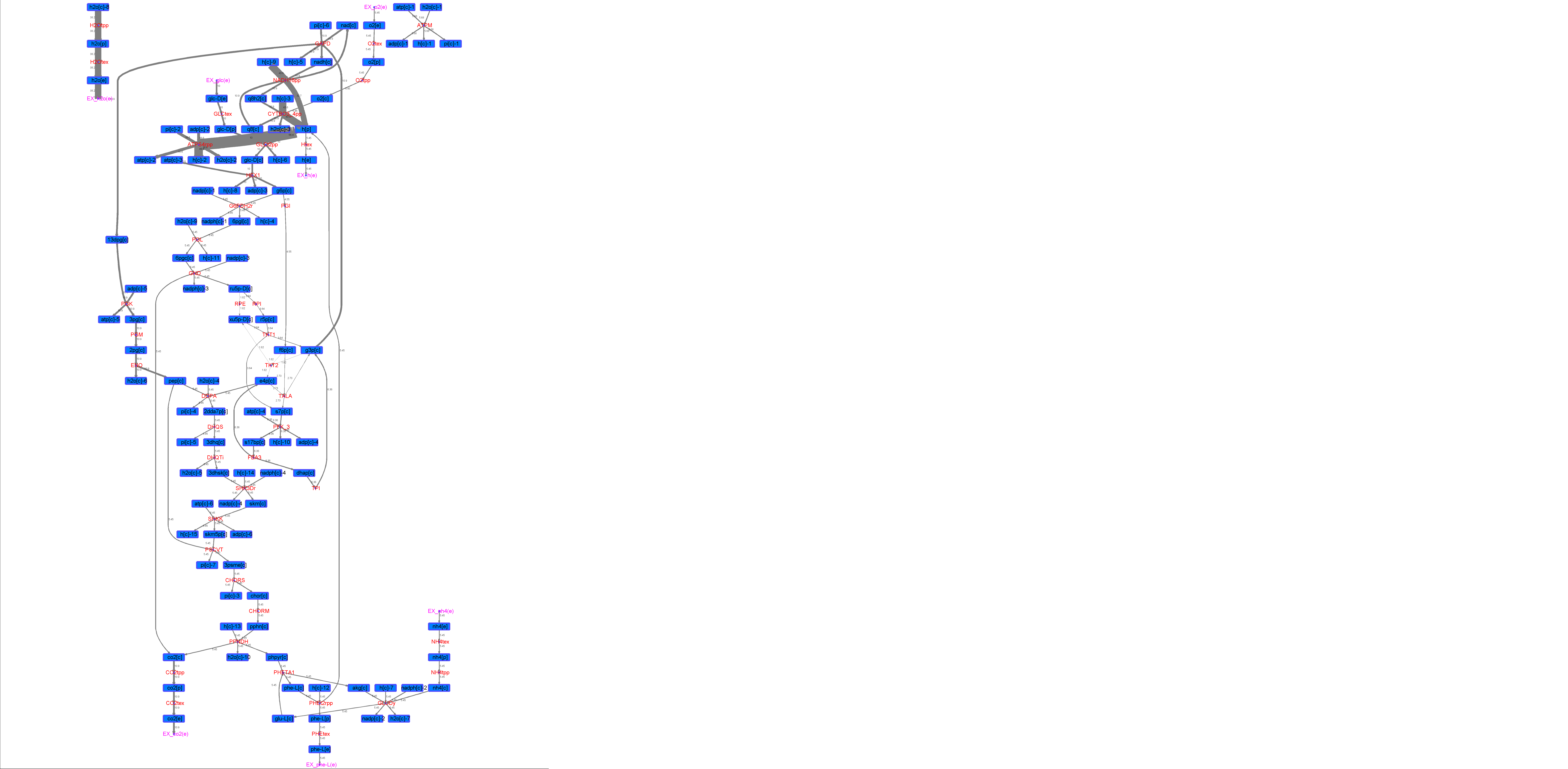

### R00352_L-Try.png

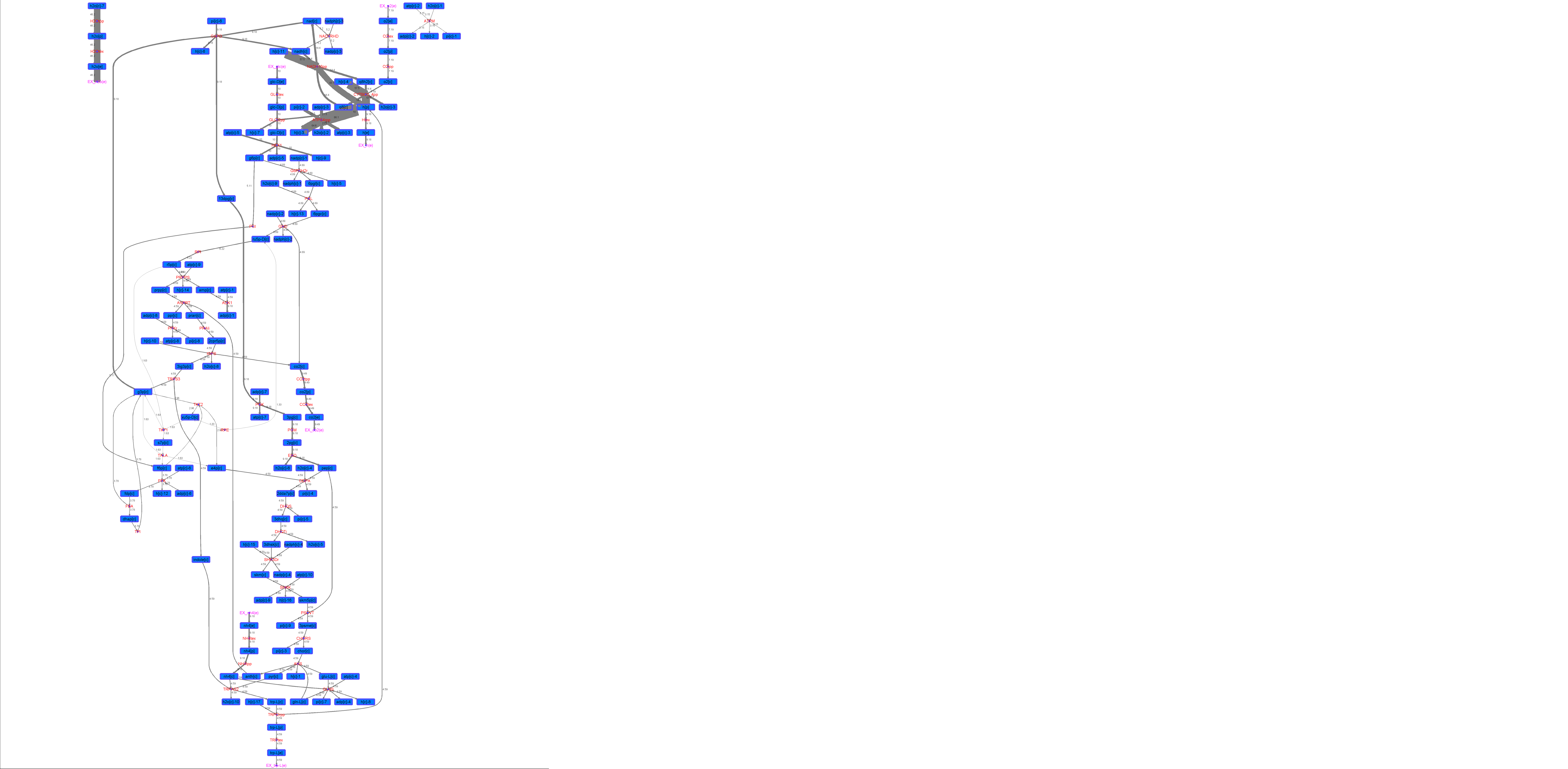

### R00352_Lys.png

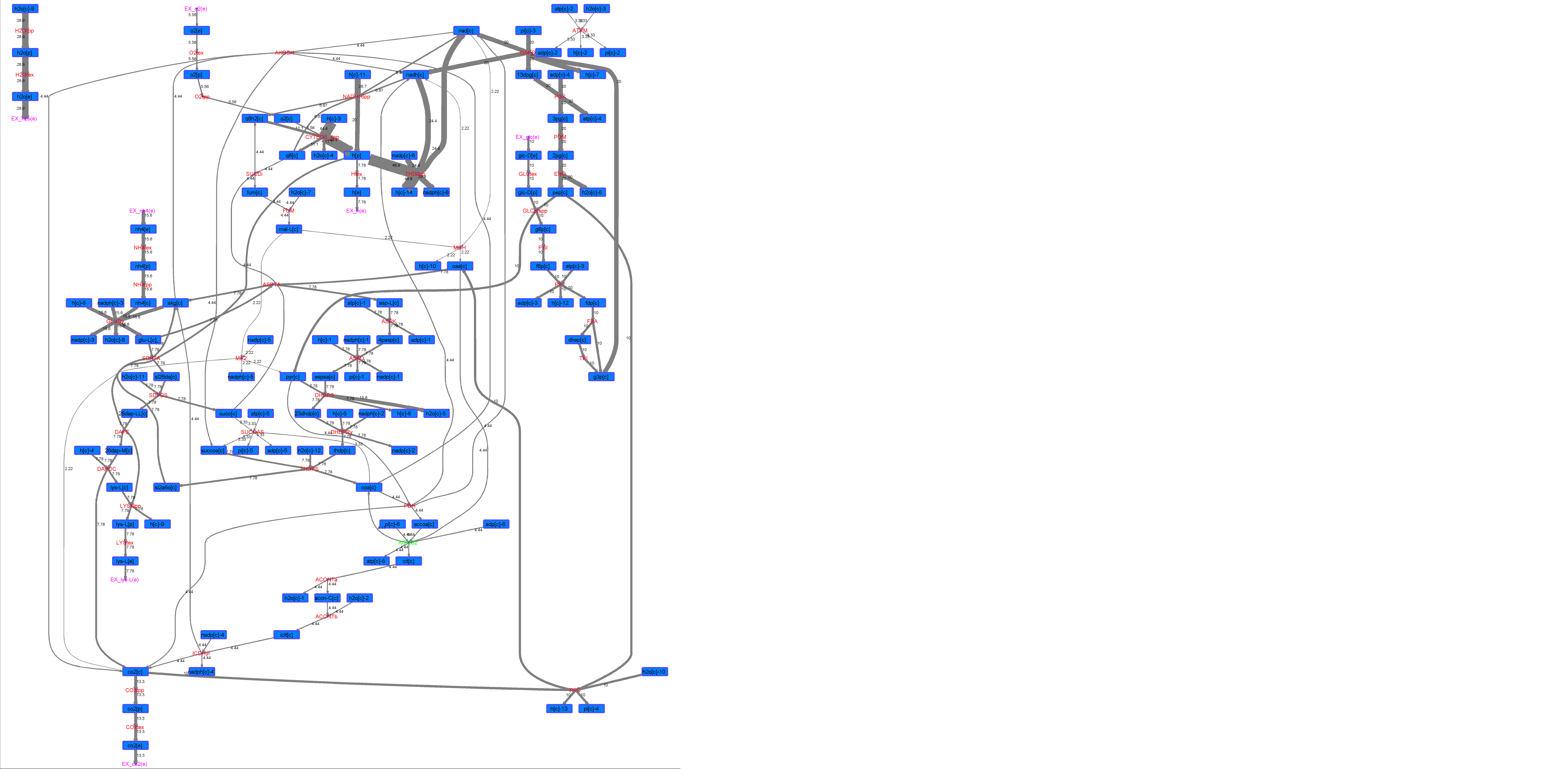

### R00352_Suc.png

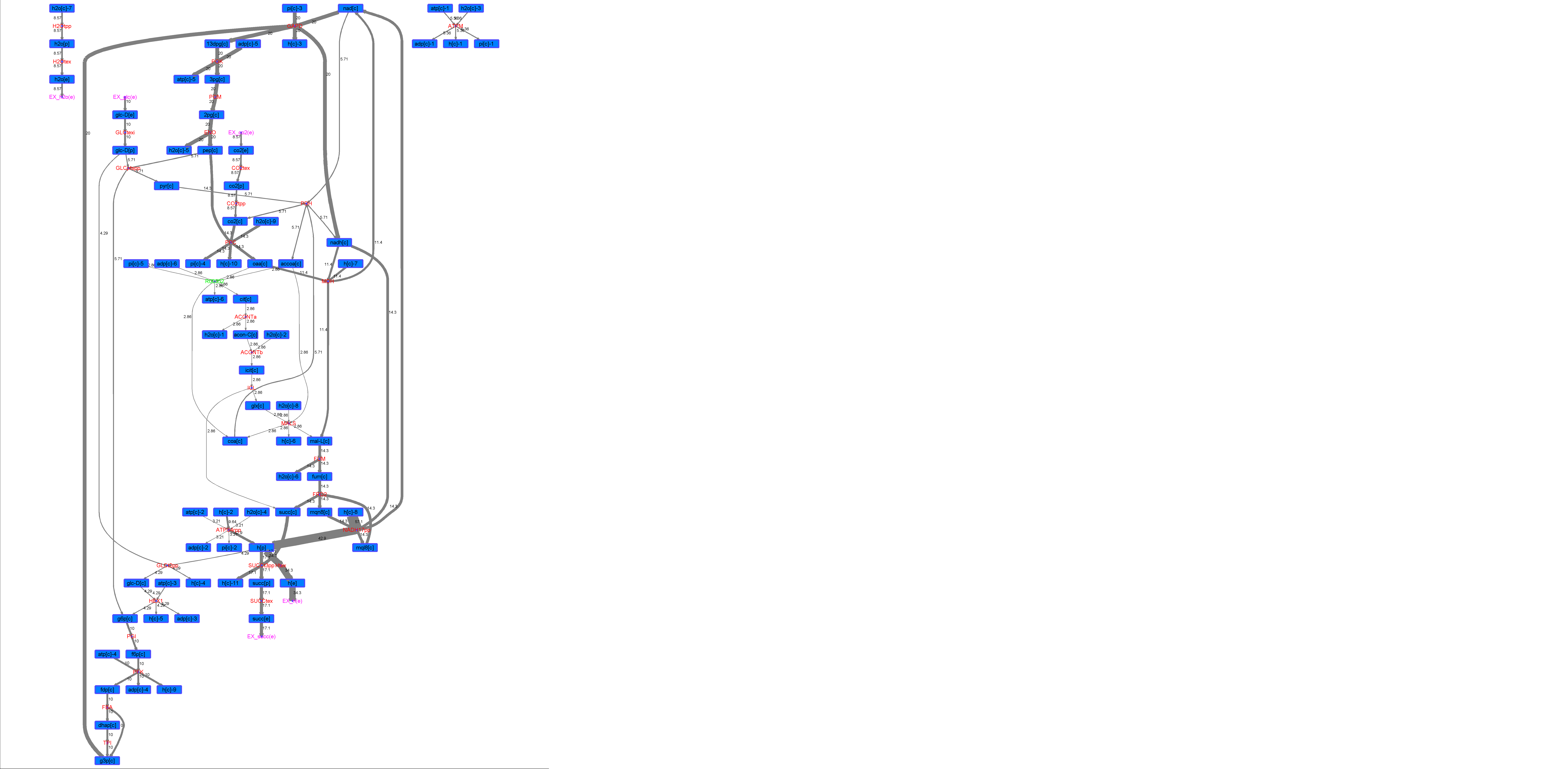

### R00352_Thr.png

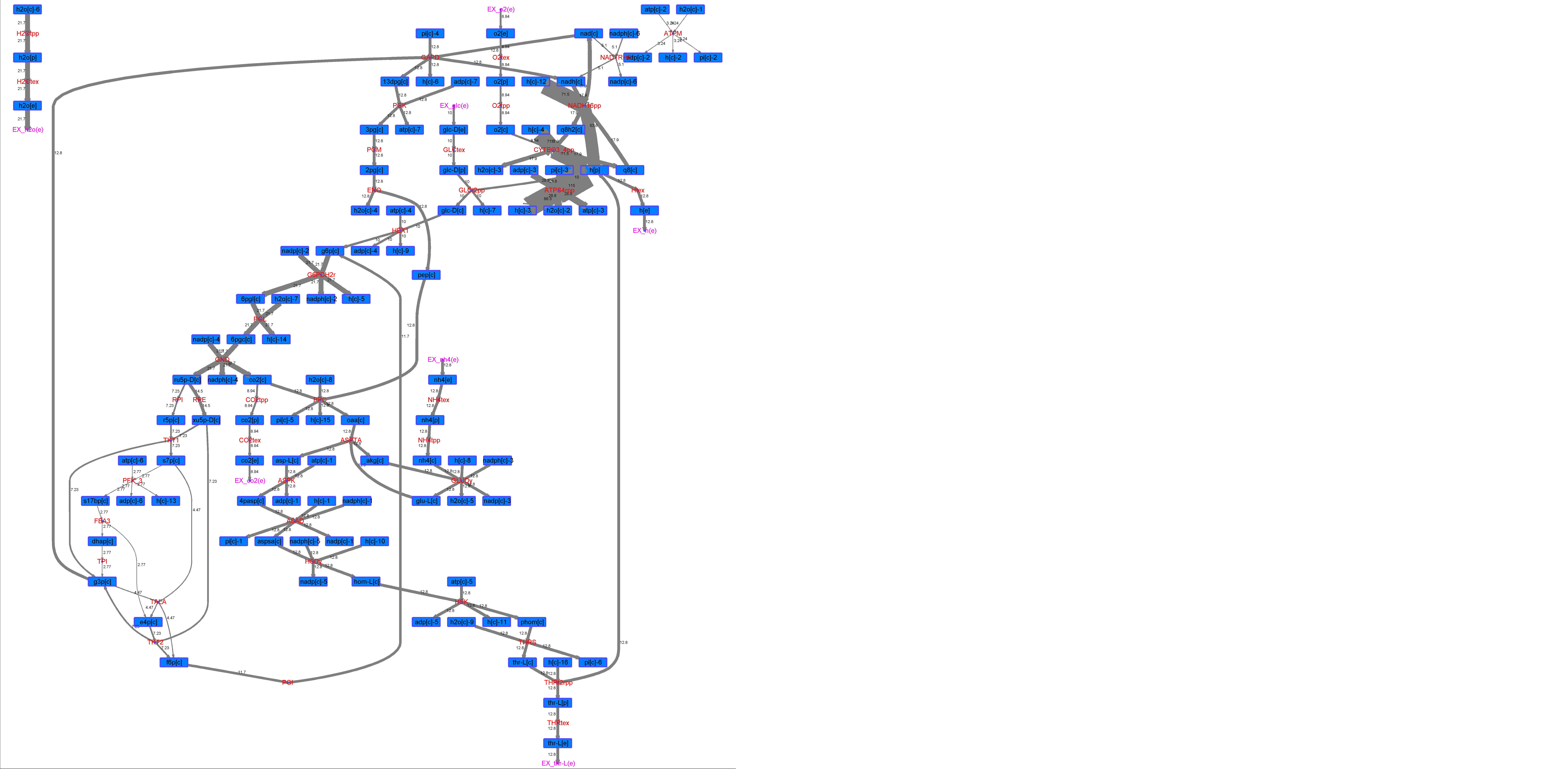

### R00396_Ace.png

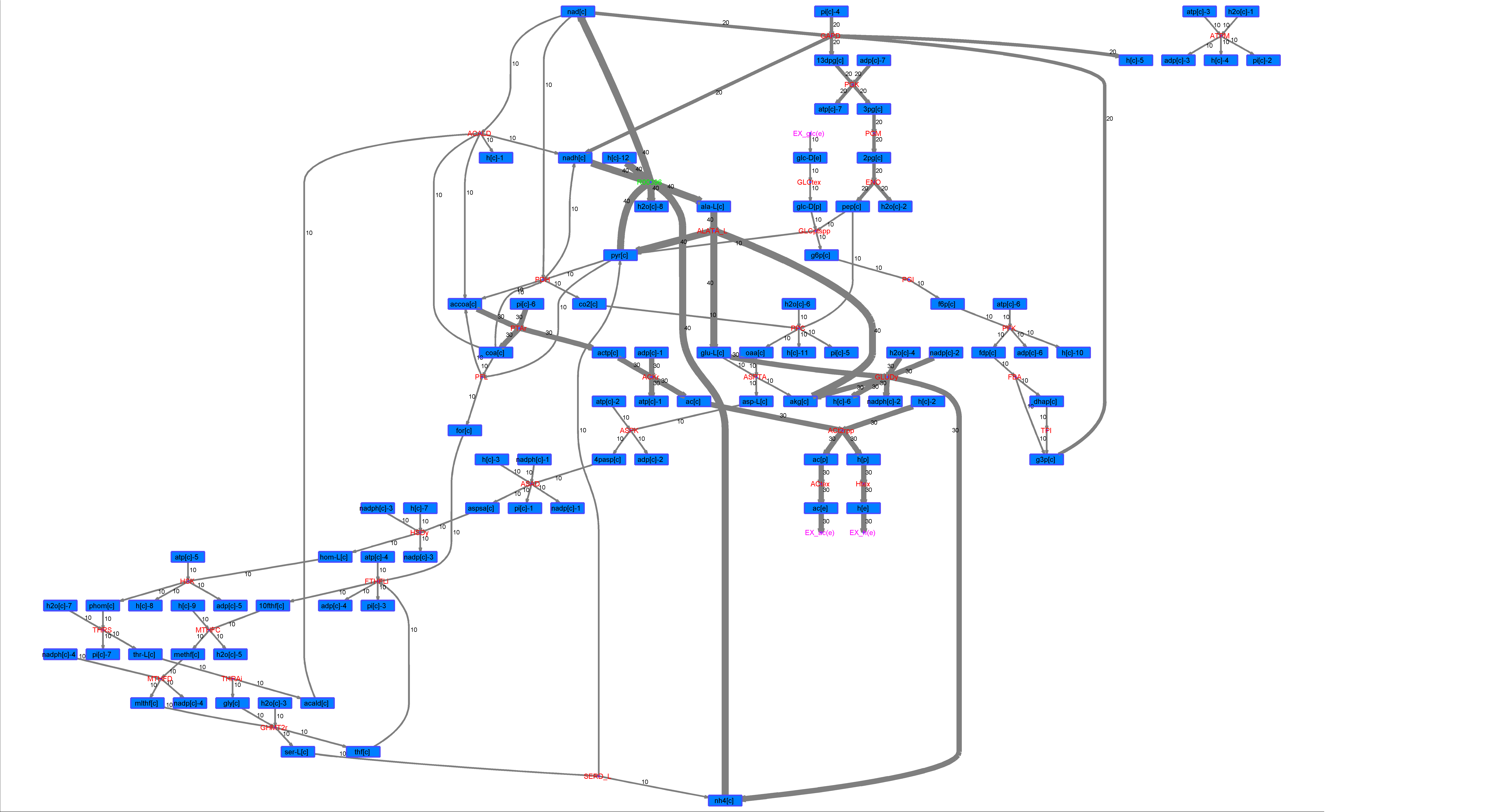

### R00396_For.png

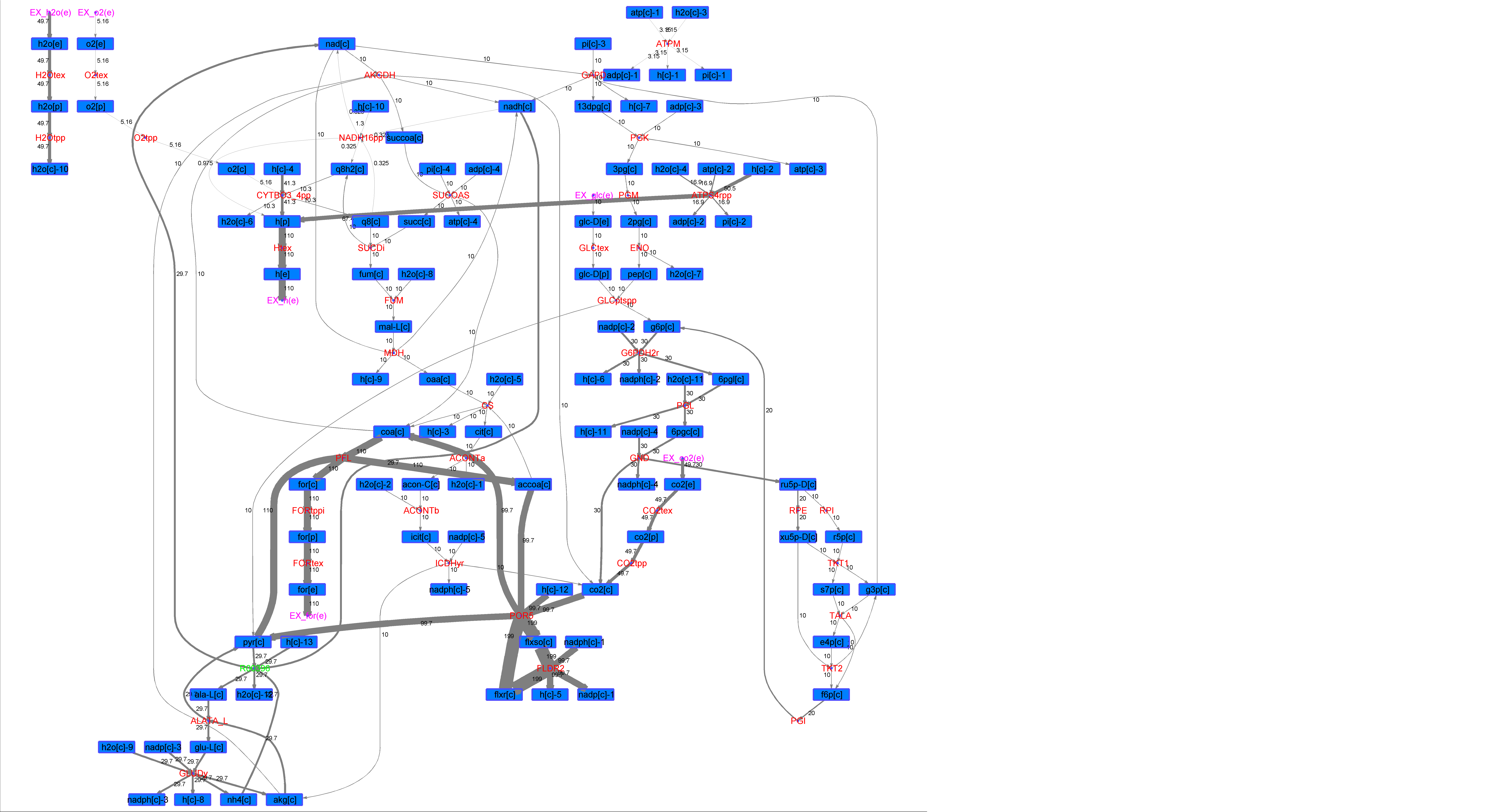

### R00396_Glu.png

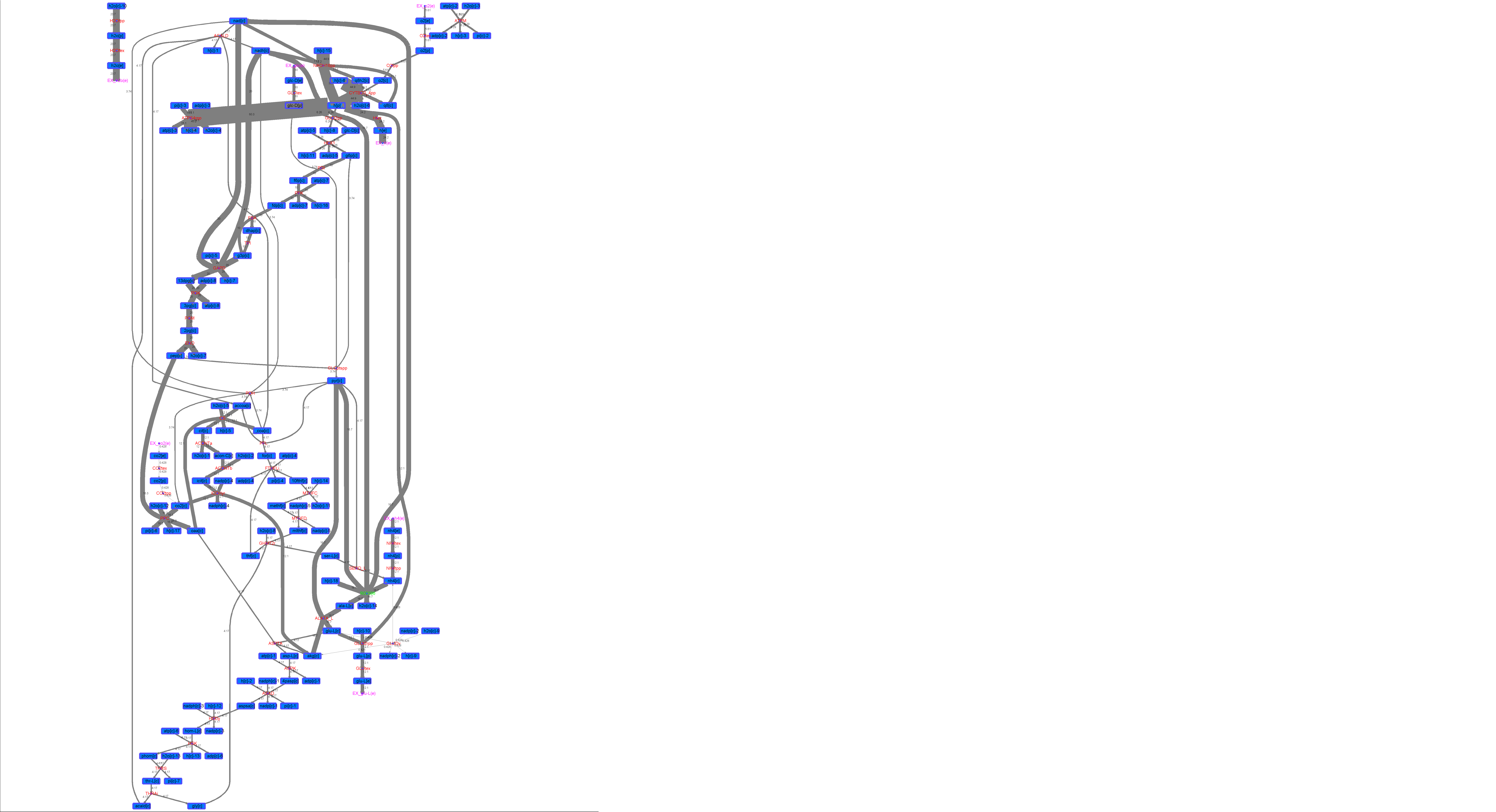

### R00396_L-ala.png

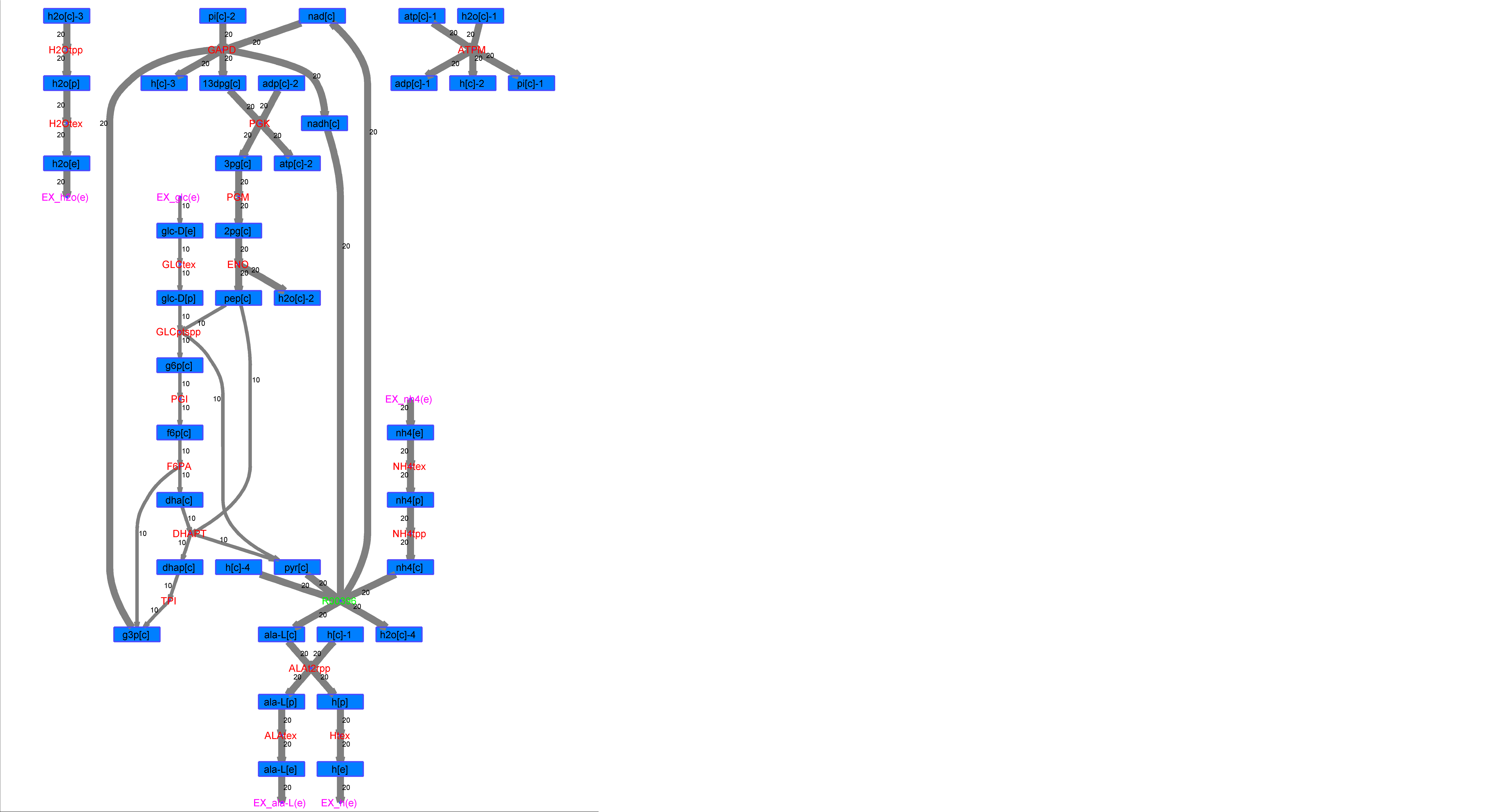

### R00396_L-Mal.png

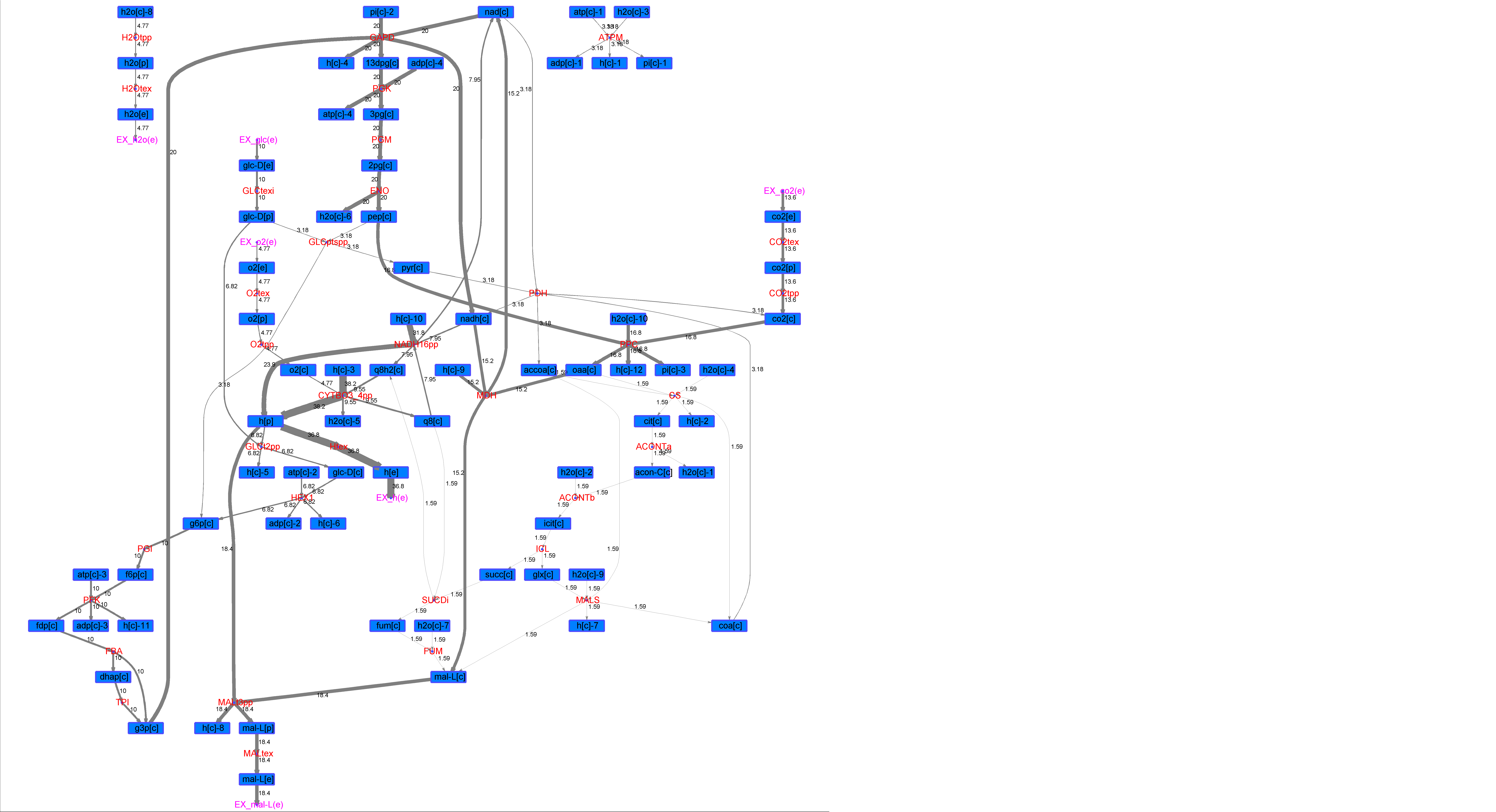

### R00396_L-Phe.png

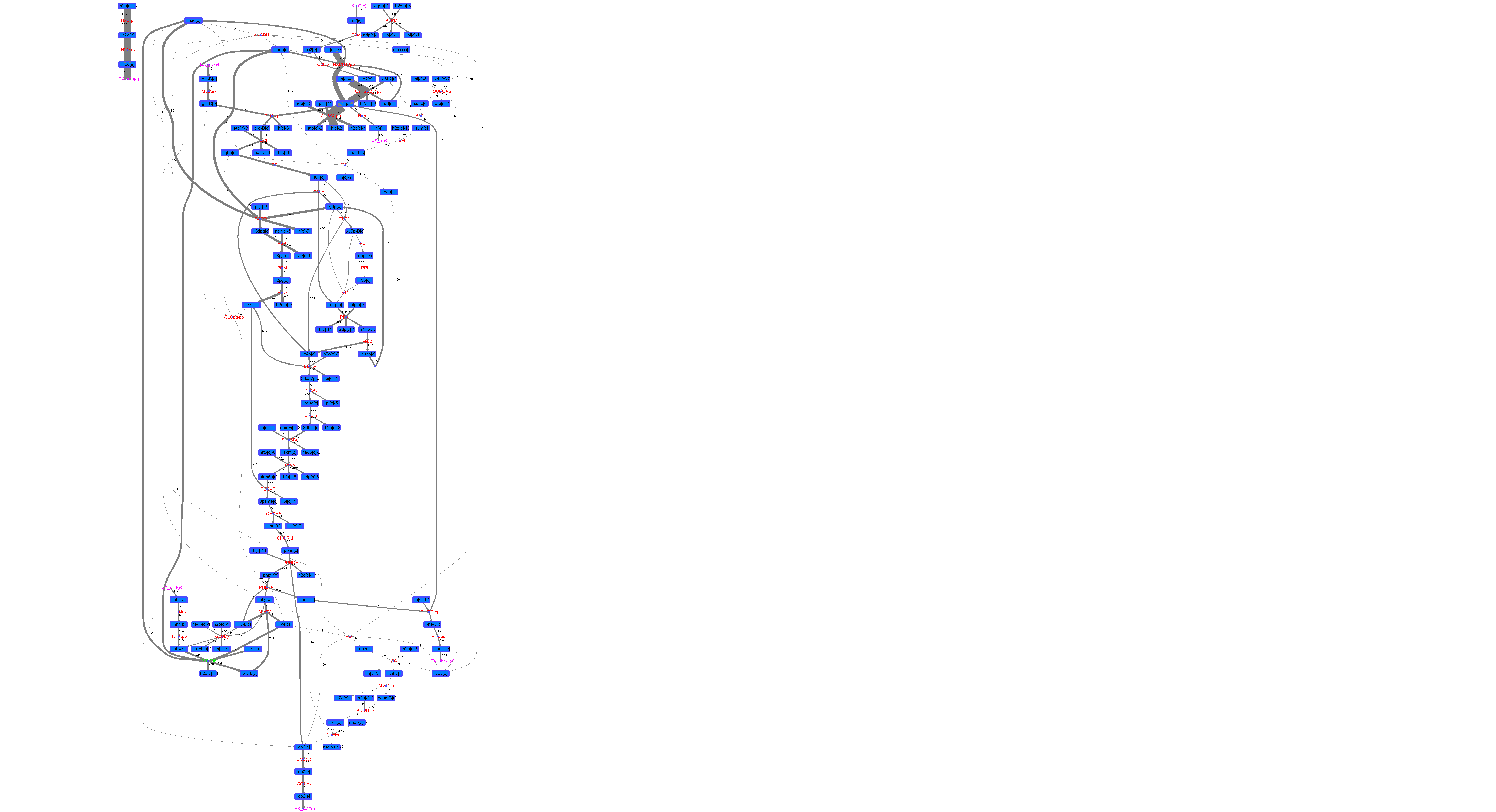

### R00396_L-Try.png

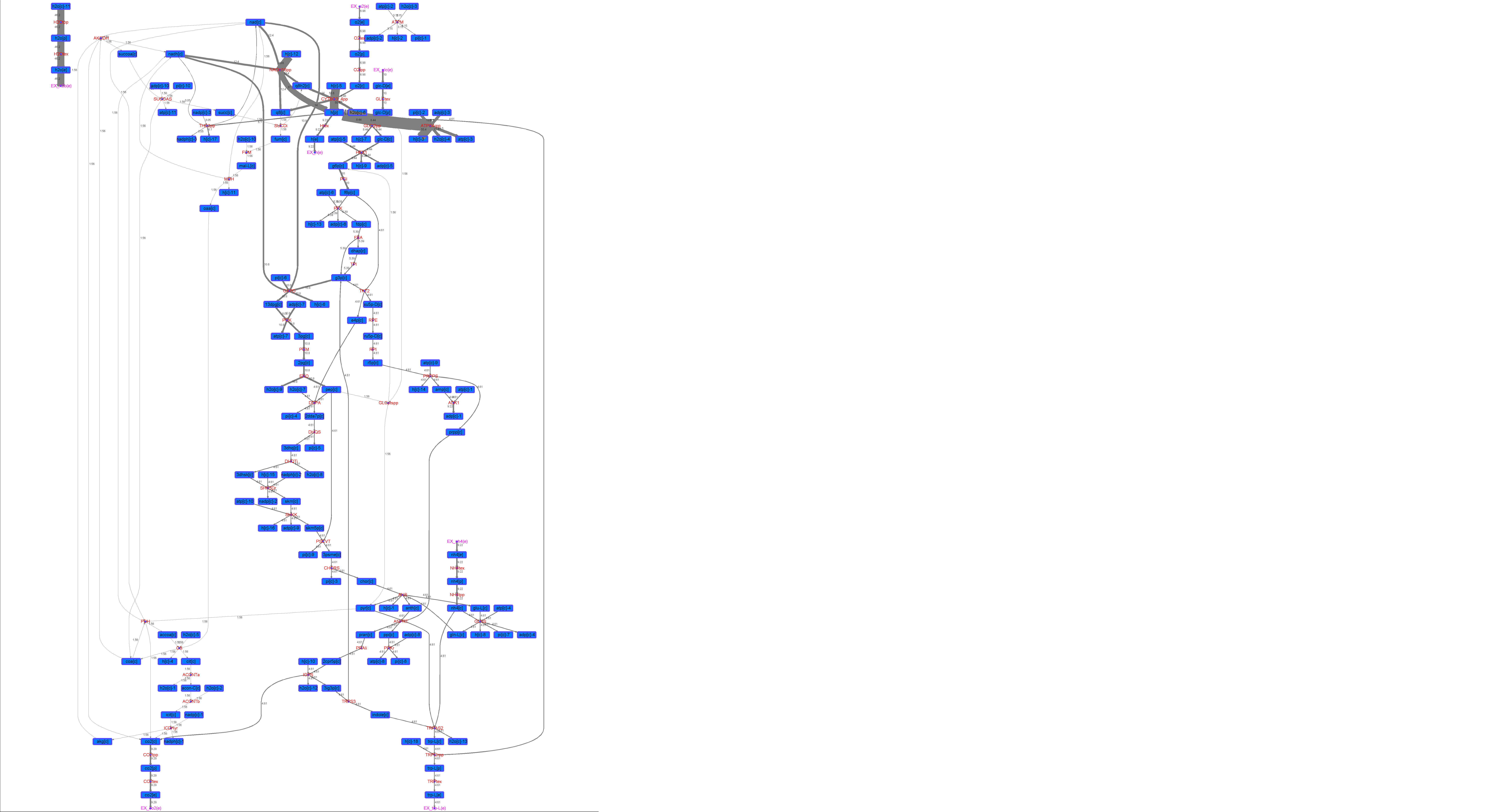

### R00396_Lys.png

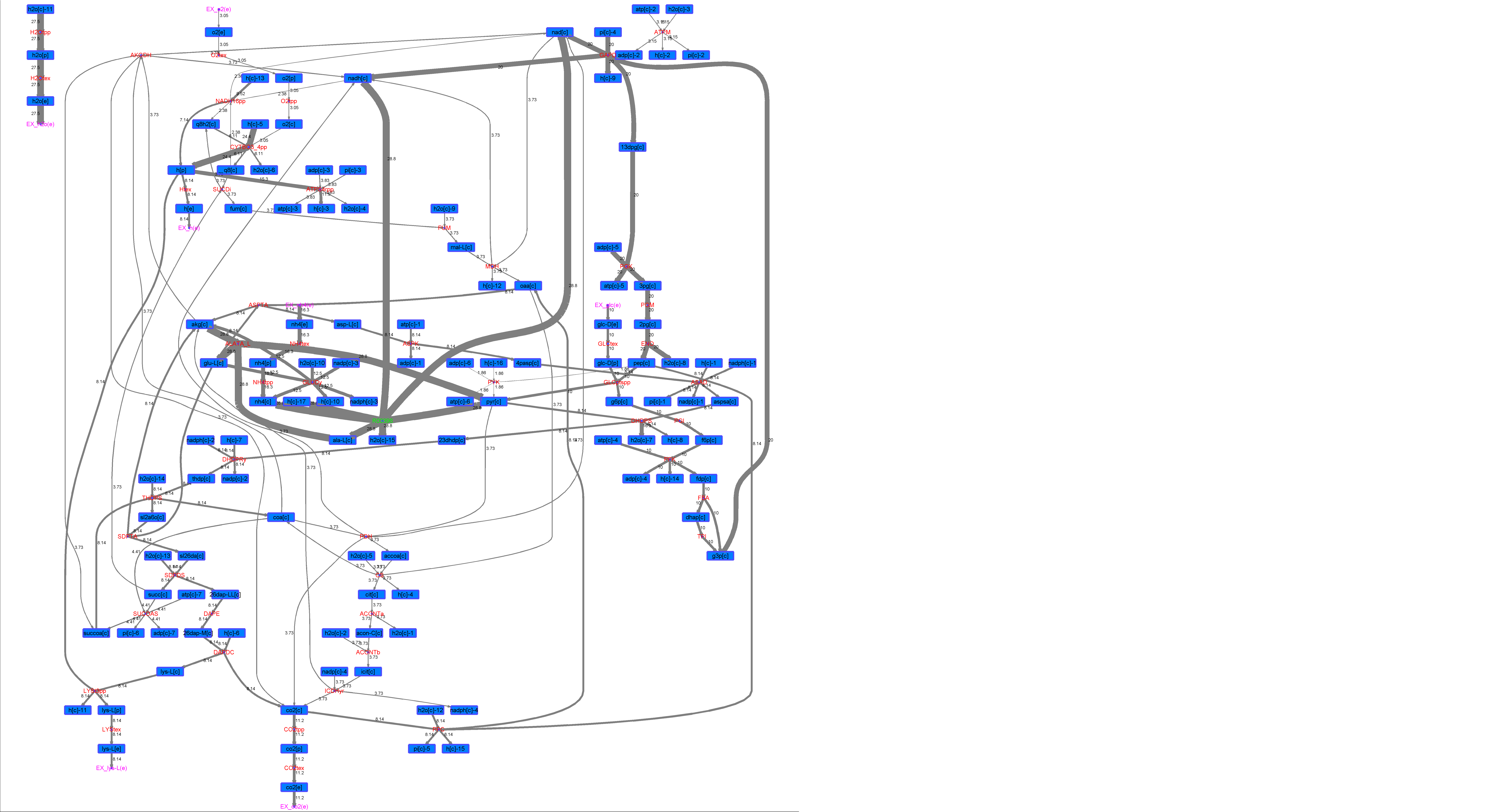

### R00396_Suc.png

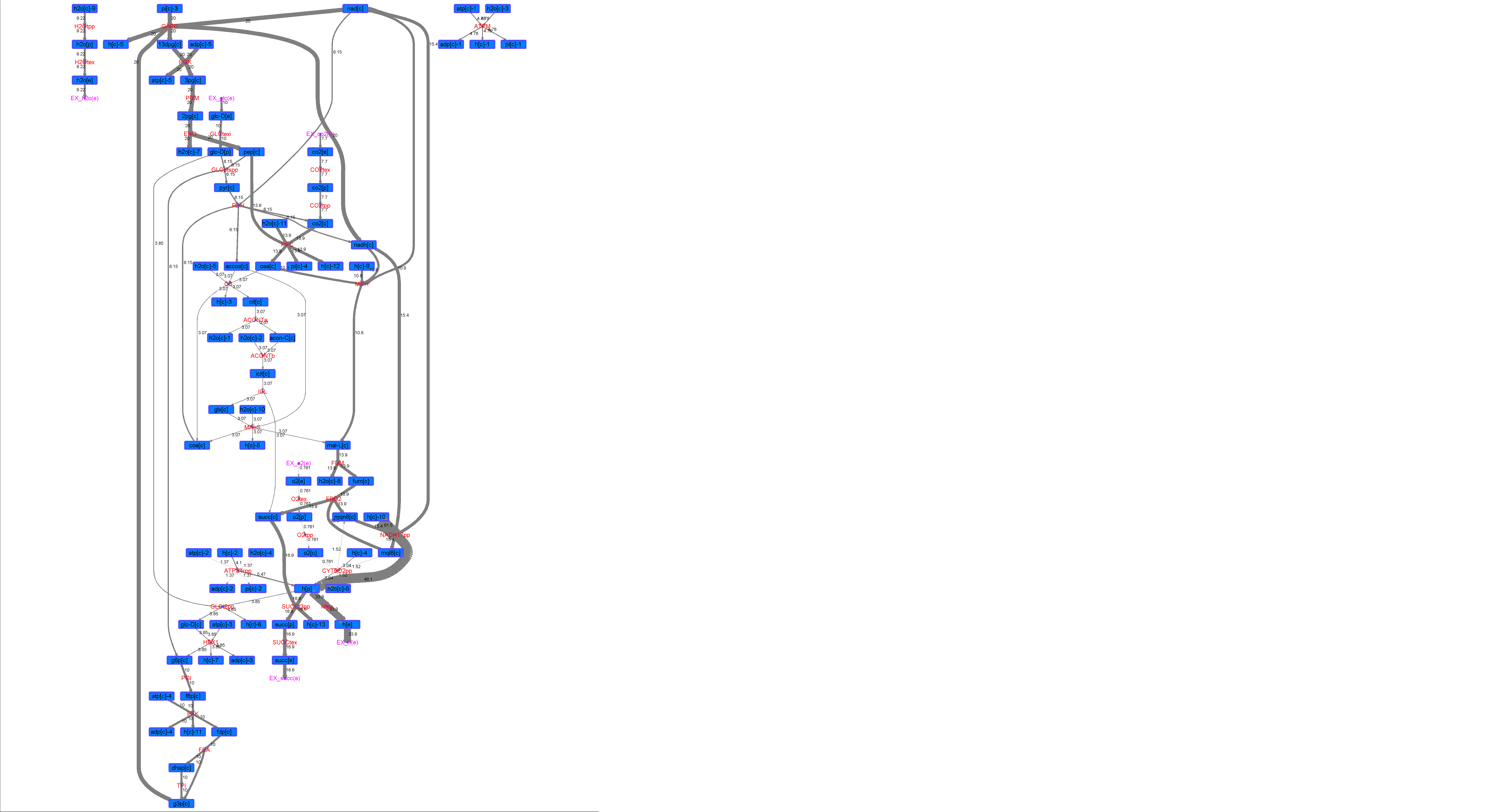

### R00396_Thr.png

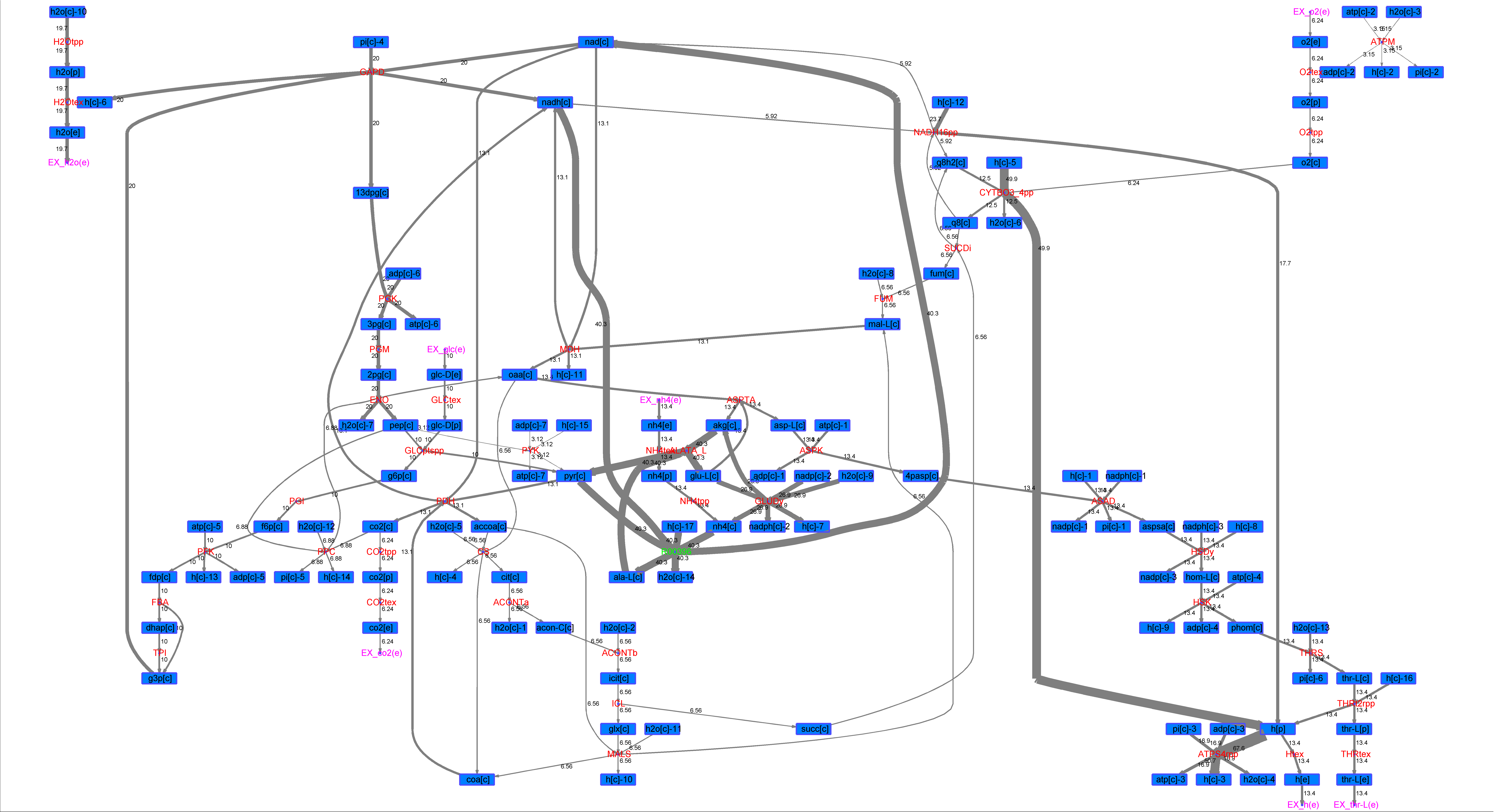

### R00397_Ace.png

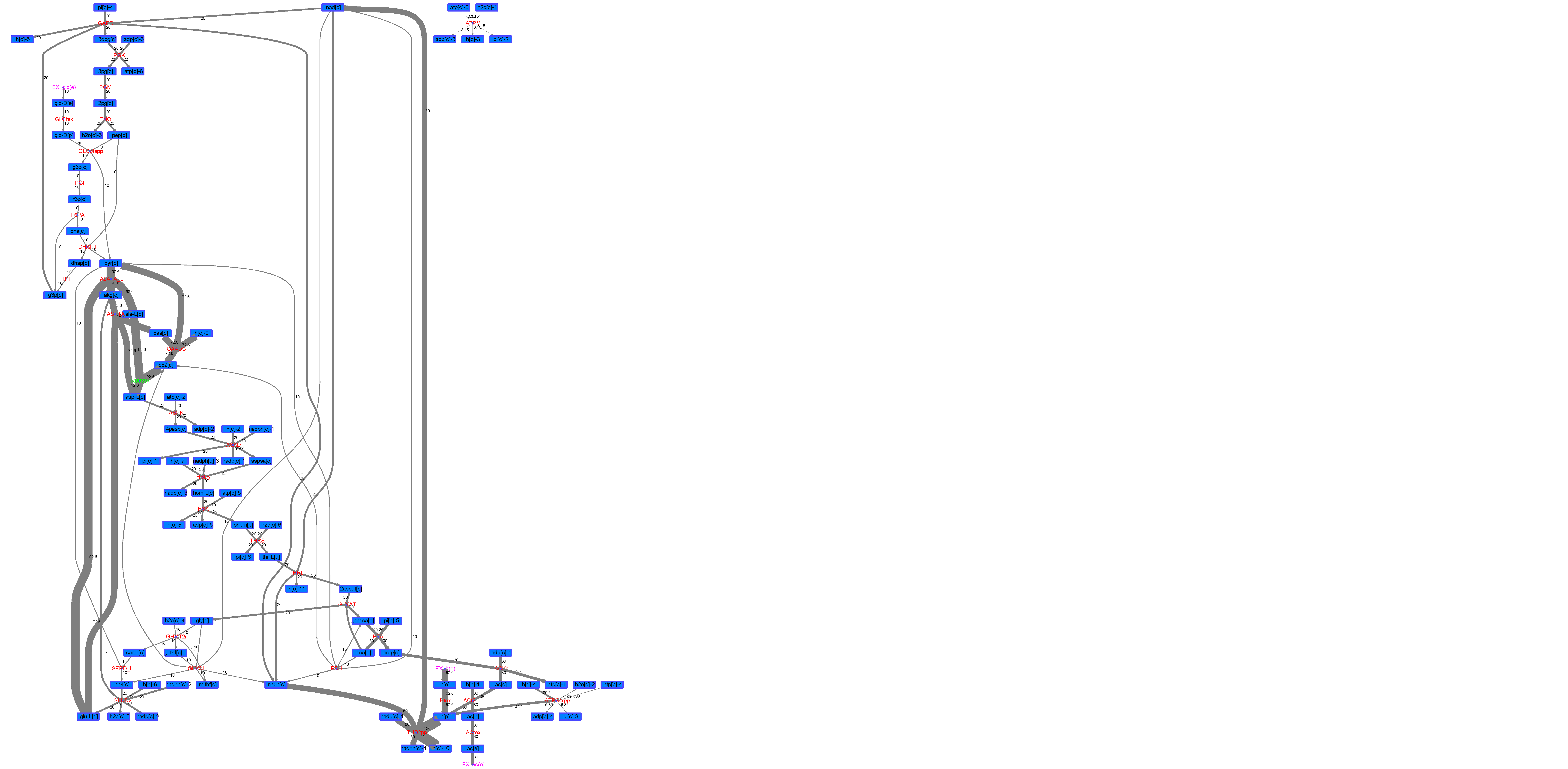

### R00397_For.png

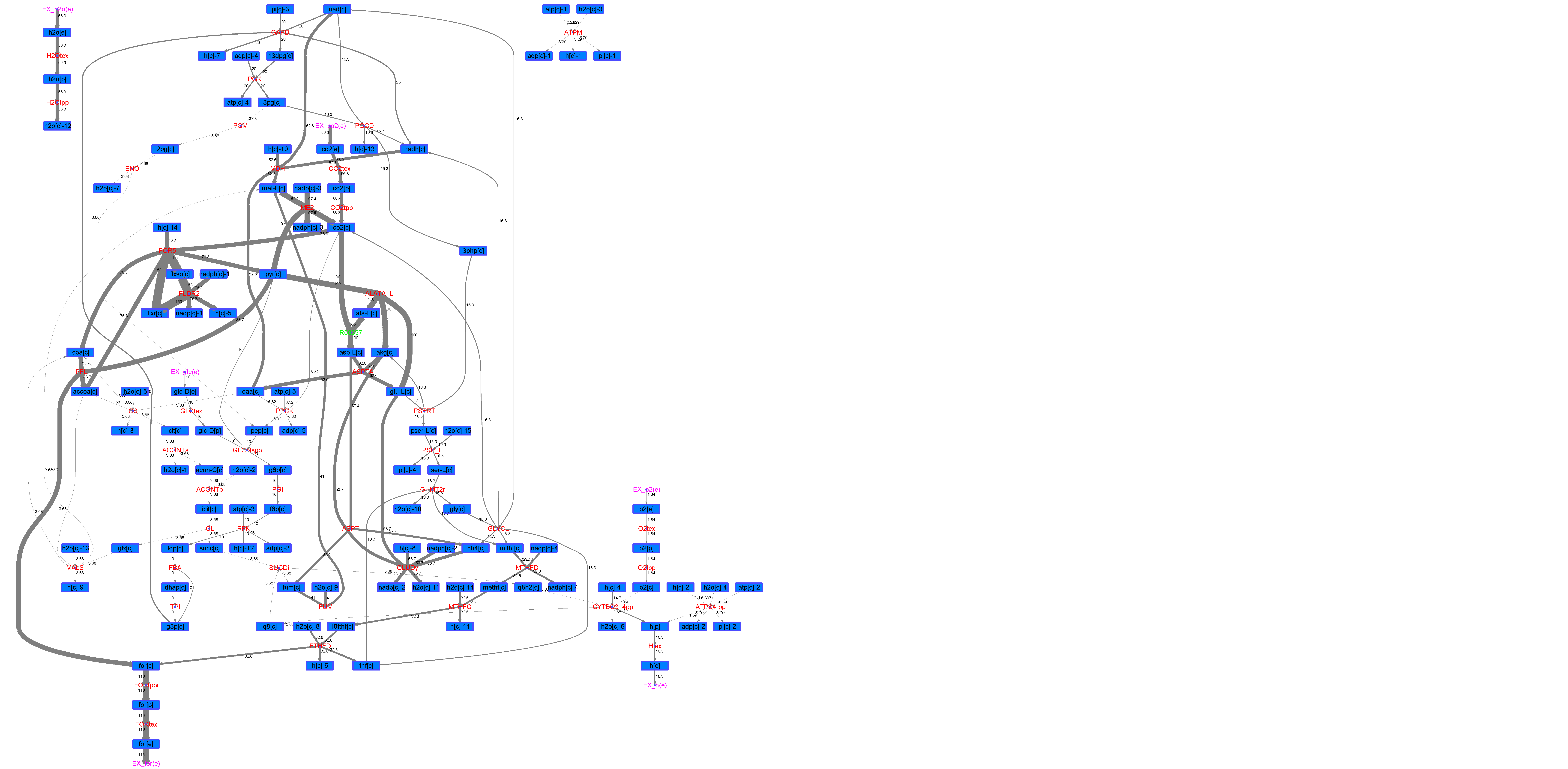

### R00397_Glu.png

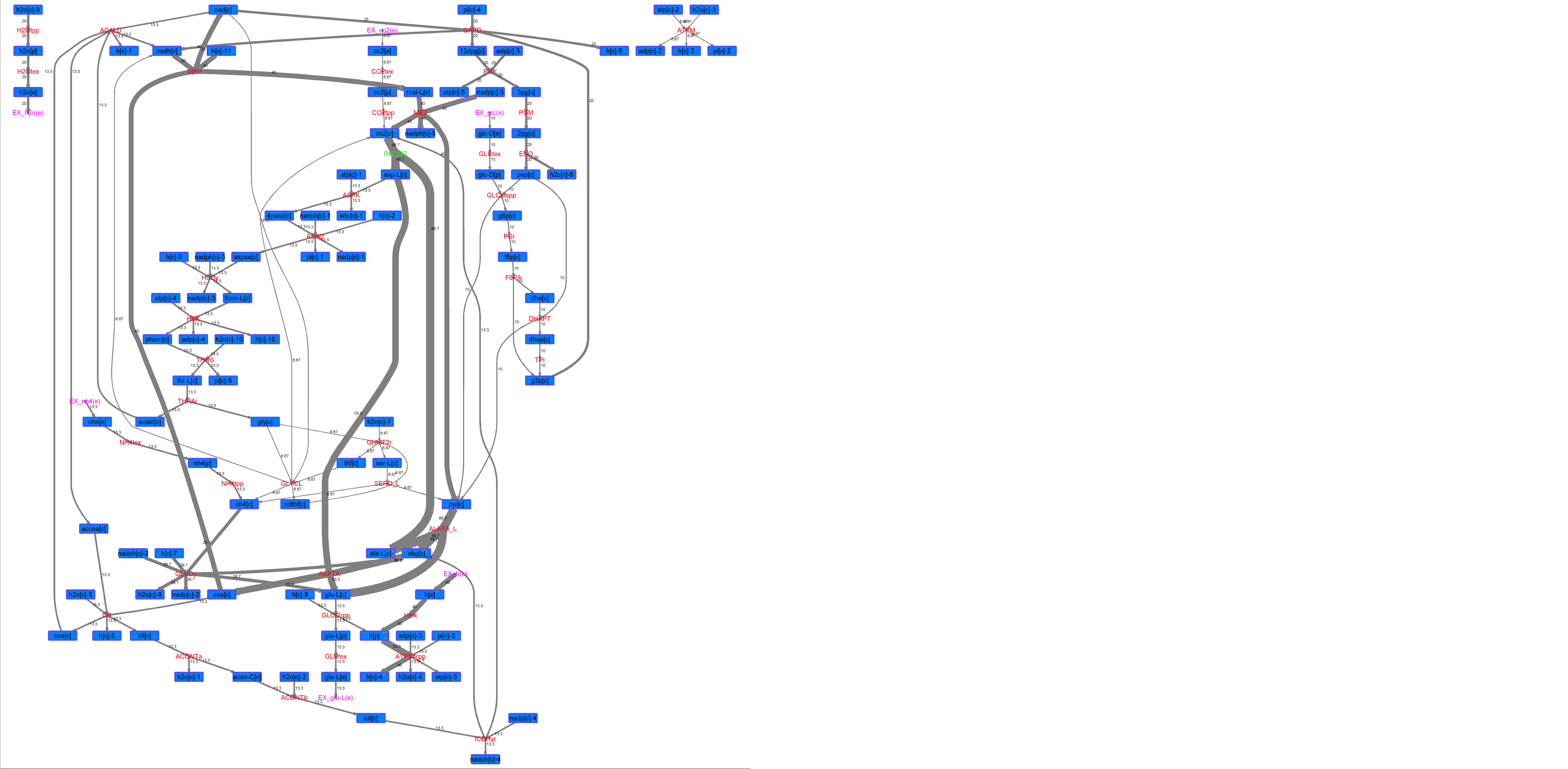

### R00397_L-ala.png

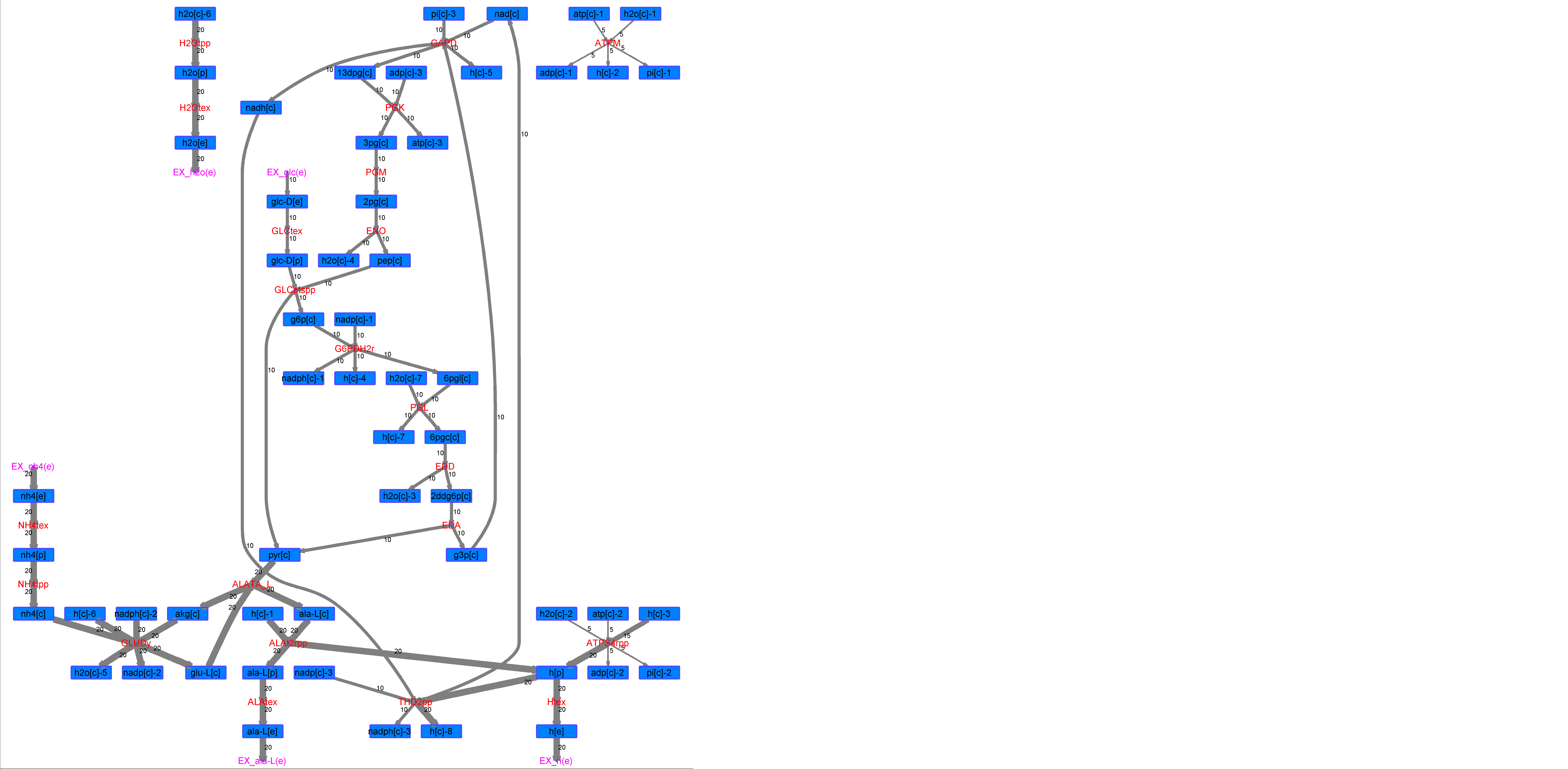

### R00397_L-Mal.png

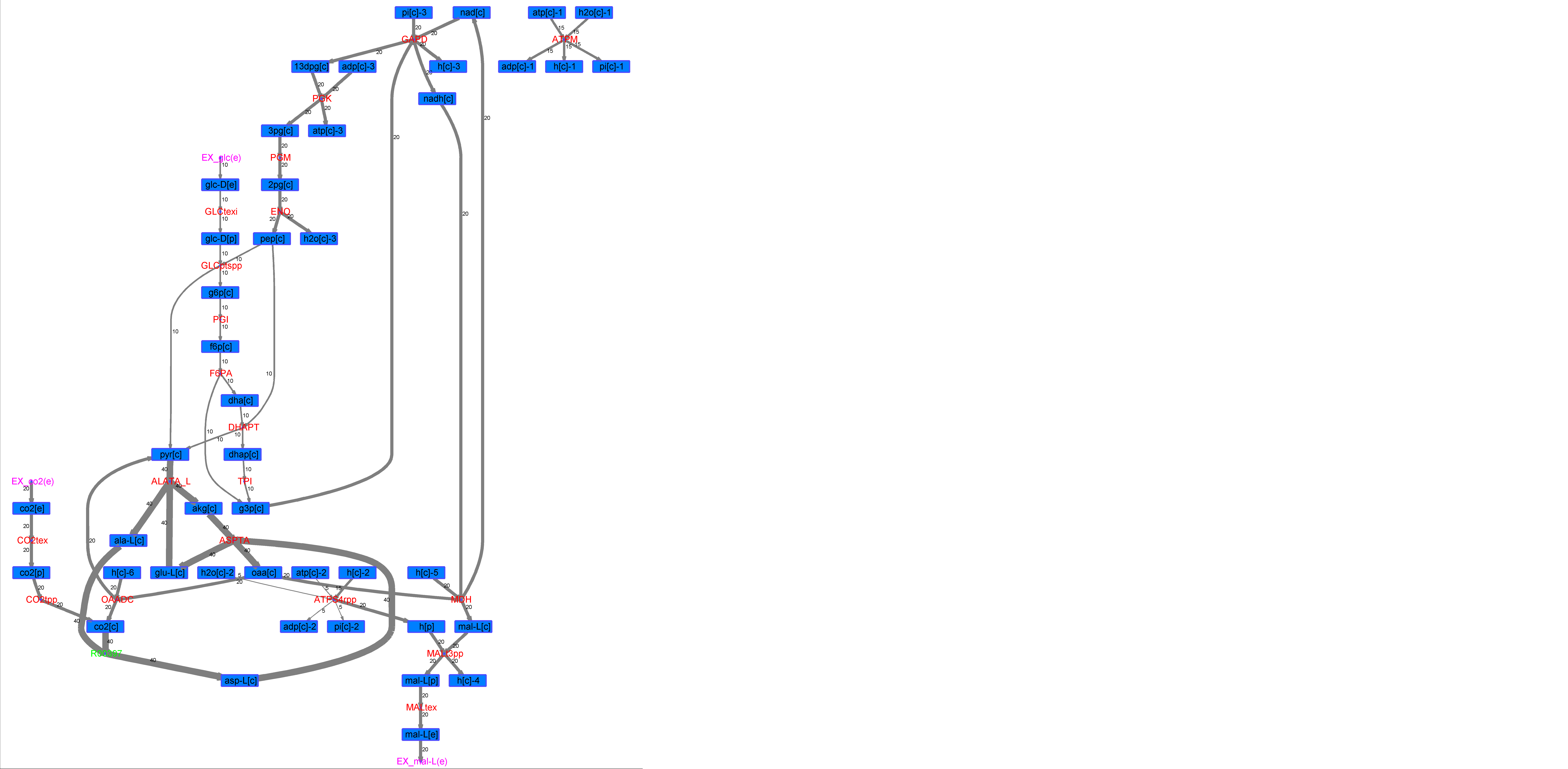

### R00397_L-Phe.png

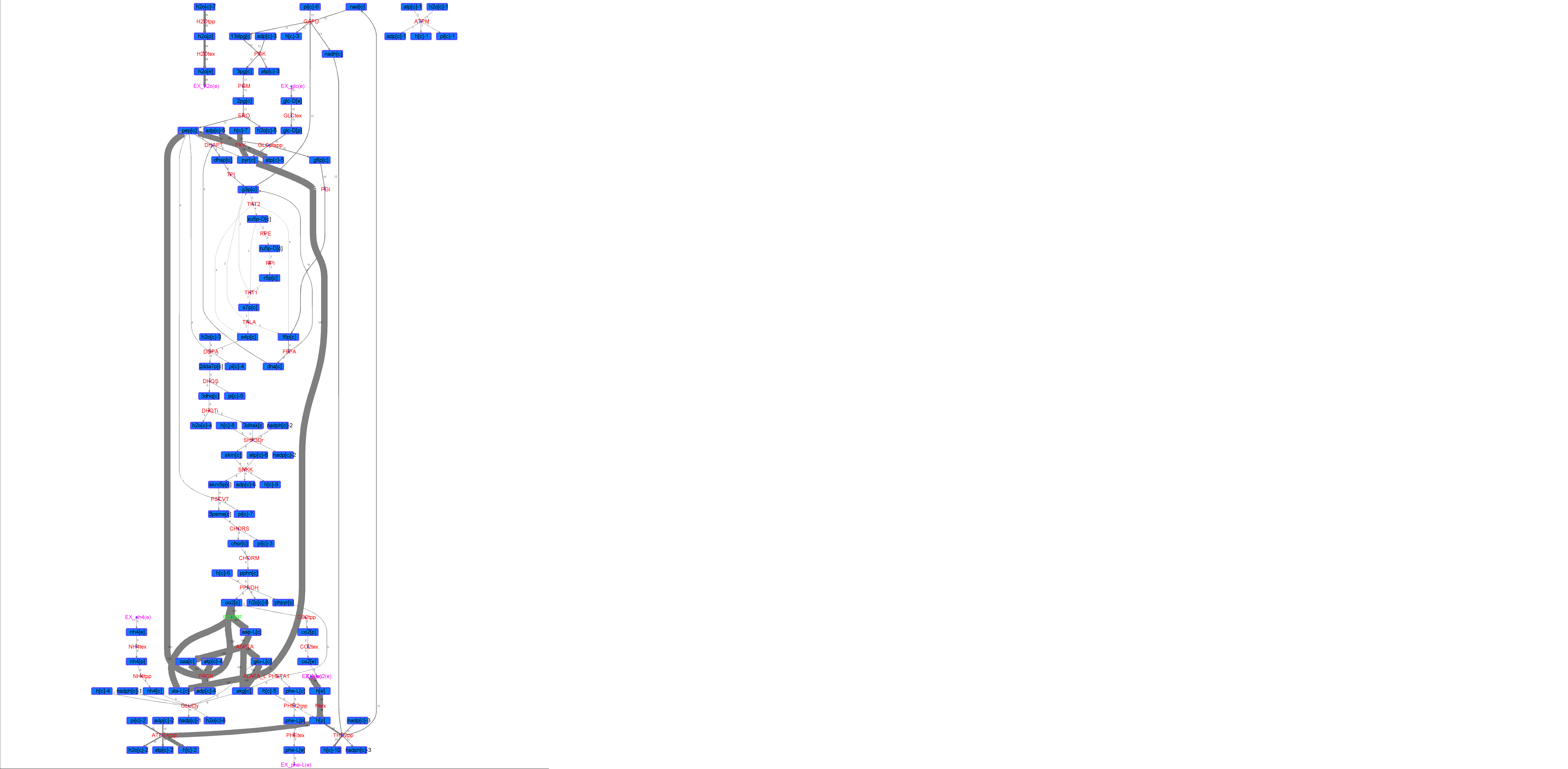

### R00397_L-Try.png

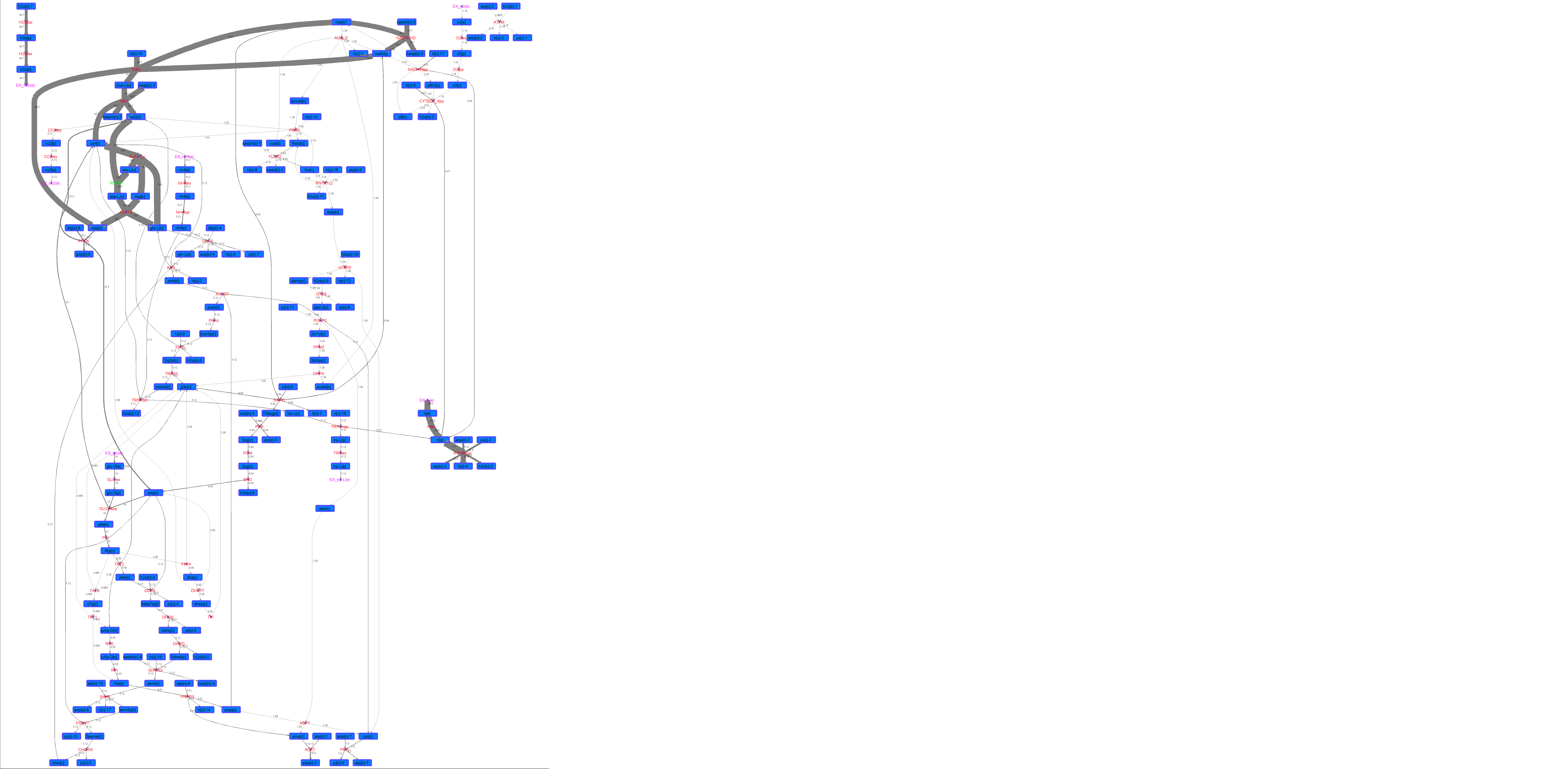

### R00397_Lys.png

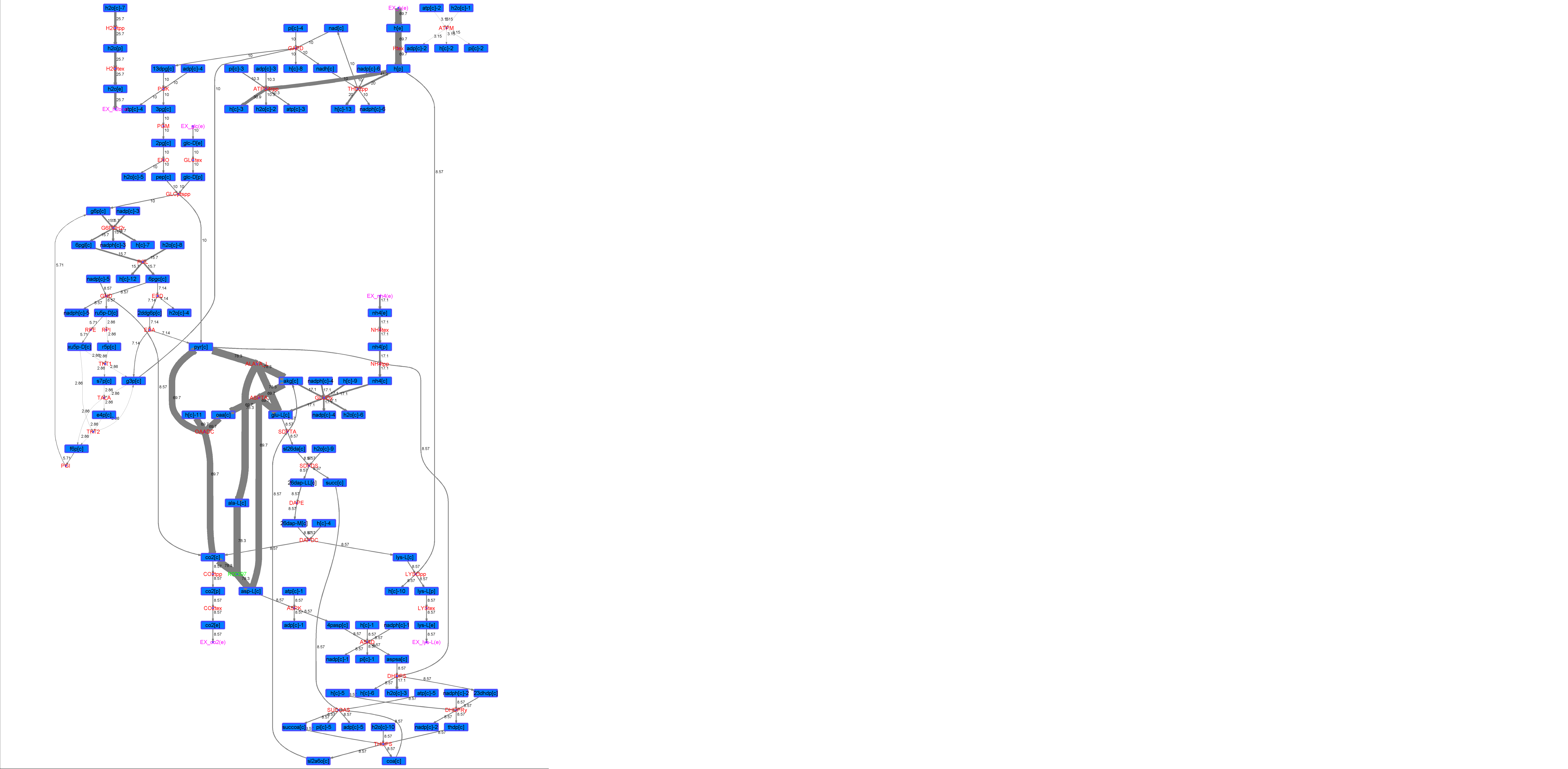

### R00397_Suc.png

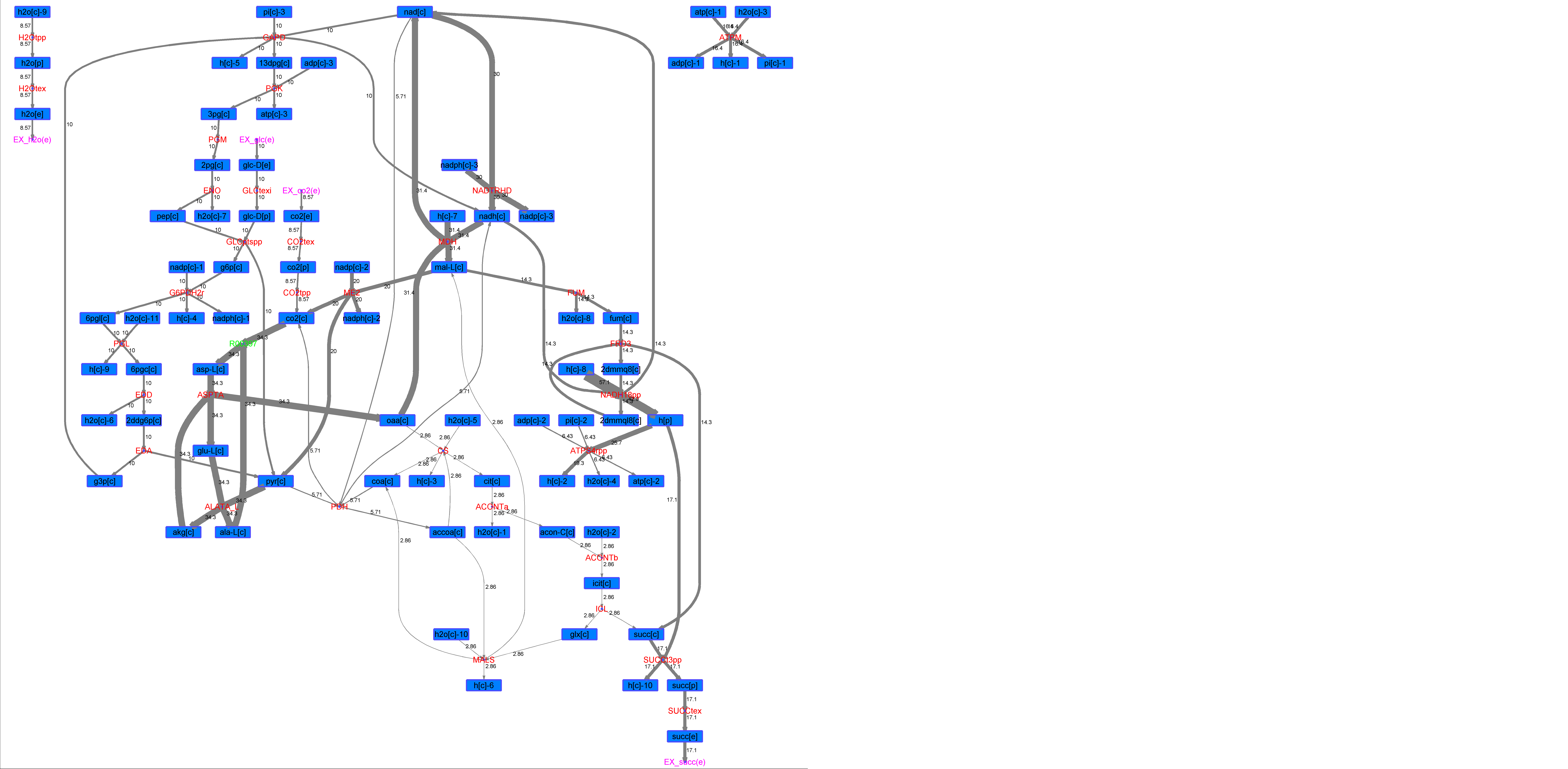

### R00397_Thr.png

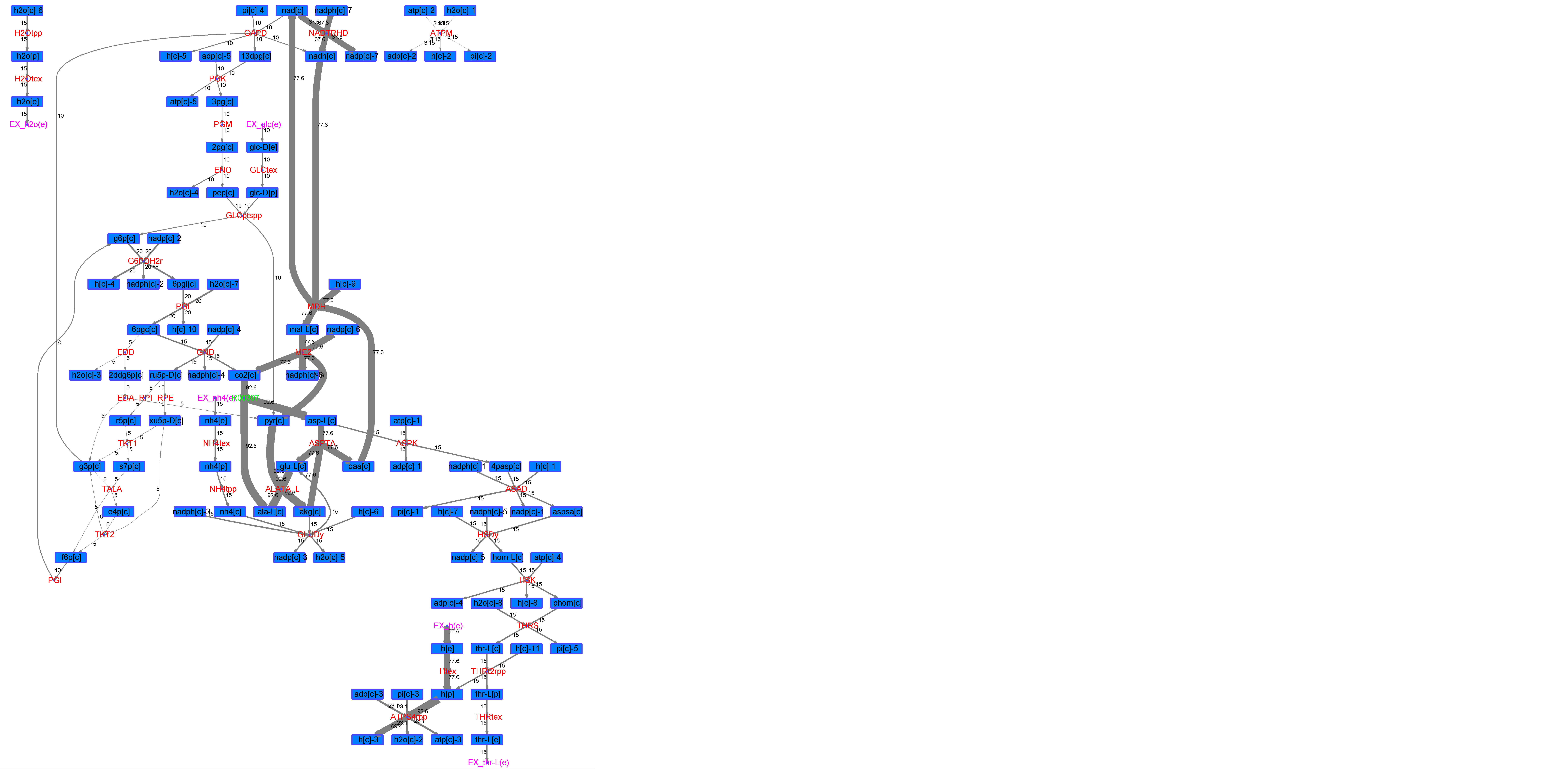

### R00430_Ace.png

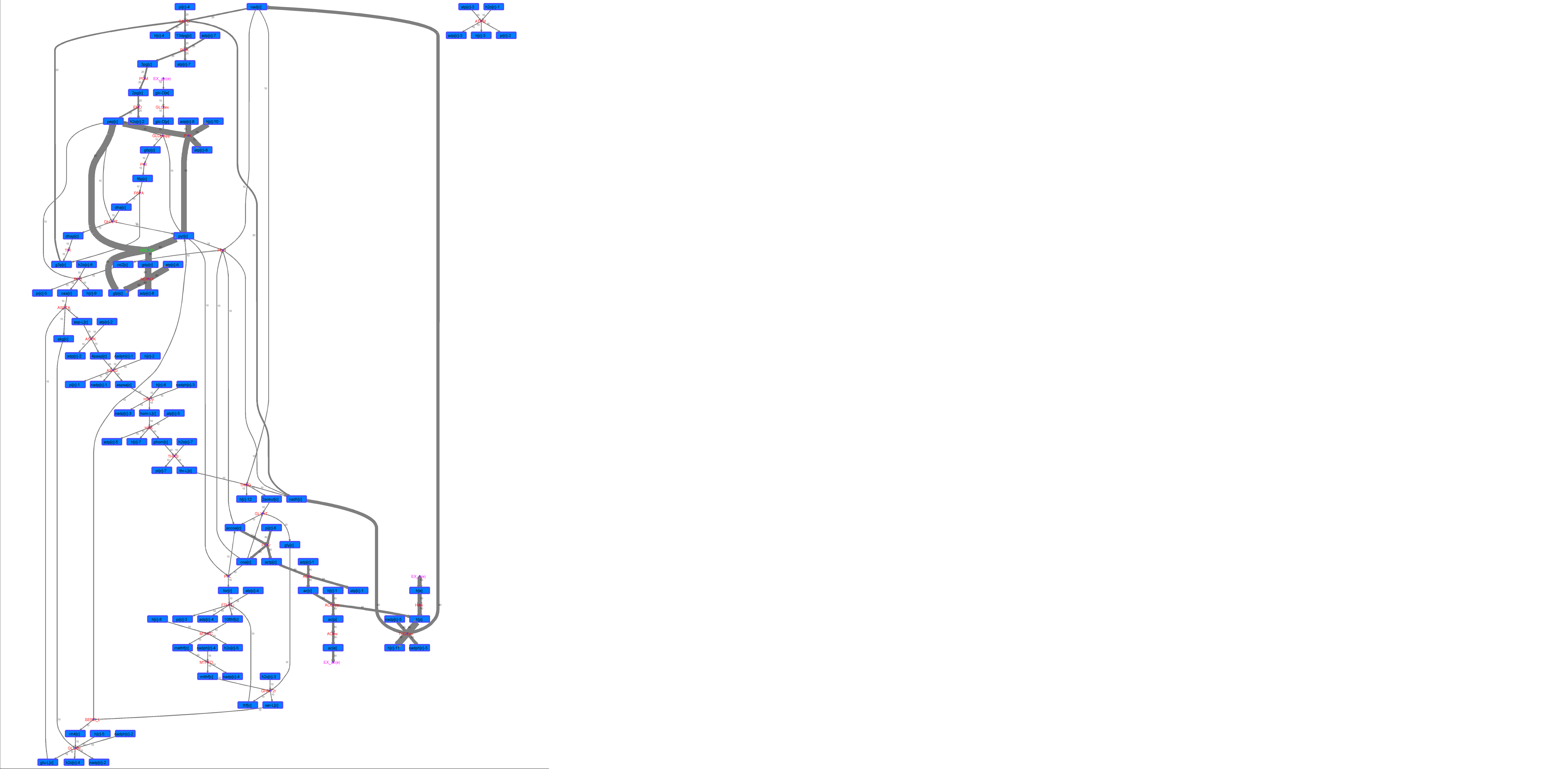

### R00430_For.png

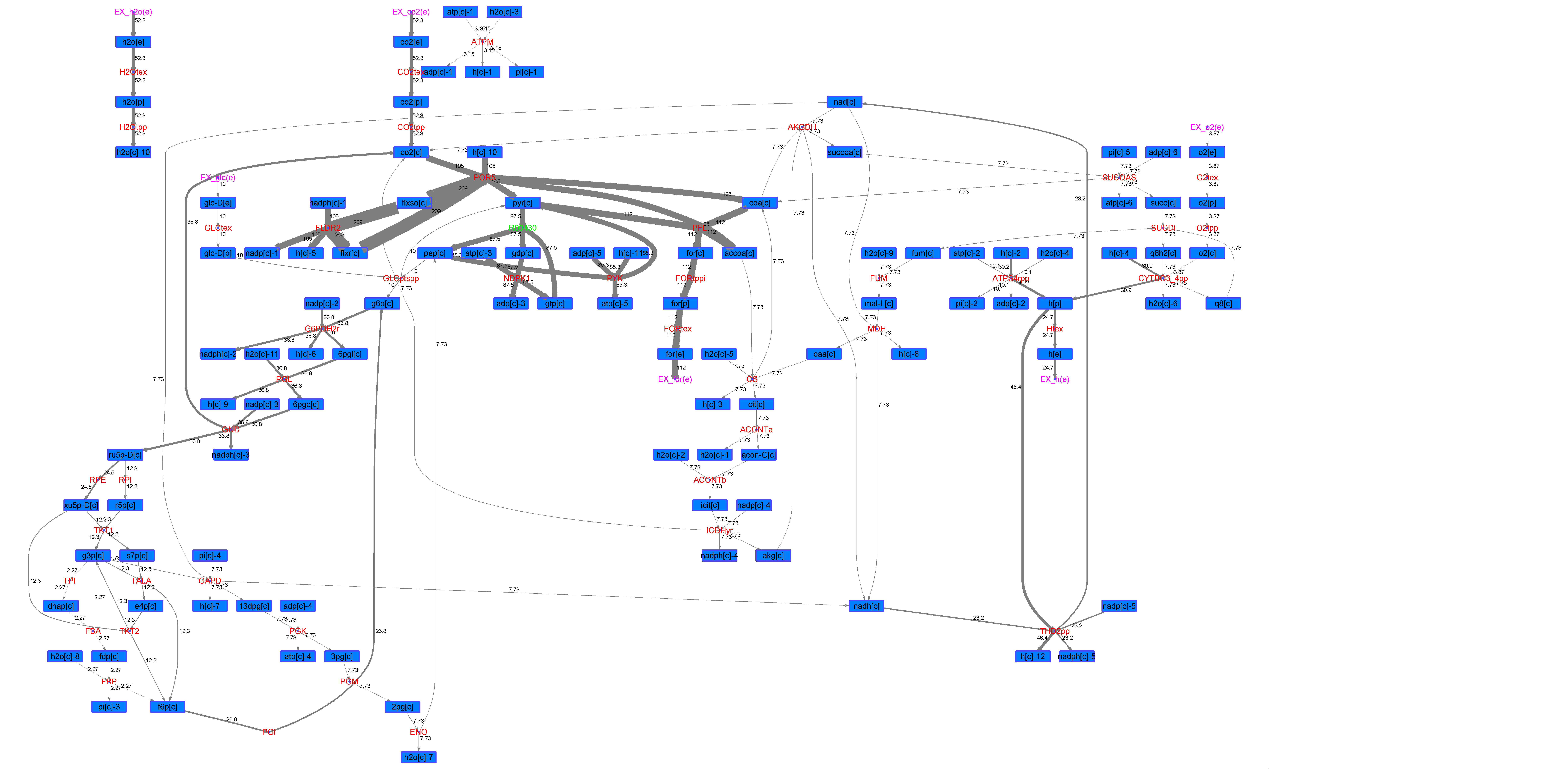

### R00430_Glu.png

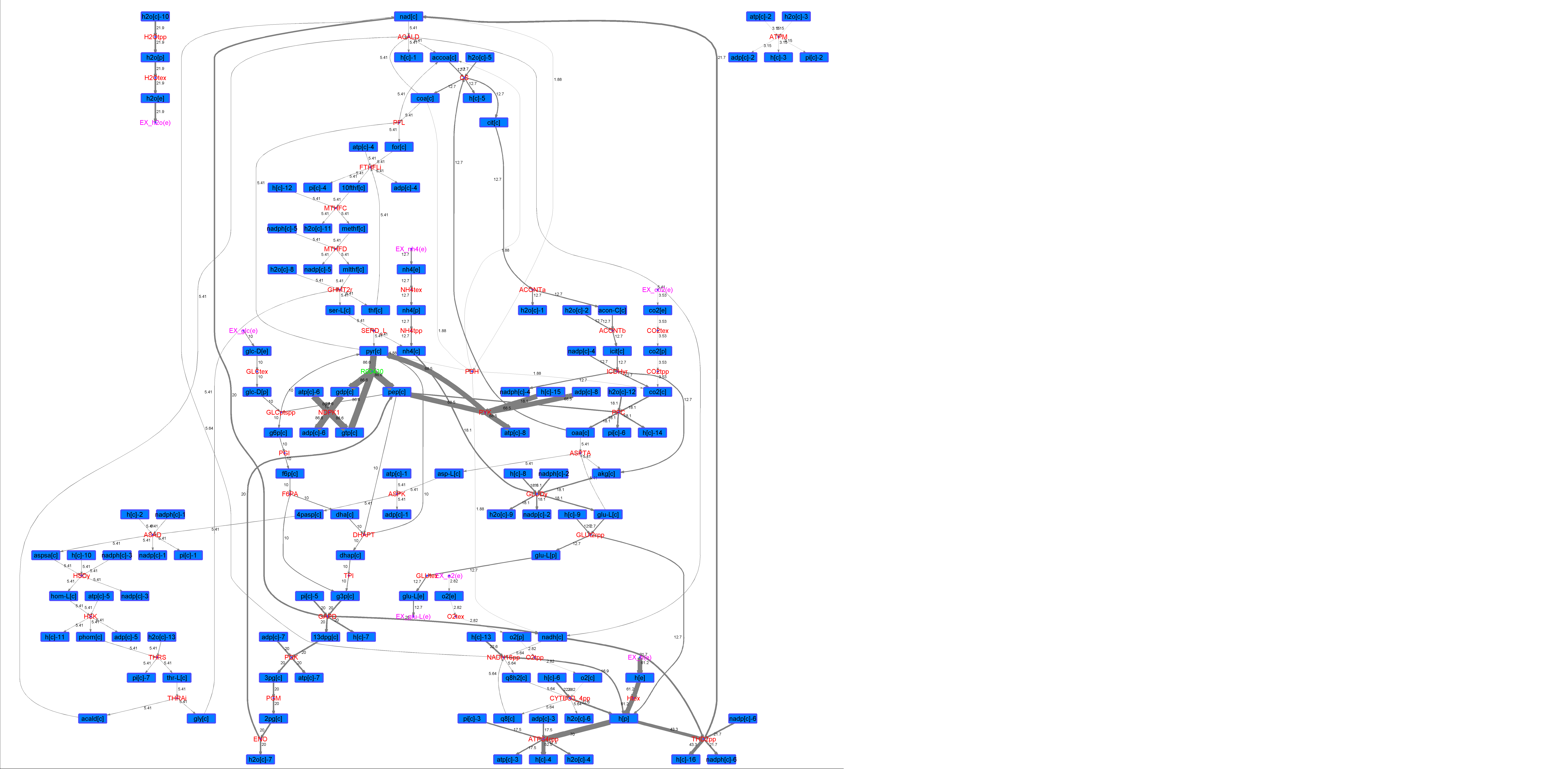

### R00430_L-ala.png

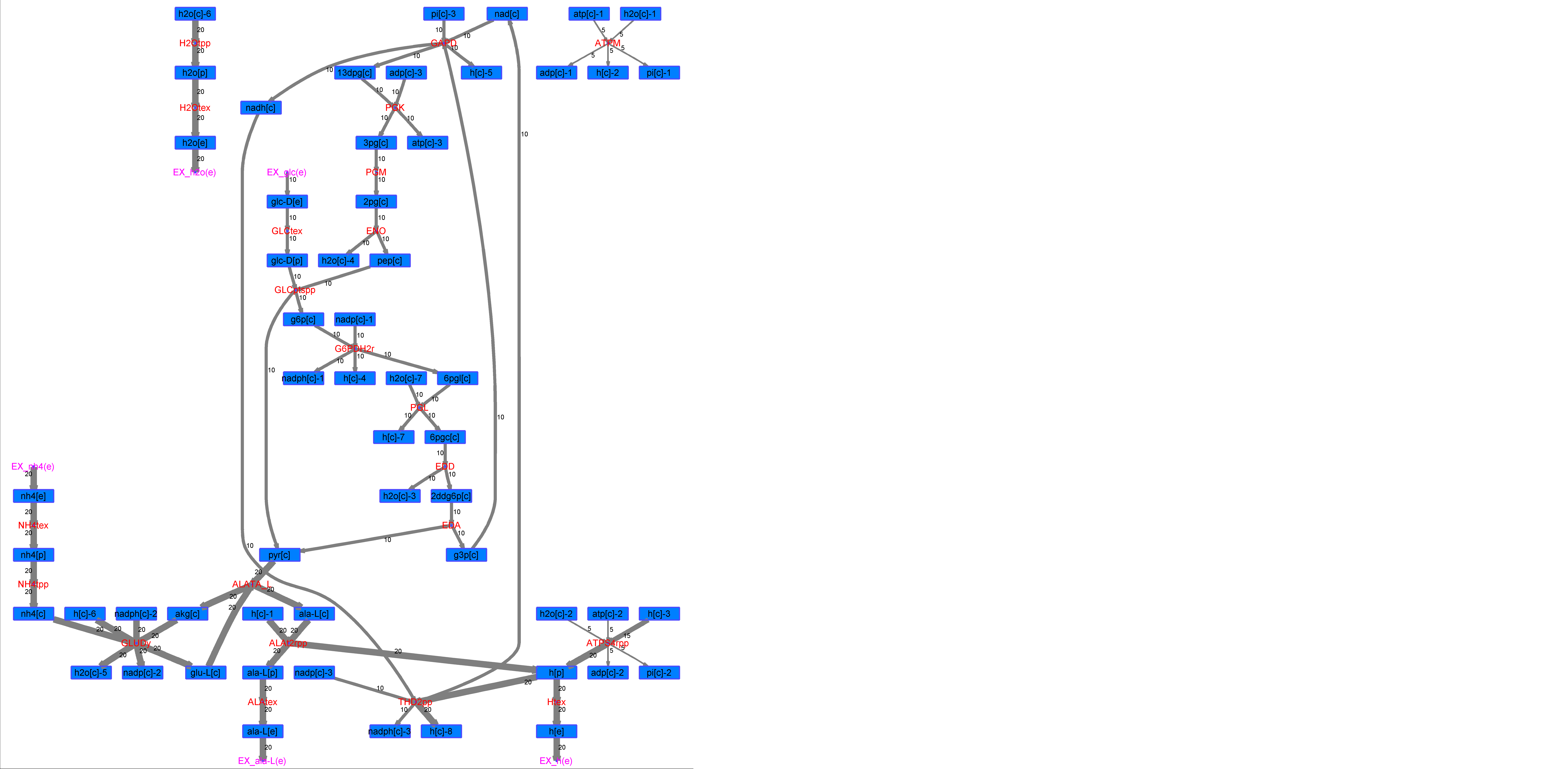

### R00430_L-Mal.png

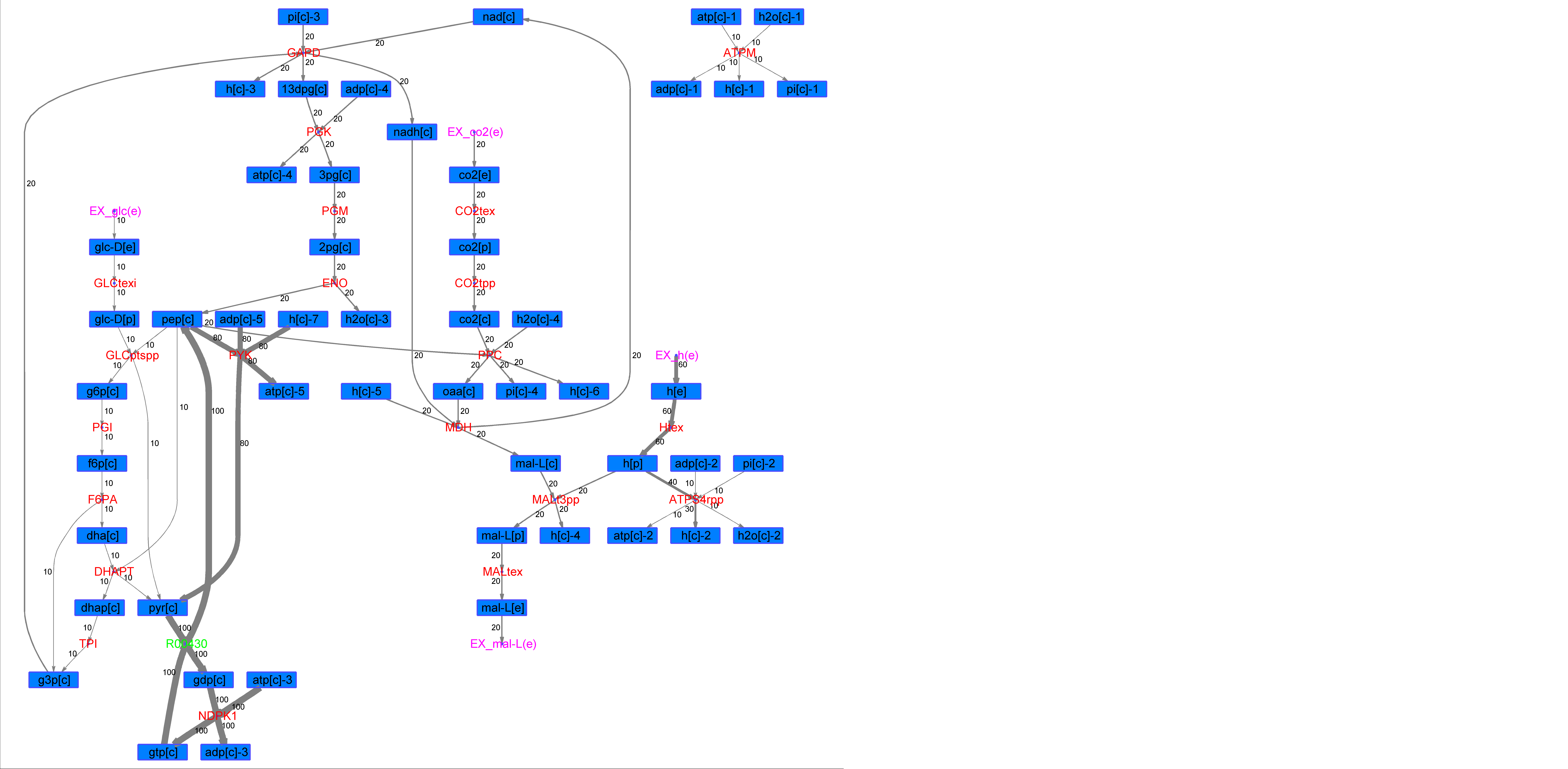

### R00430_L-Phe.png

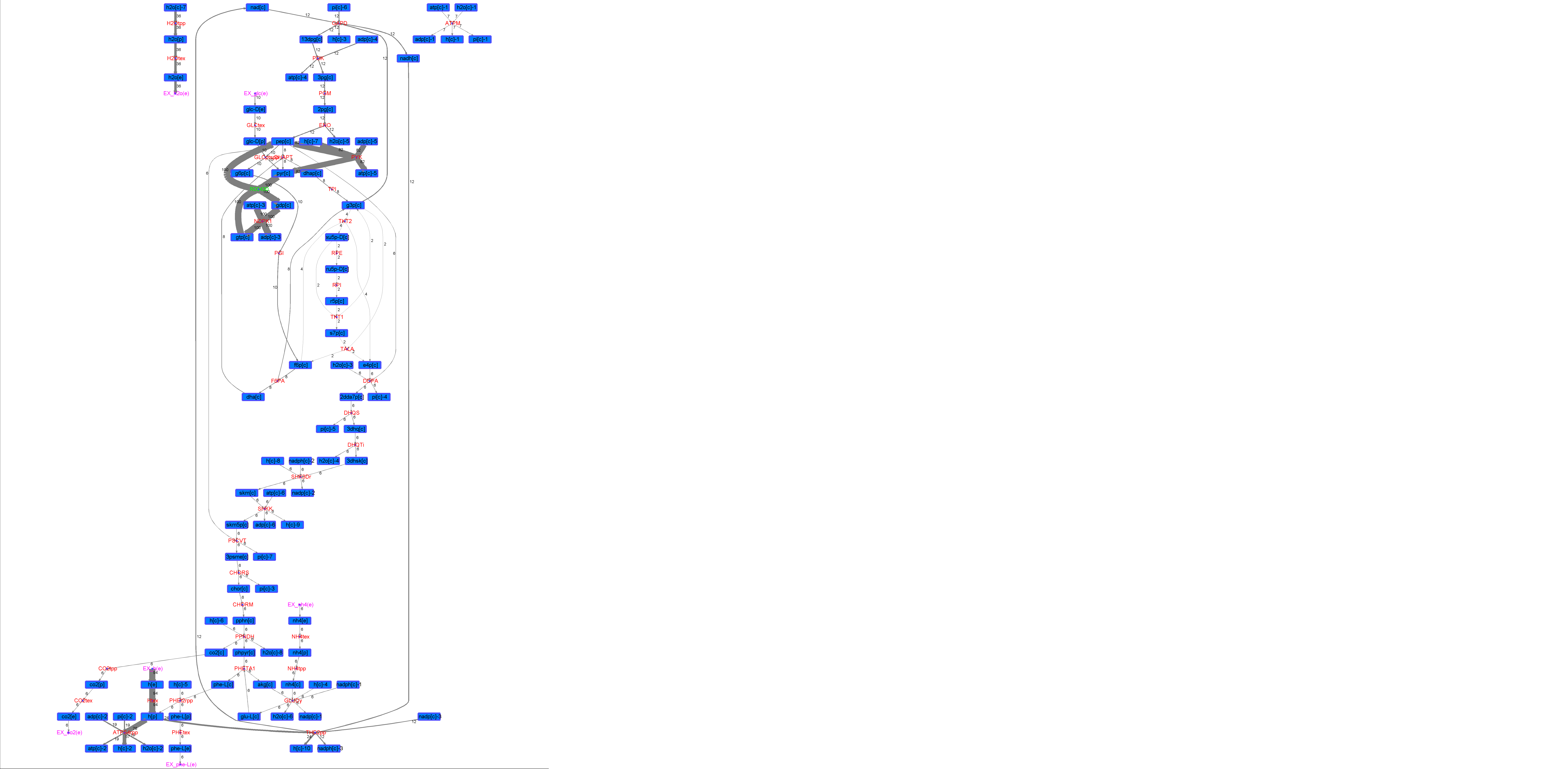

### R00430_L-Try.png

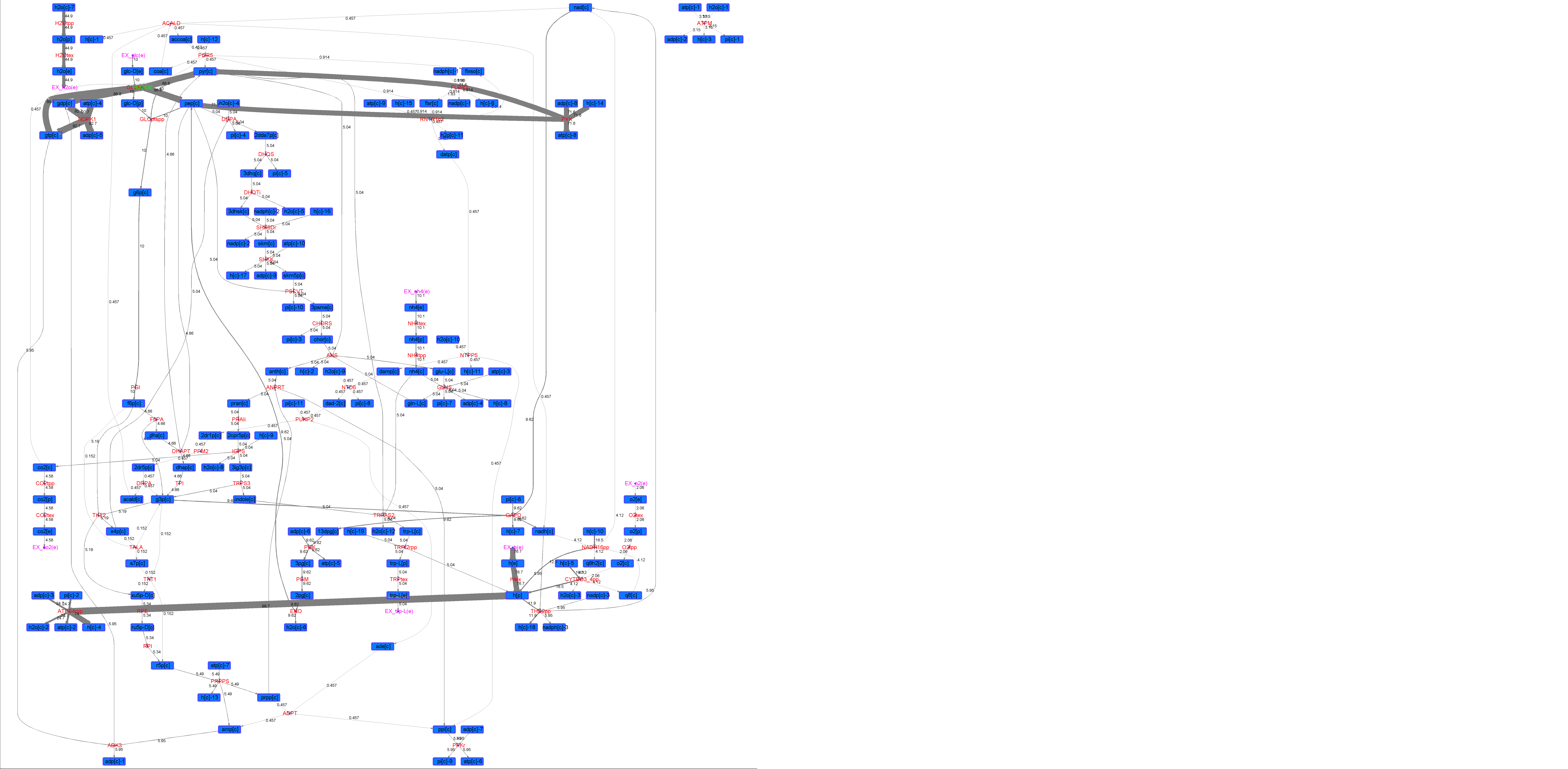

### R00430_Lys.png

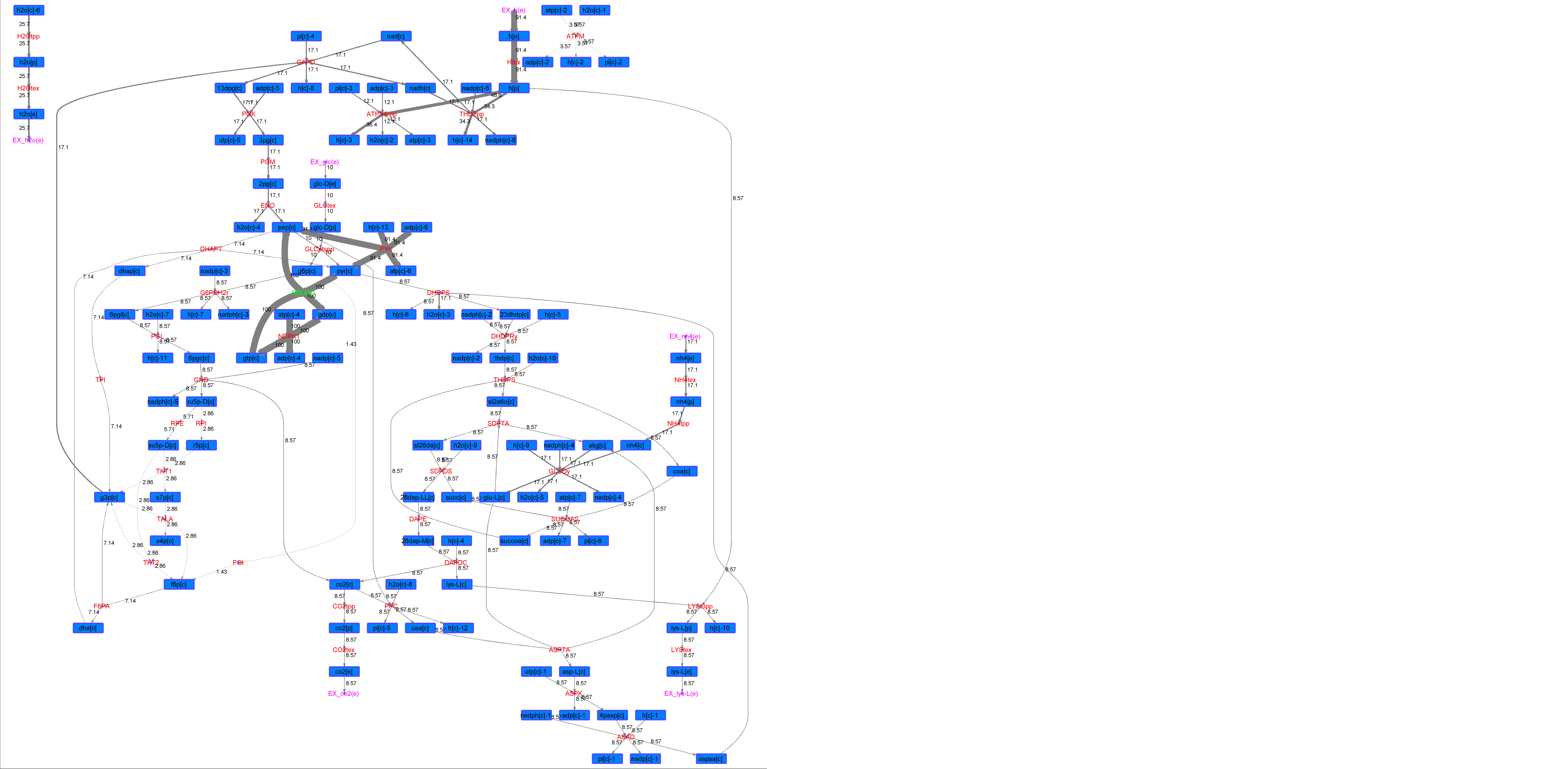

### R00430_Suc.png

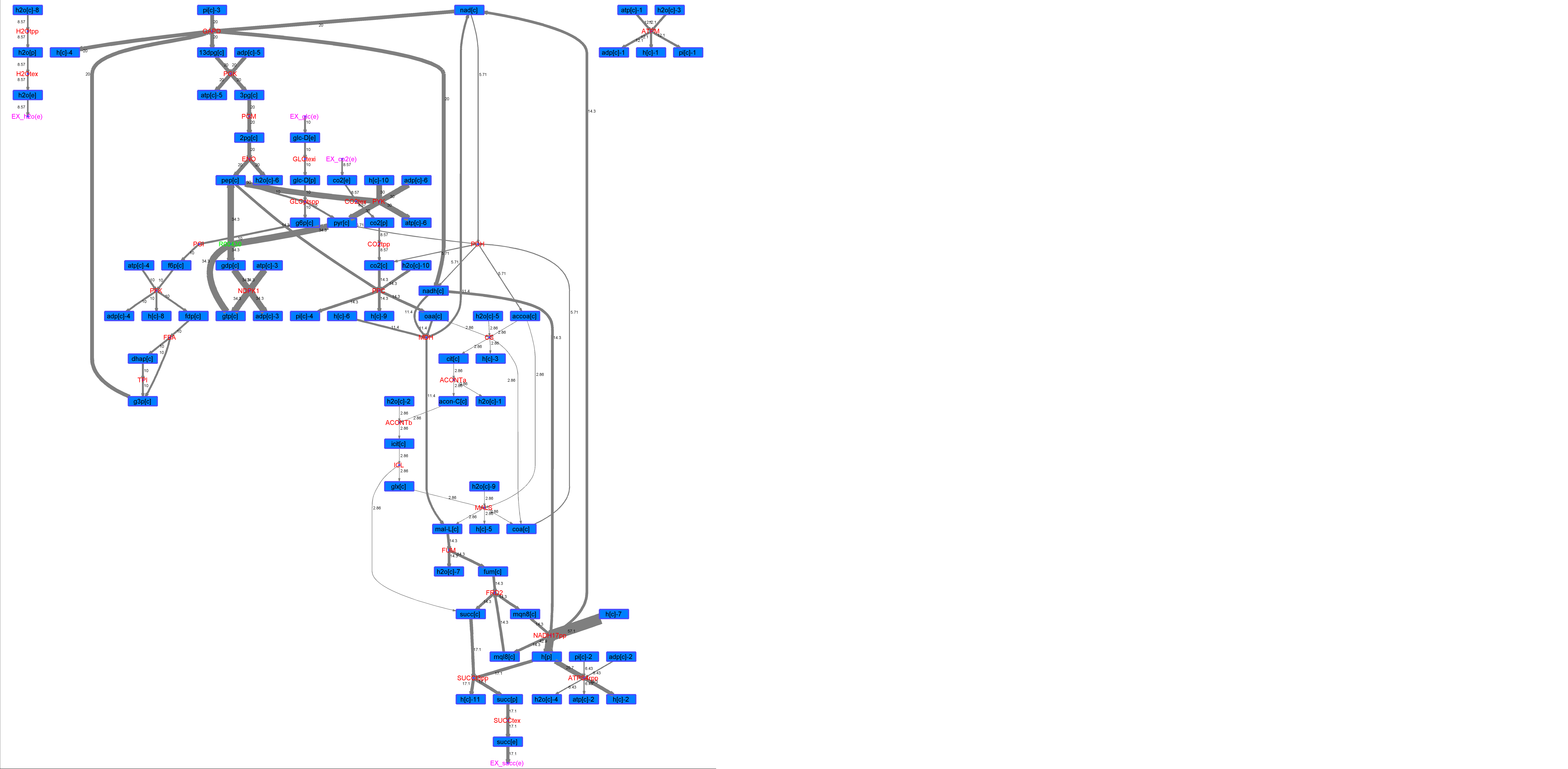

### R00430_Thr.png

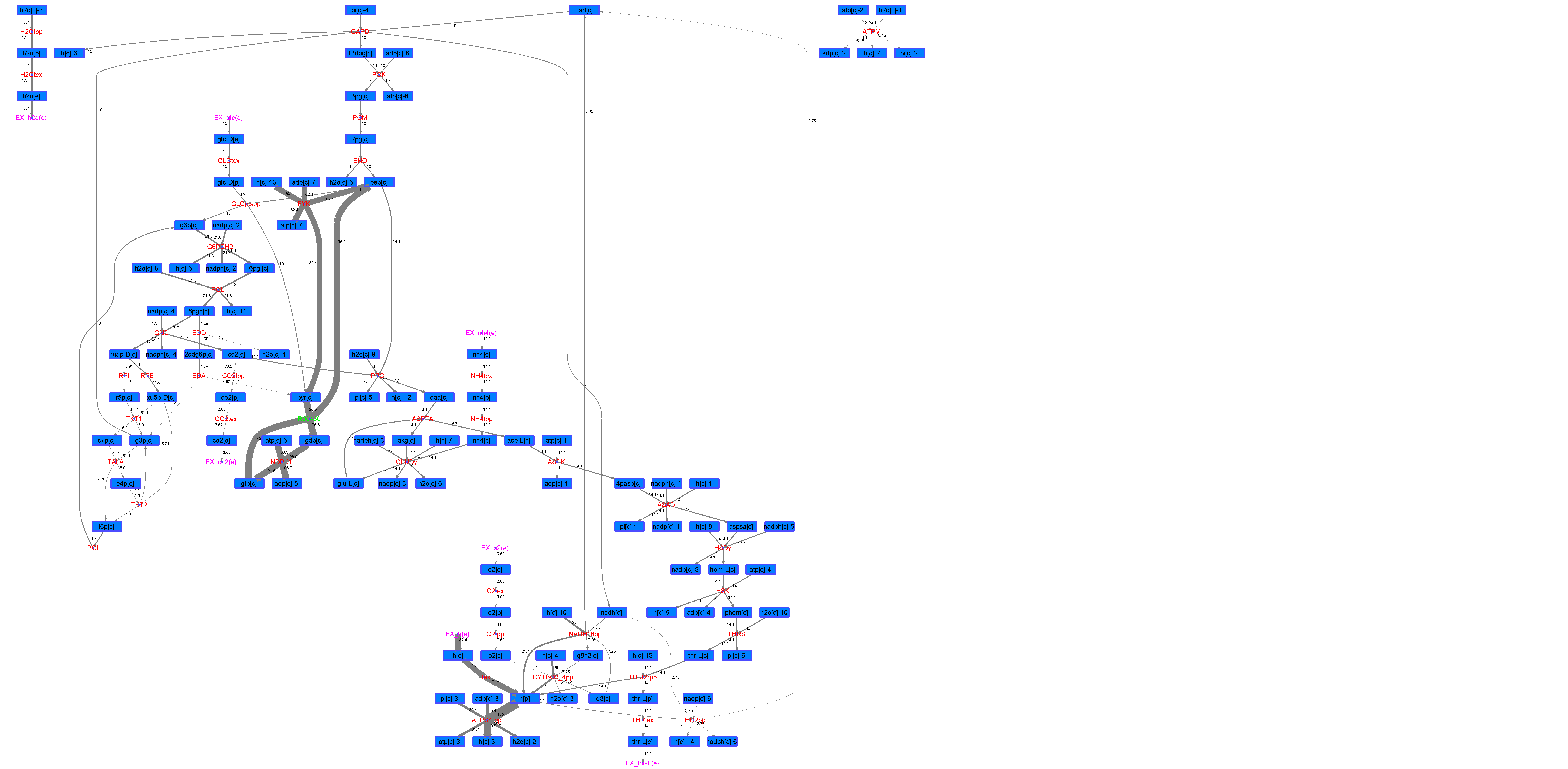

### R00431_Ace.png

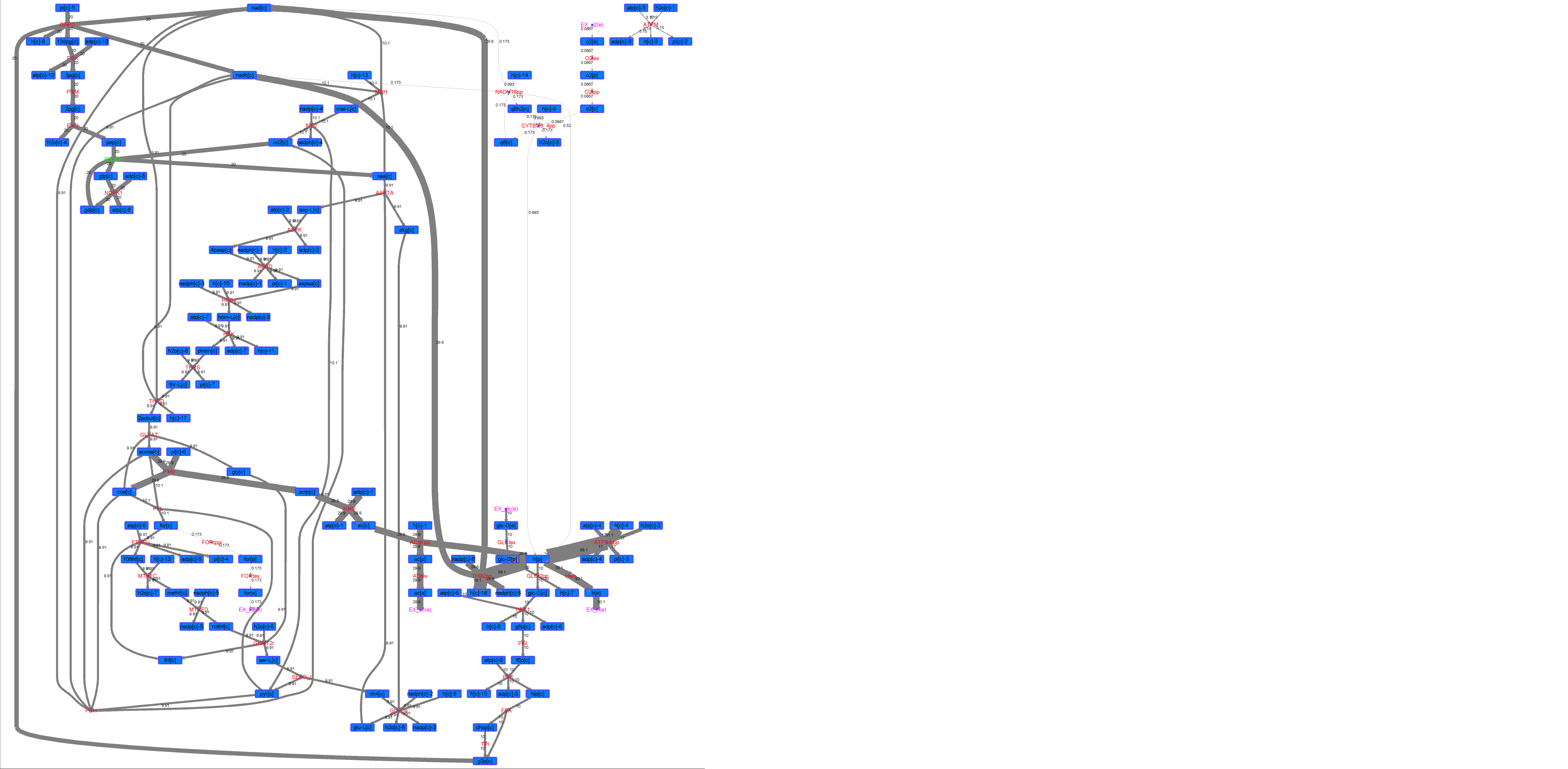

### R00431_For.png

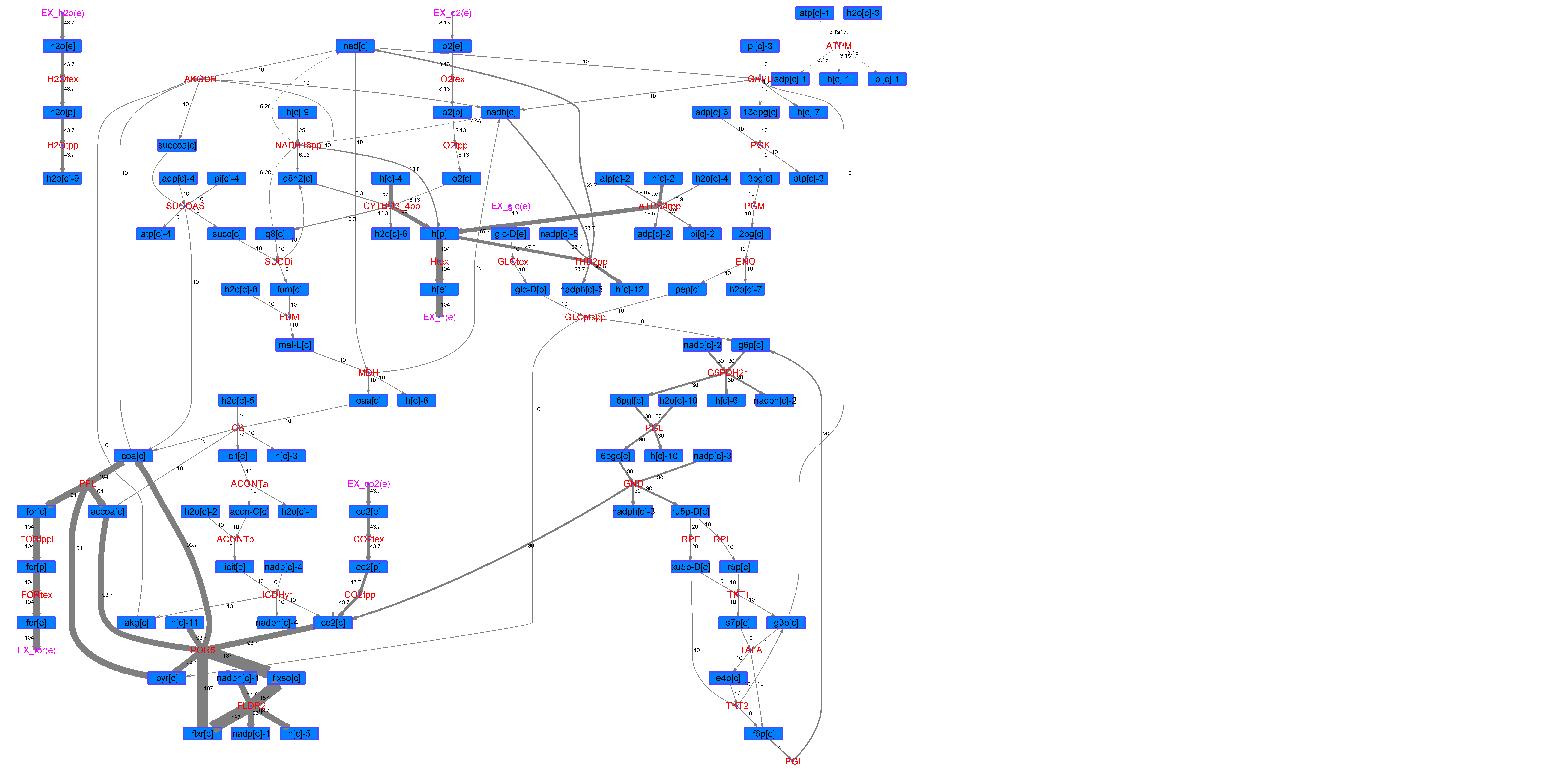

### R00431_Glu.png

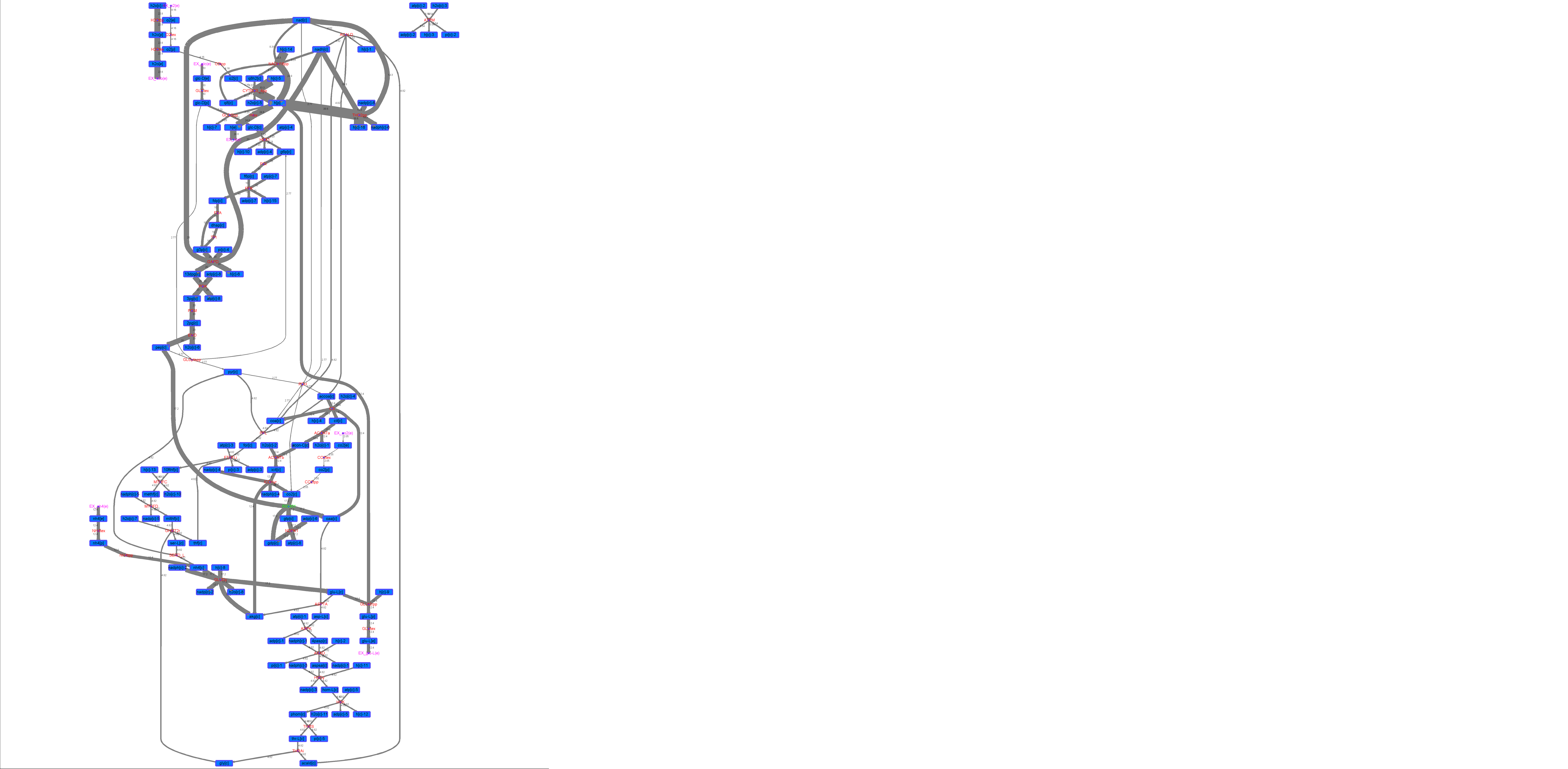

### R00431_L-ala.png

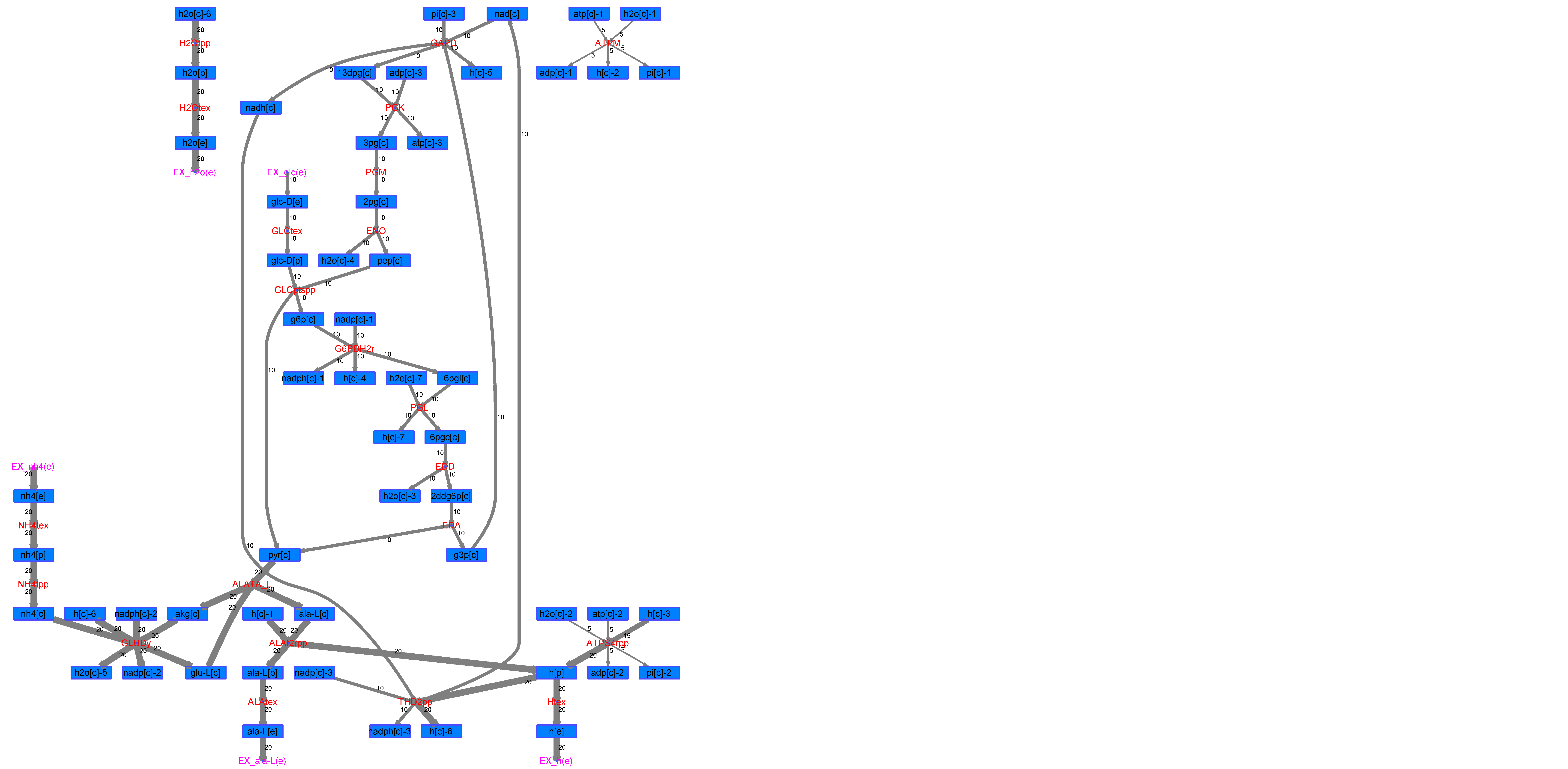

### R00431_L-Mal.png

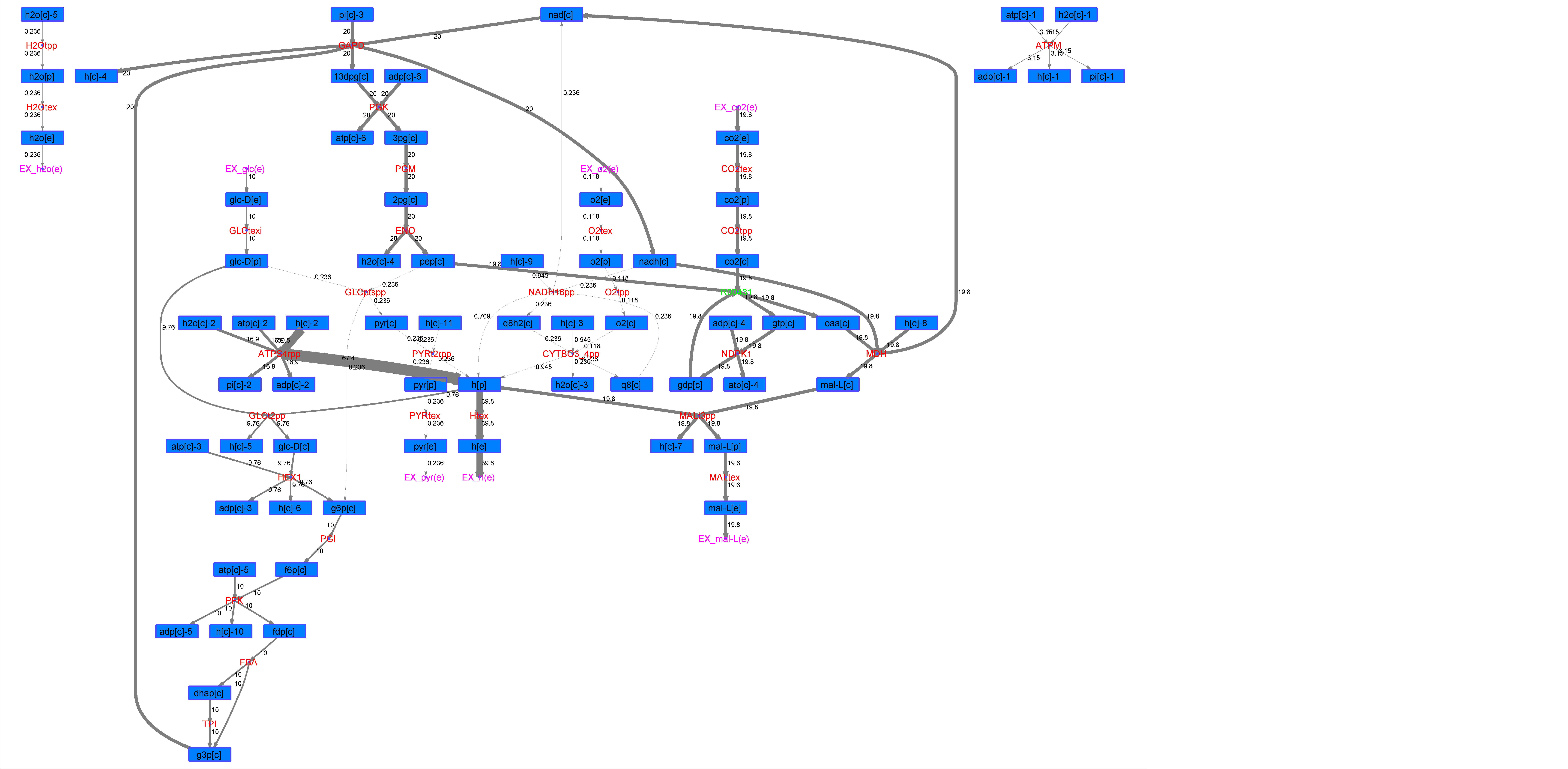

### R00431_L-Phe.png

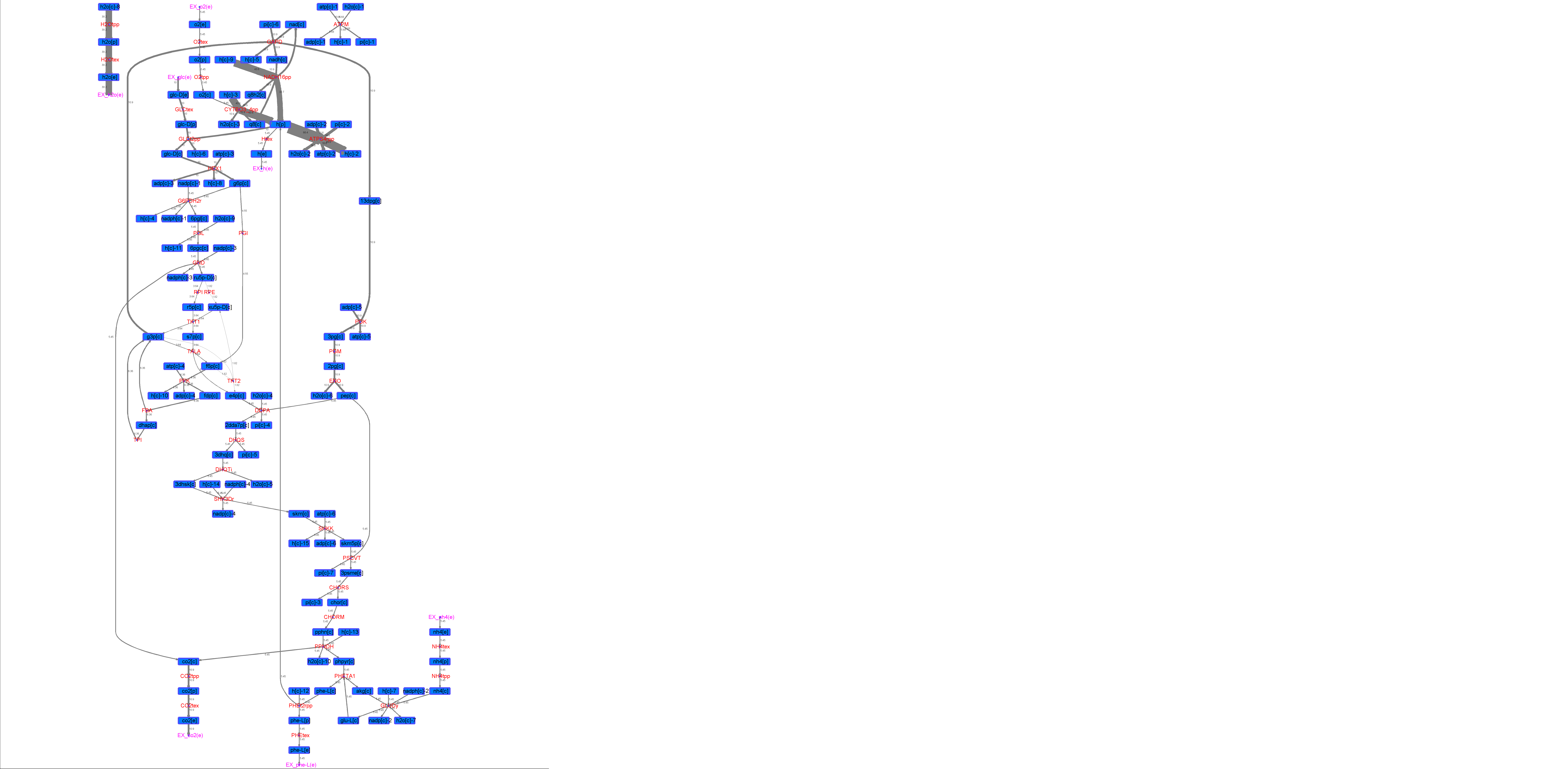

### R00431_L-Try.png

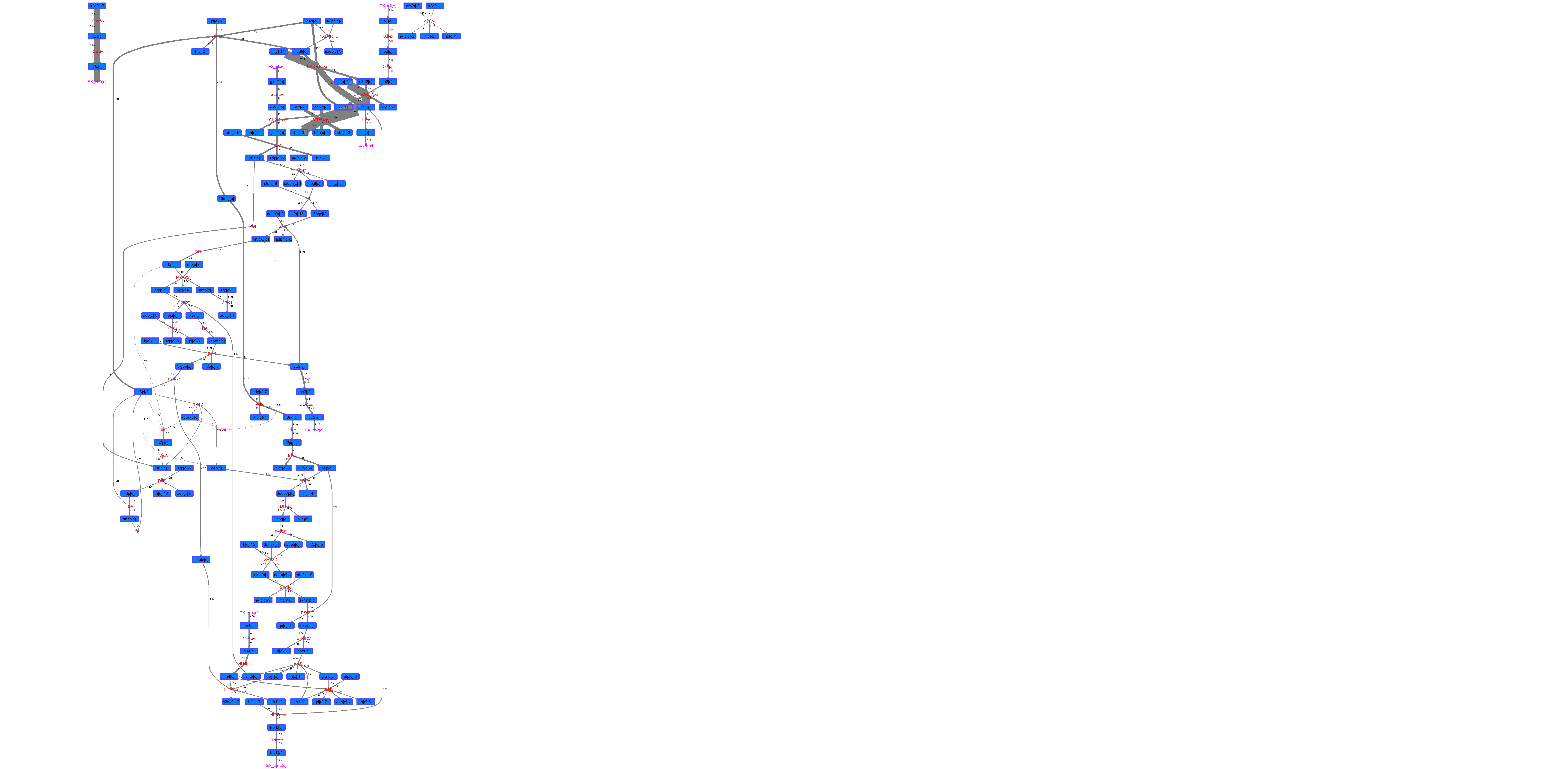

### R00431_Lys.png

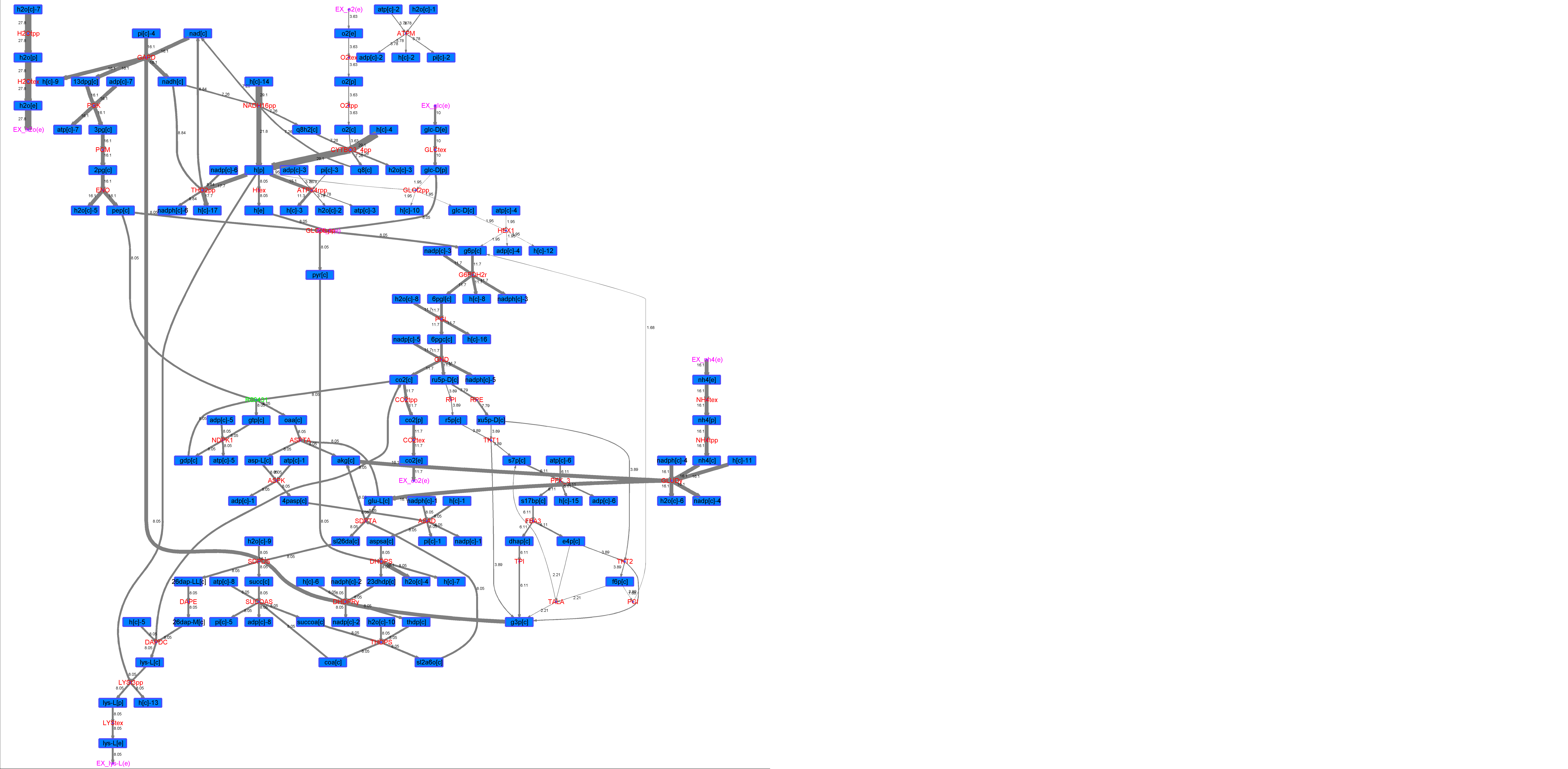

### R00431_Suc.png

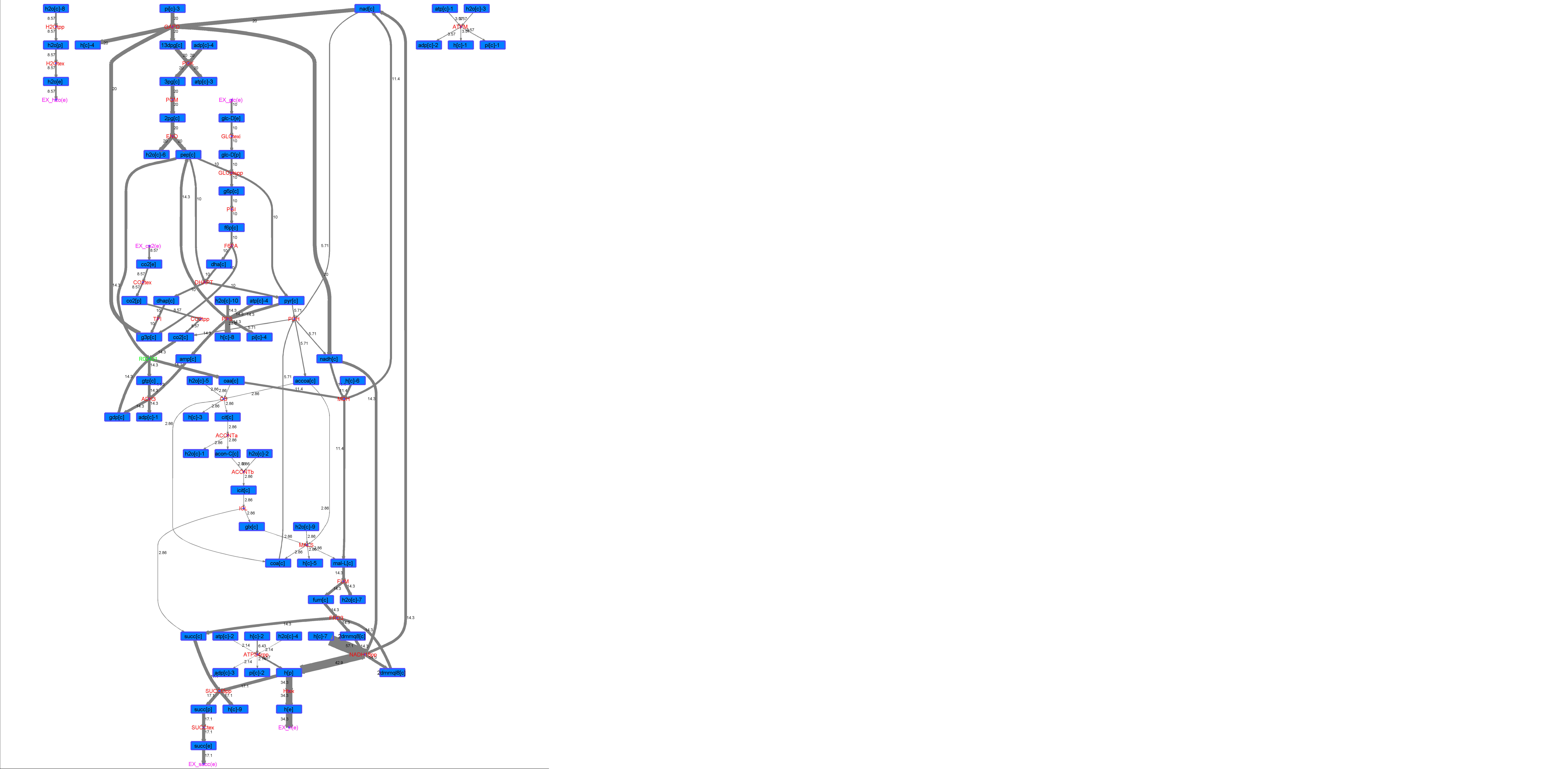

### R00431_Thr.png

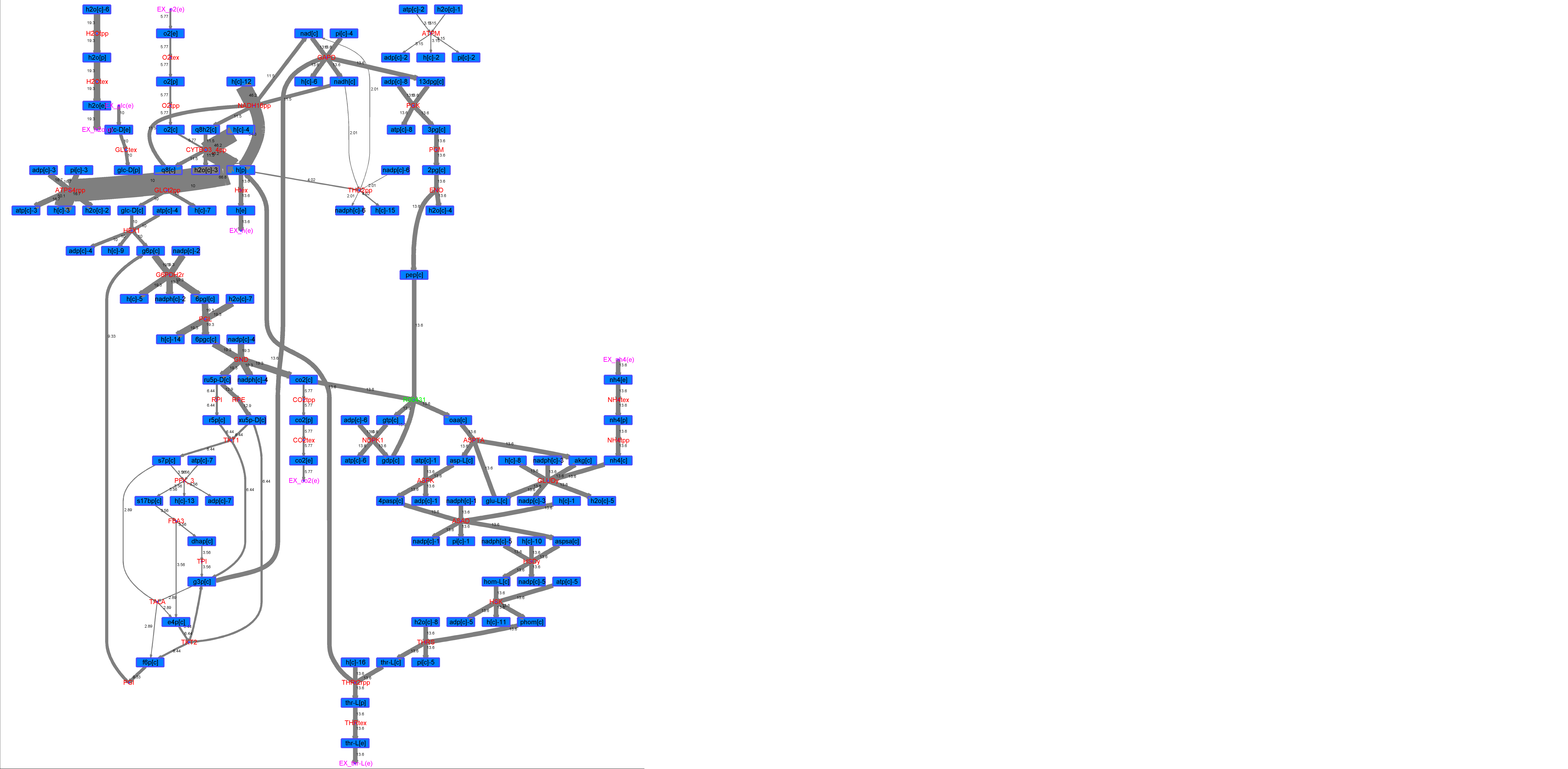

### R00502_Ace.png

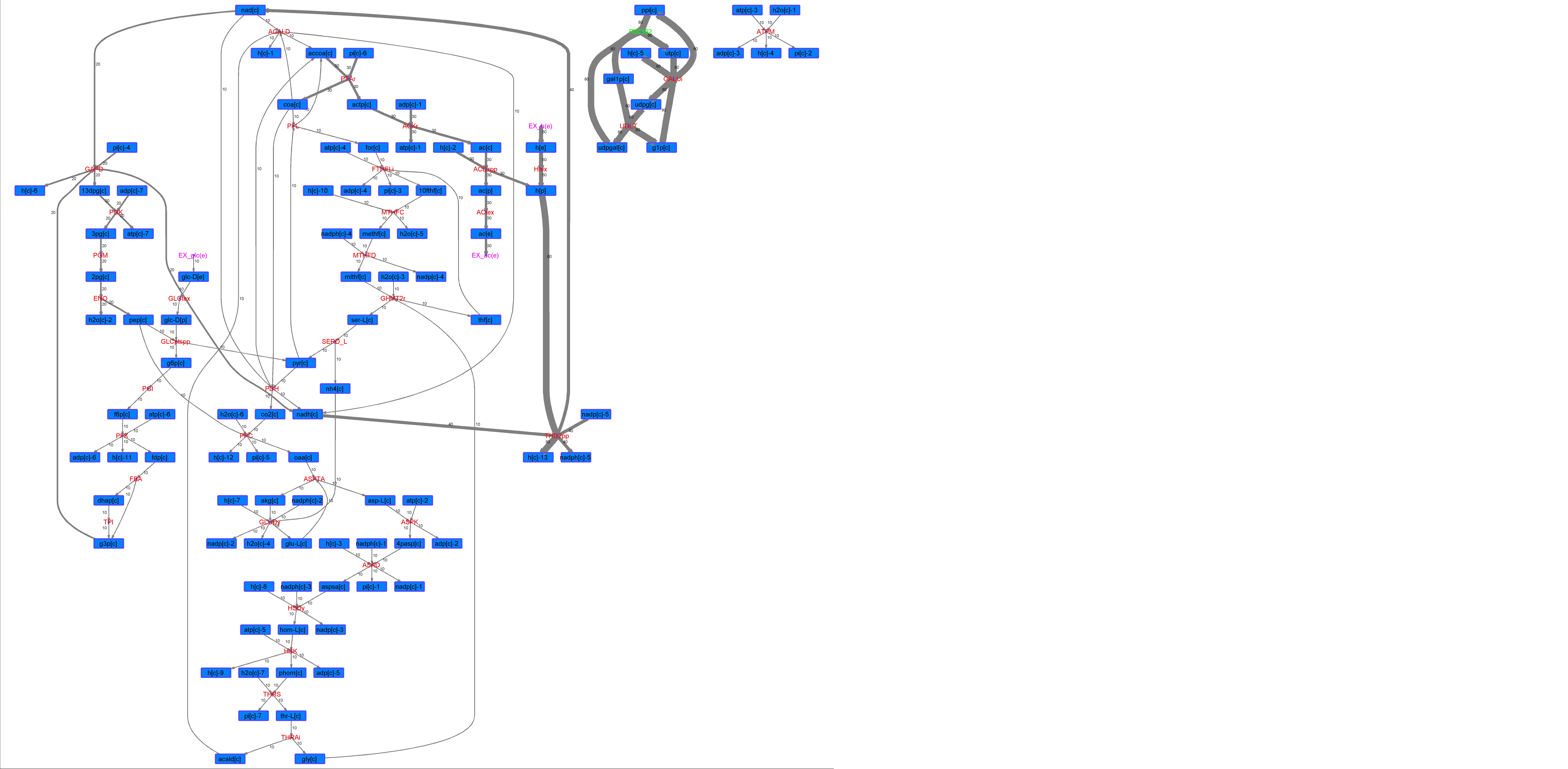

### R00502_For.png

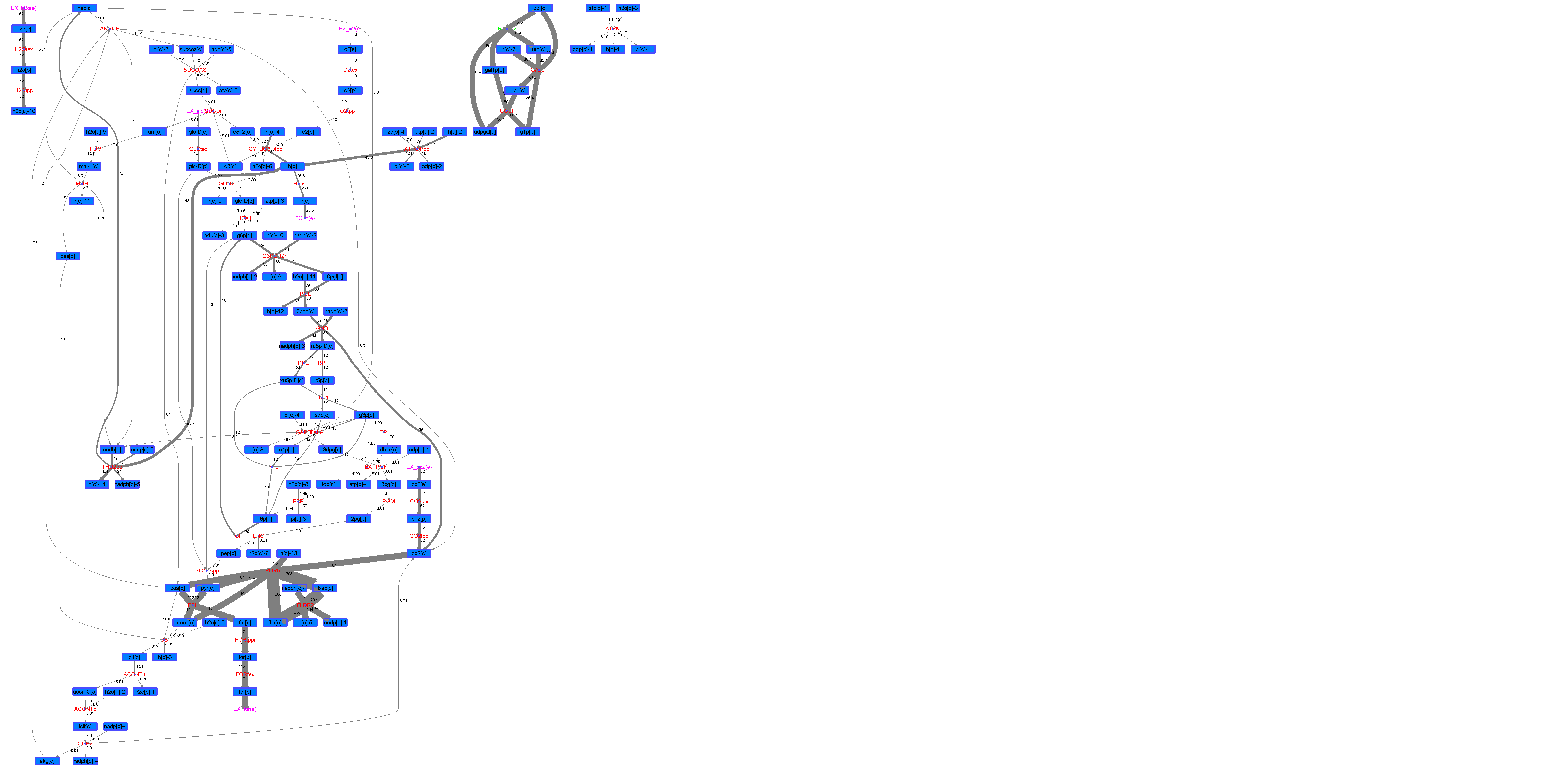

### R00502_Glu.png

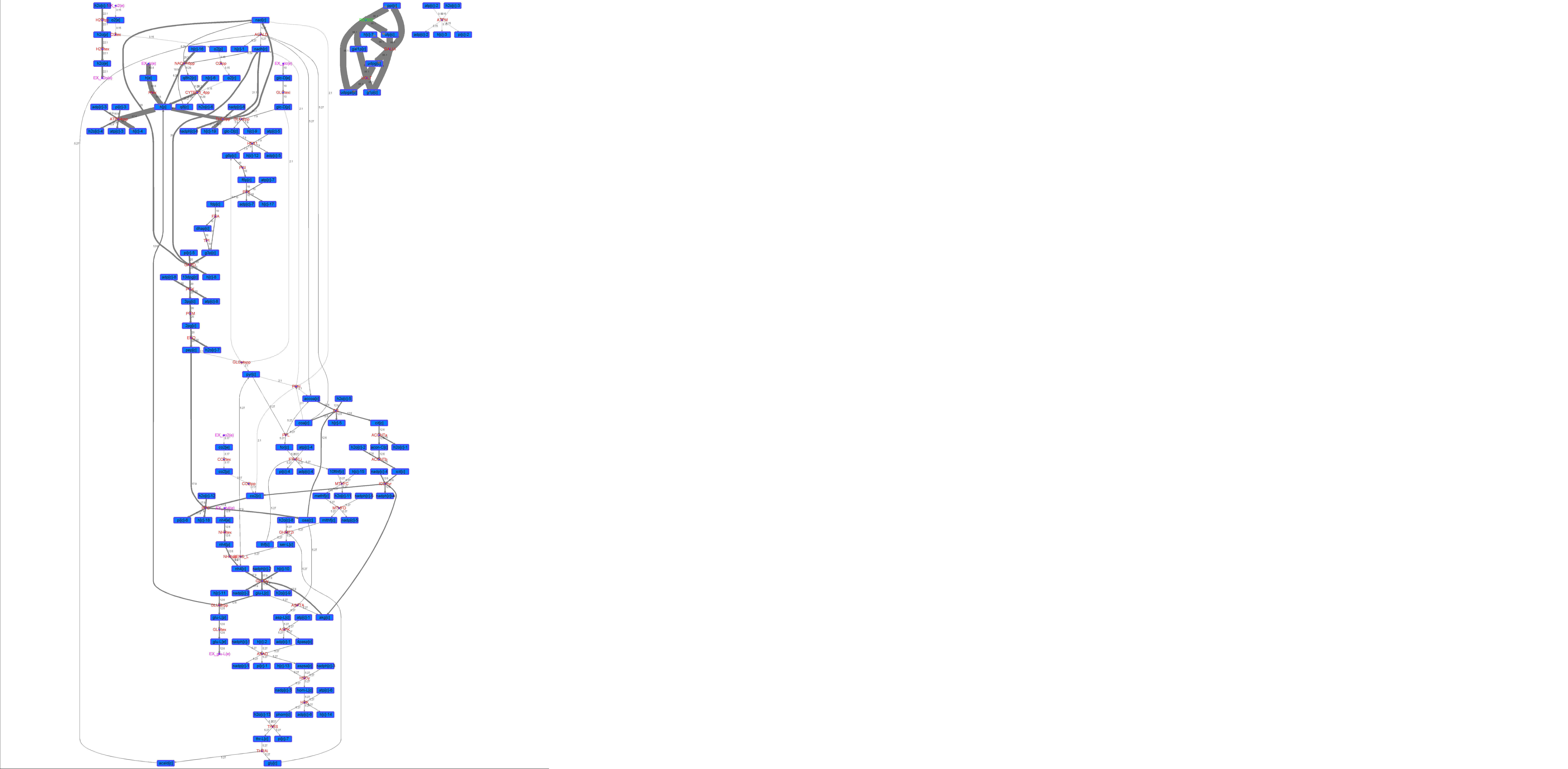

### R00502_L-ala.png

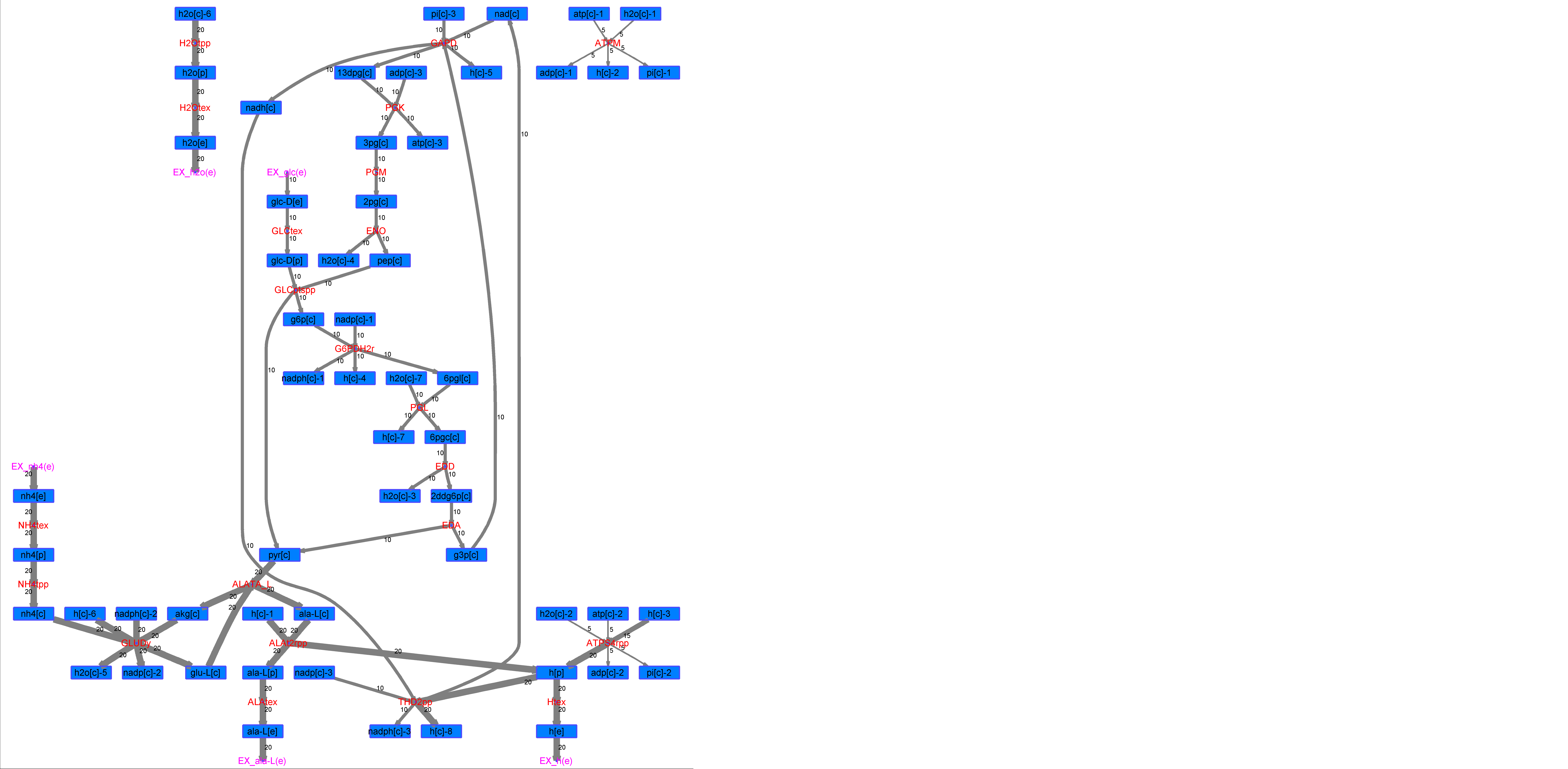

### R00502_L-Mal.png

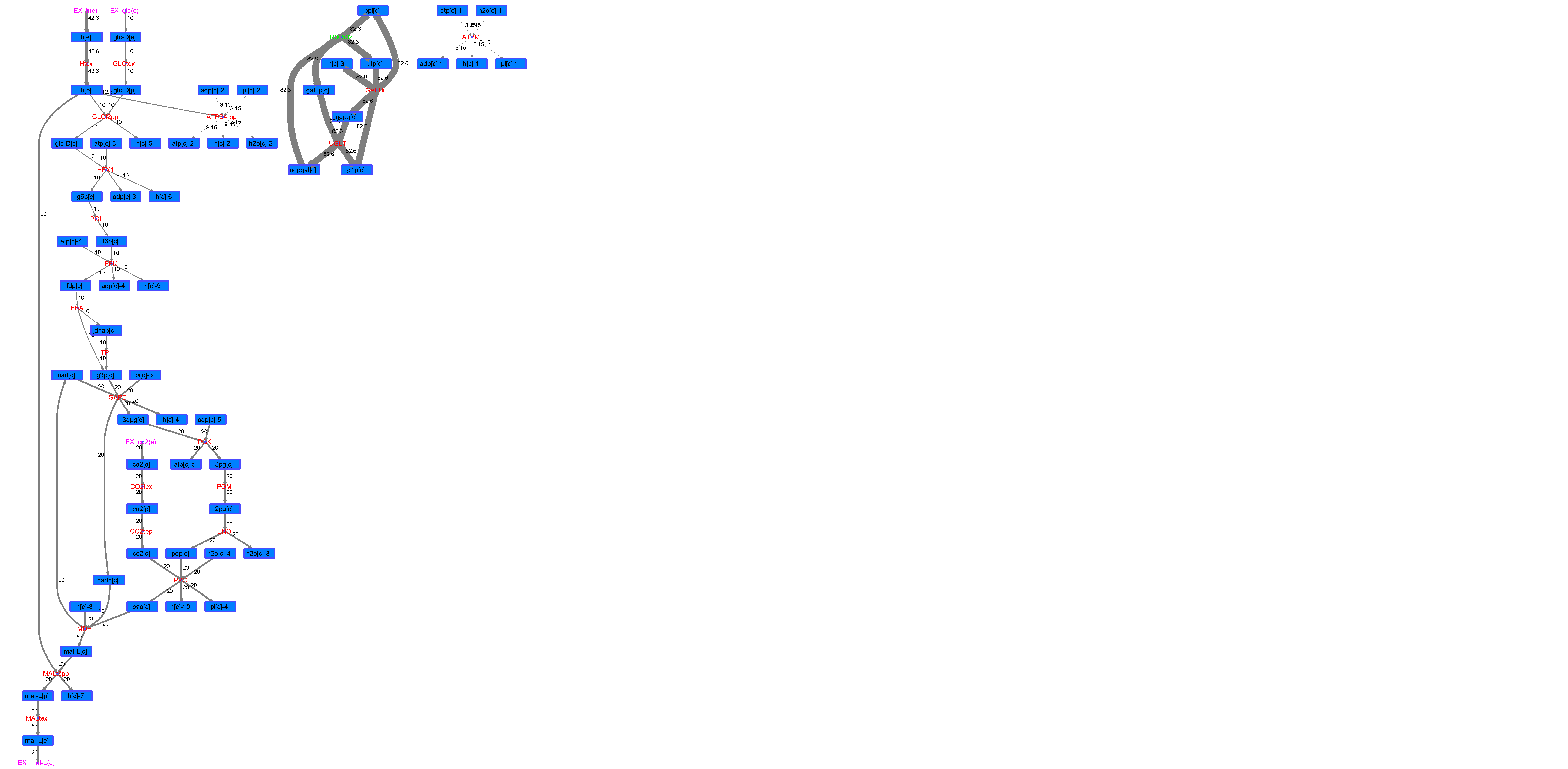

### R00502_L-Phe.png

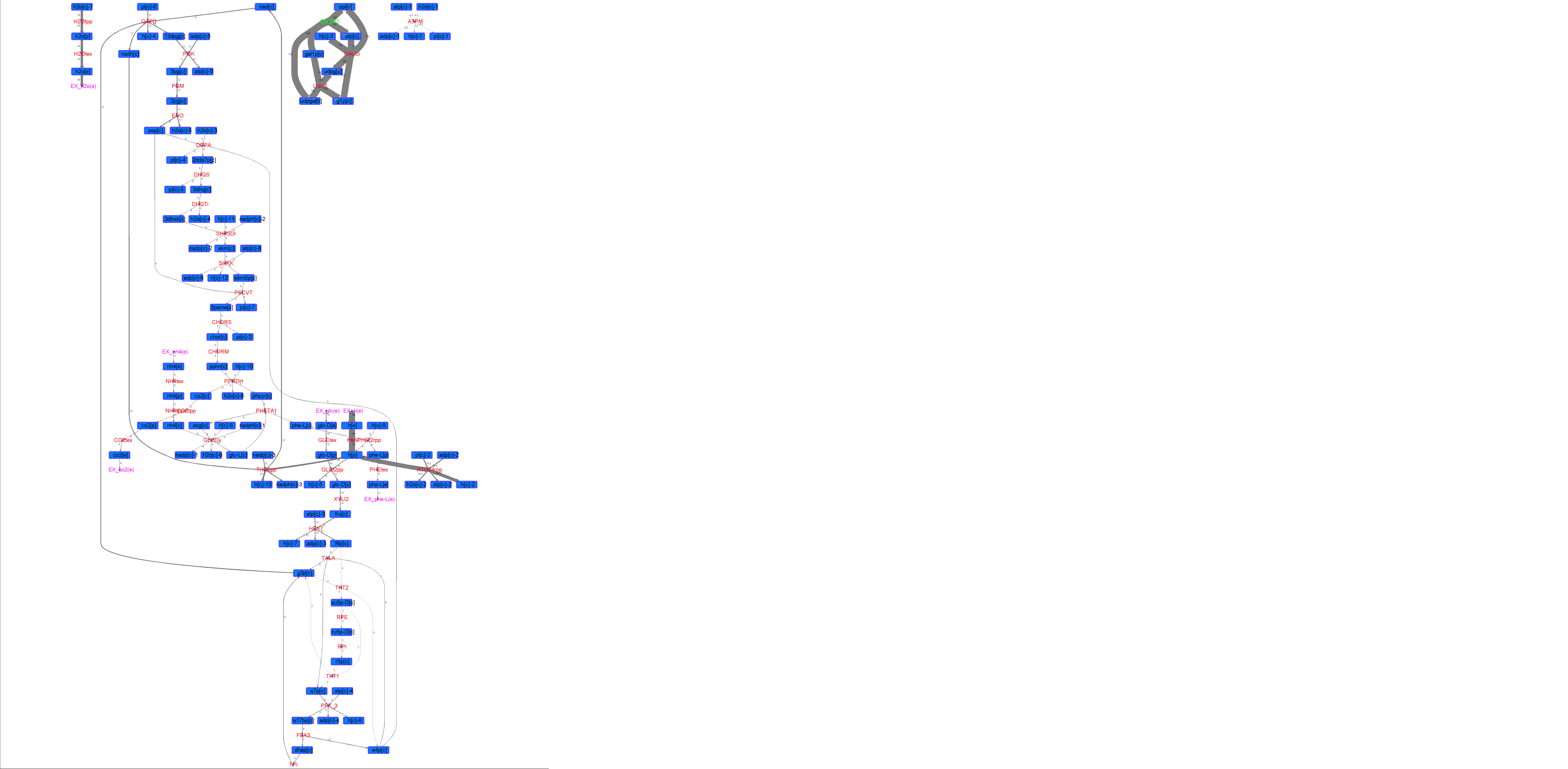

### R00502_L-Try.png

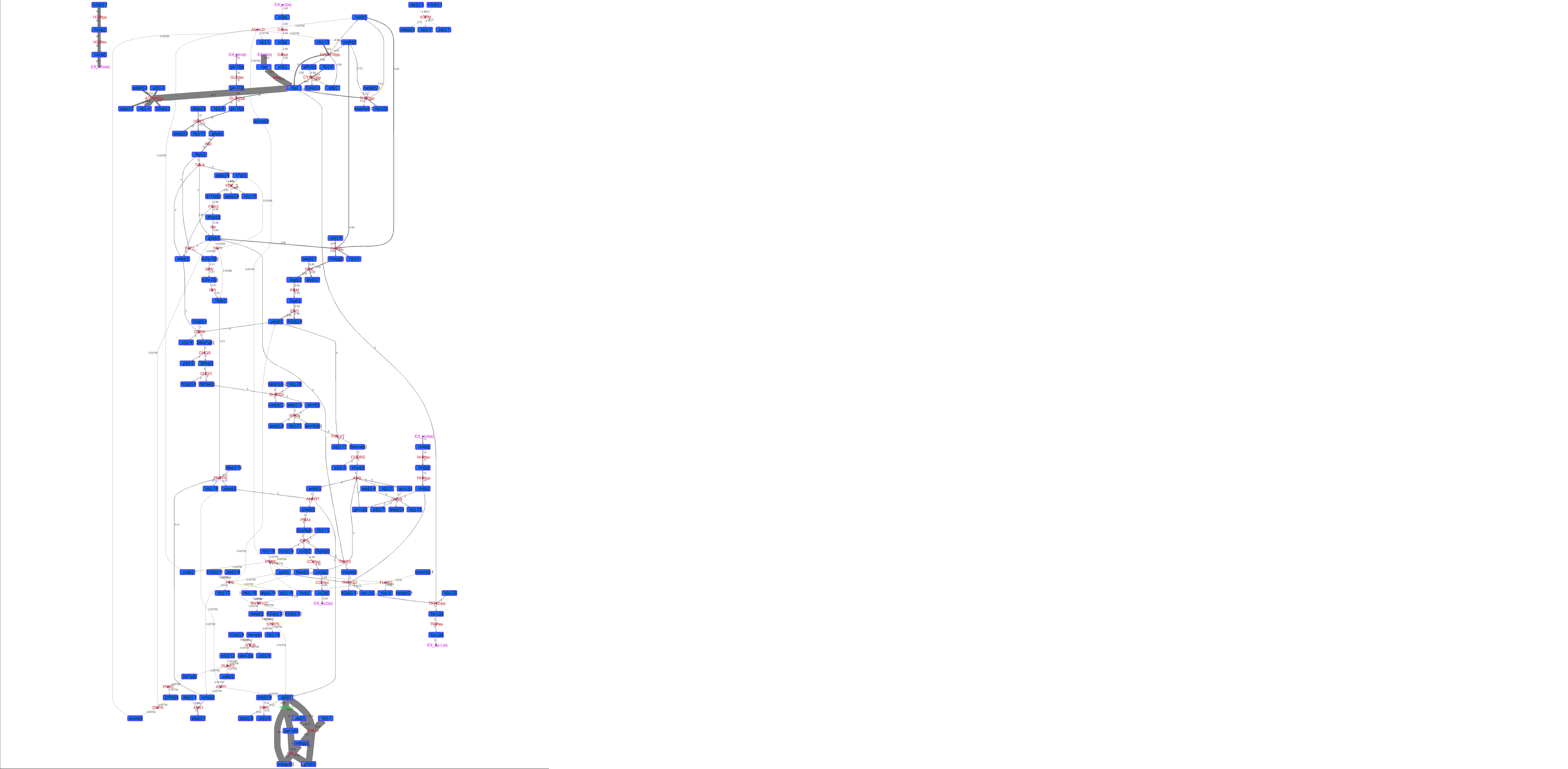

### R00502_Lys.png

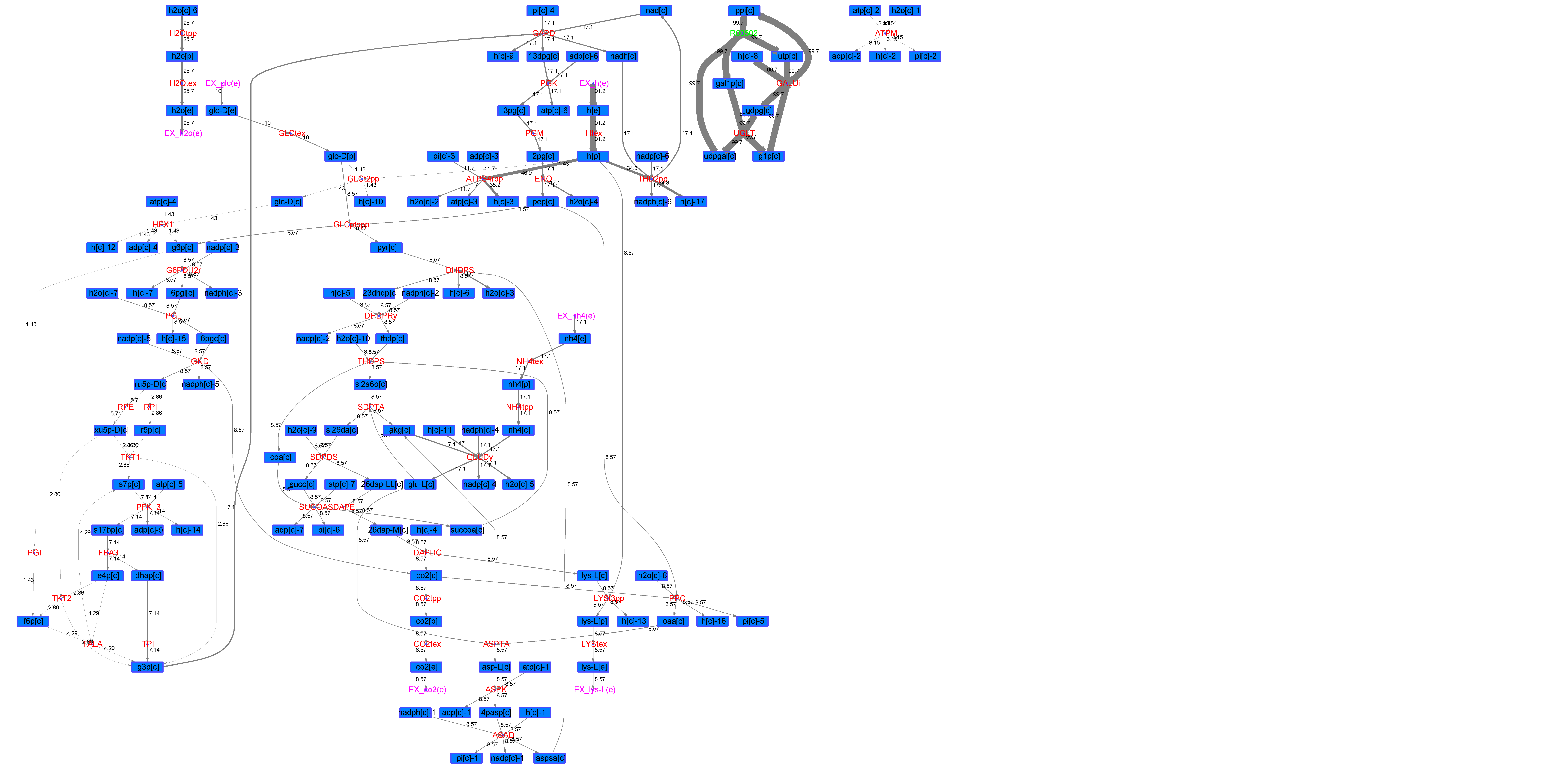

### R00502_Suc.png

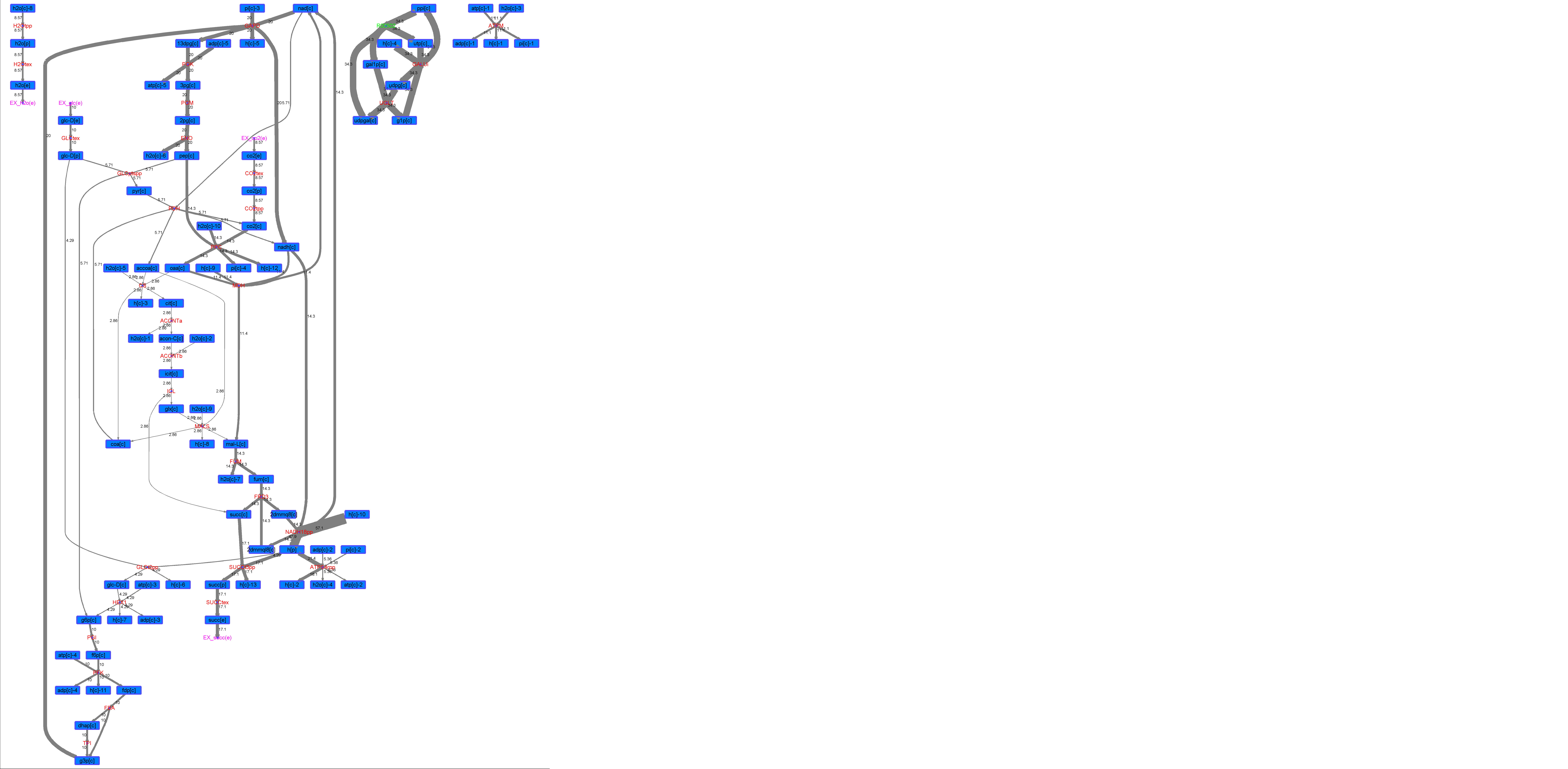

### R00502_Thr.png

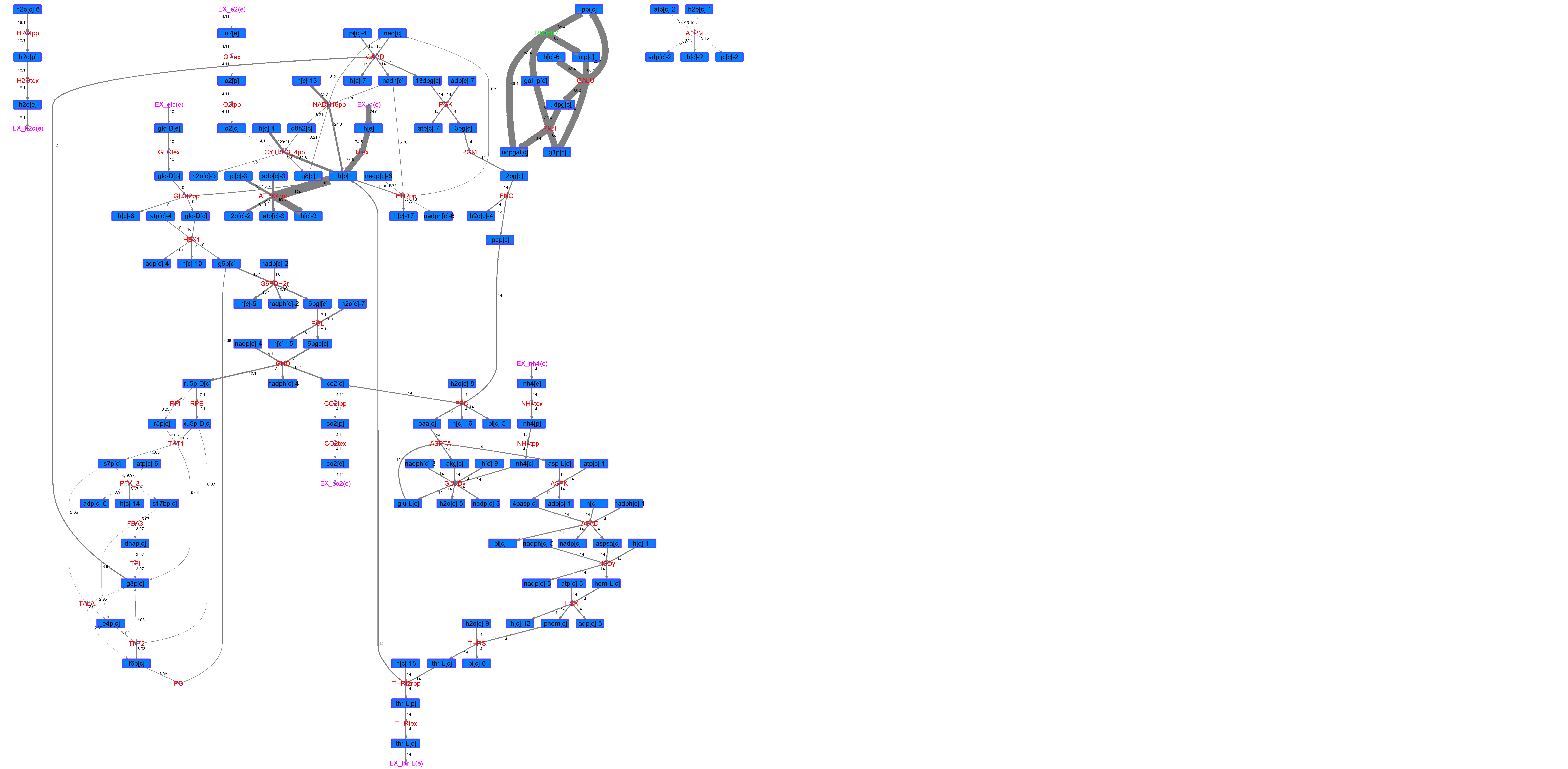

### R00519_Ace.png

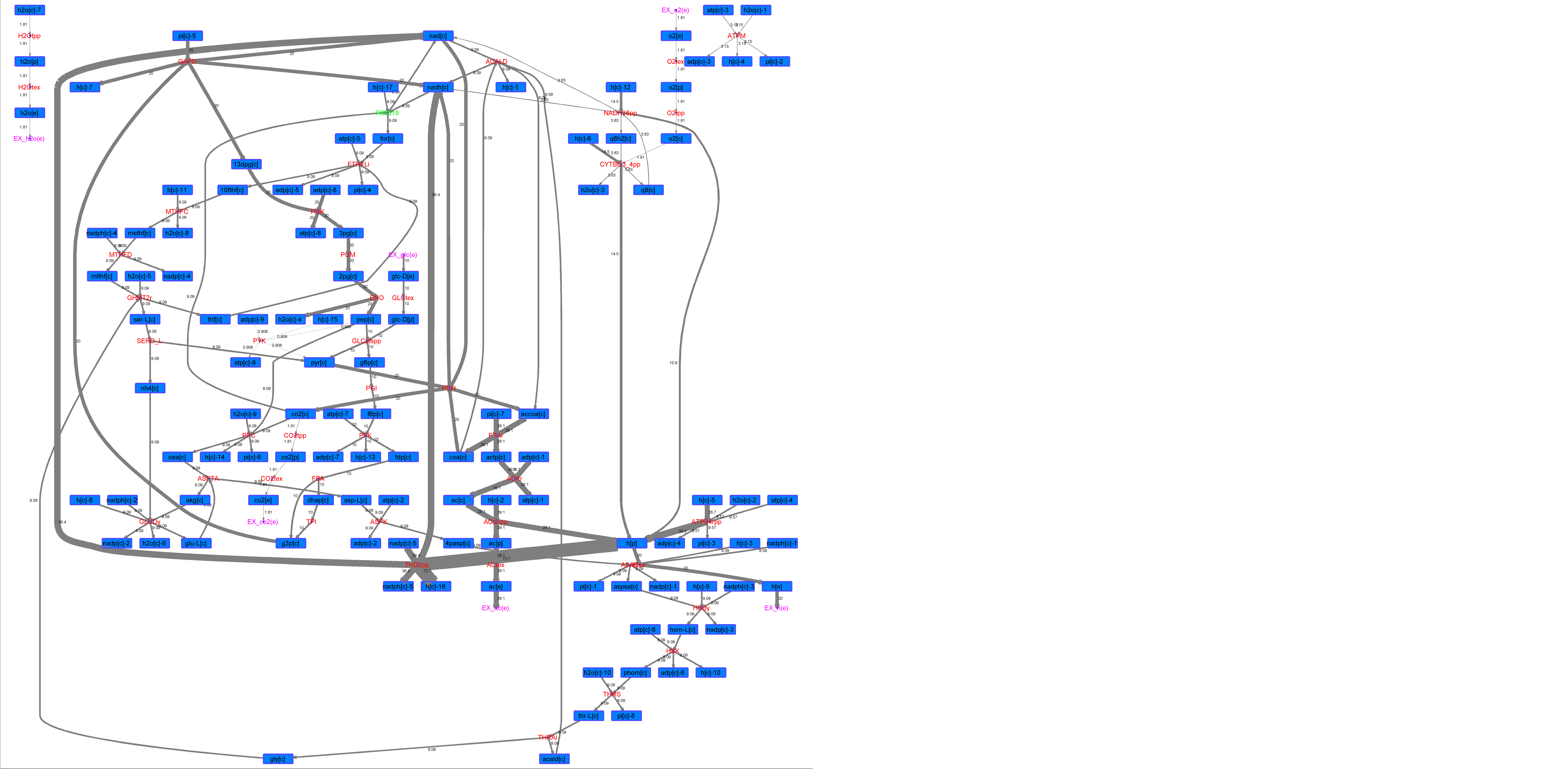

### R00519_For.png

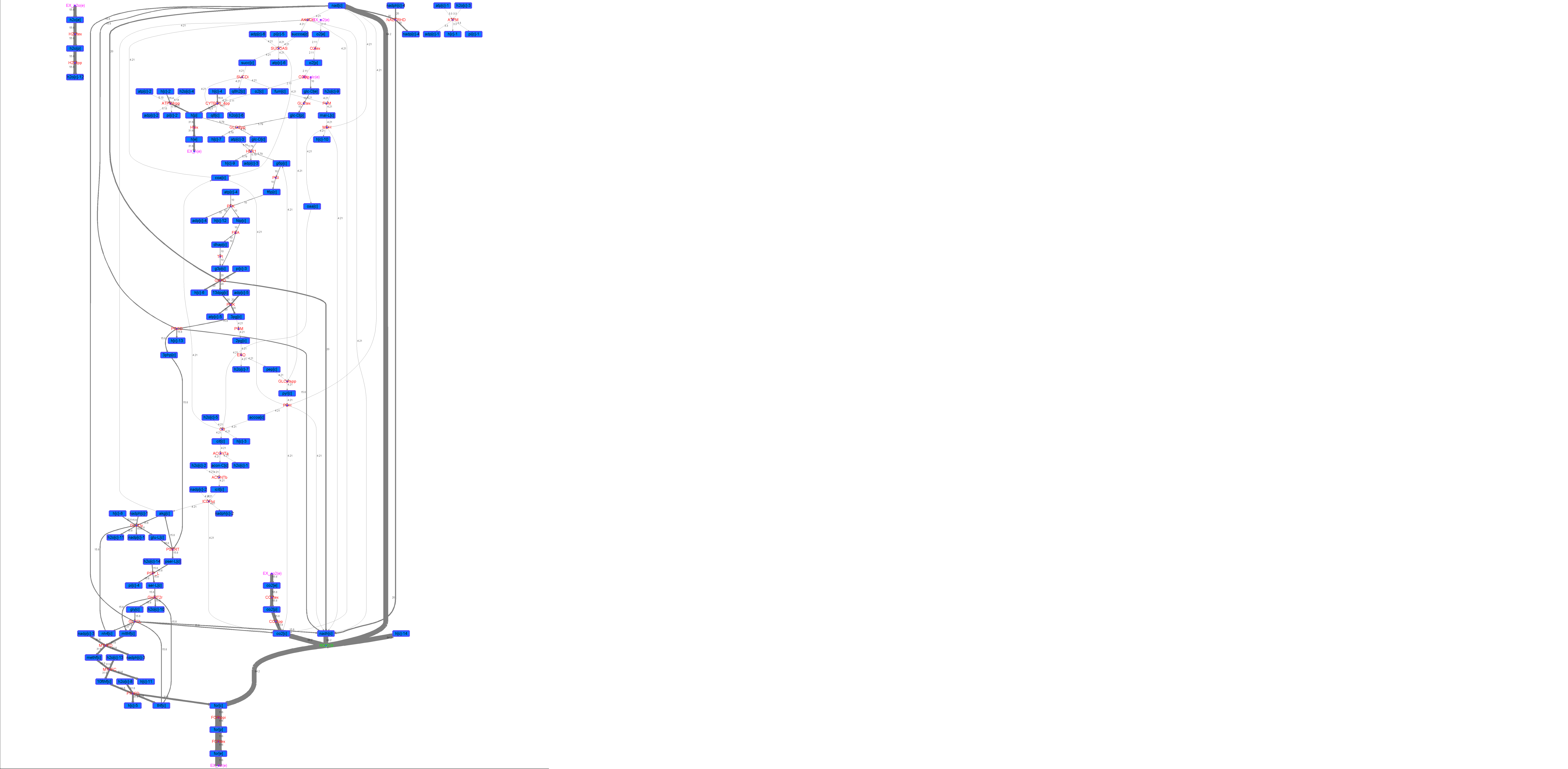

### R00519_Glu.png

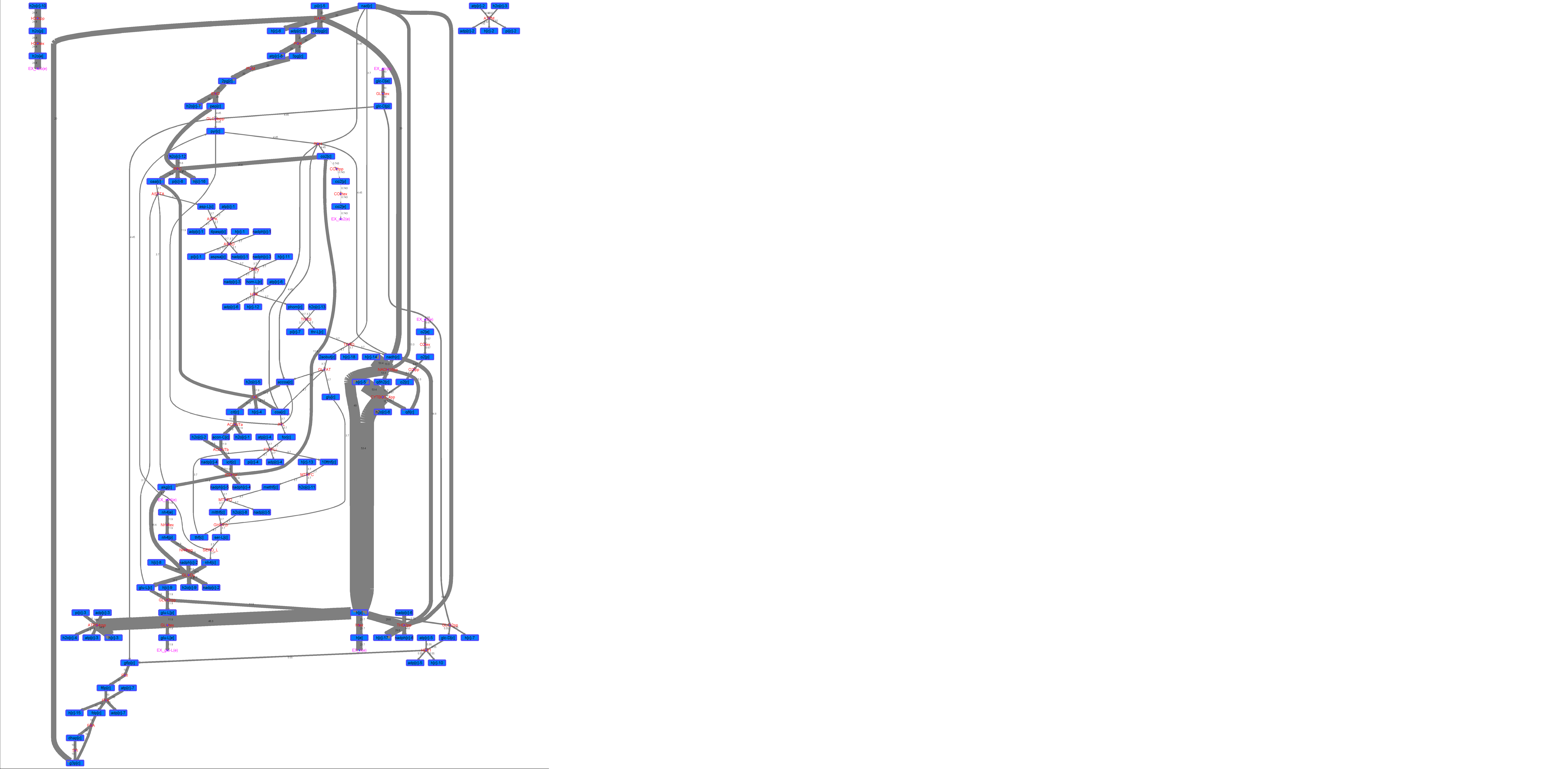

### R00519_L-ala.png

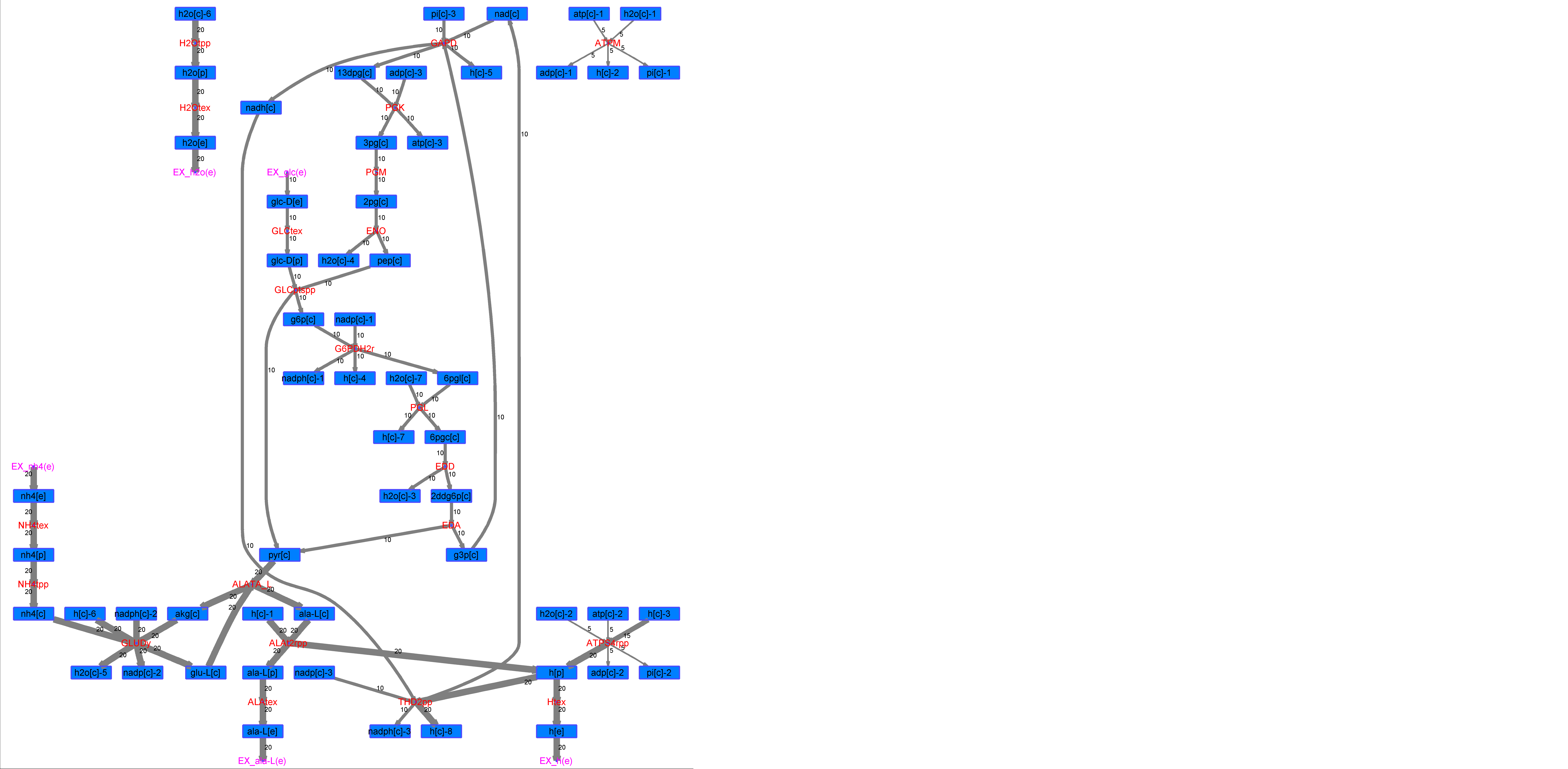

### R00519_L-Mal.png

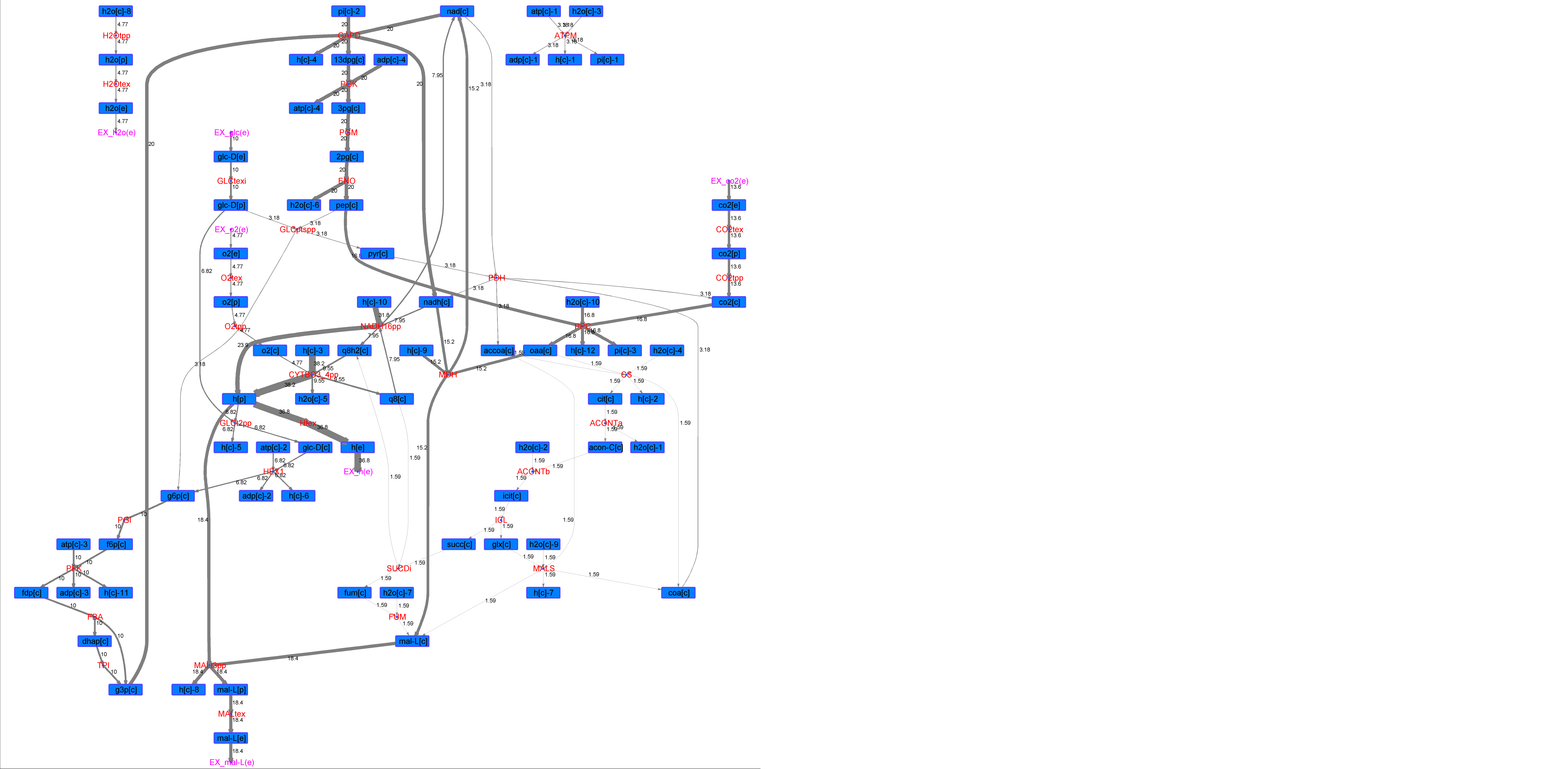

### R00519_L-Phe.png

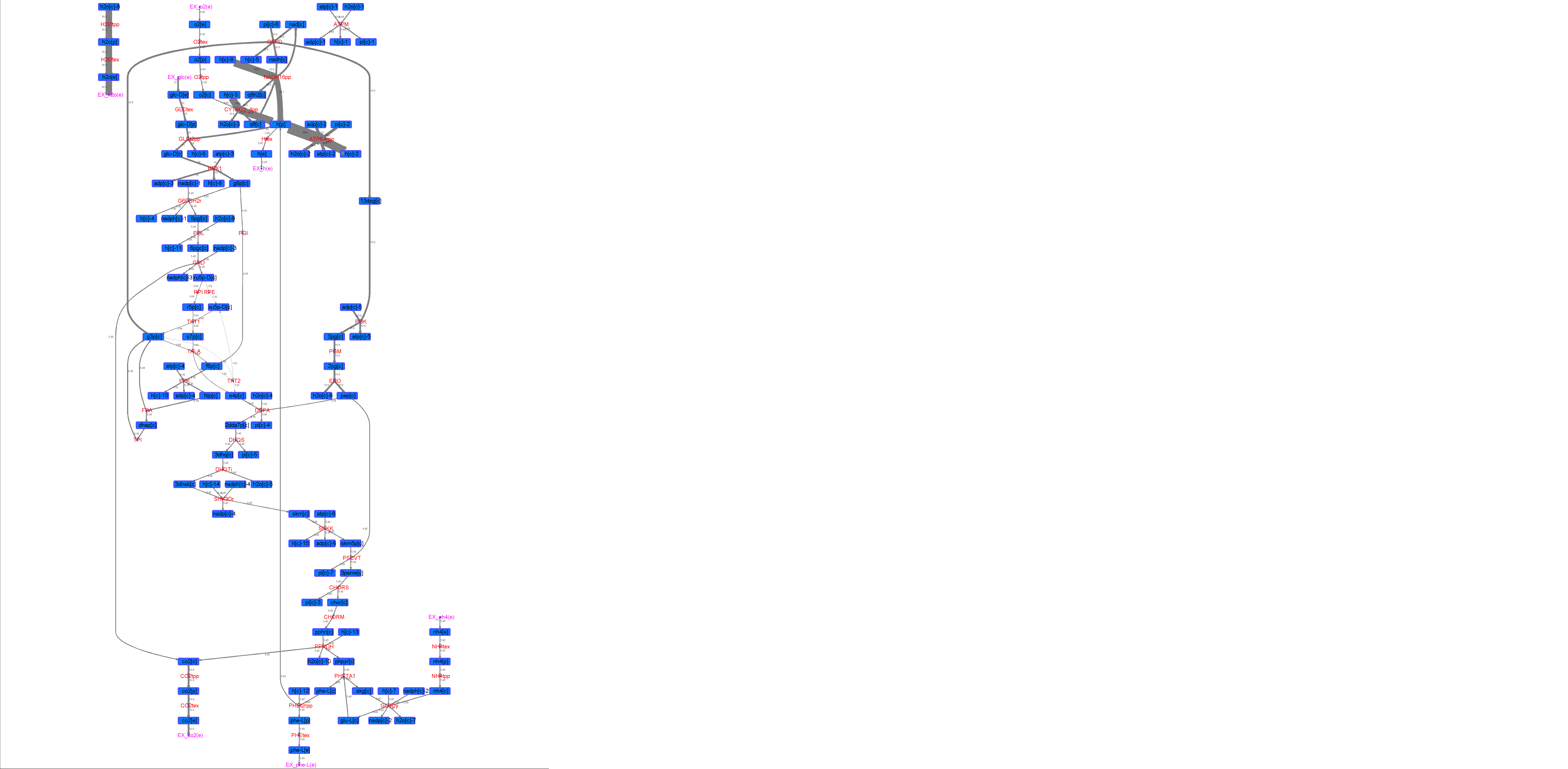

### R00519_L-Try.png

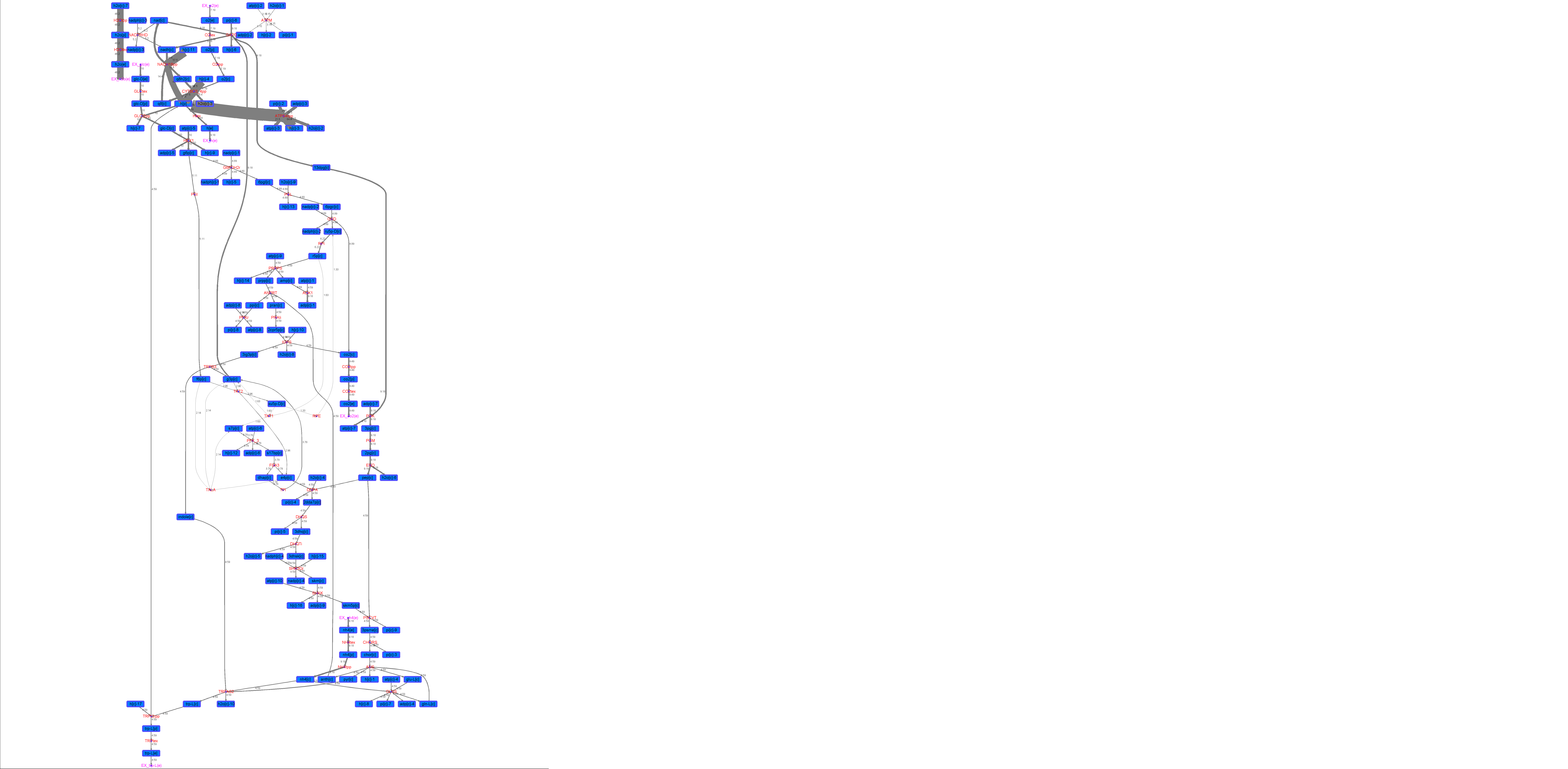

### R00519_Lys.png

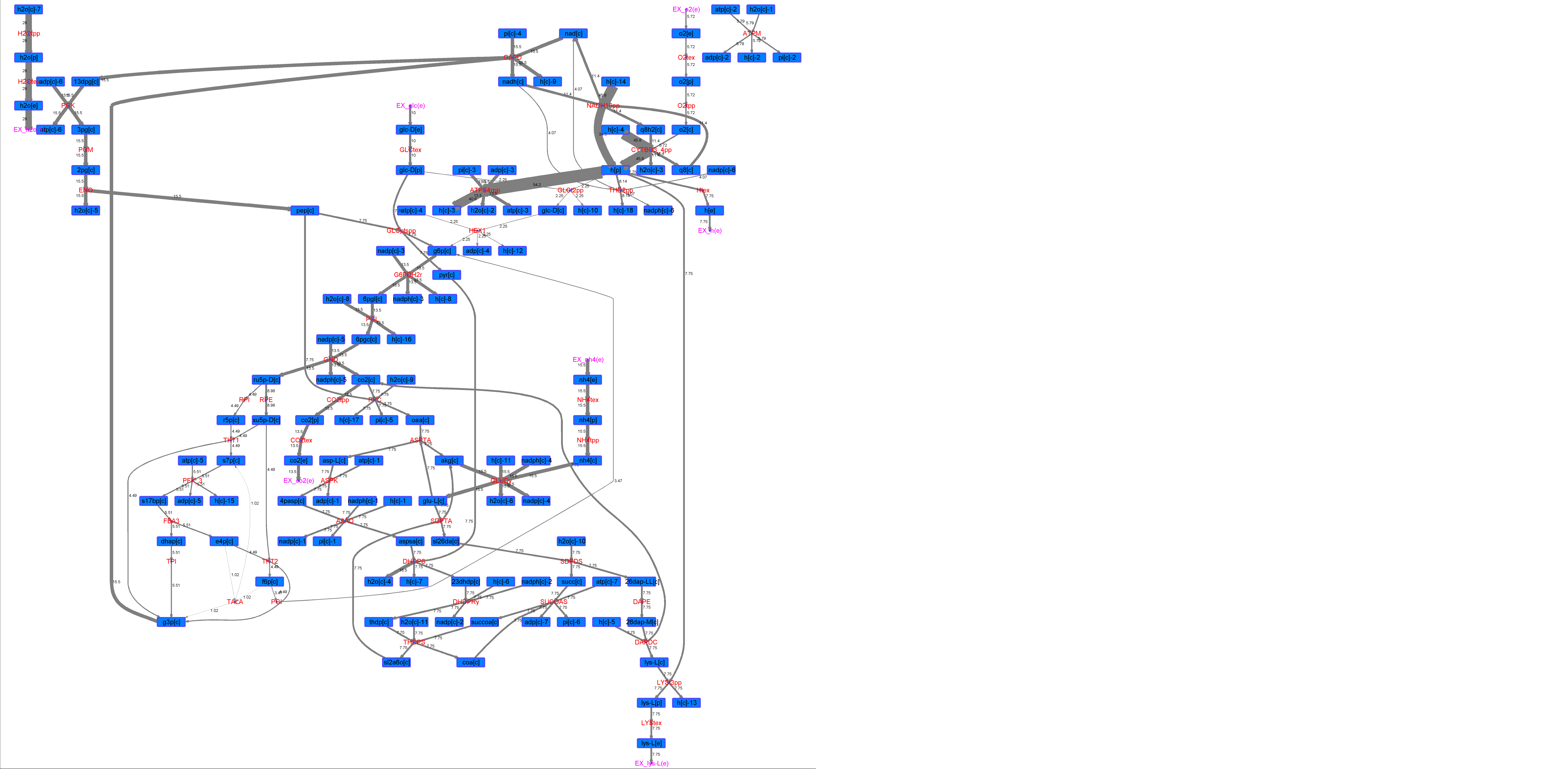

### R00519_Suc.png

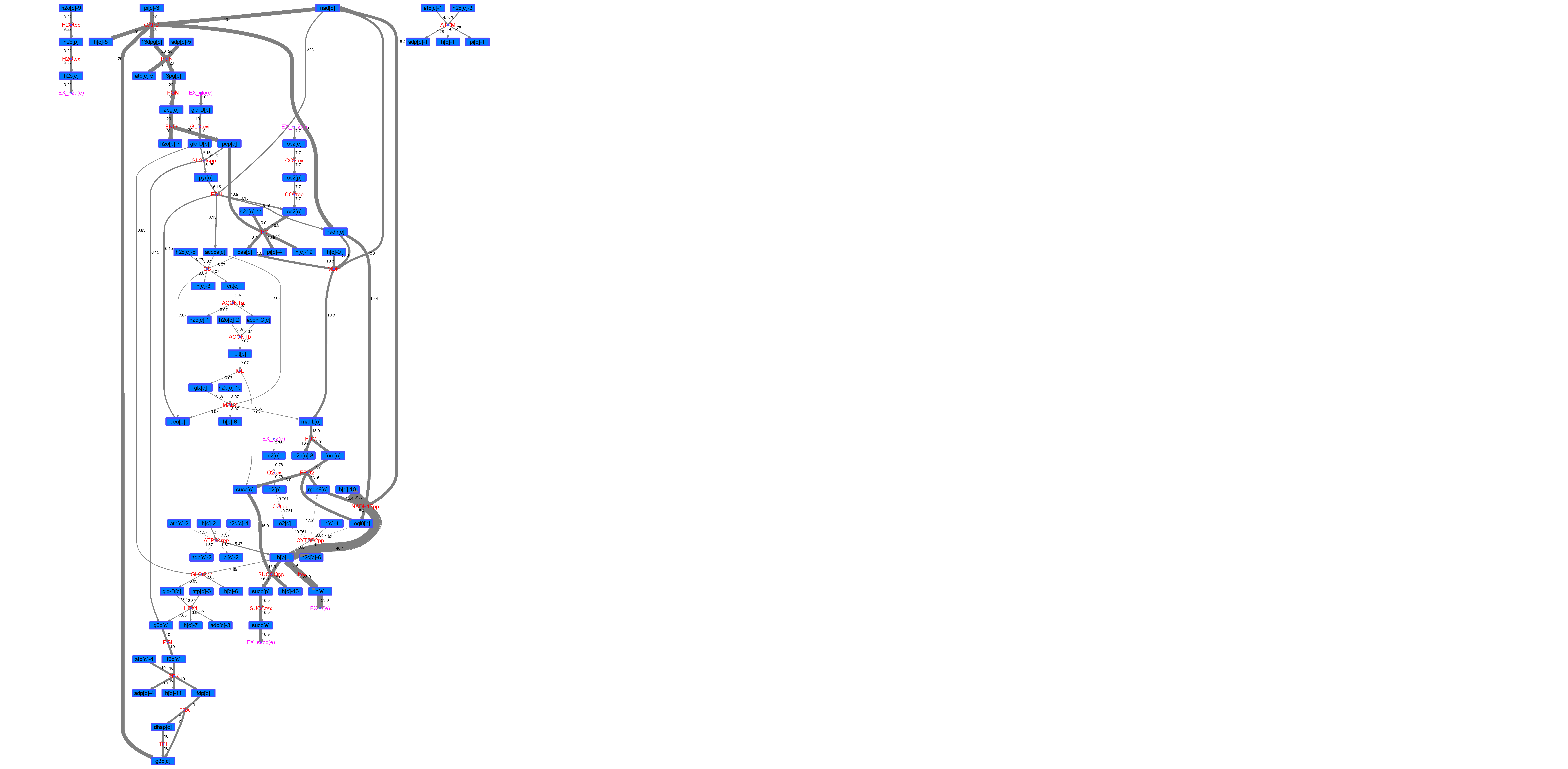

### R00519_Thr.png

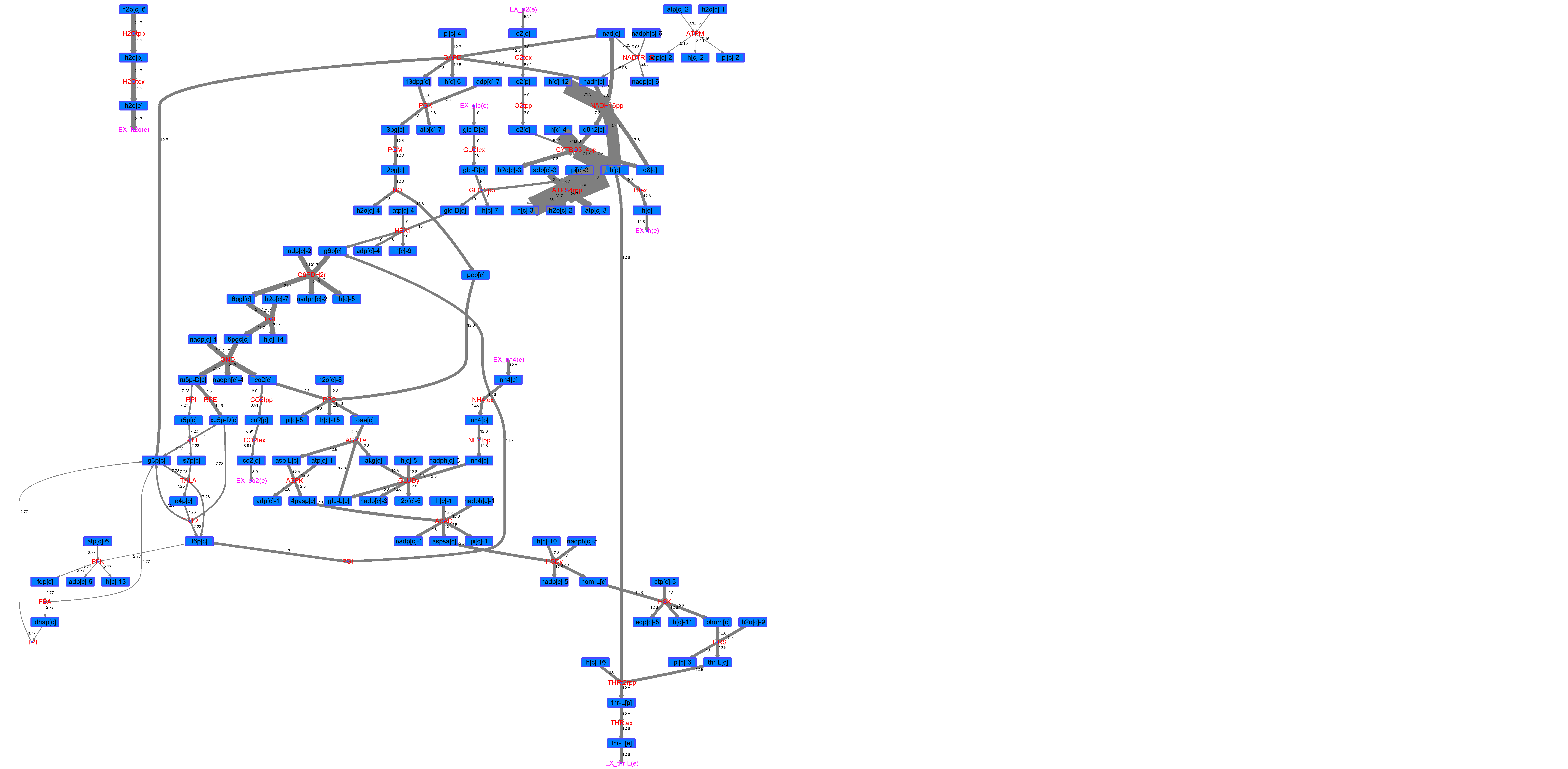

### R00522_Ace.png

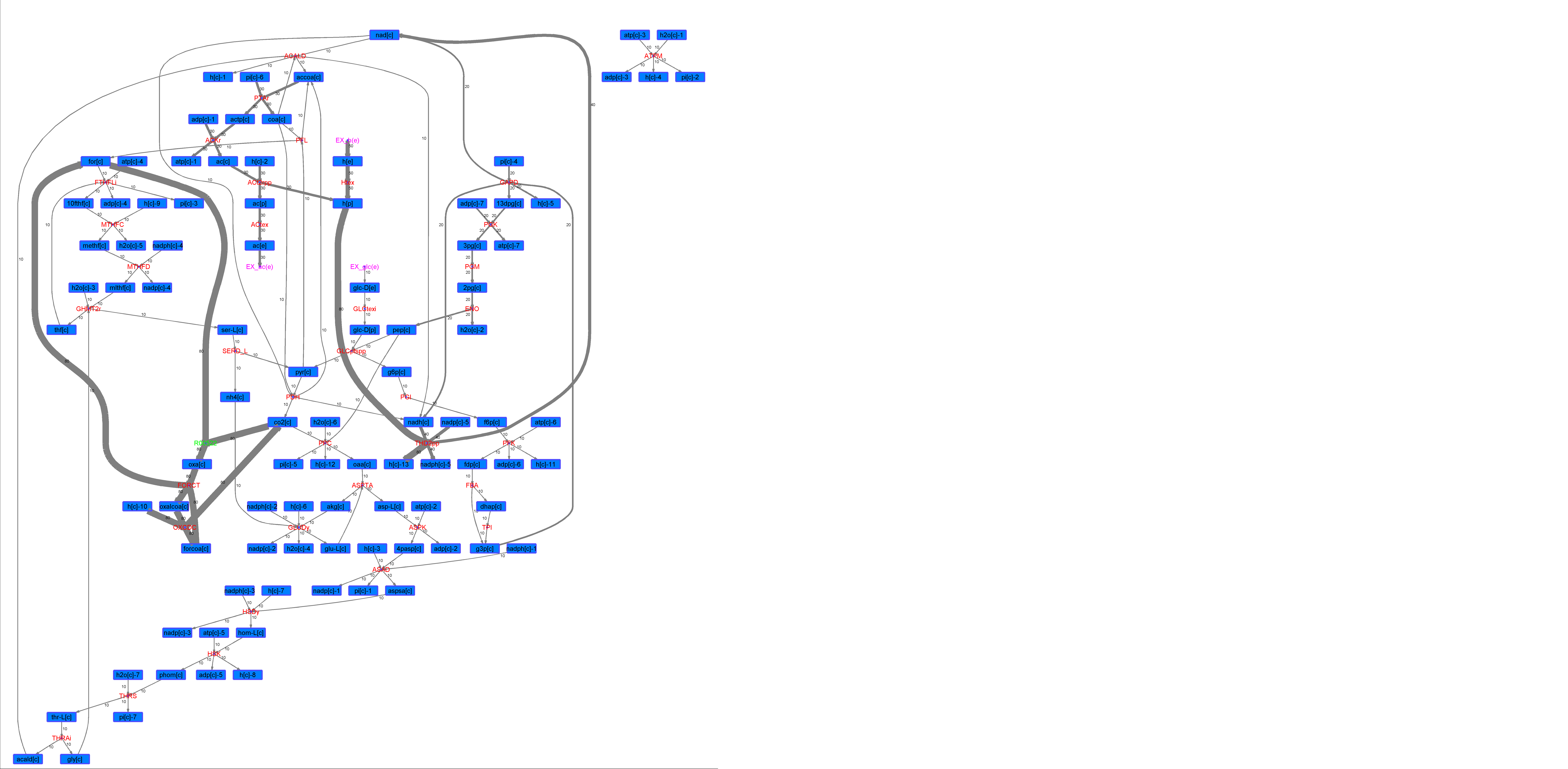

### R00522_For.png

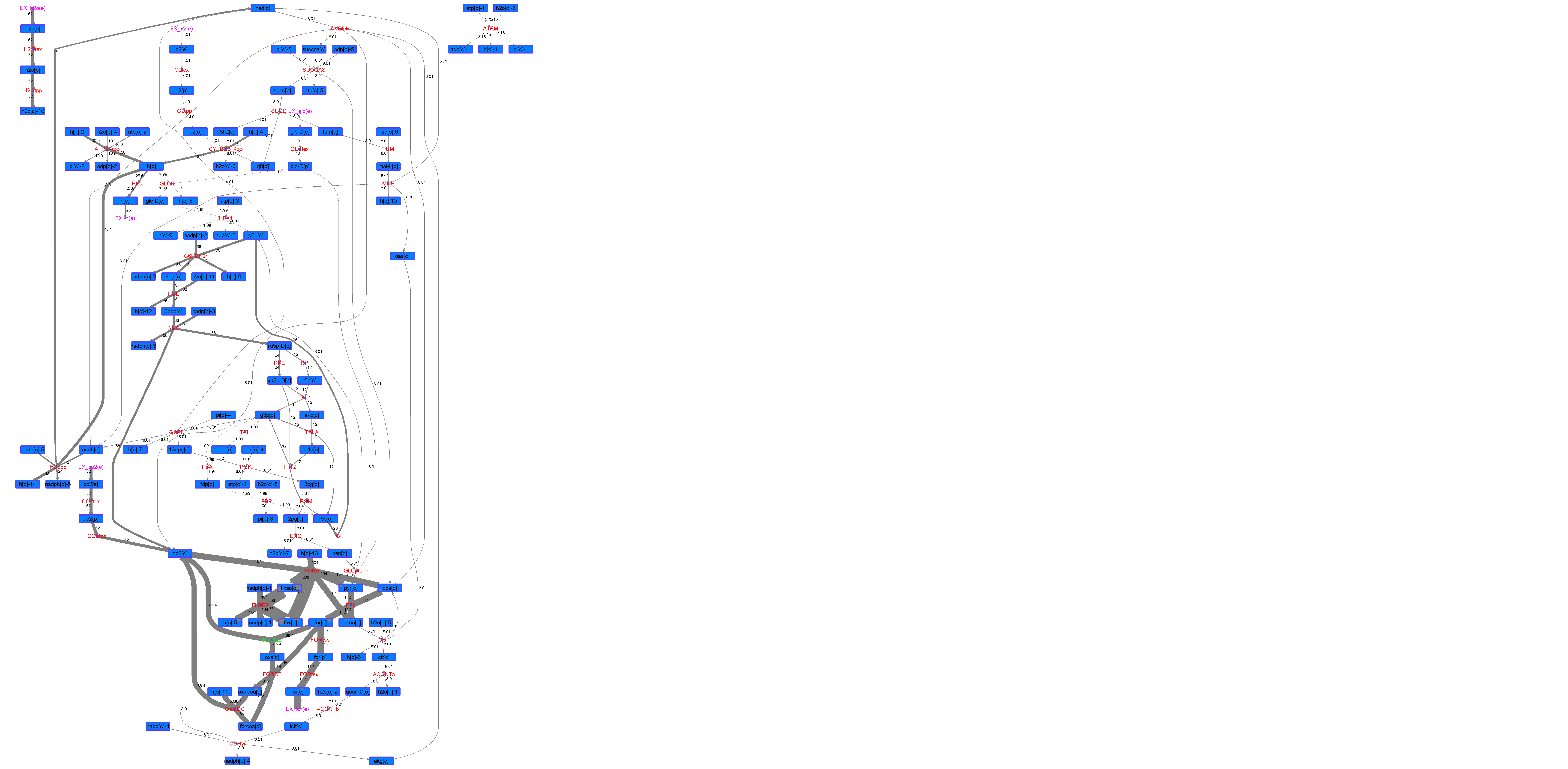

### R00522_Glu.png

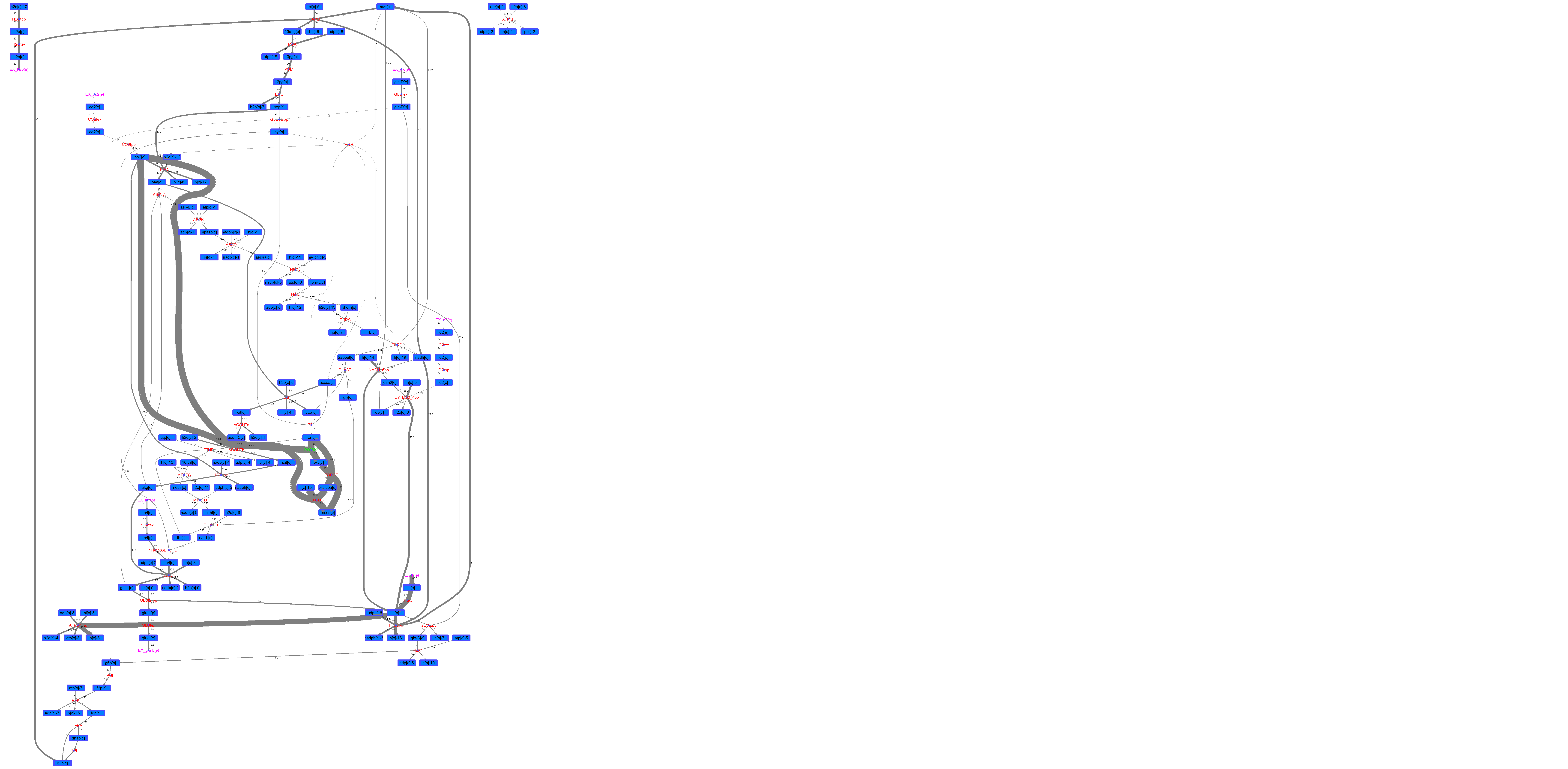

### R00522_L-ala.png

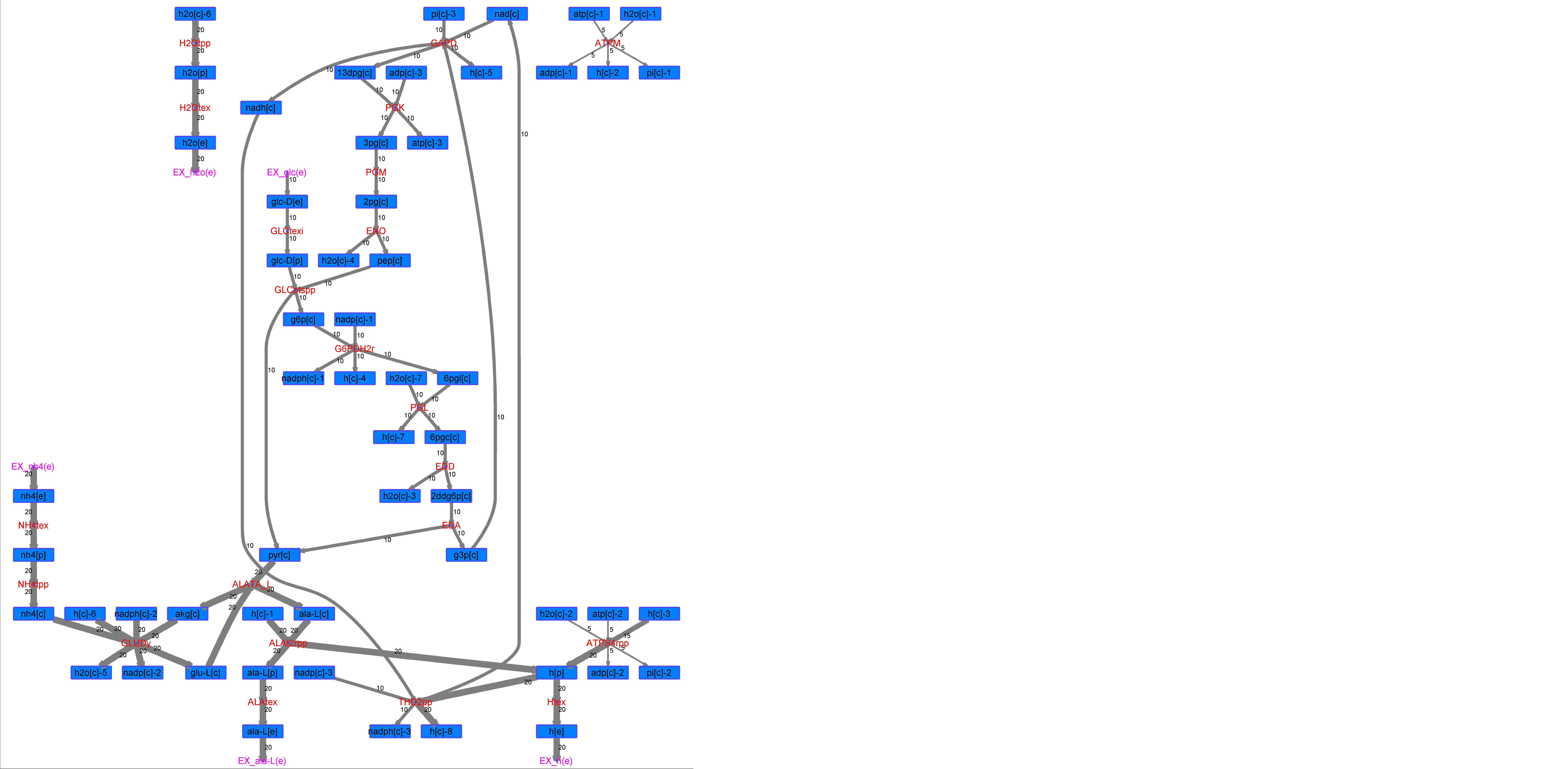

### R00522_L-Mal.png

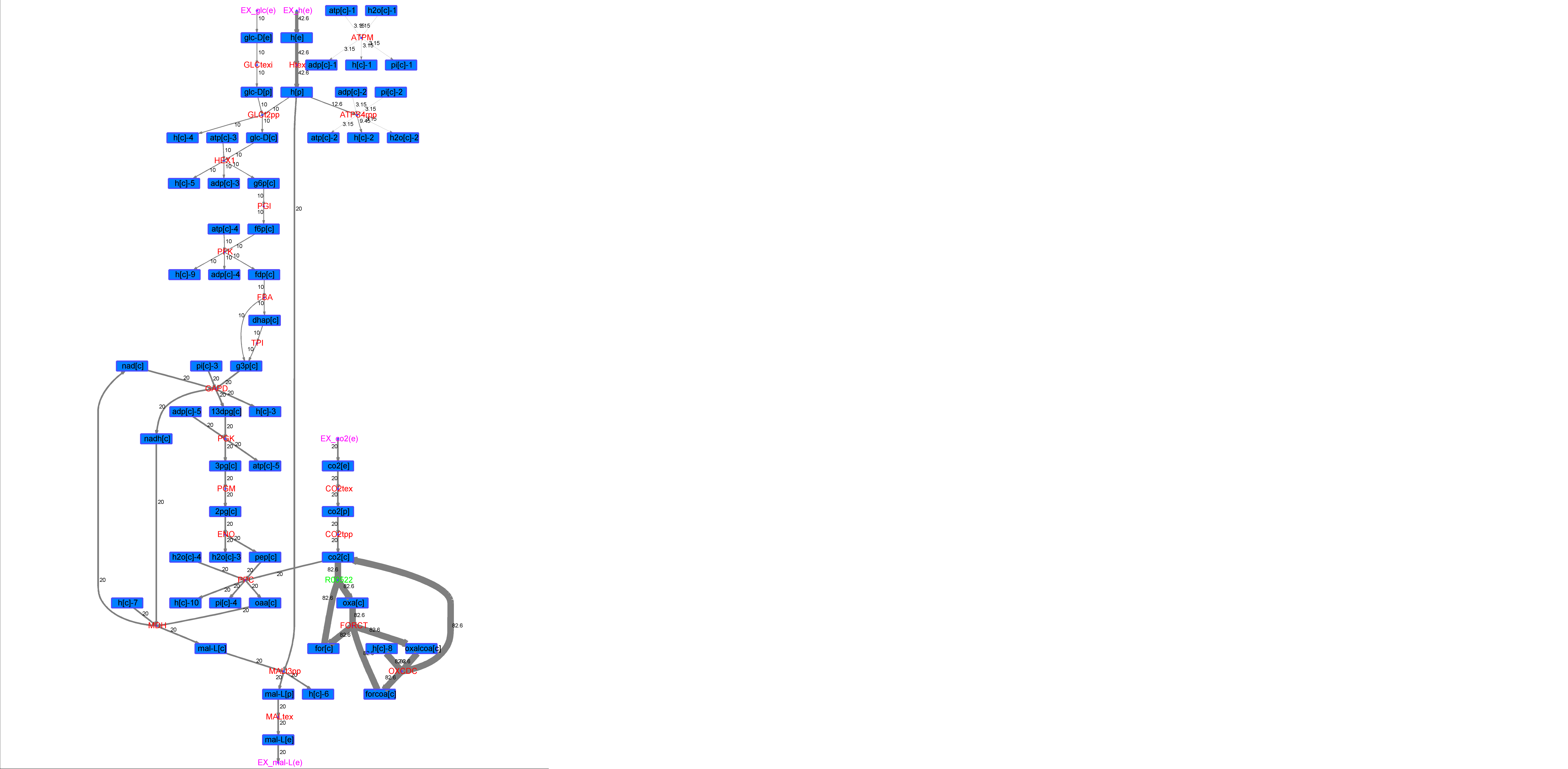

### R00522_L-Phe.png

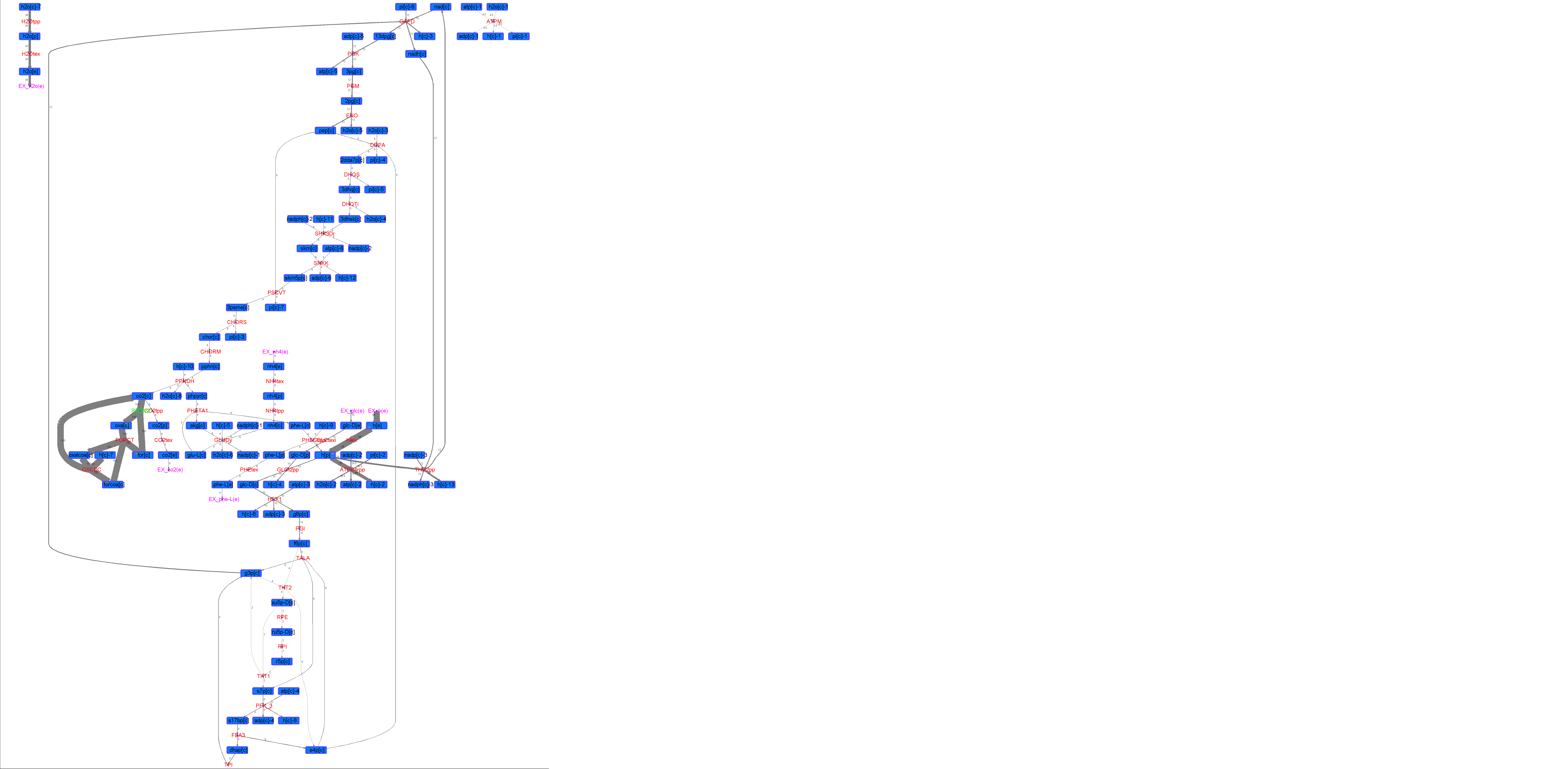

### R00522_L-Try.png

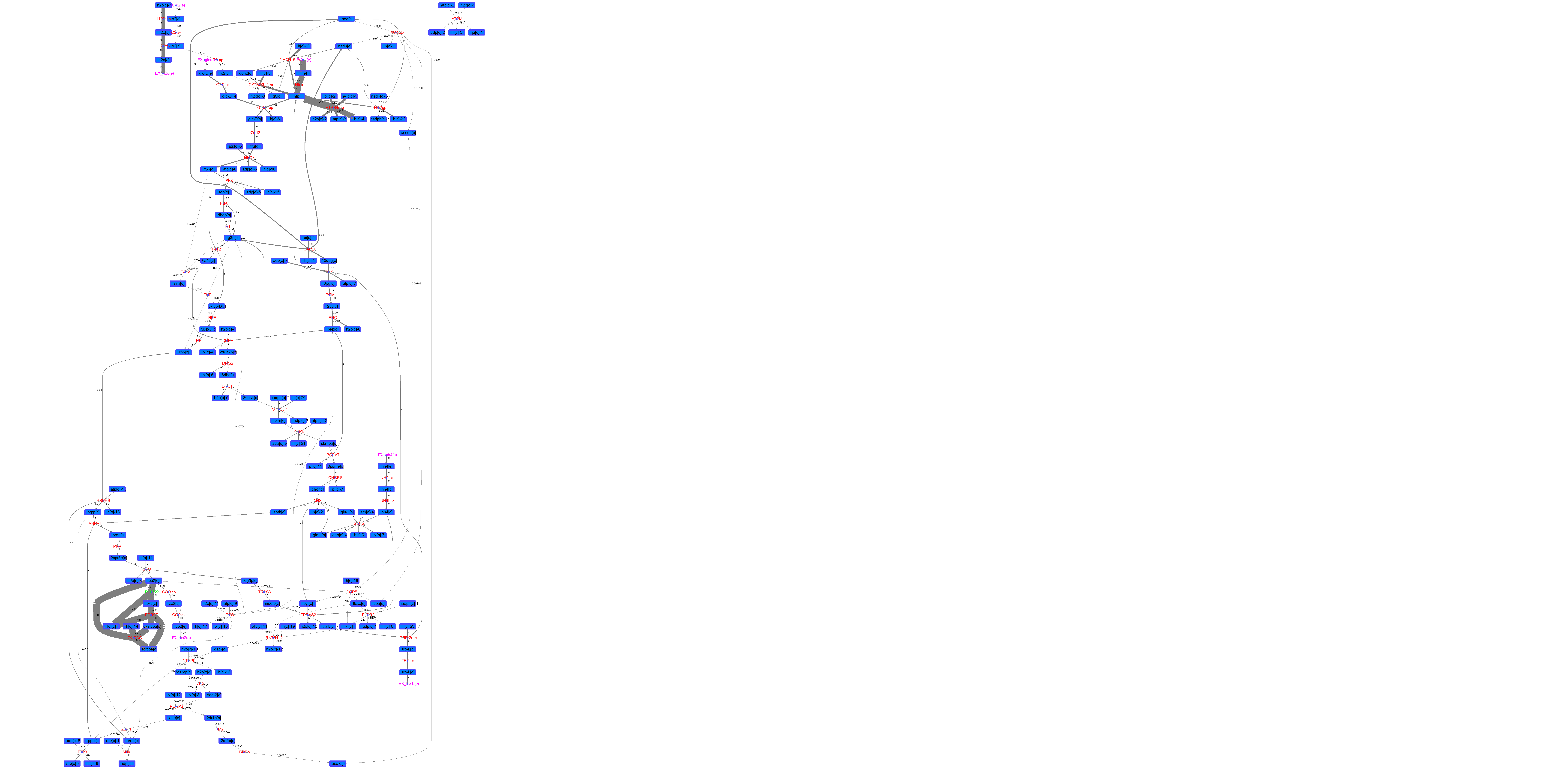

### R00522_Lys.png

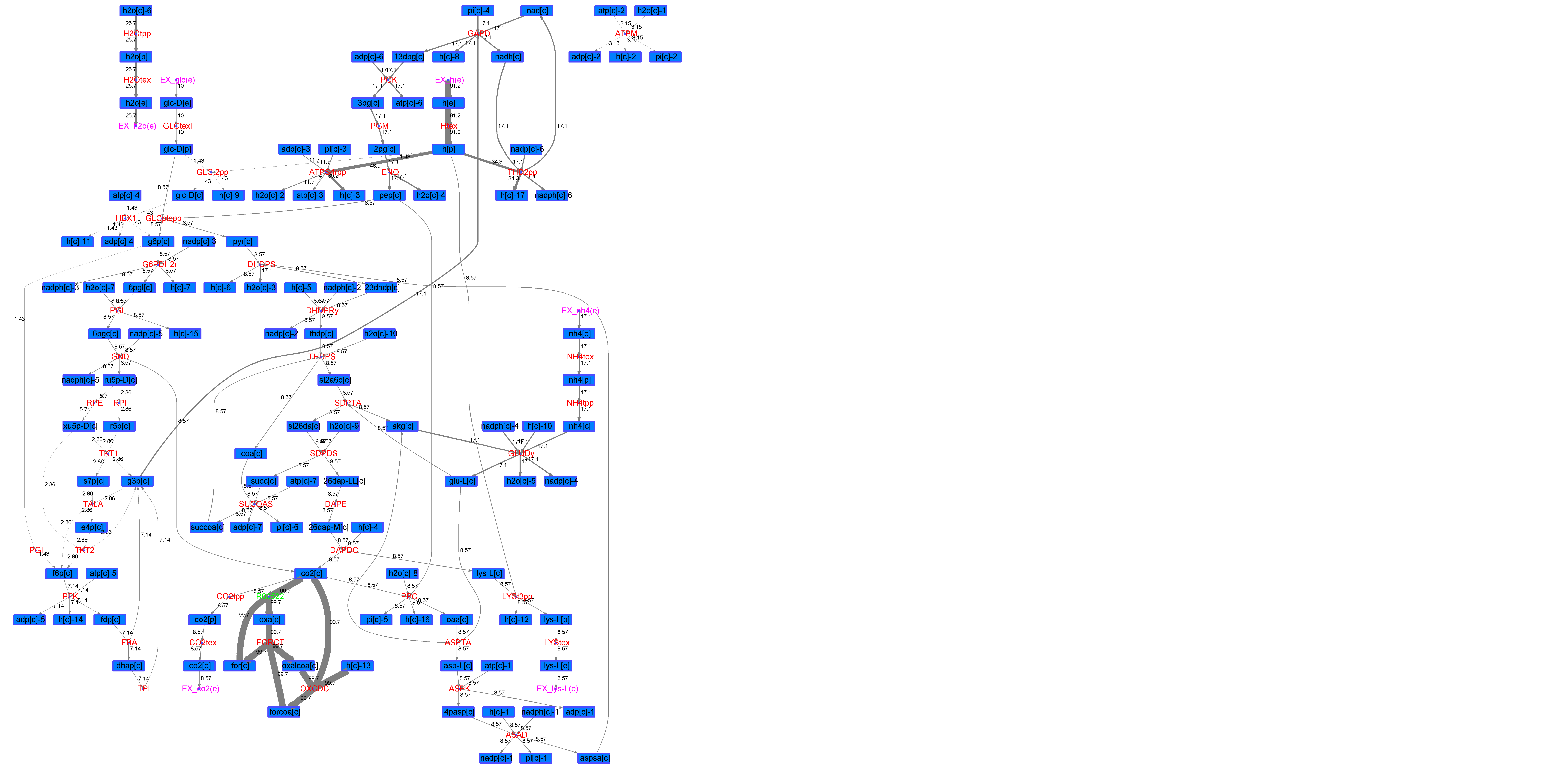

### R00522_Suc.png

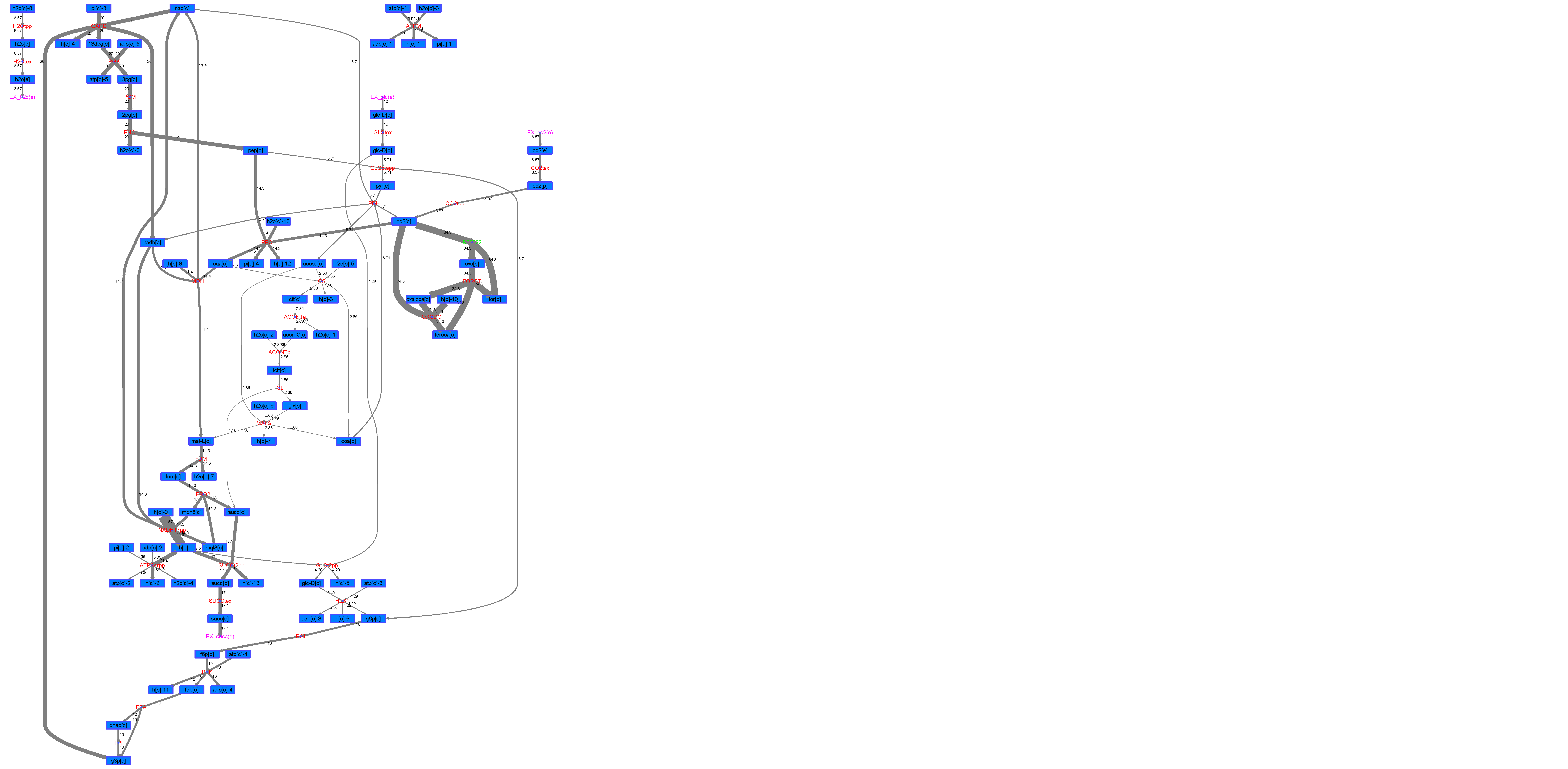

### R00522_Thr.png

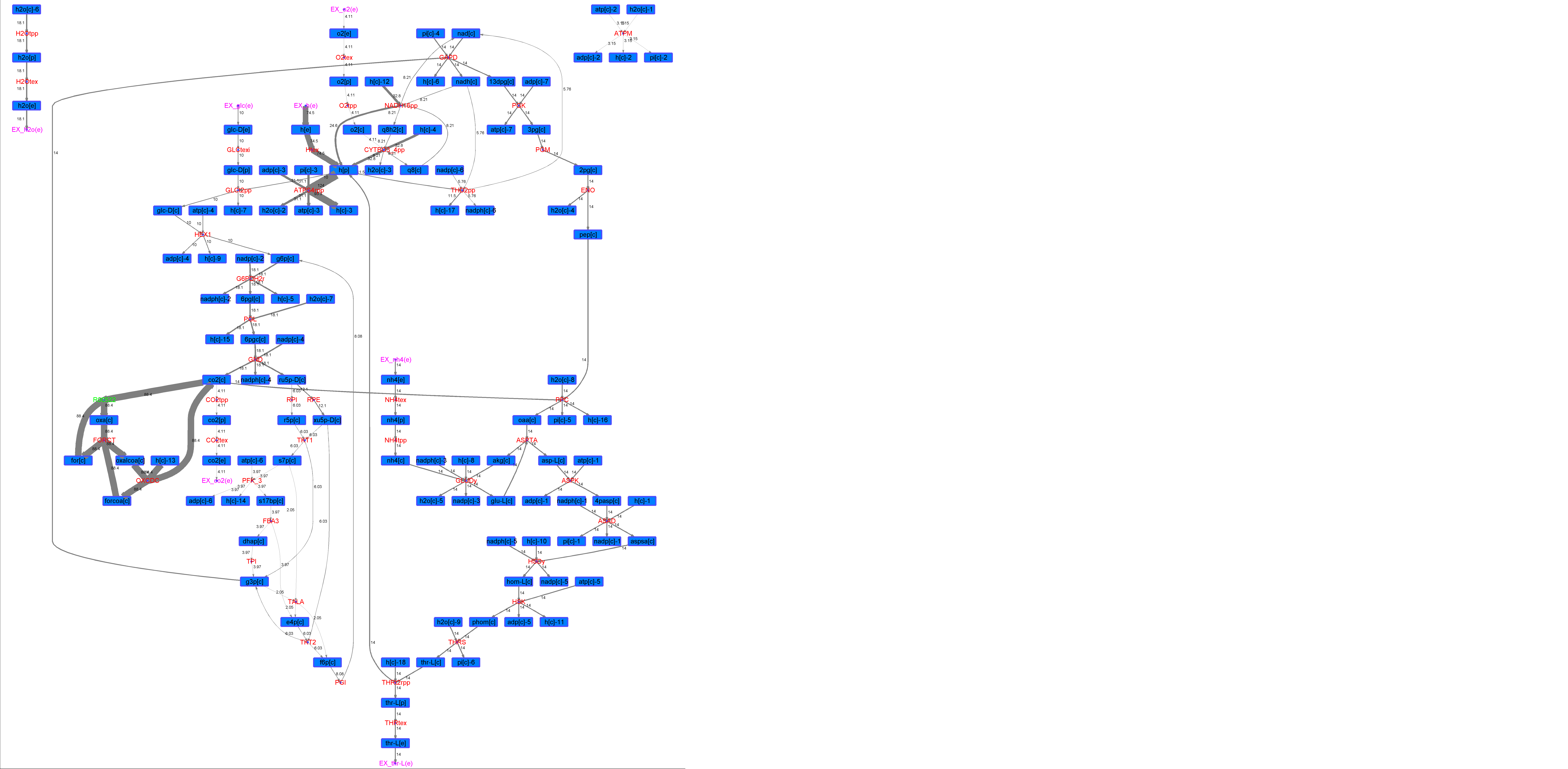

### R00572_Ace.png

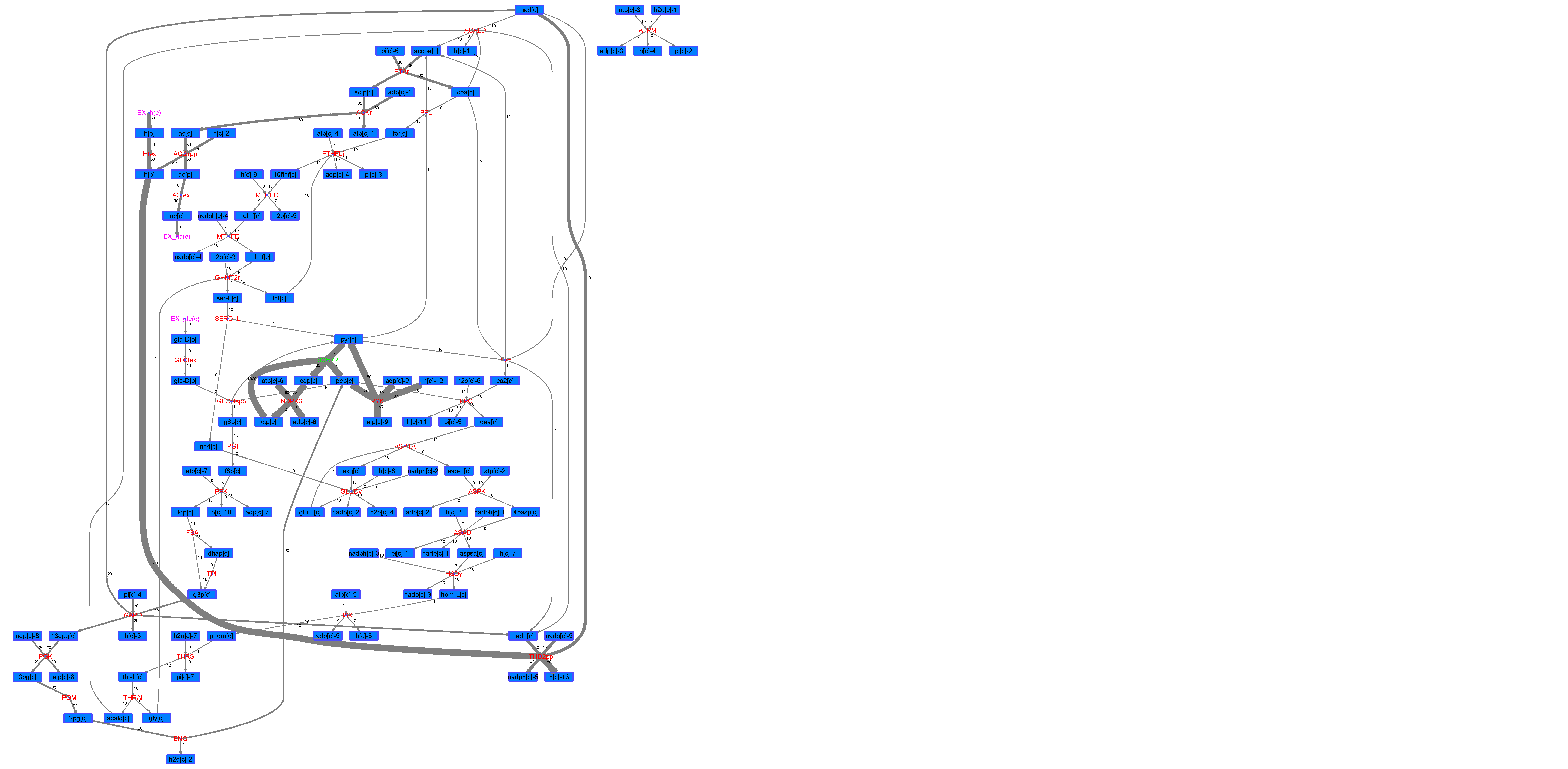

### R00572_For.png

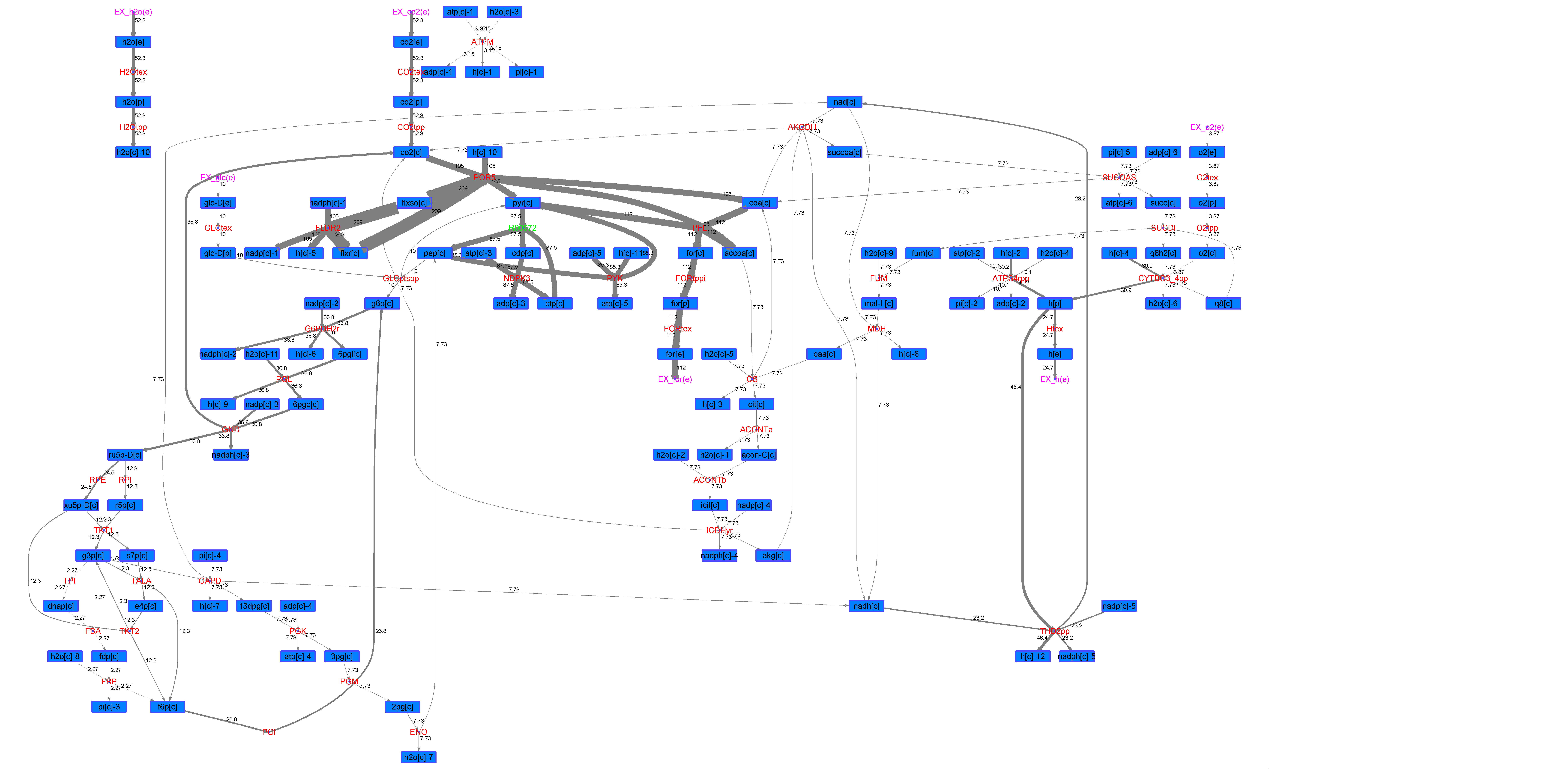

### R00572_Glu.png

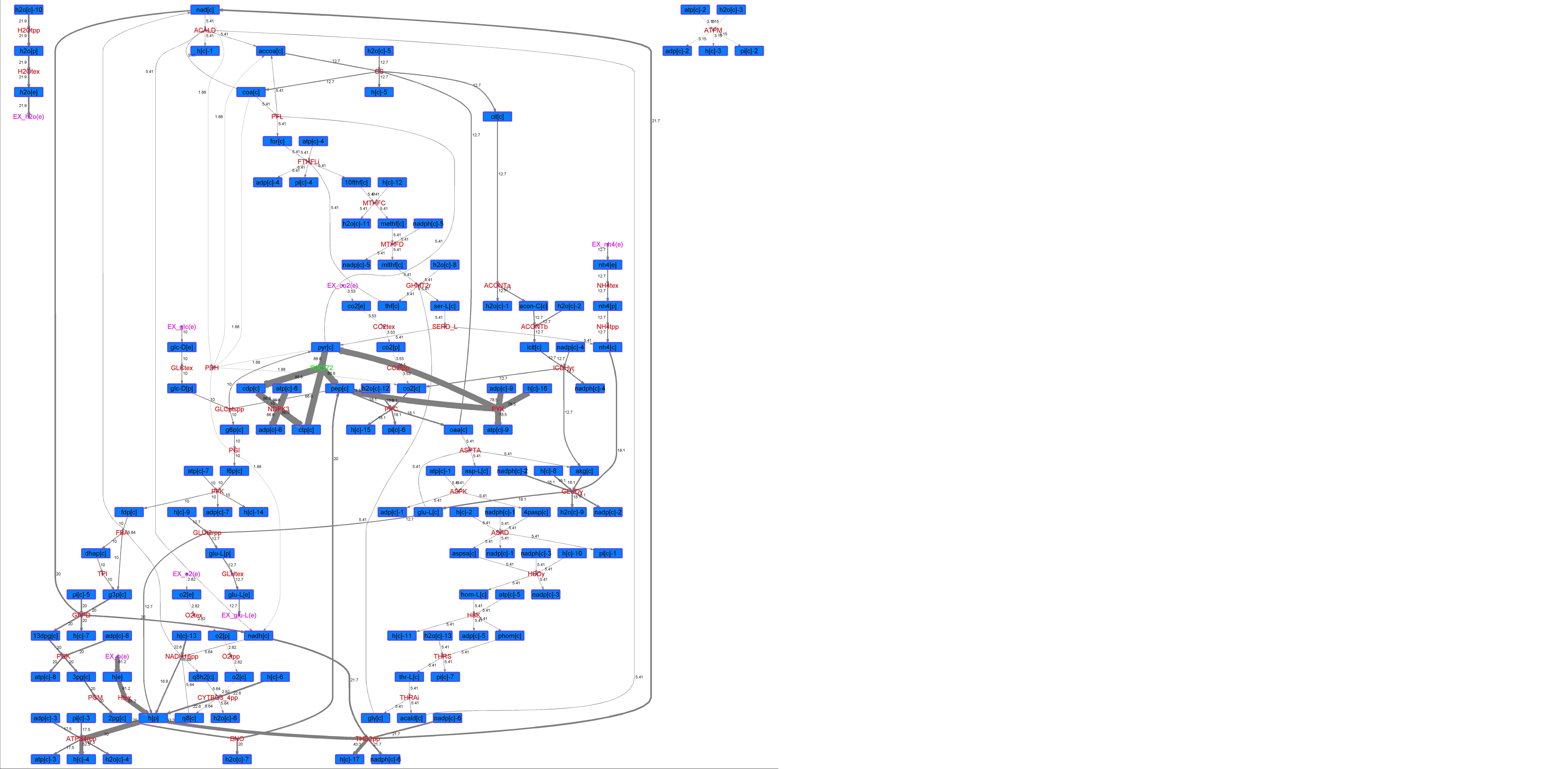

### R00572_L-ala.png

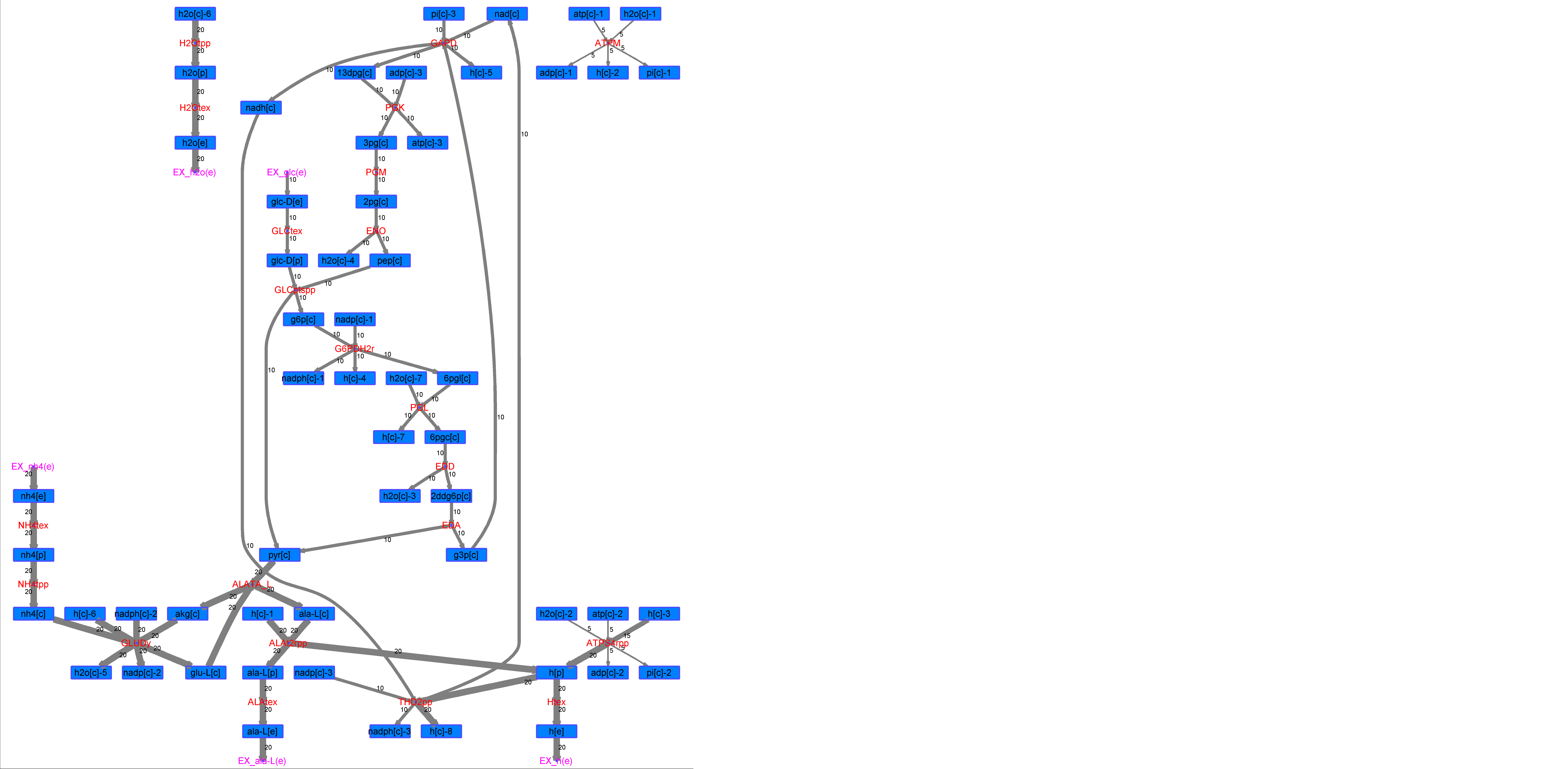

### R00572_L-Mal.png

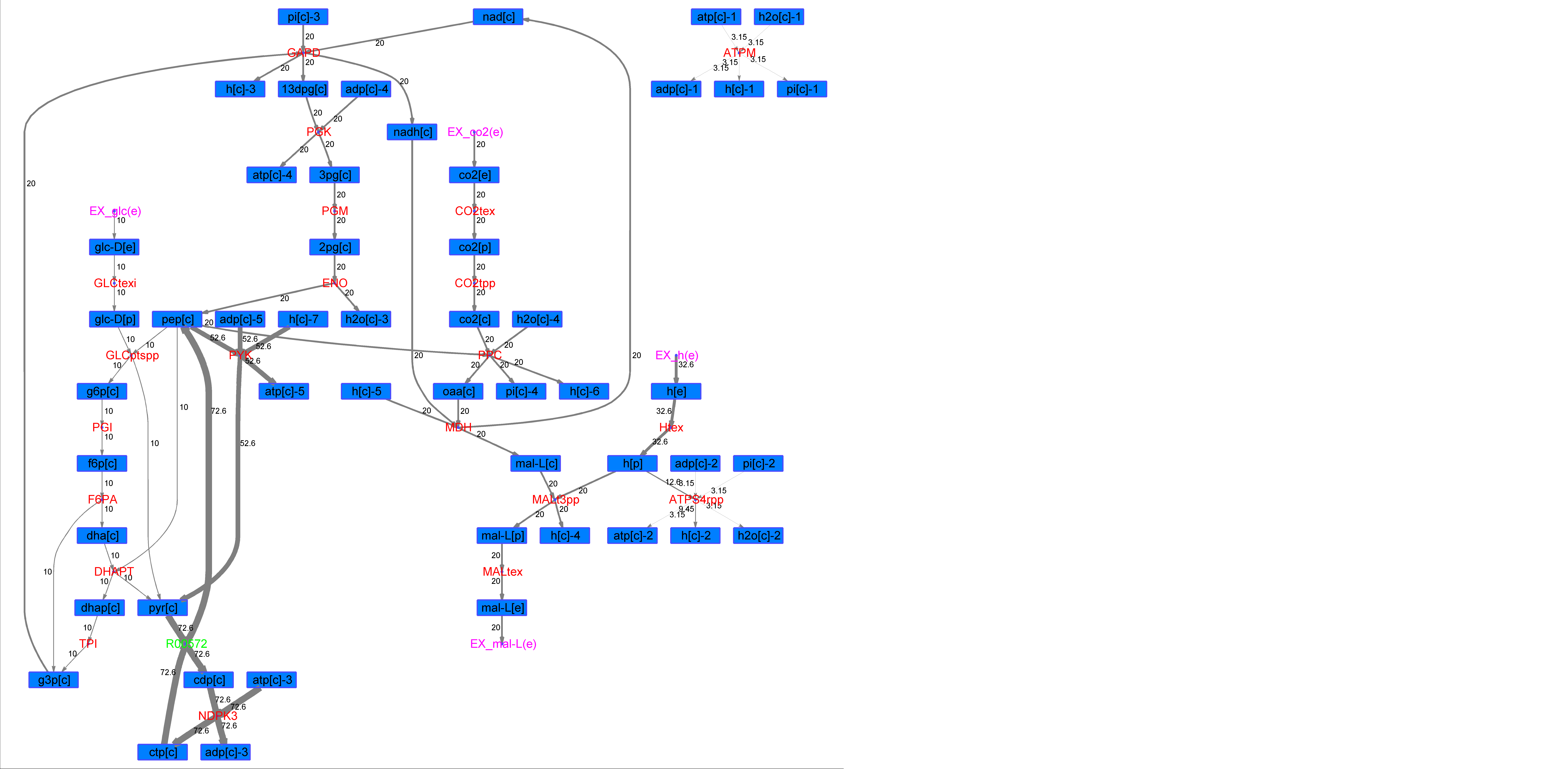

### R00572_L-Phe.png

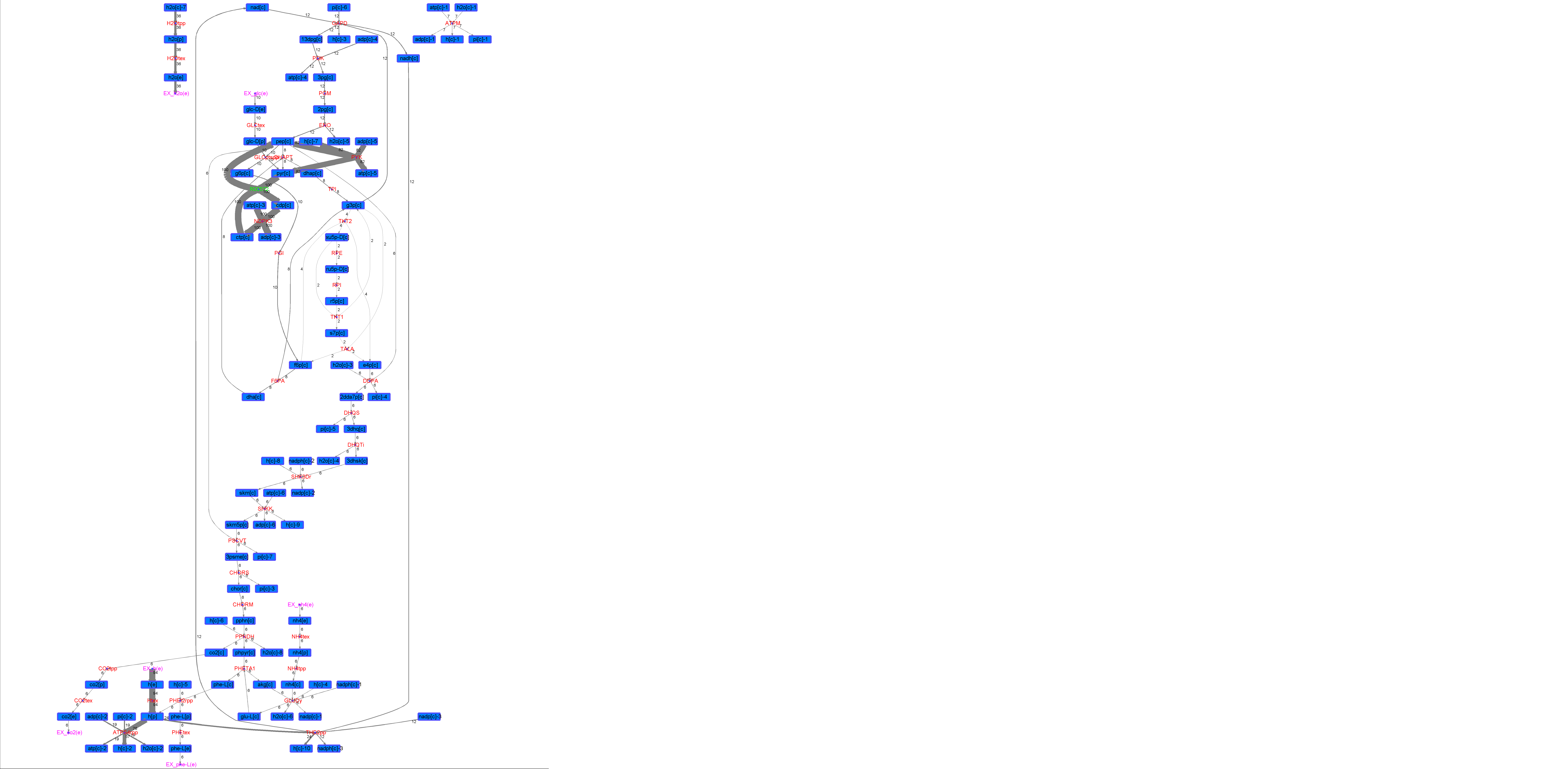

### R00572_L-Try.png

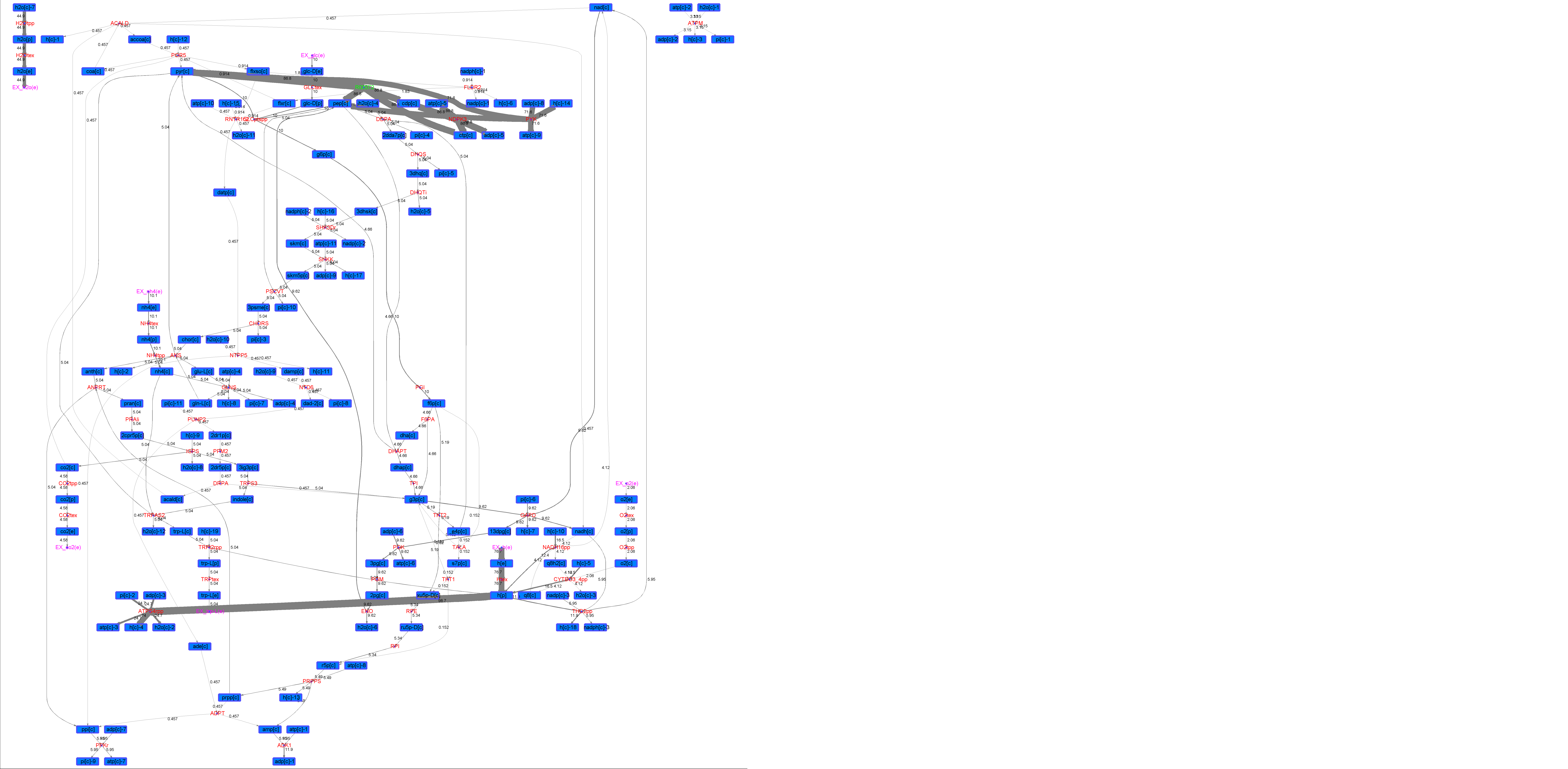

### R00572_Lys.png

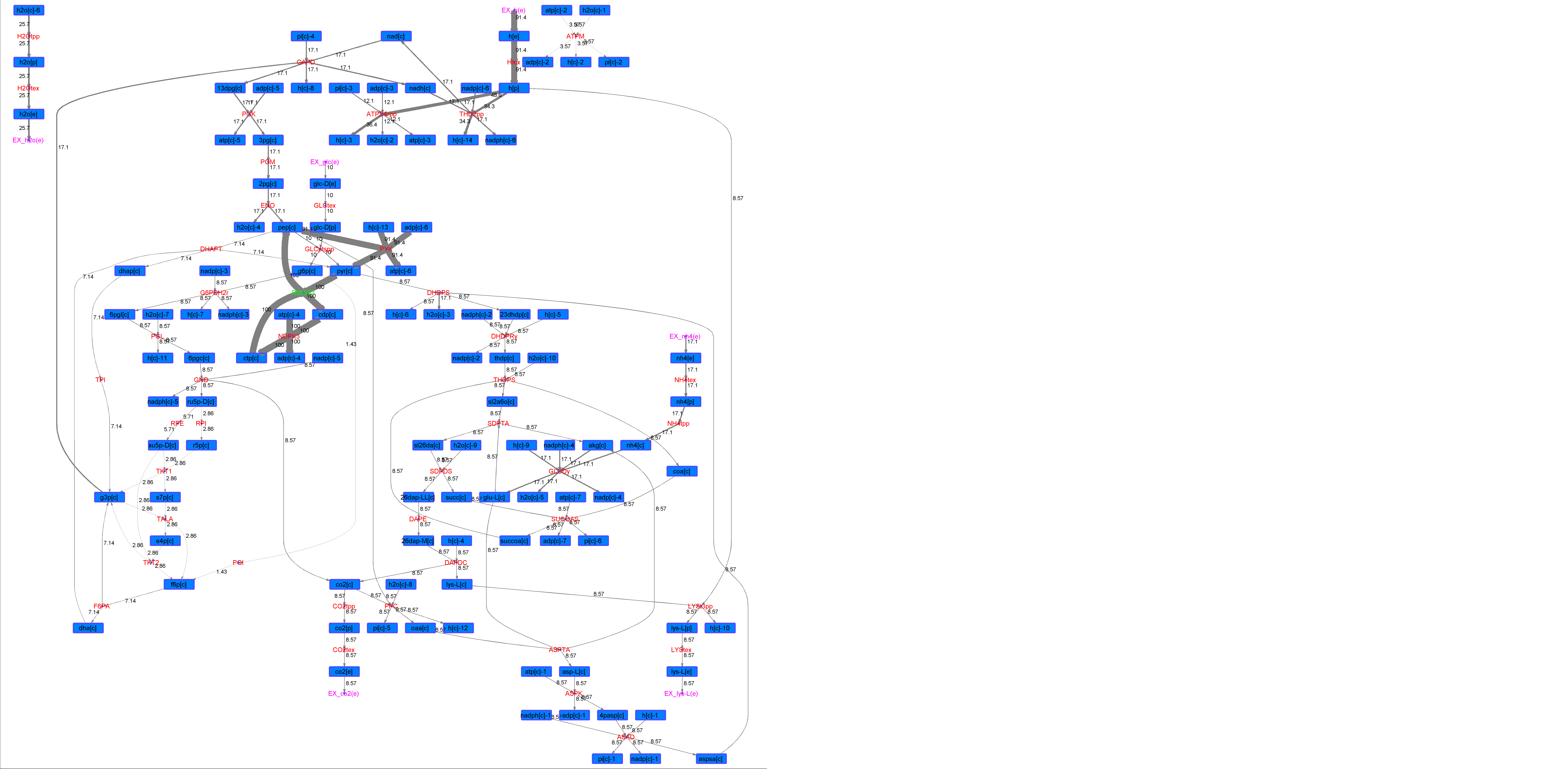

### R00572_Suc.png

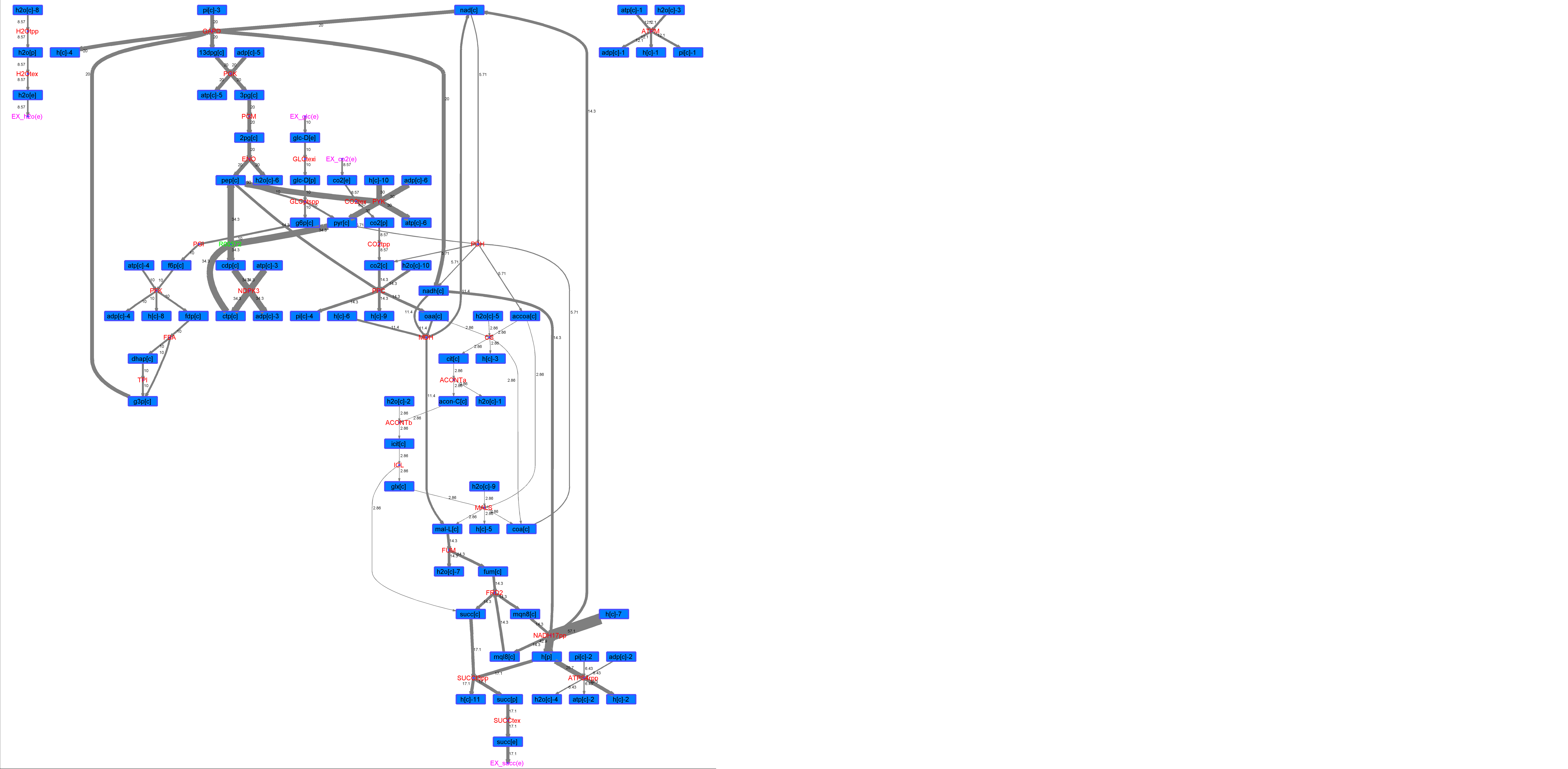

### R00572_Thr.png

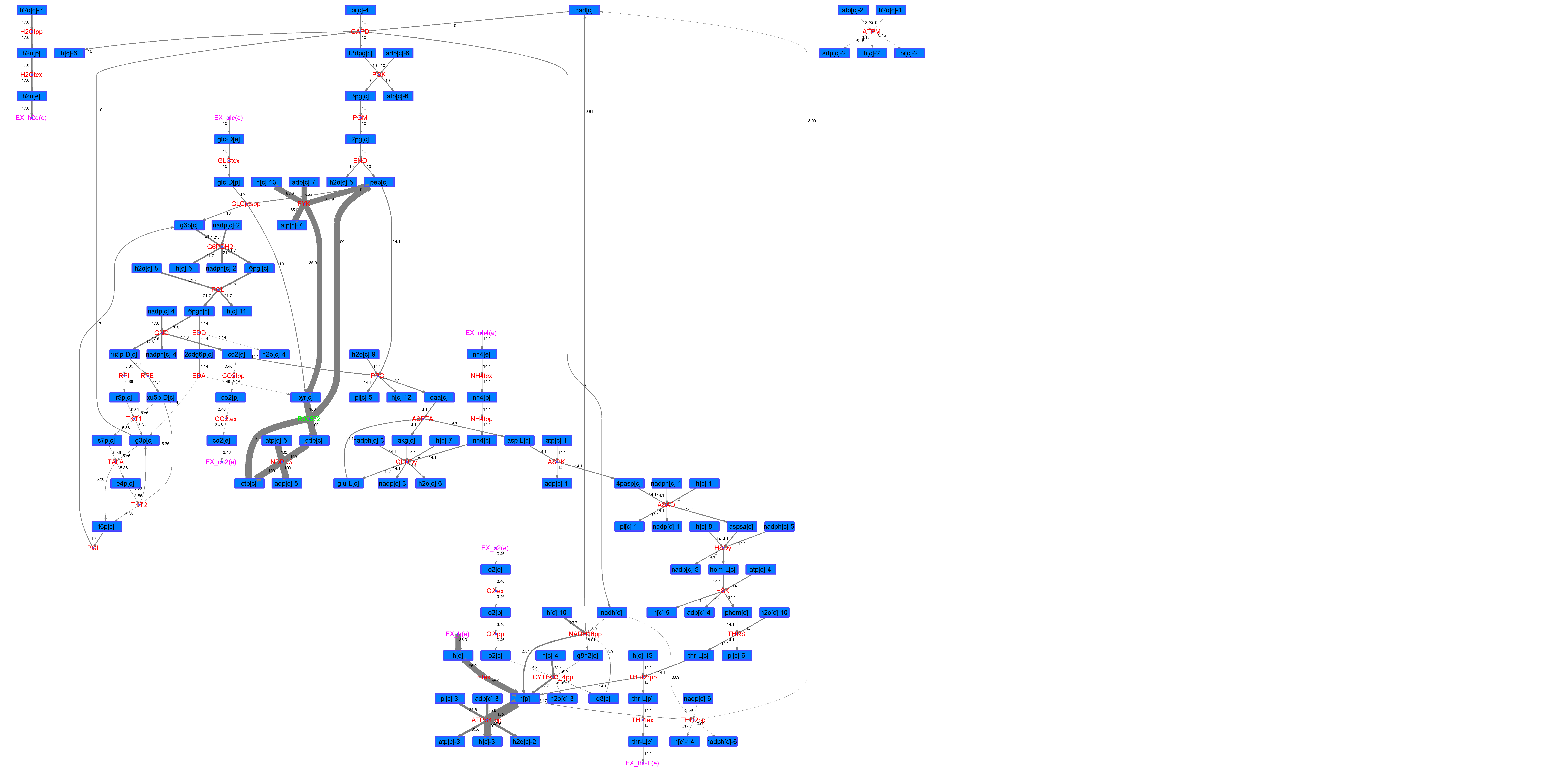

### R00585_Ace.png

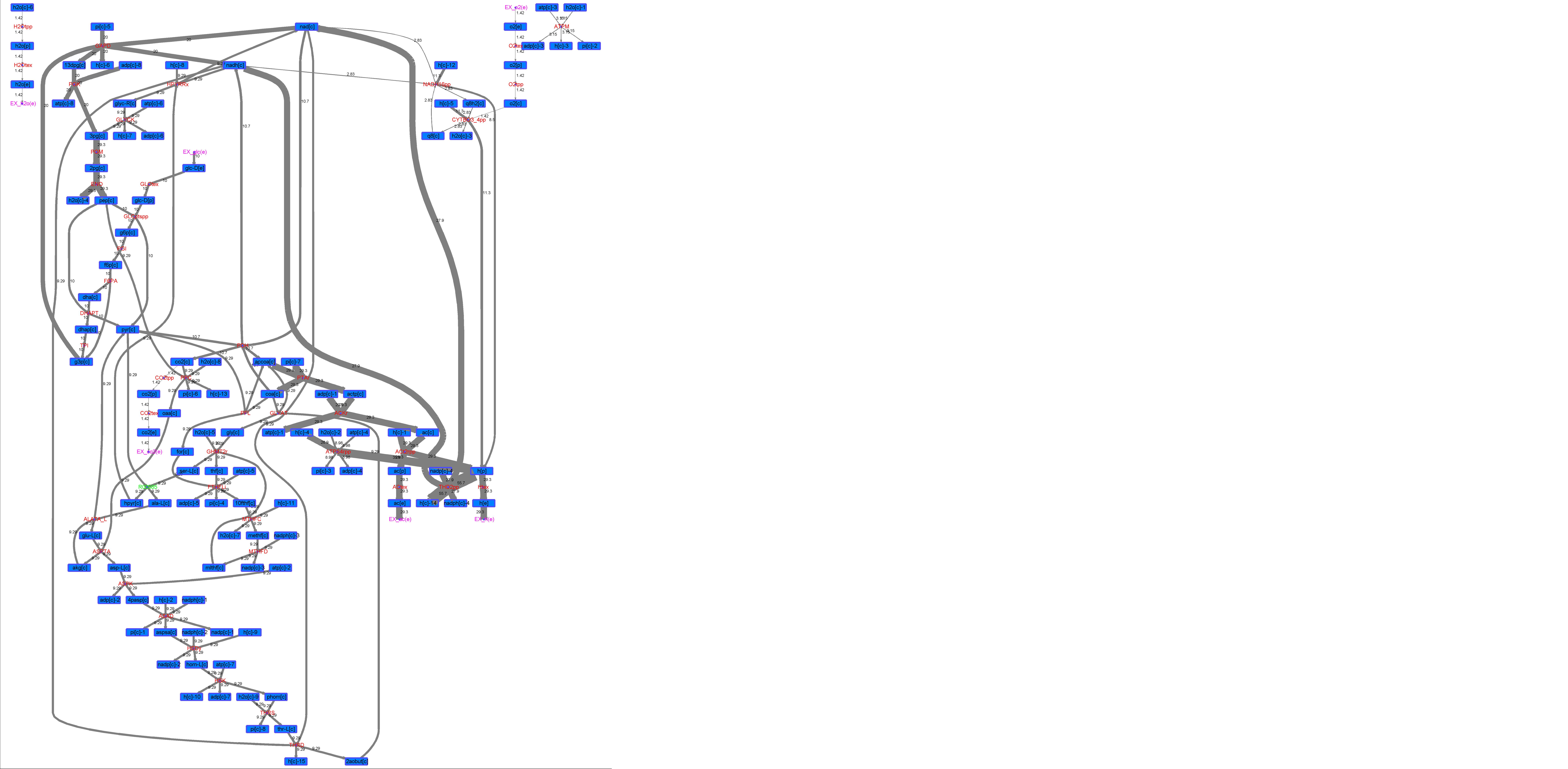

### R00585_For.png

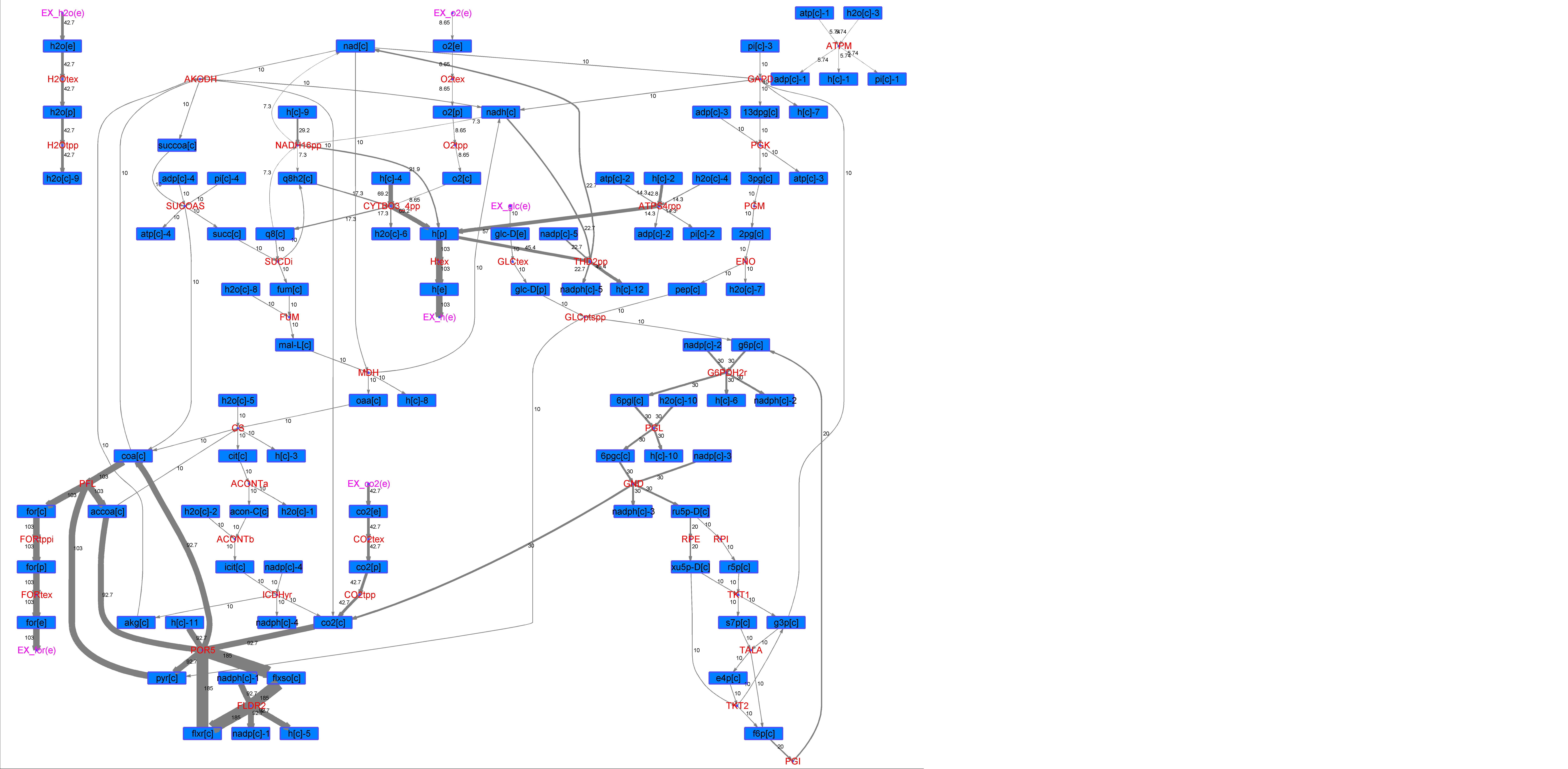

### R00585_Glu.png

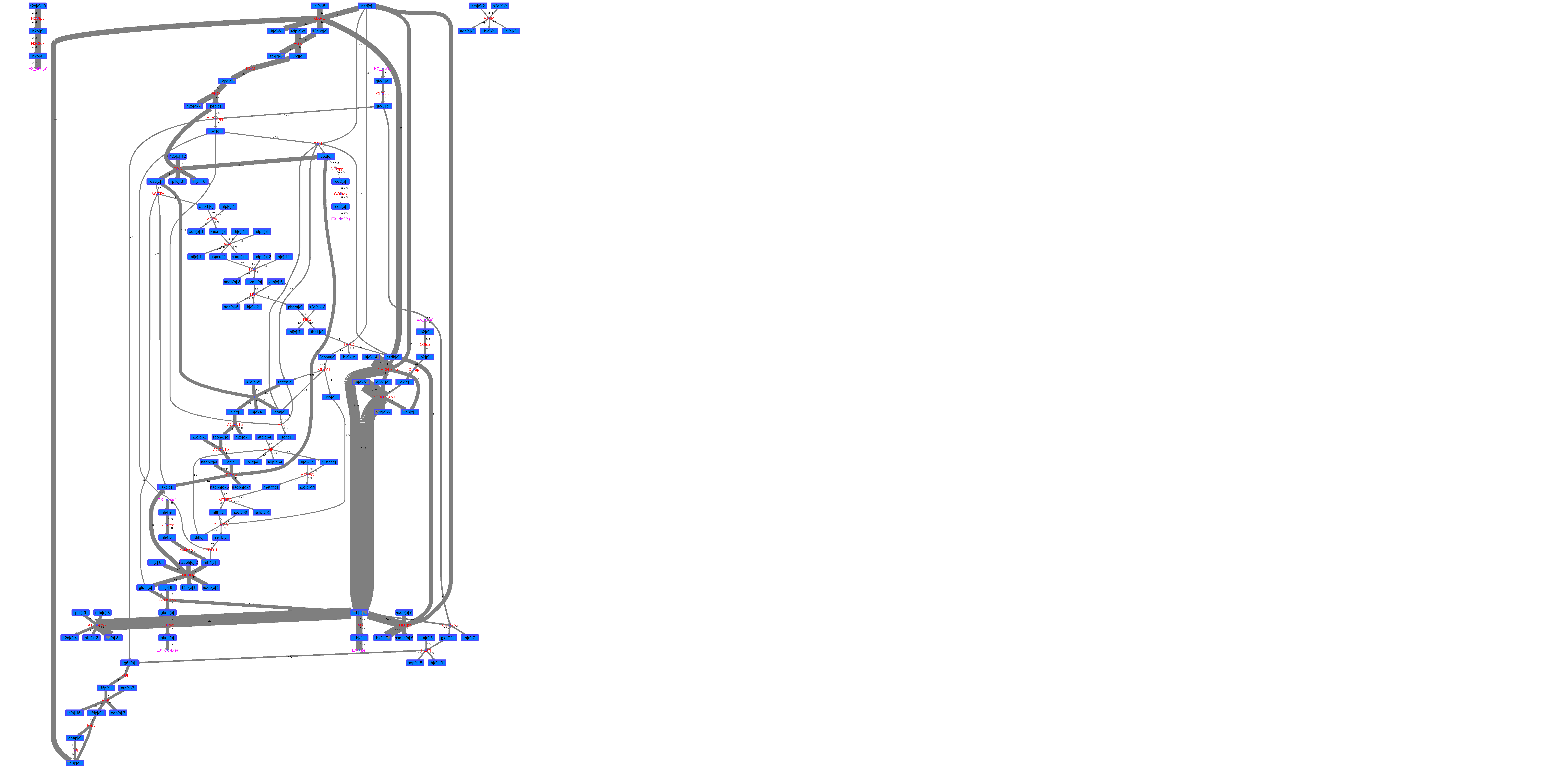

### R00585_L-ala.png

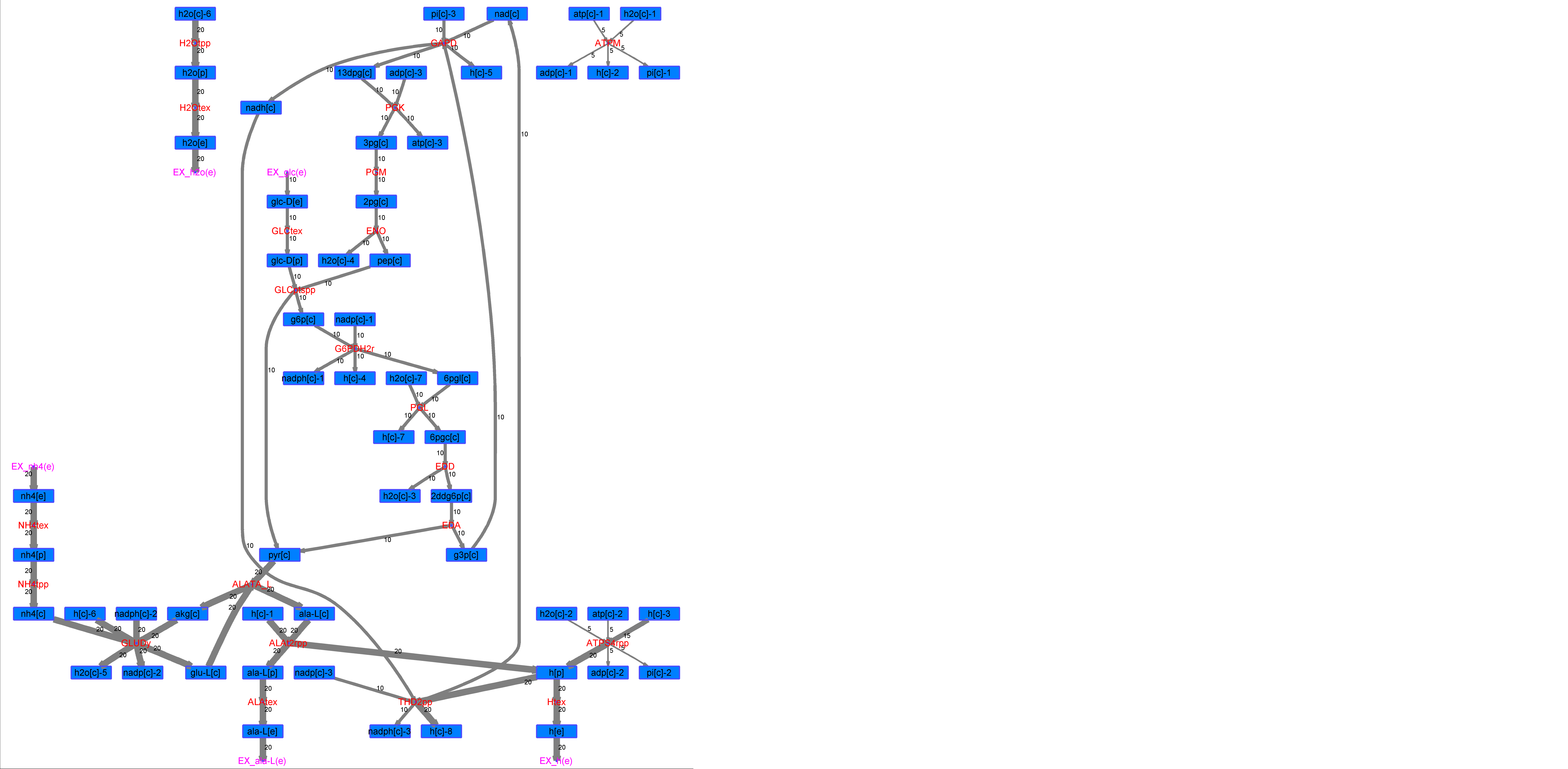

### R00585_L-Mal.png

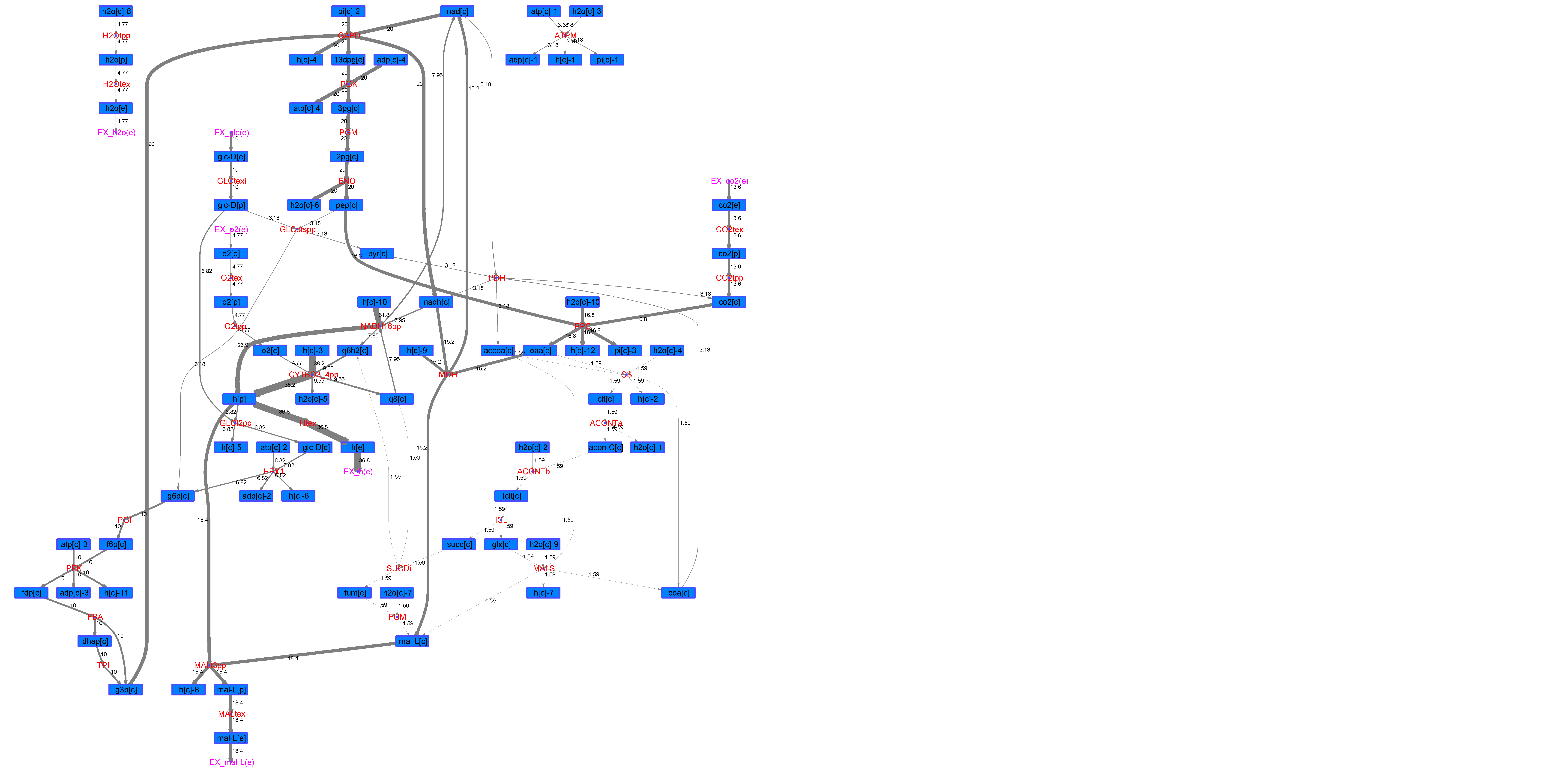

### R00585_L-Phe.png

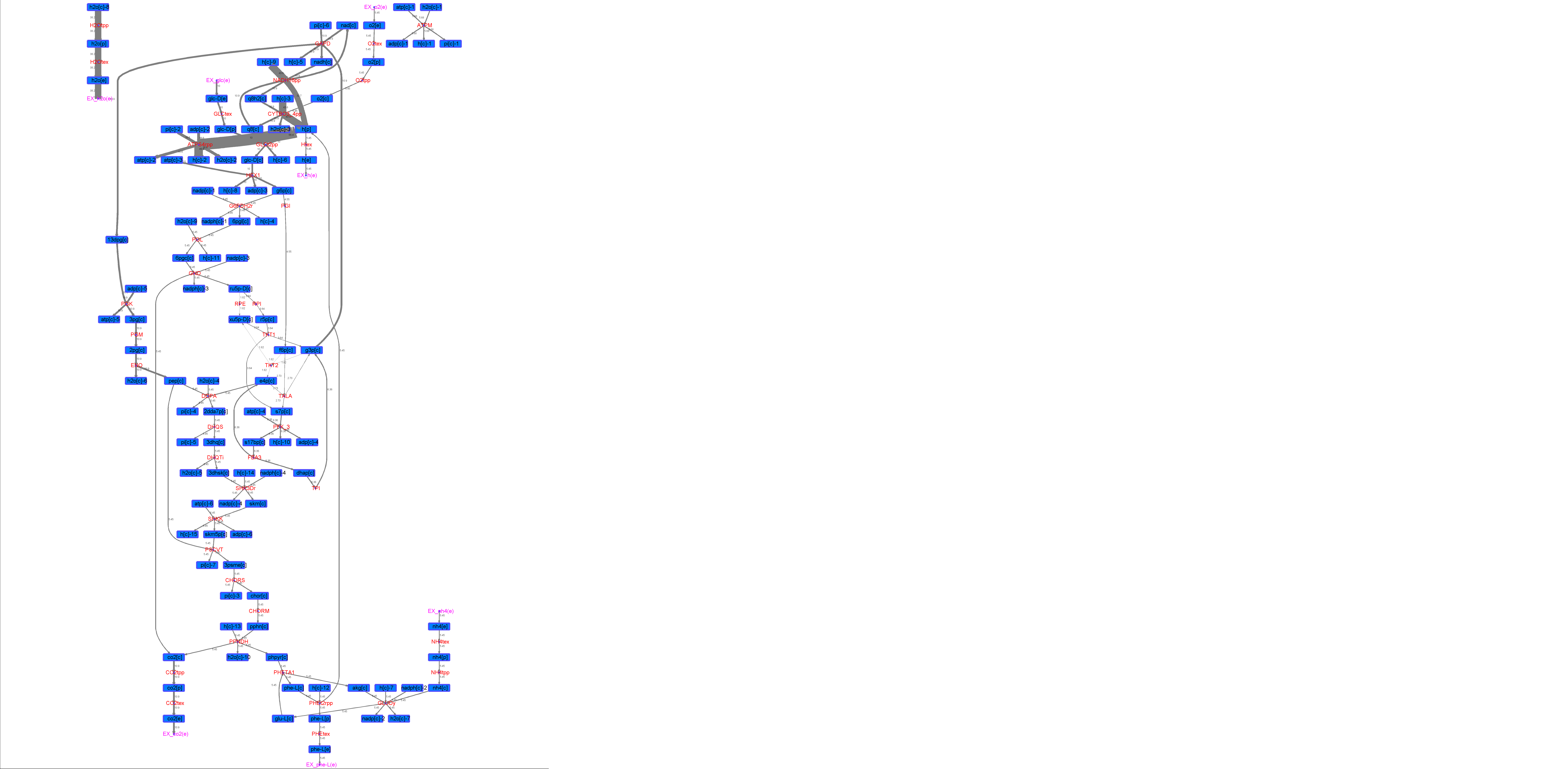

### R00585_L-Try.png

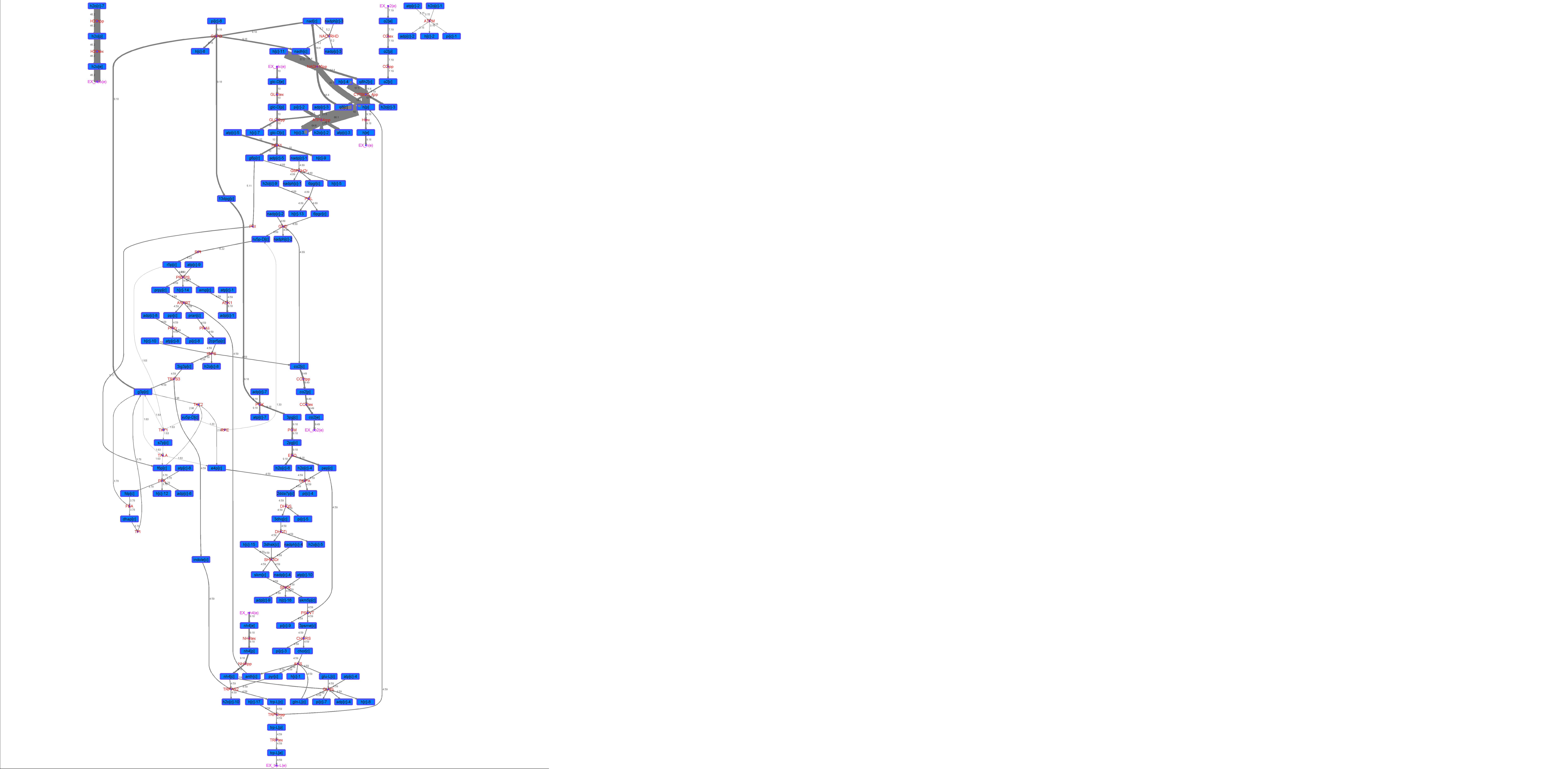

### R00585_Lys.png

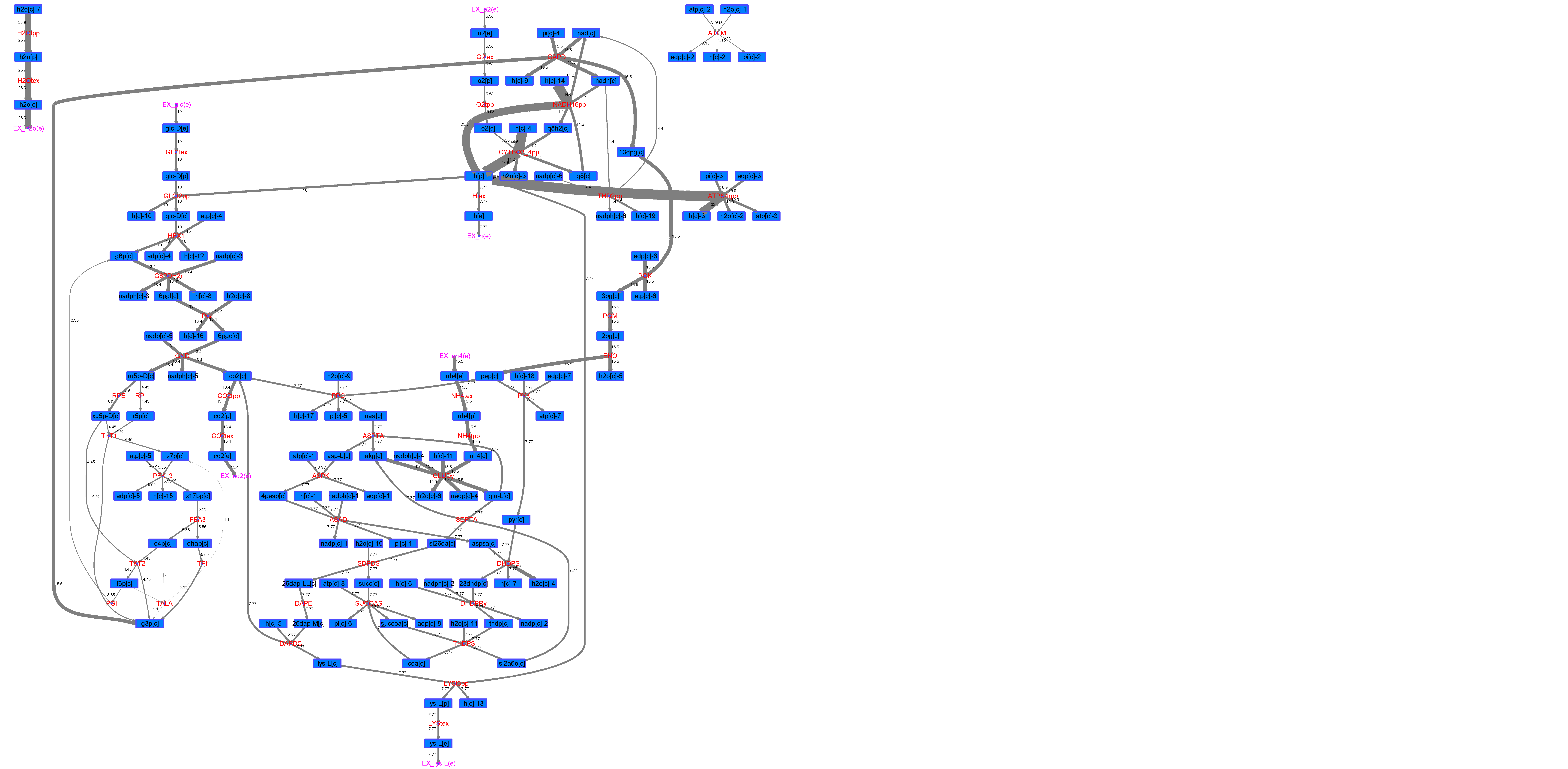

### R00585_Suc.png

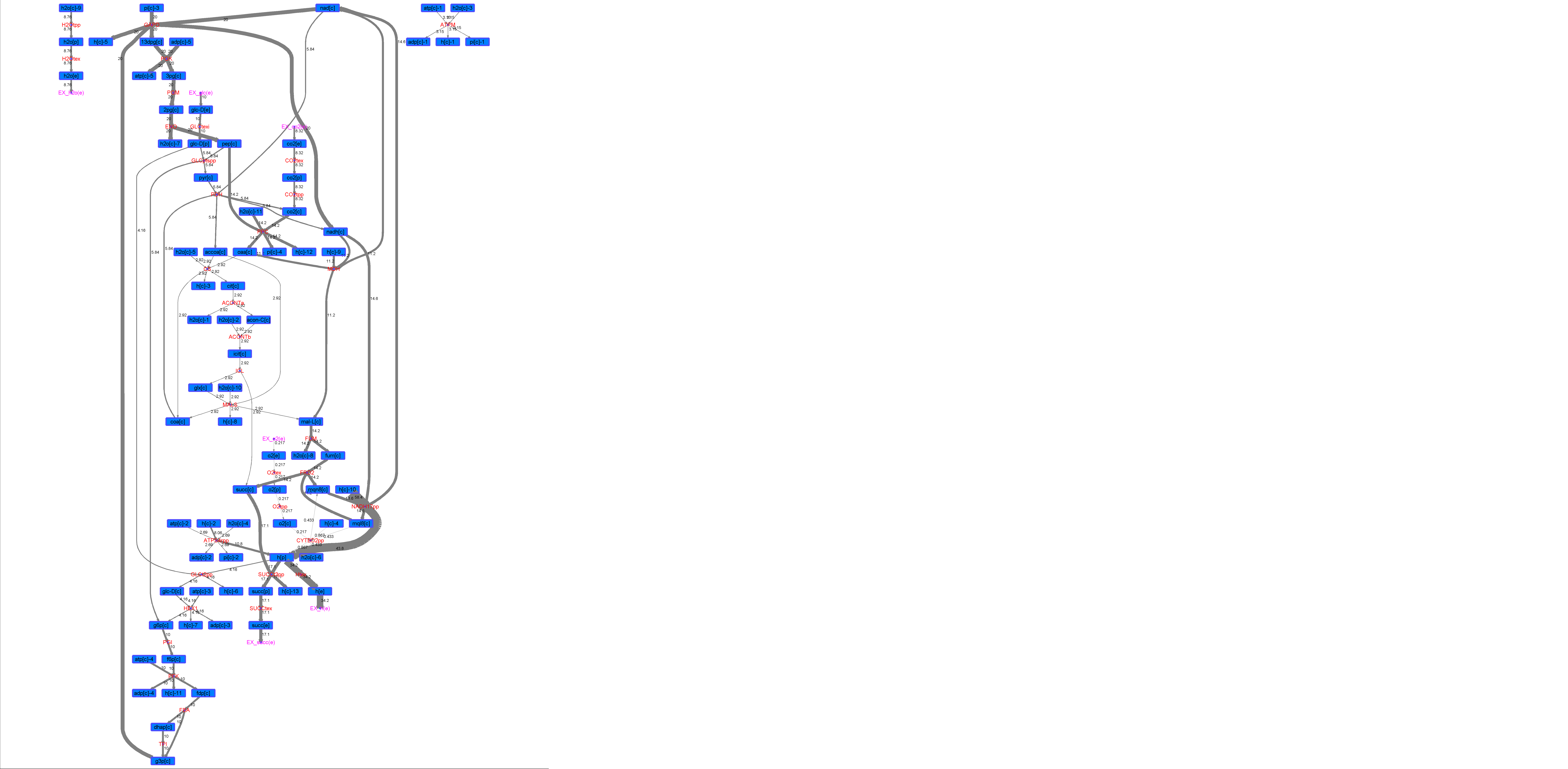

### R00585_Thr.png

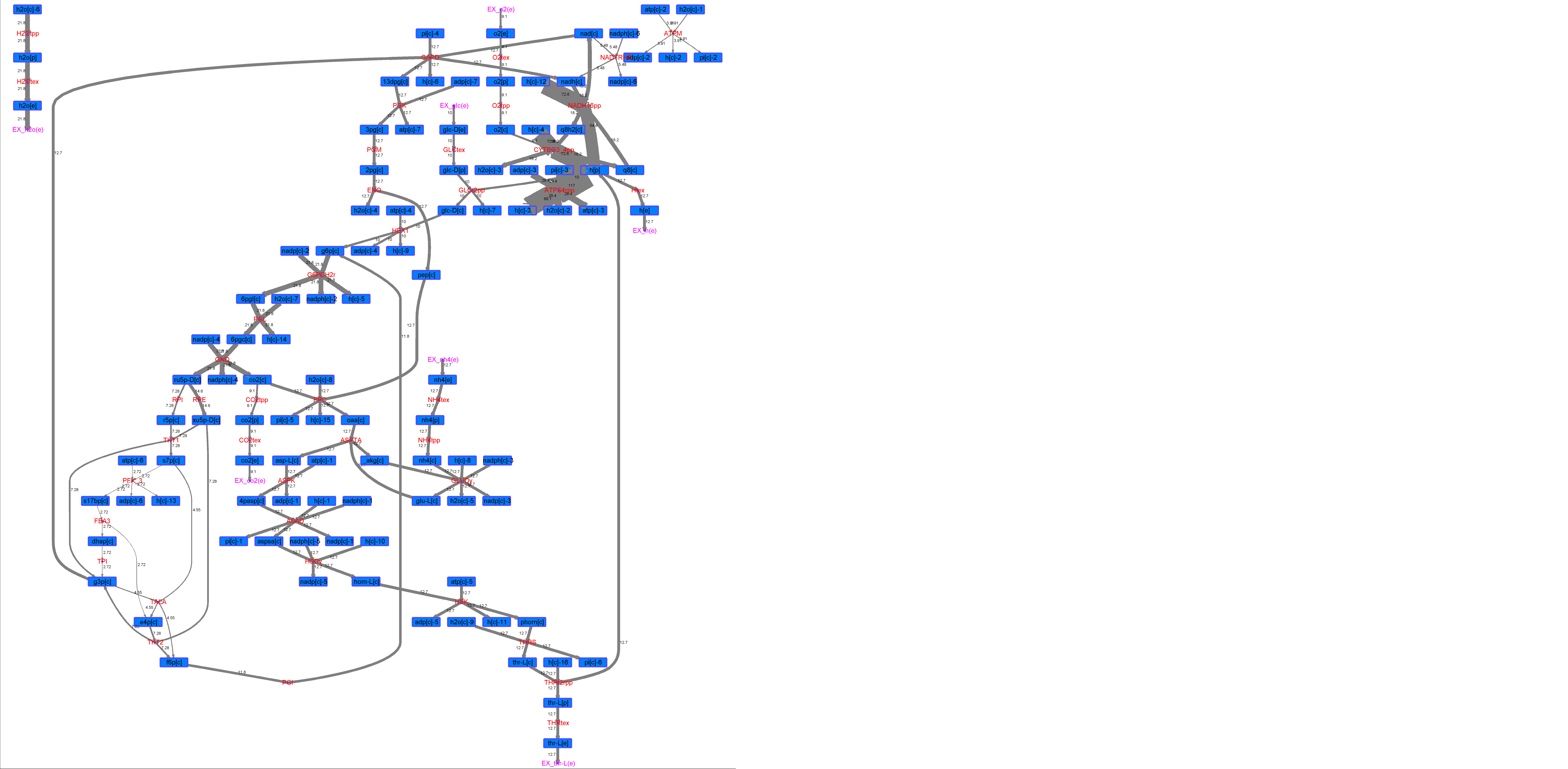
