## Supplementary figures and images for "The Enzymes that beyond Non-Oxidative Glycolysis"

### R00094_Ace.png

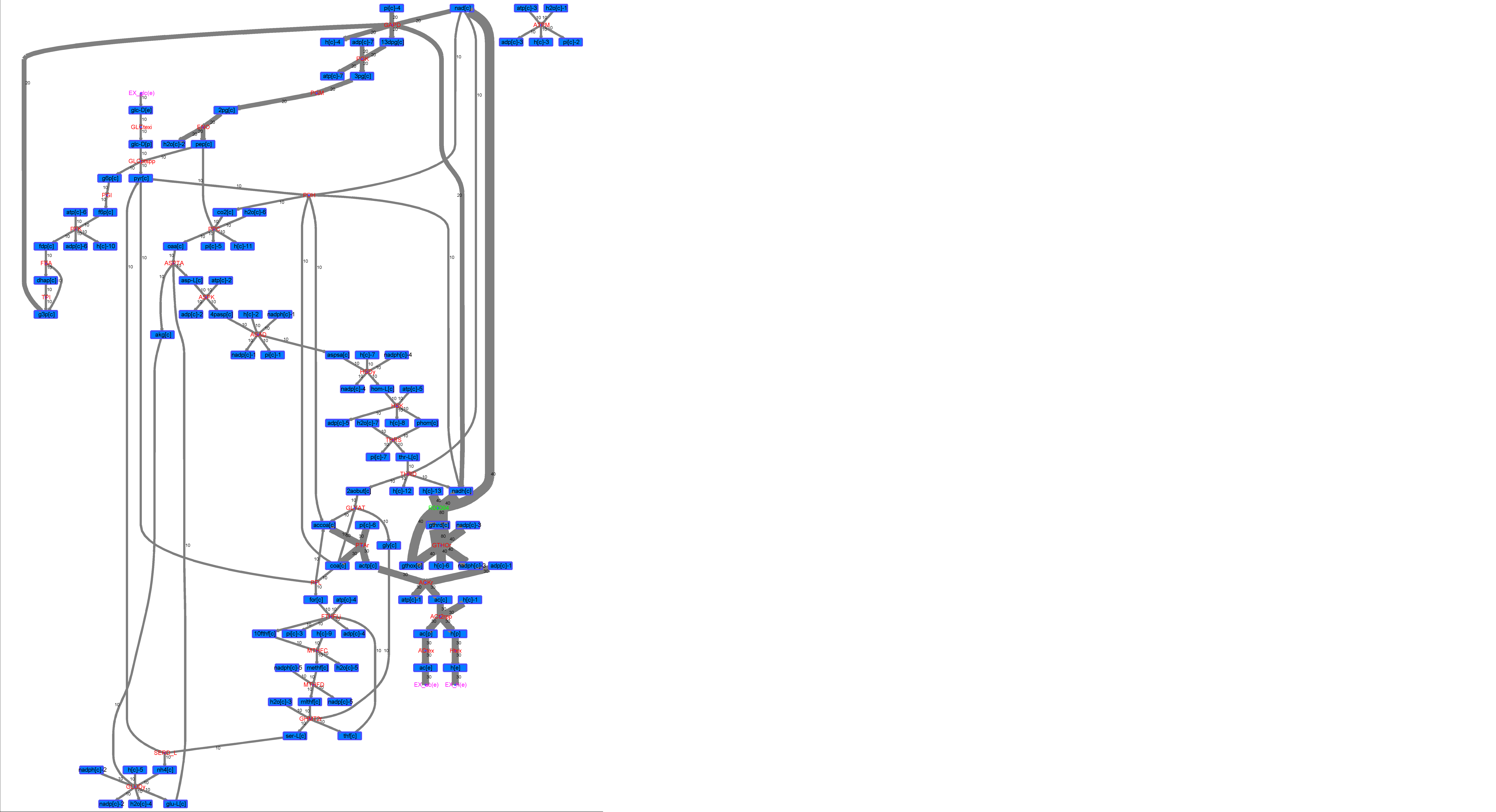

### R00094_For.png

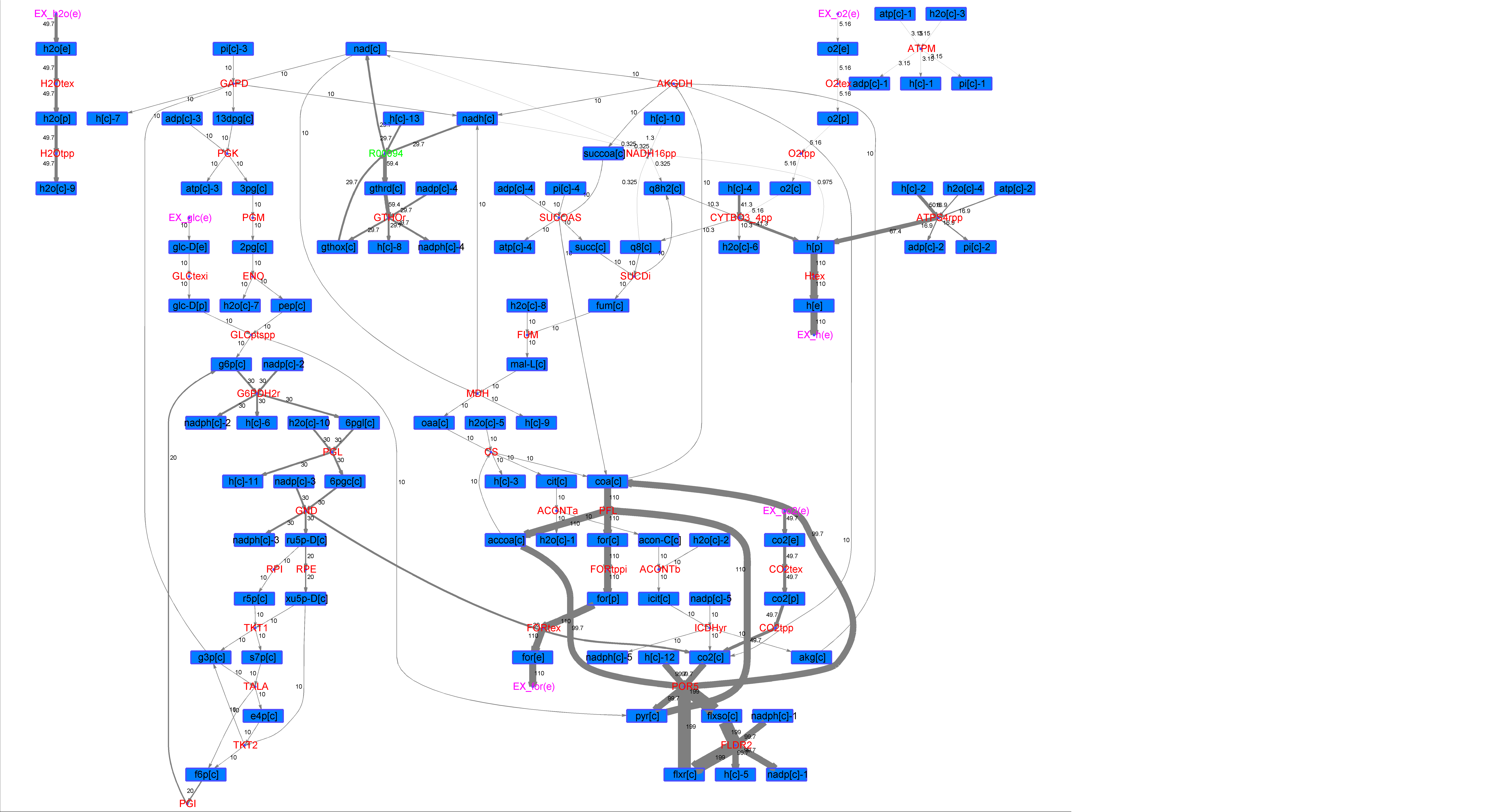

### R00094_Glu.png

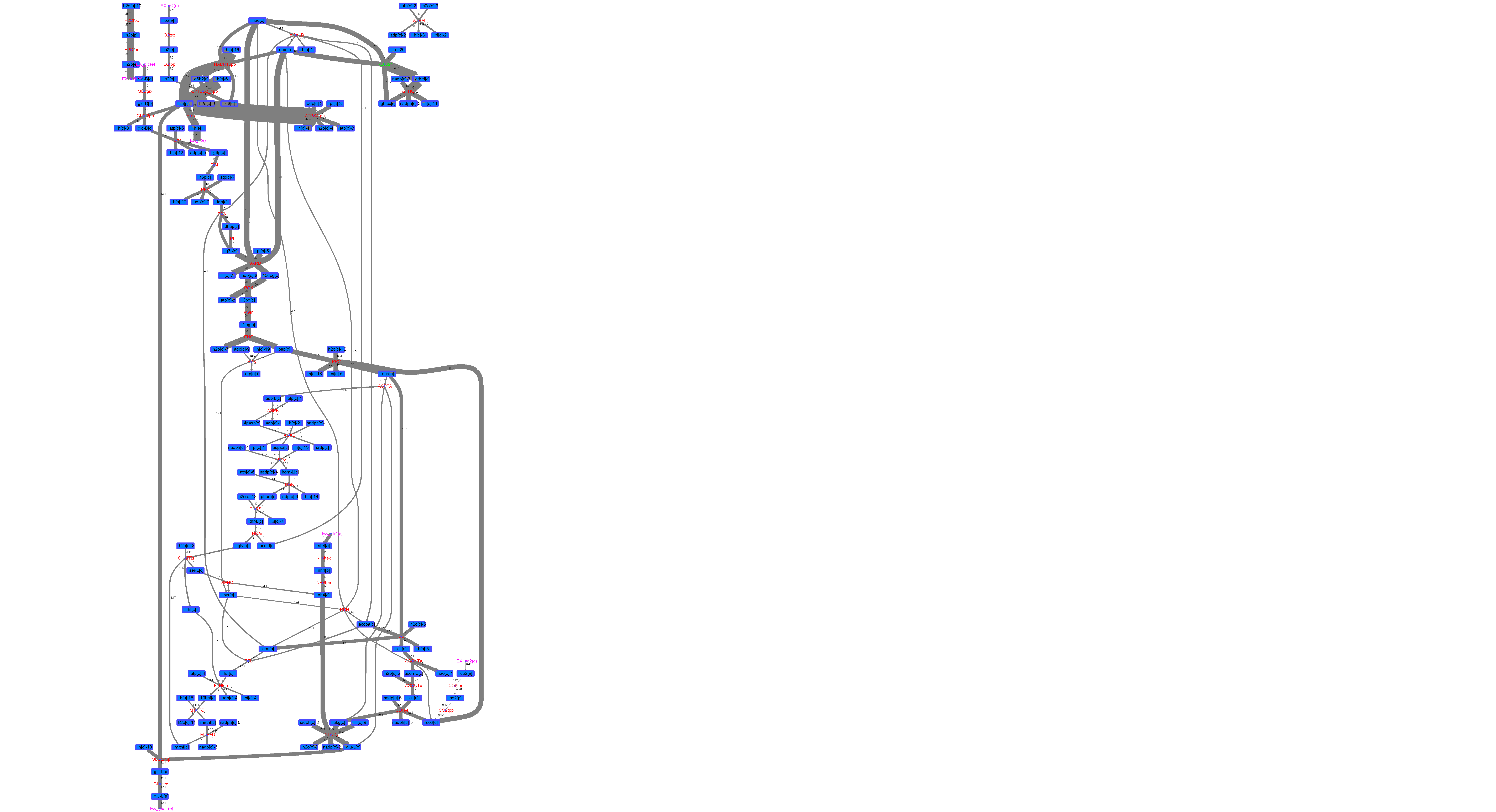

### R00094_L-ala.png

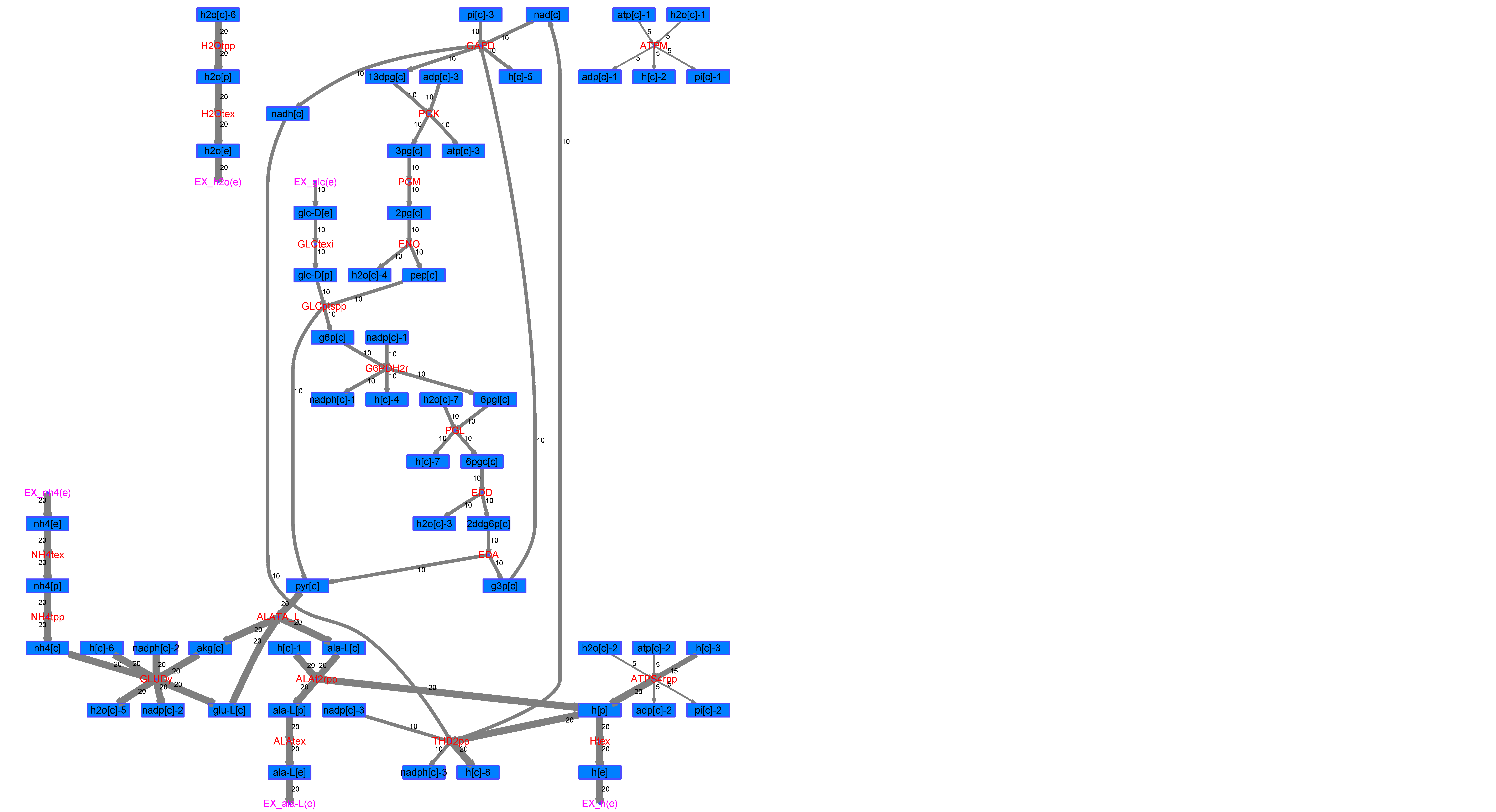

### R00094_L-Mal.png

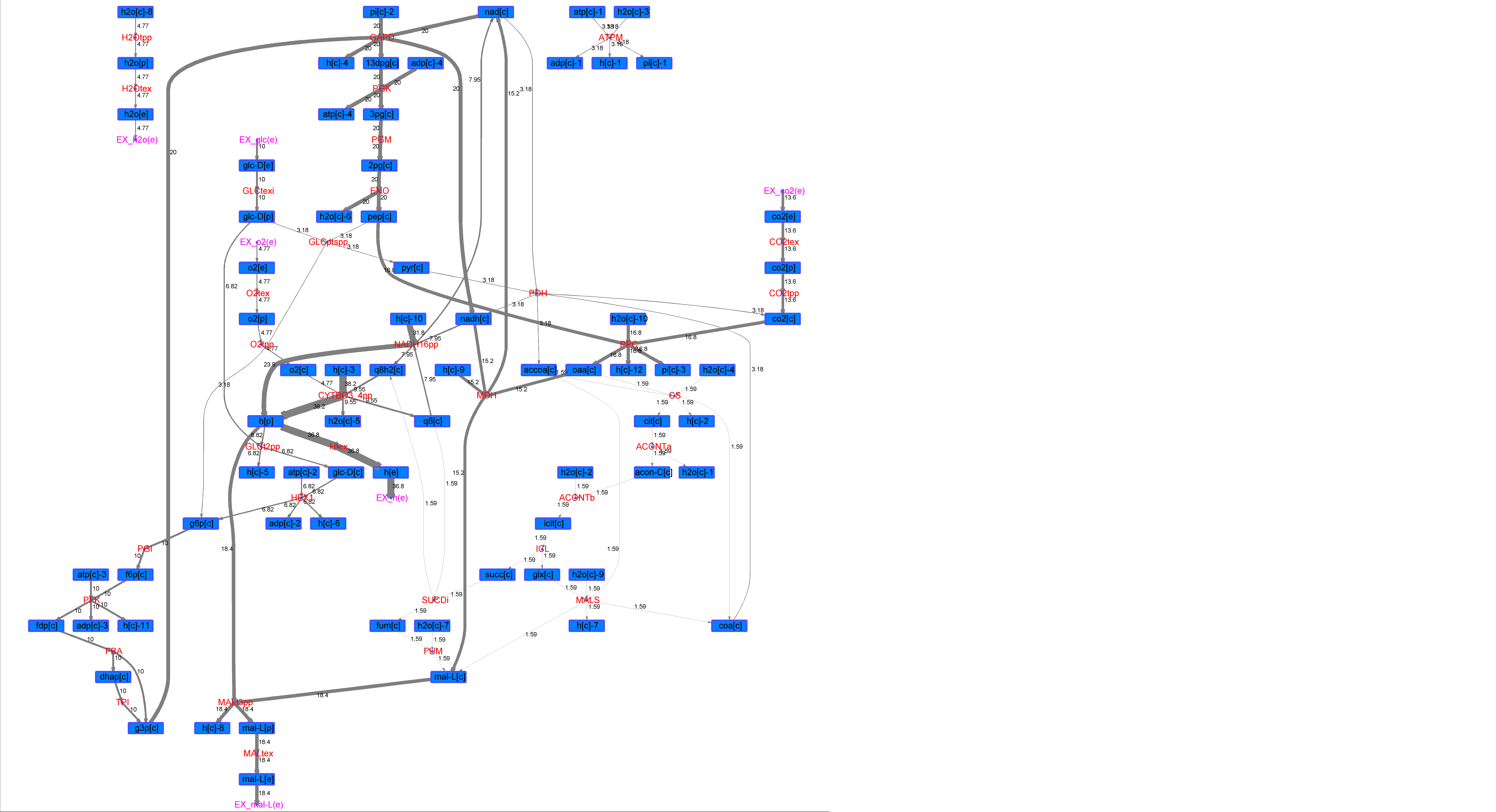

### R00094_L-Phe.png

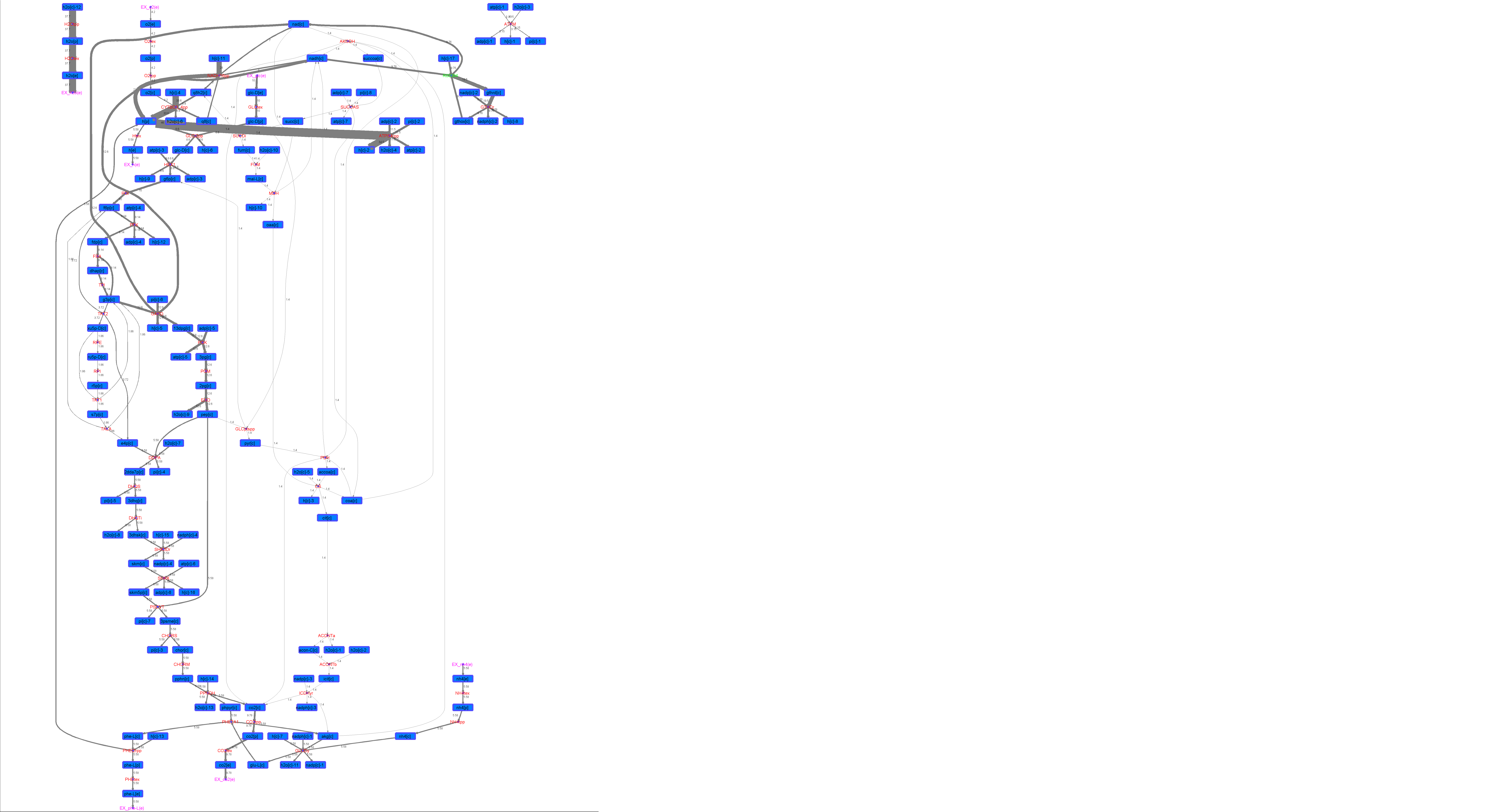

### R00094_L-Try.png

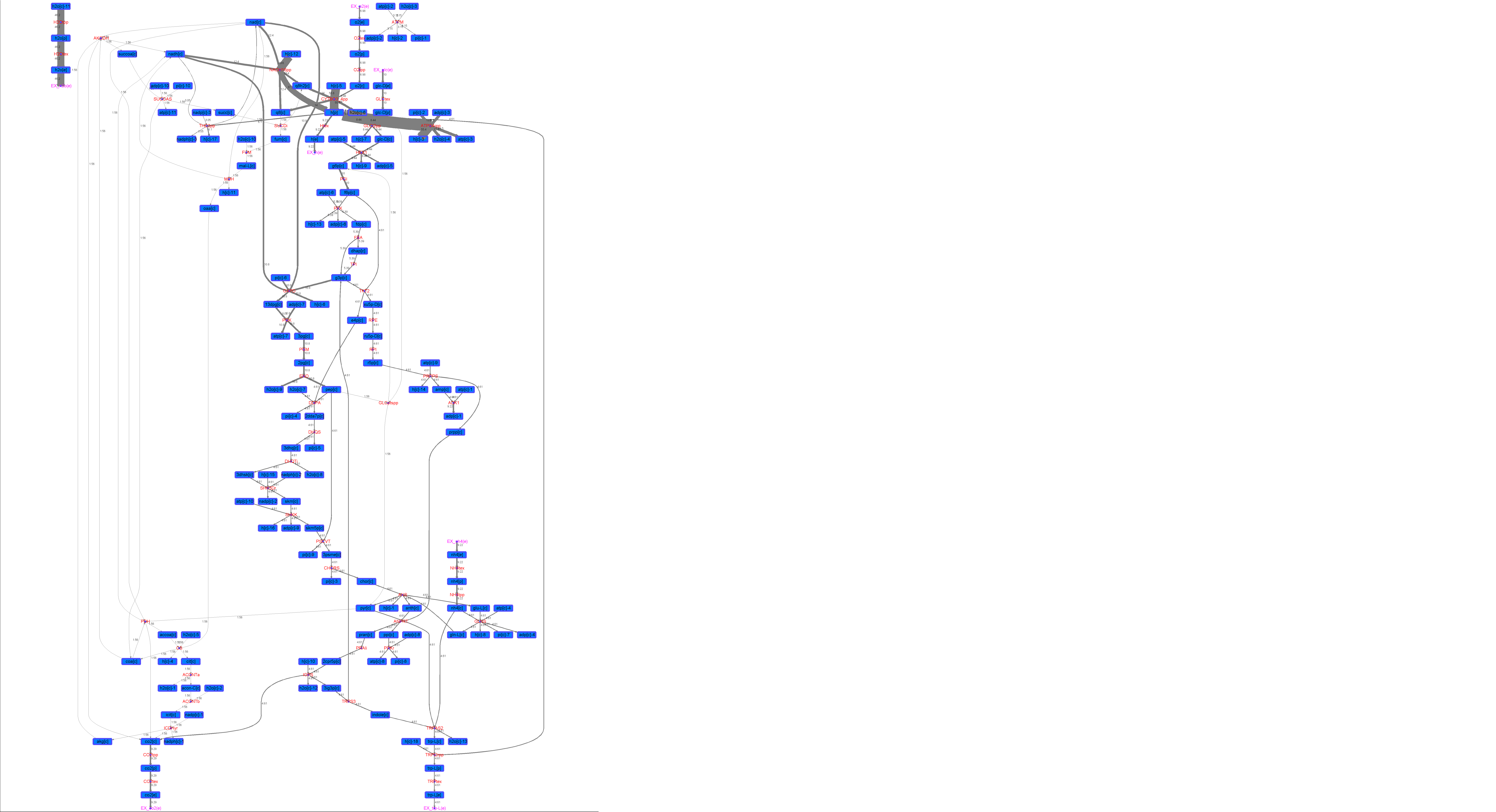

### R00094_Lys.png

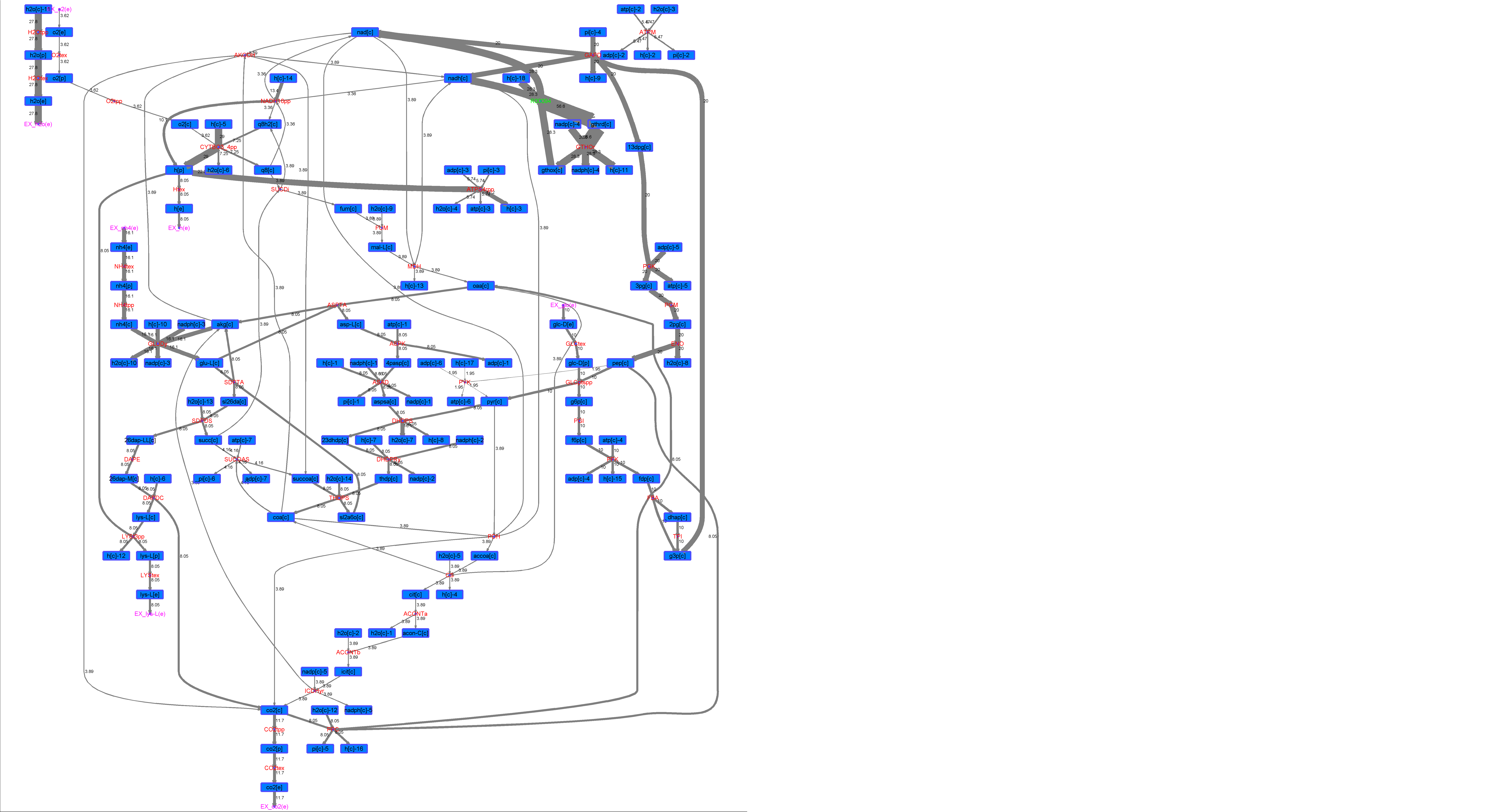

### R00094_Suc.png

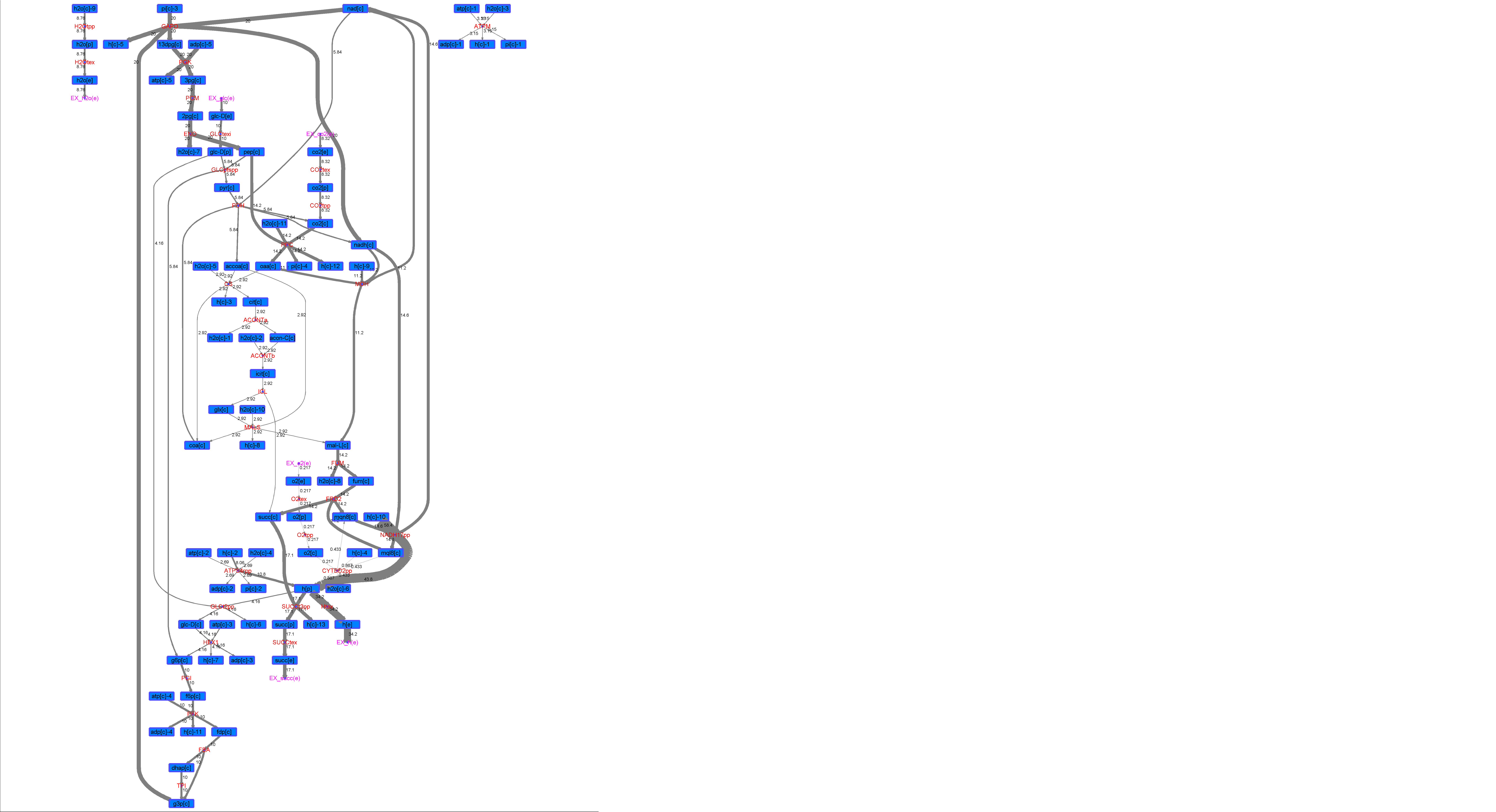

### R00094_Thr.png

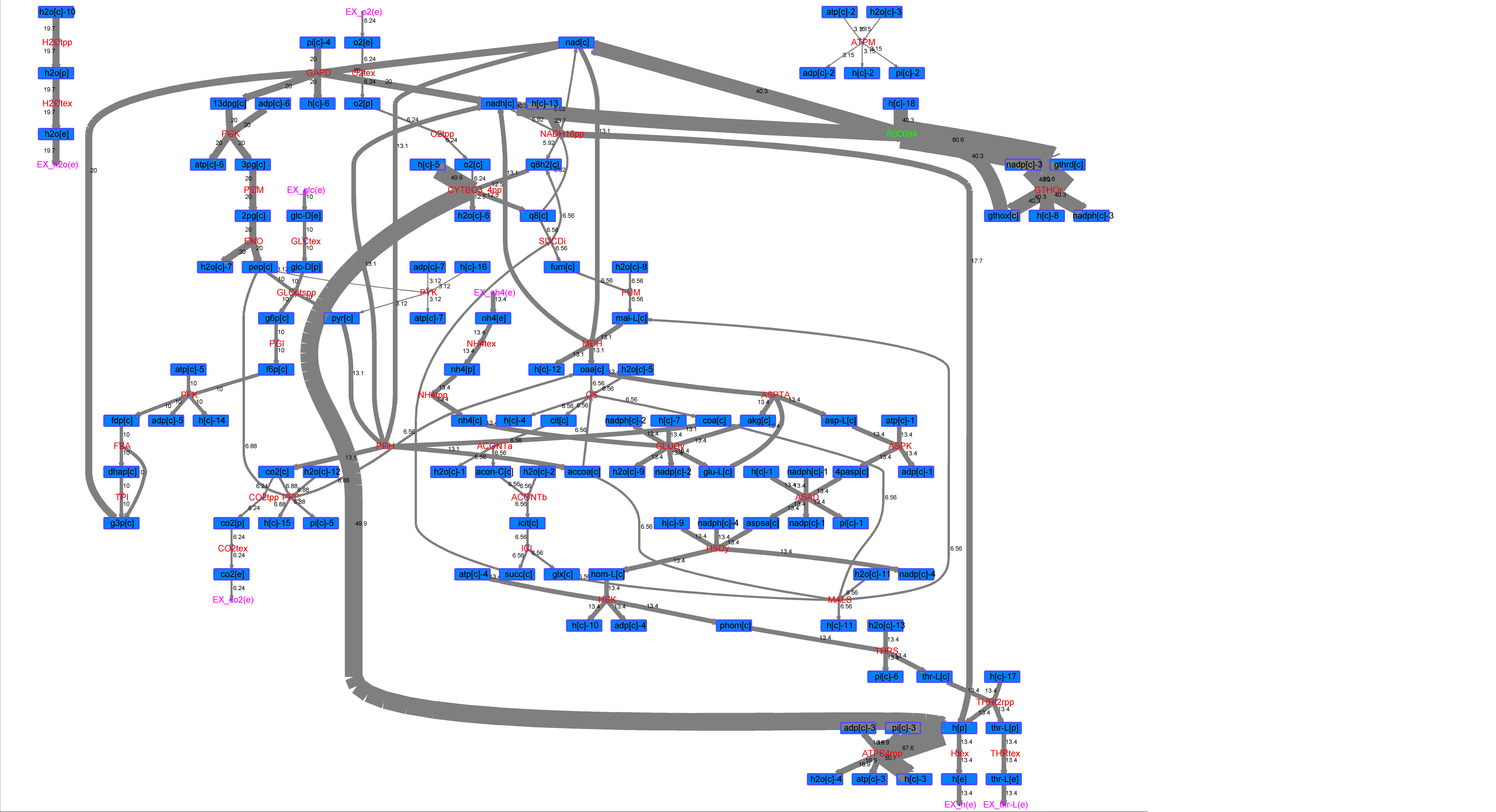

### R00126_Ace.png

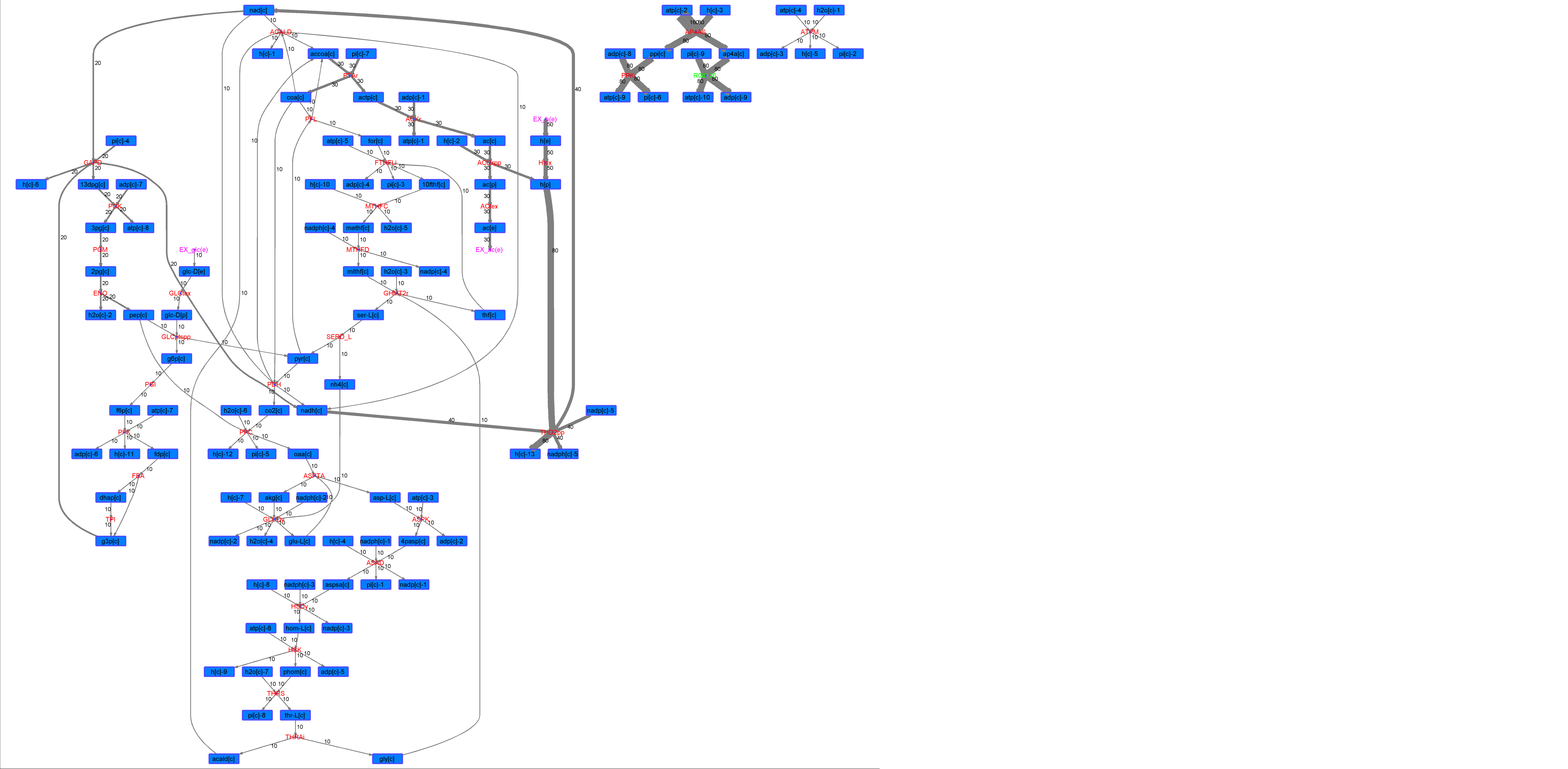

### R00126_For.png

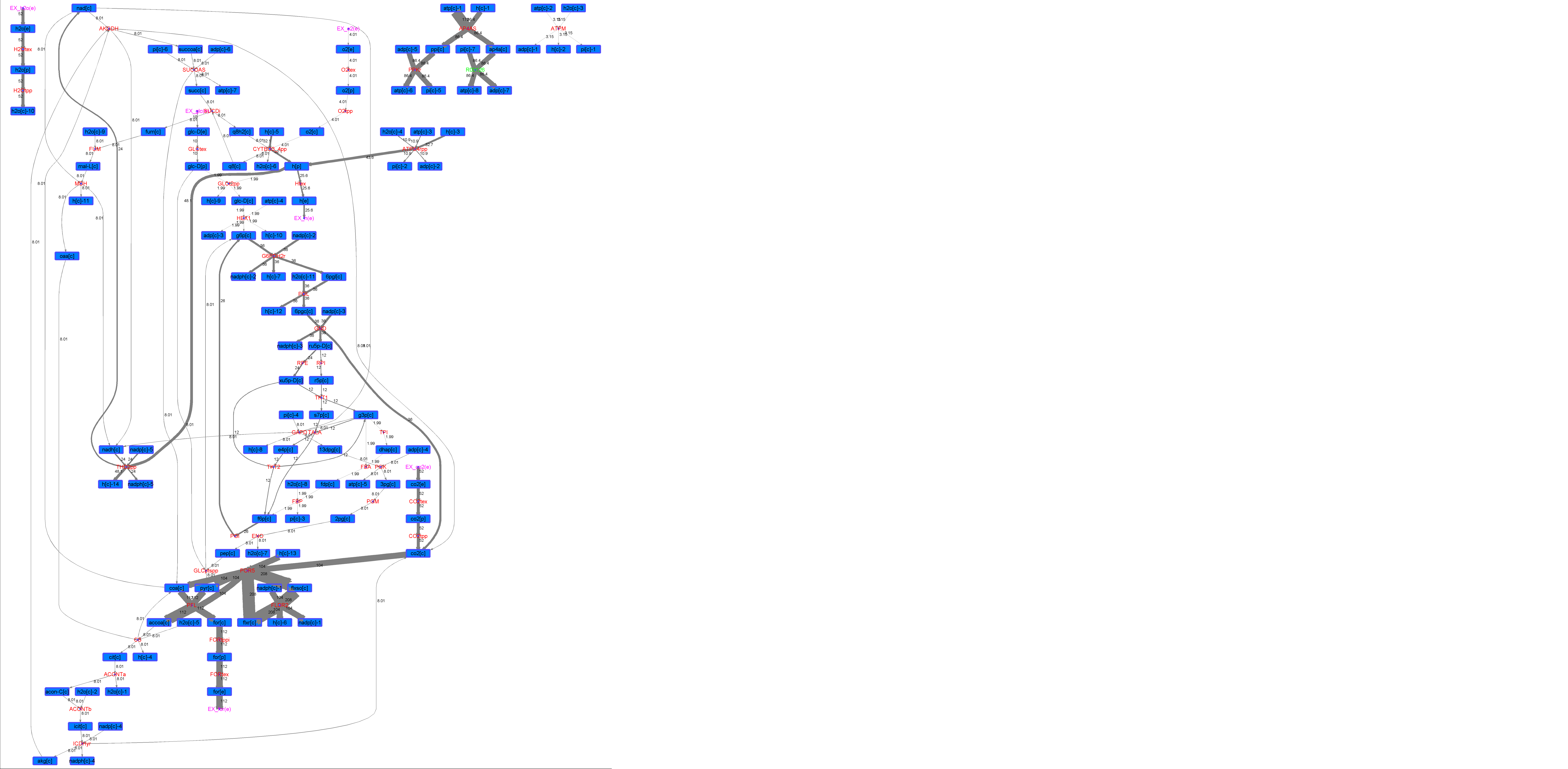

### R00126_Glu.png

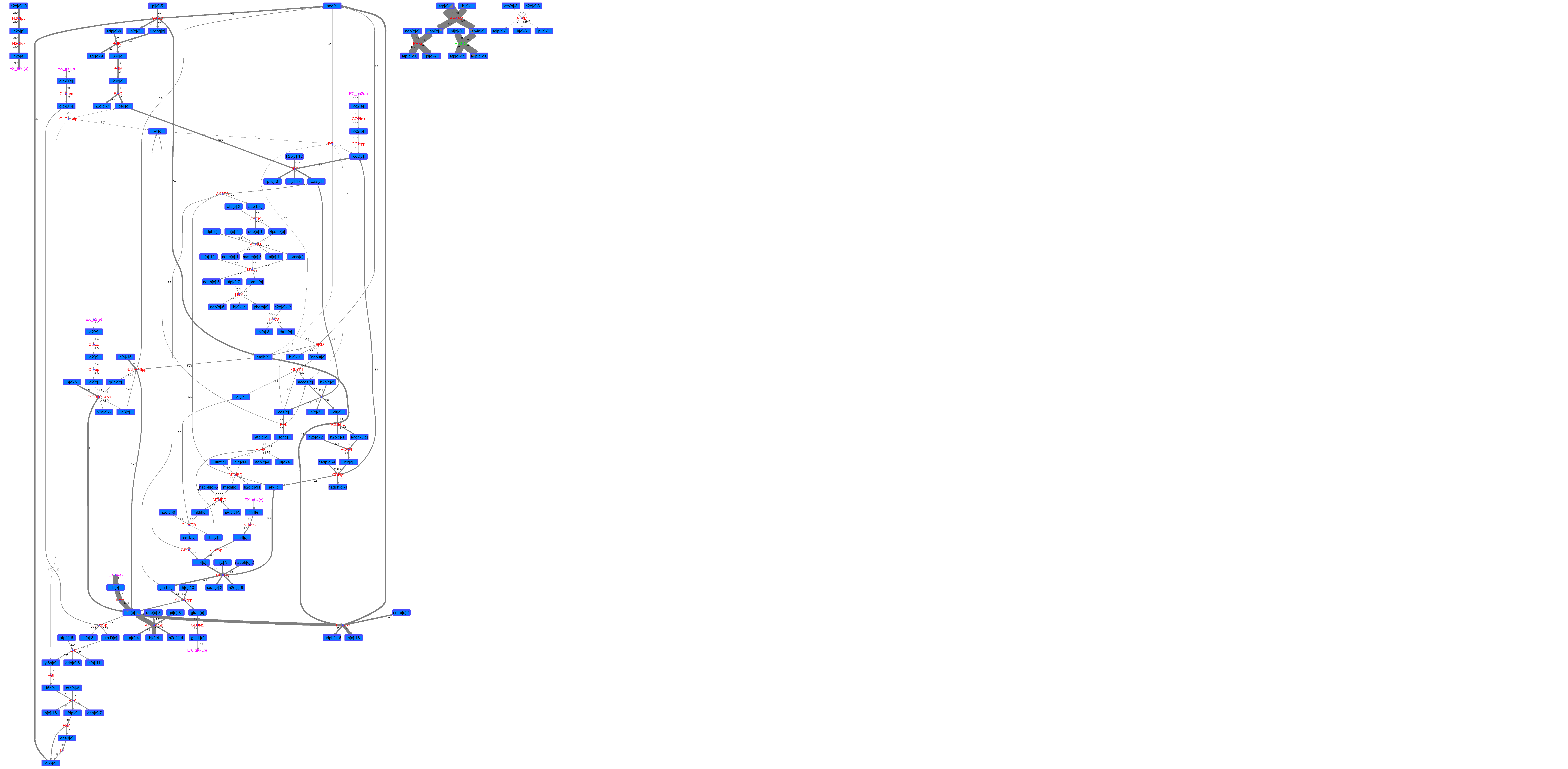

### R00126_L-ala.png

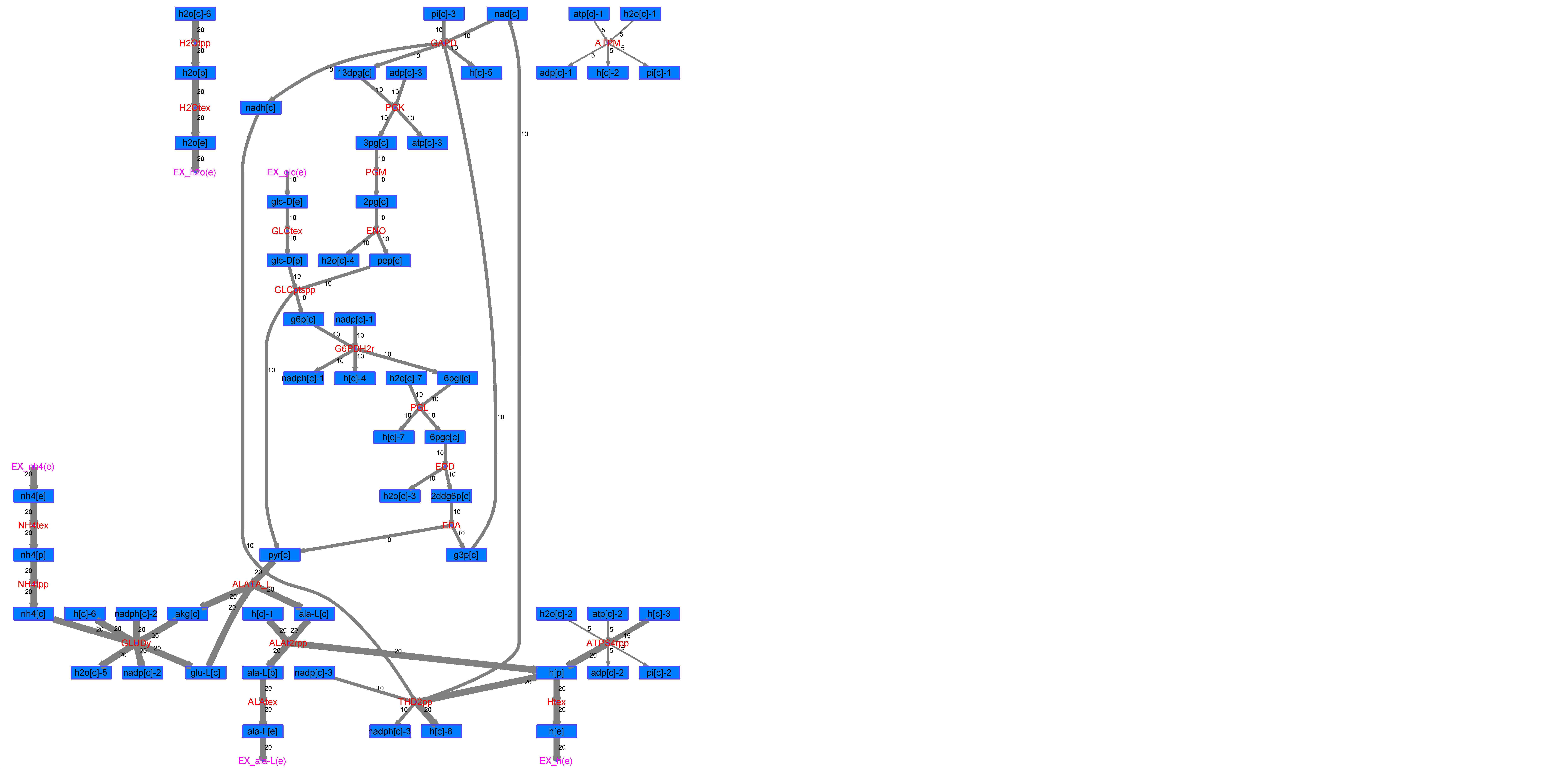

### R00126_L-Mal.png

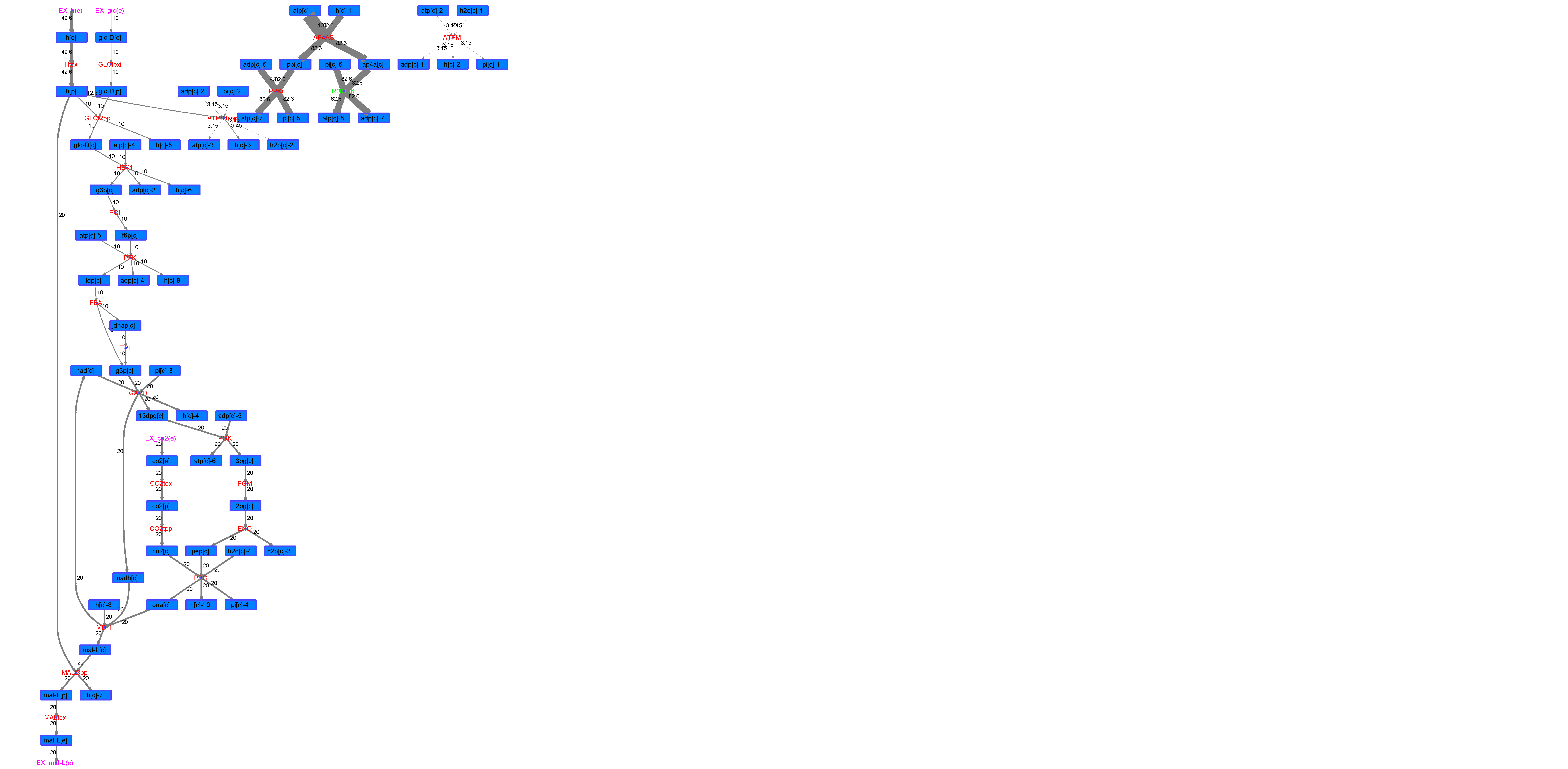

### R00126_L-Phe.png

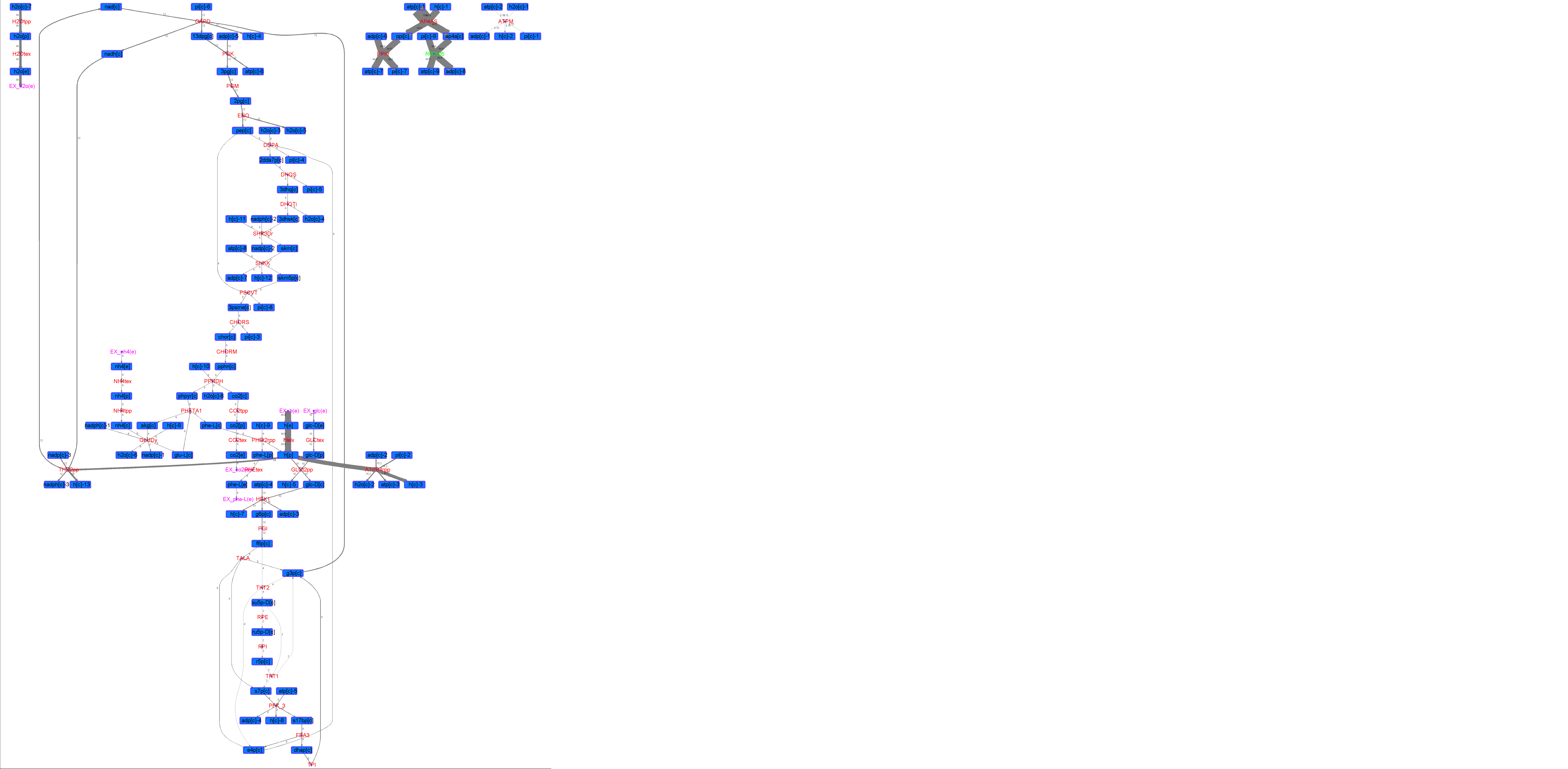

### R00126_L-Try.png

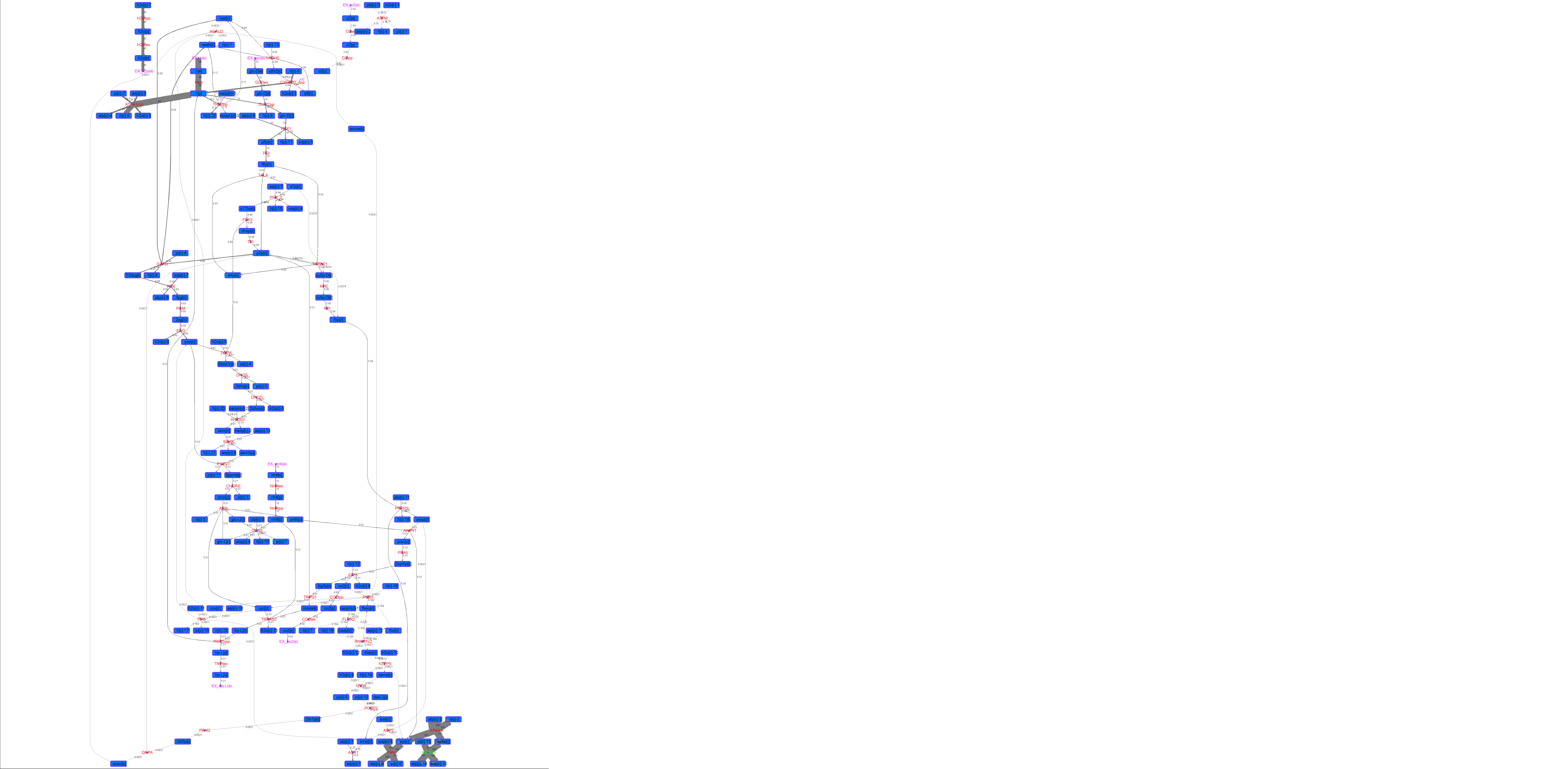

### R00126_Lys.png

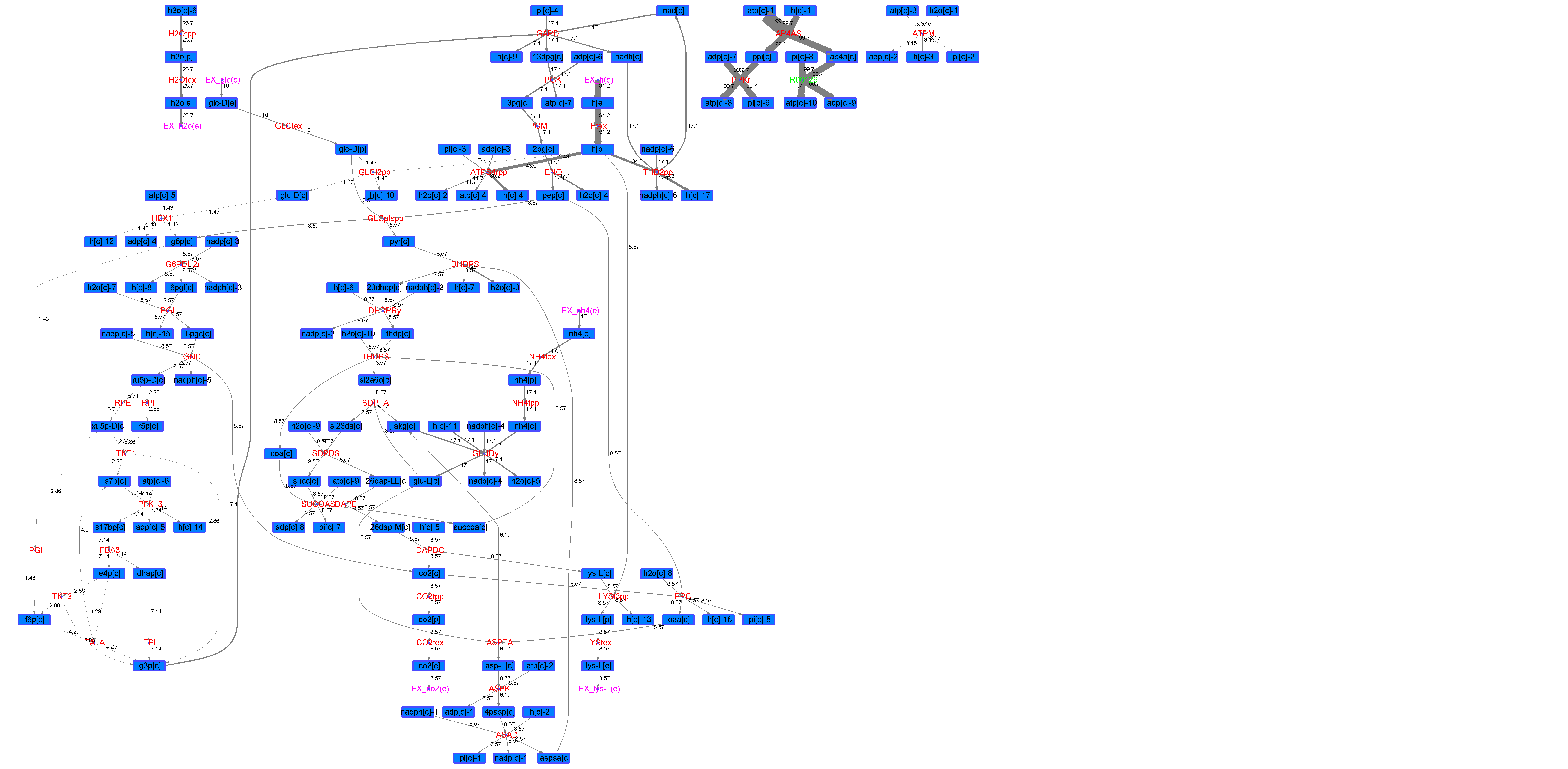

### R00126_Suc.png

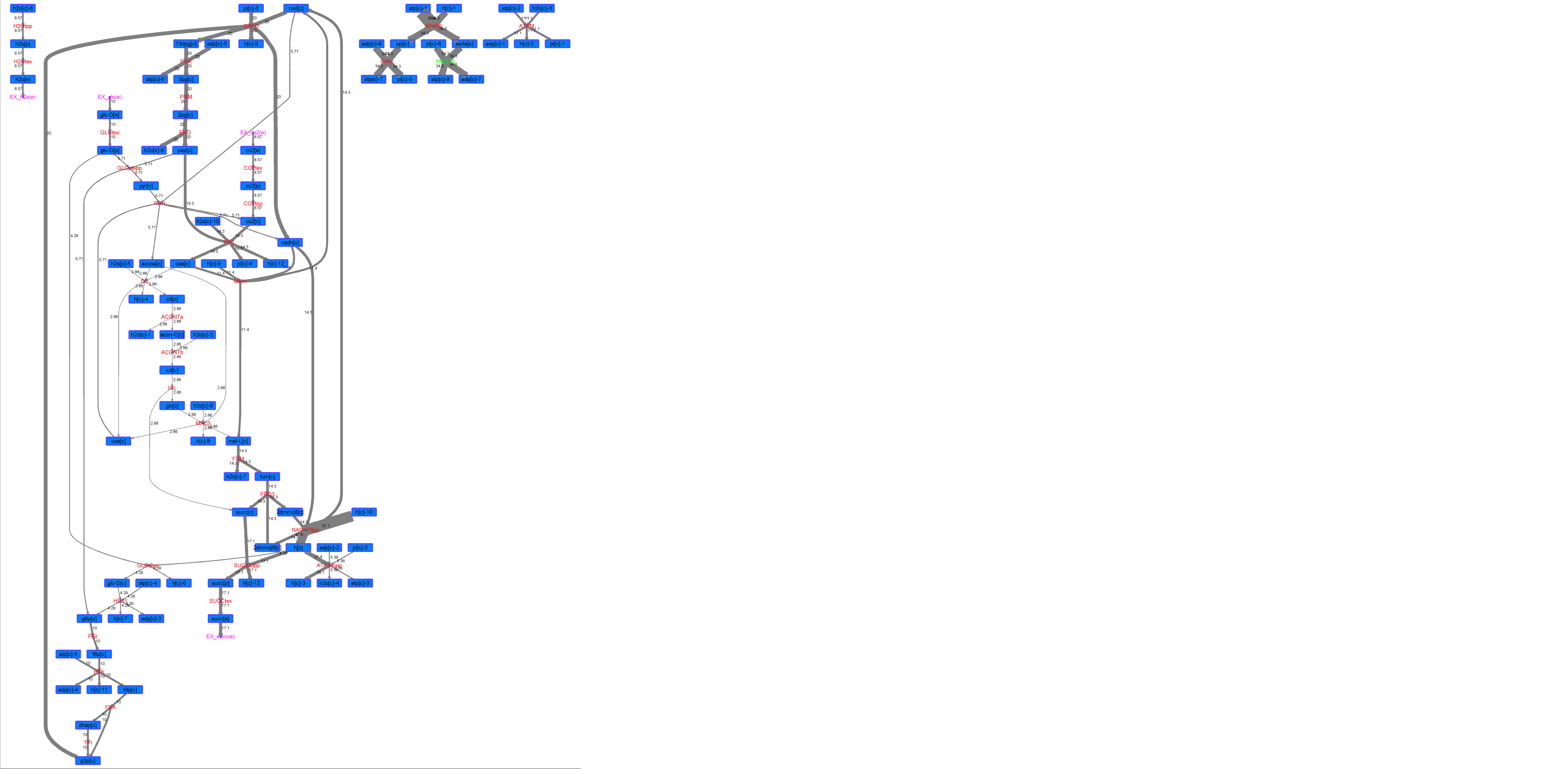

### R00126_Thr.png

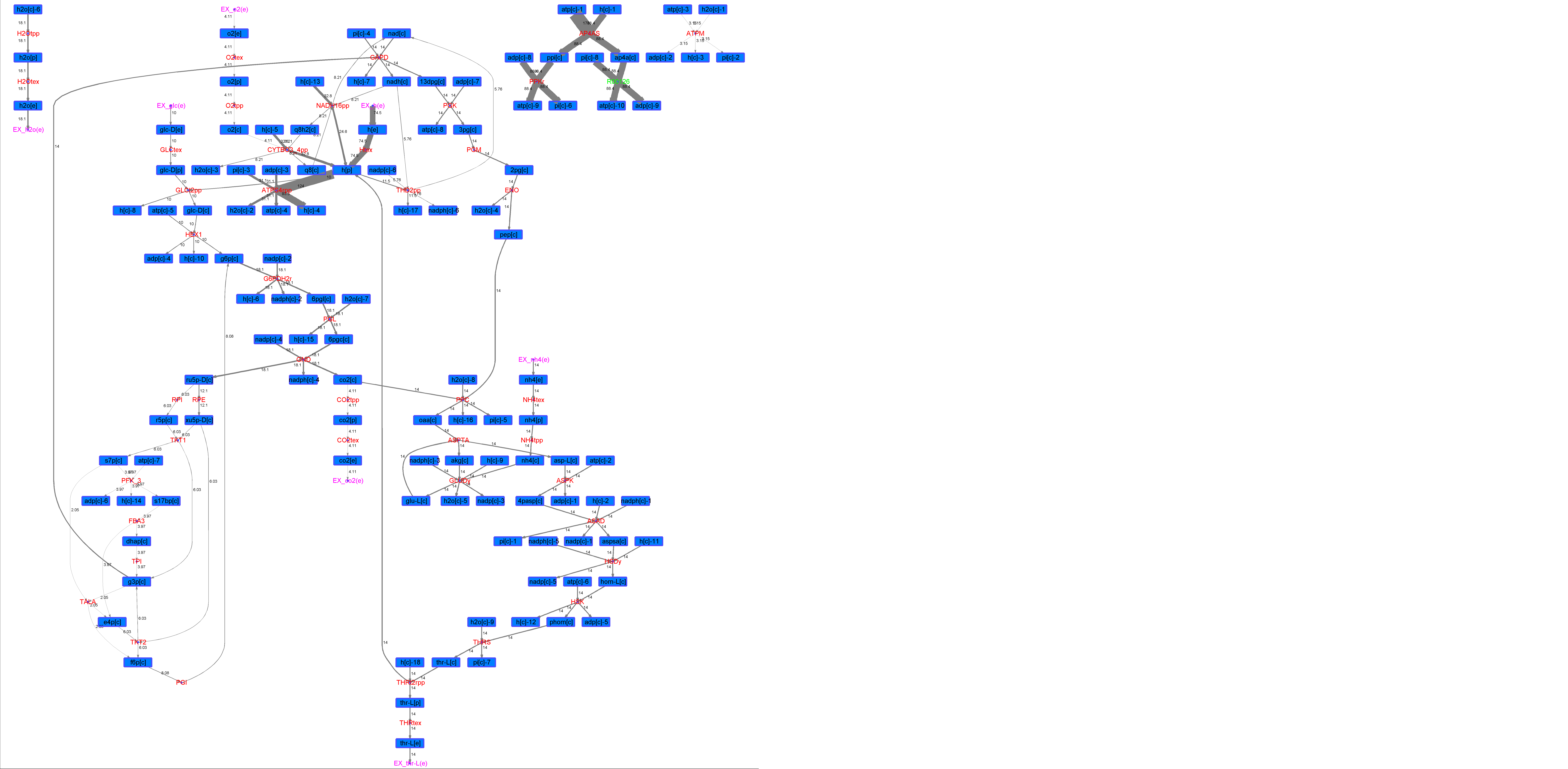

### R00149_Ace.png

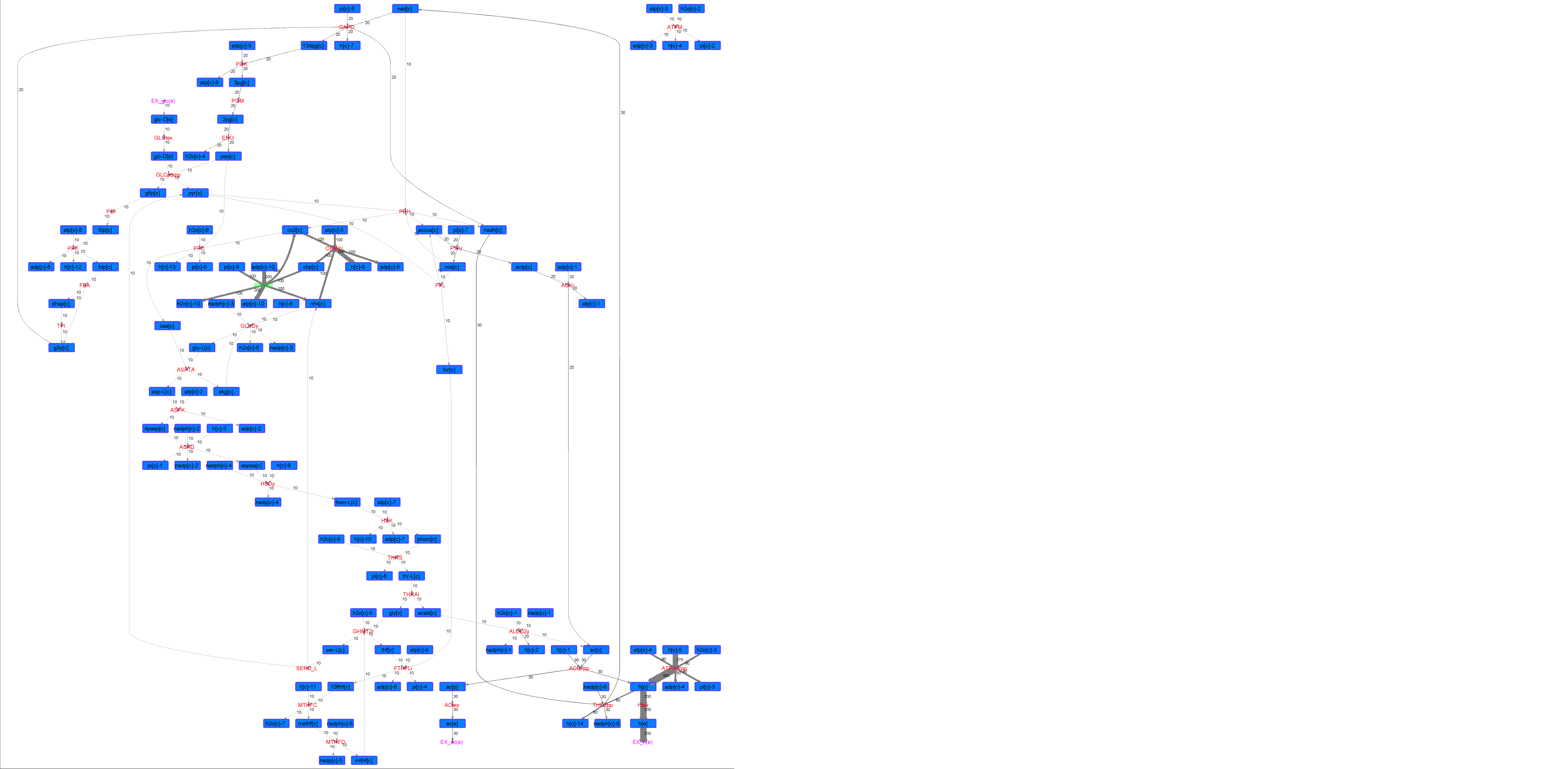

### R00149_For.png

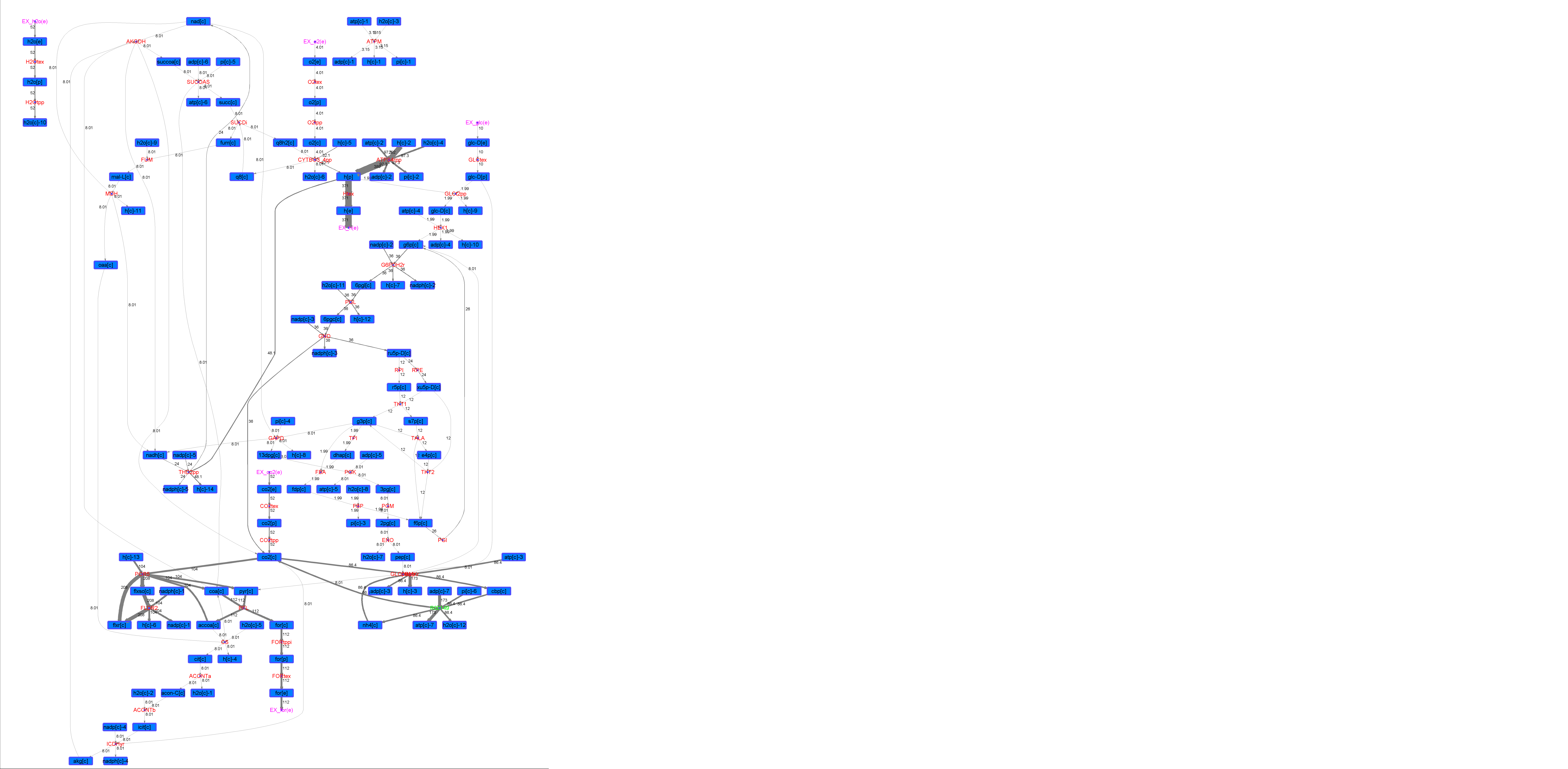

### R00149_Glu.png

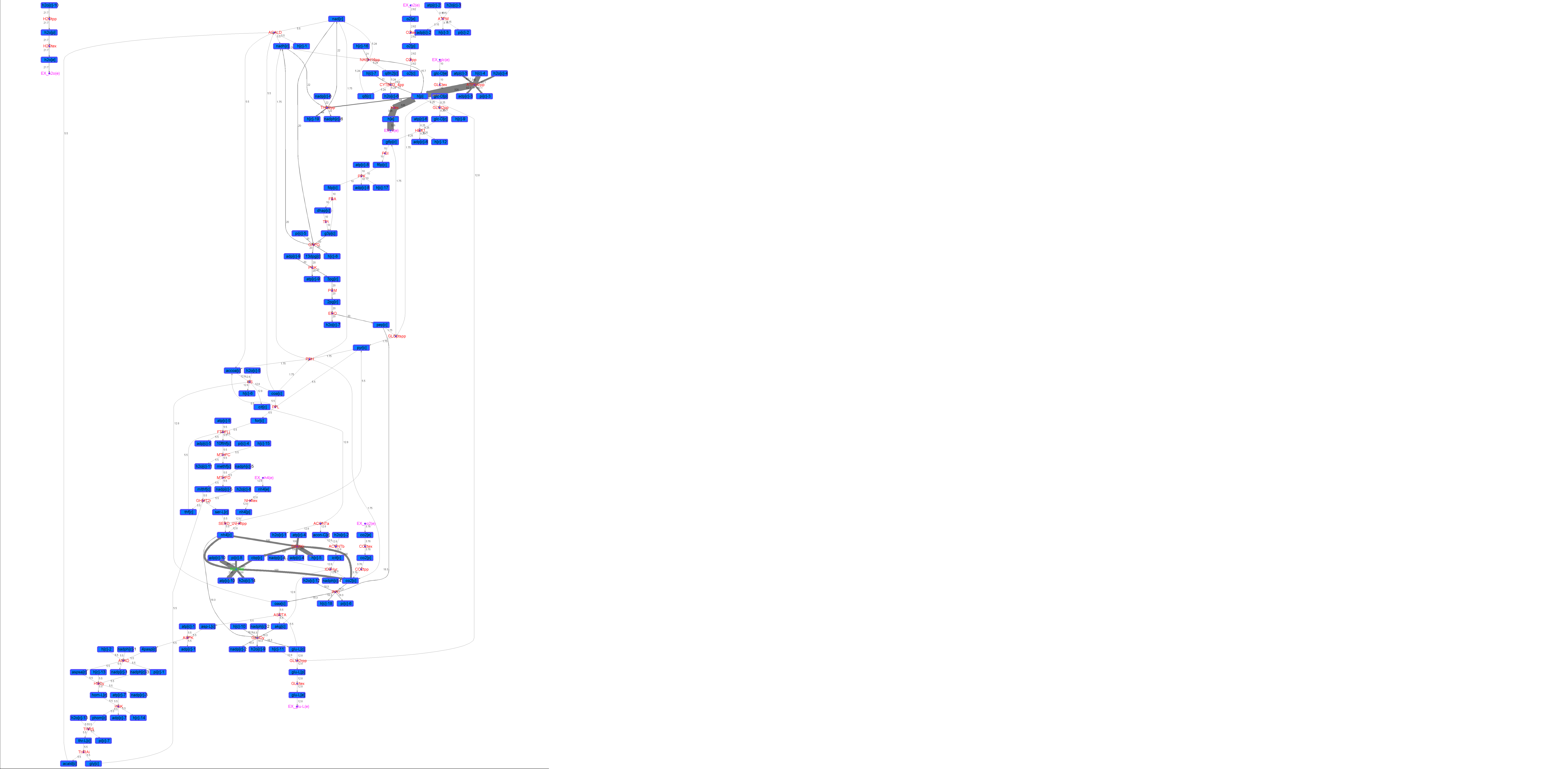

### R00149_L-ala.png

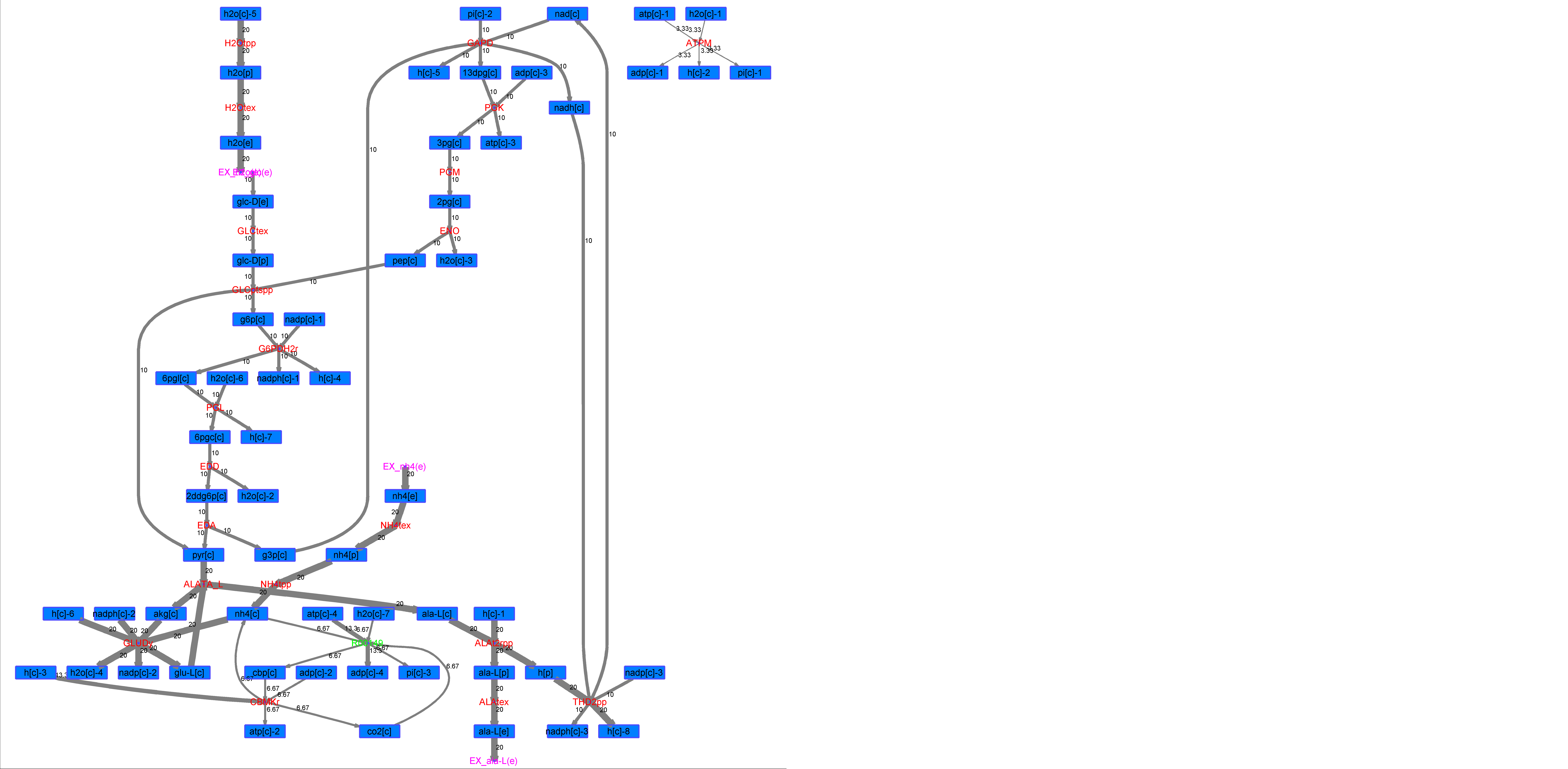

### R00149_L-Mal.png

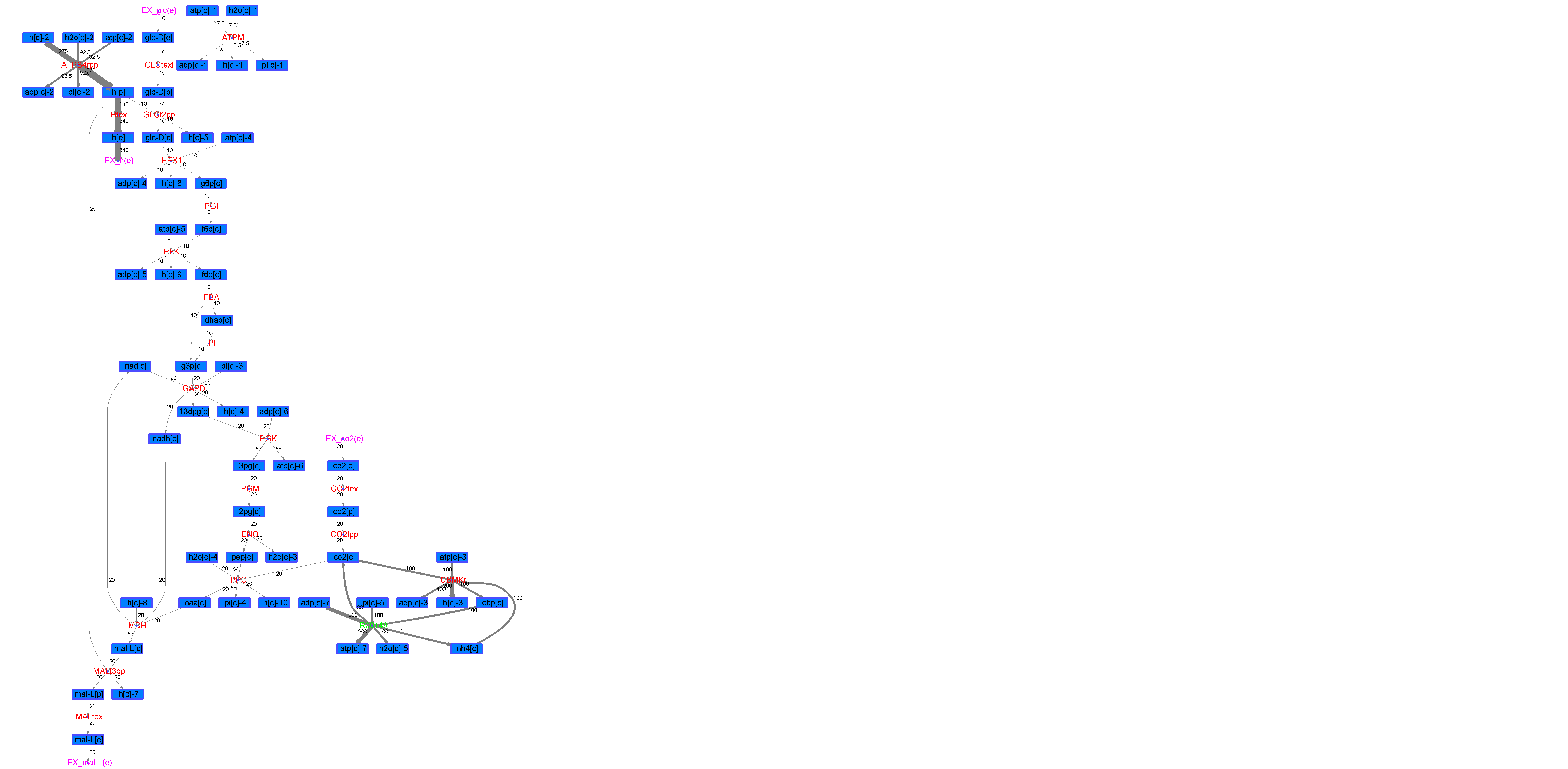

### R00149_L-Phe.png

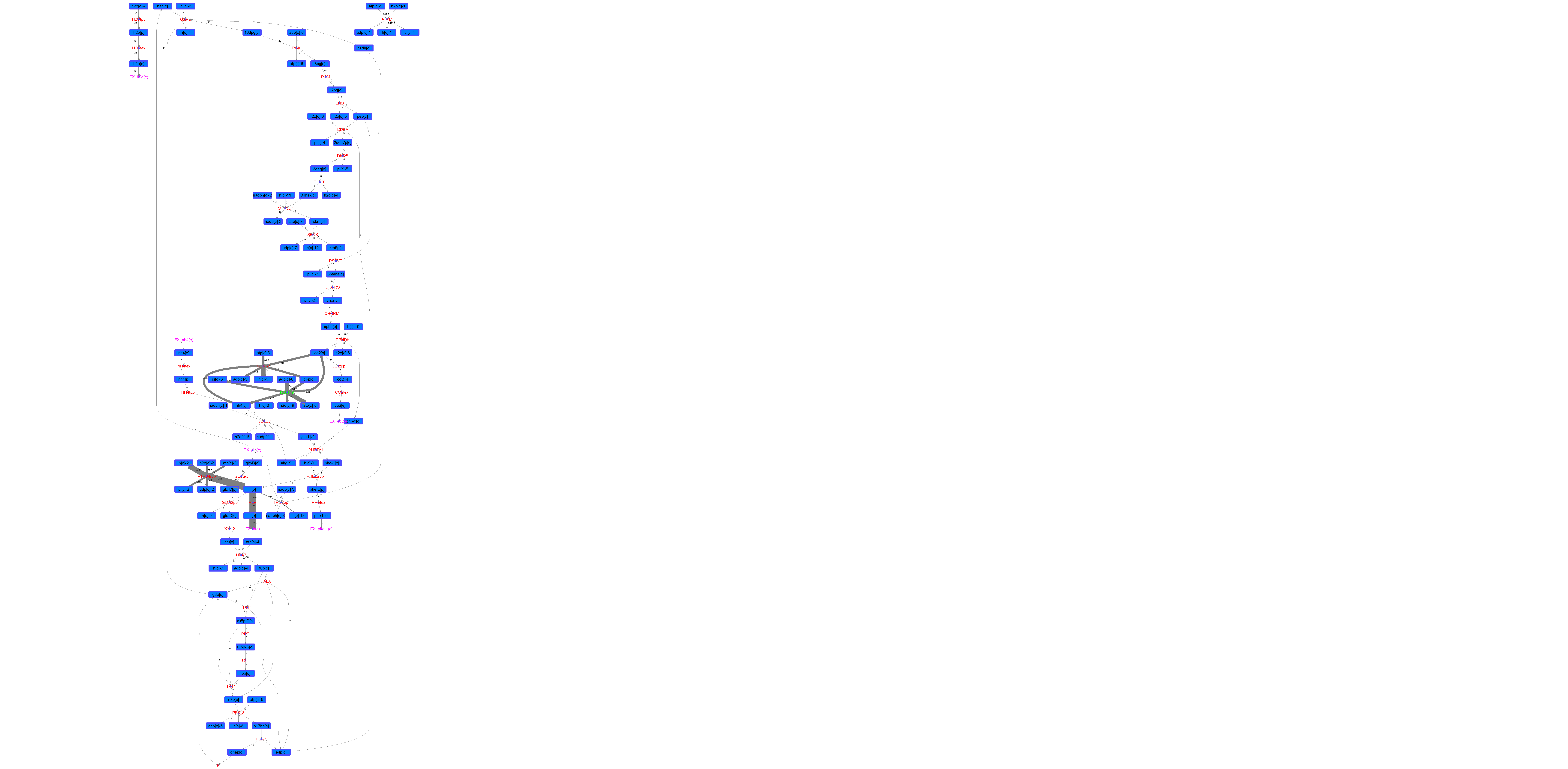

### R00149_L-Try.png

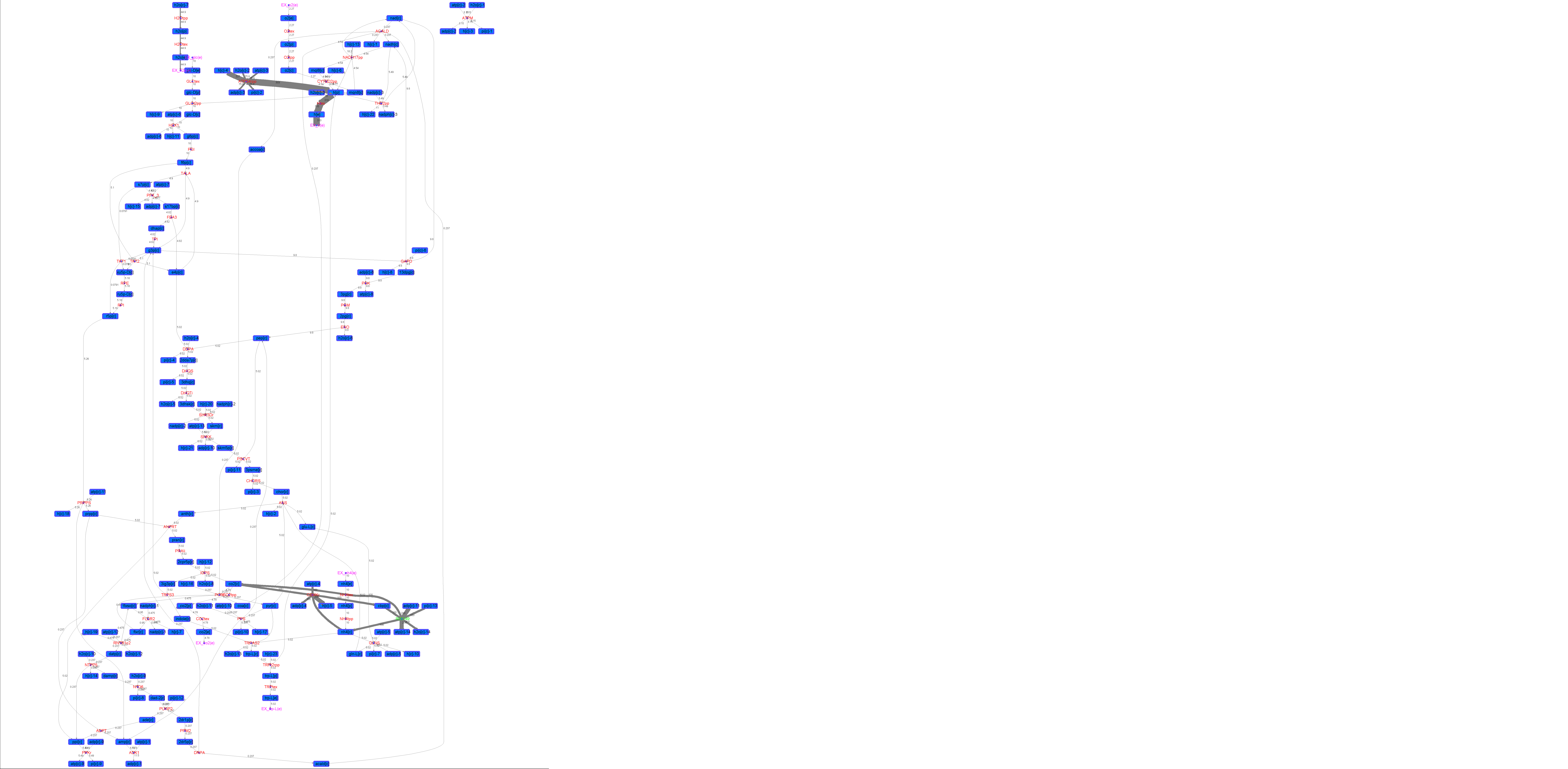

### R00149_Lys.png

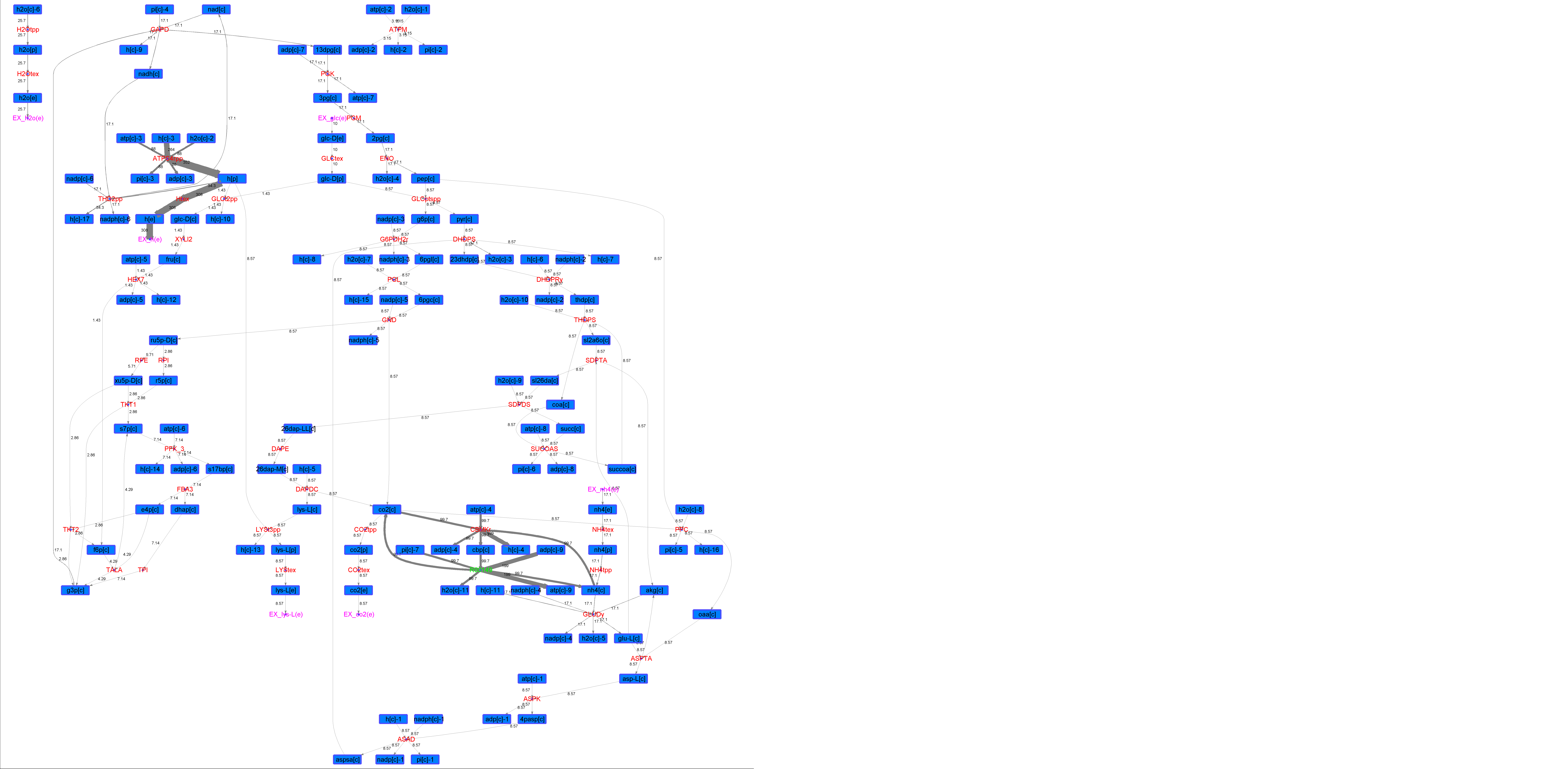

### R00149_Suc.png

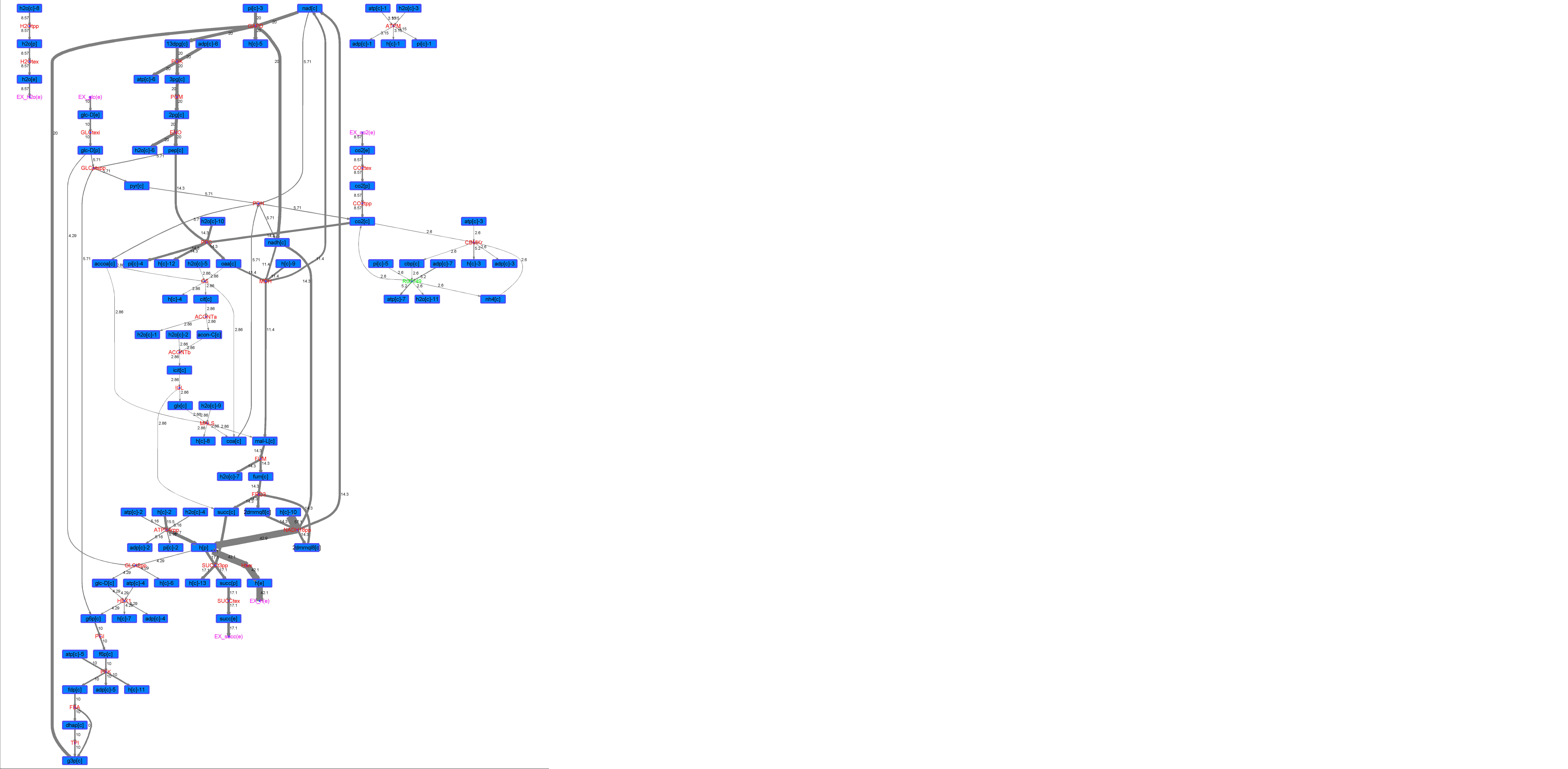

### R00149_Thr.png

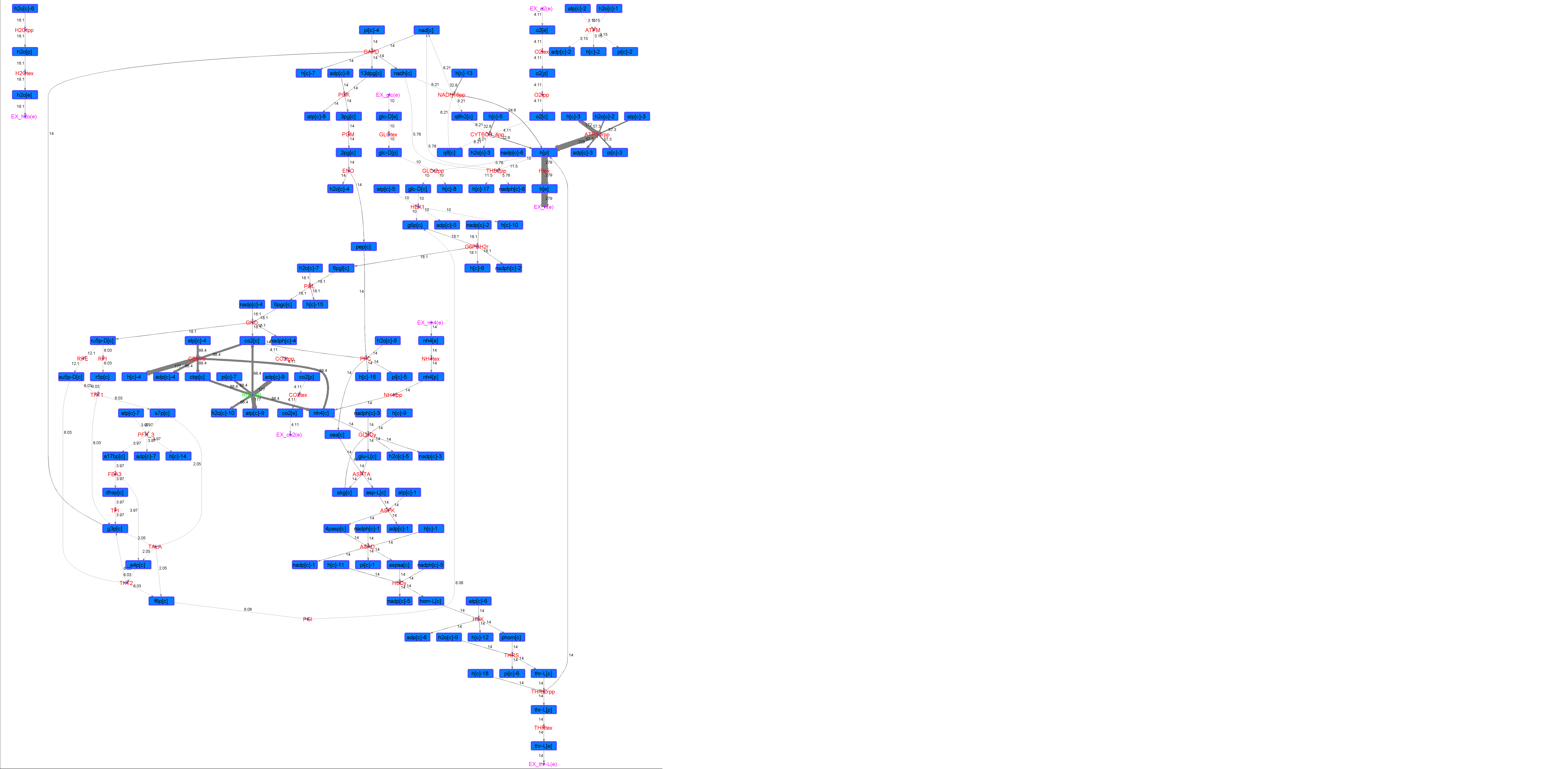

### R00206_Ace.png

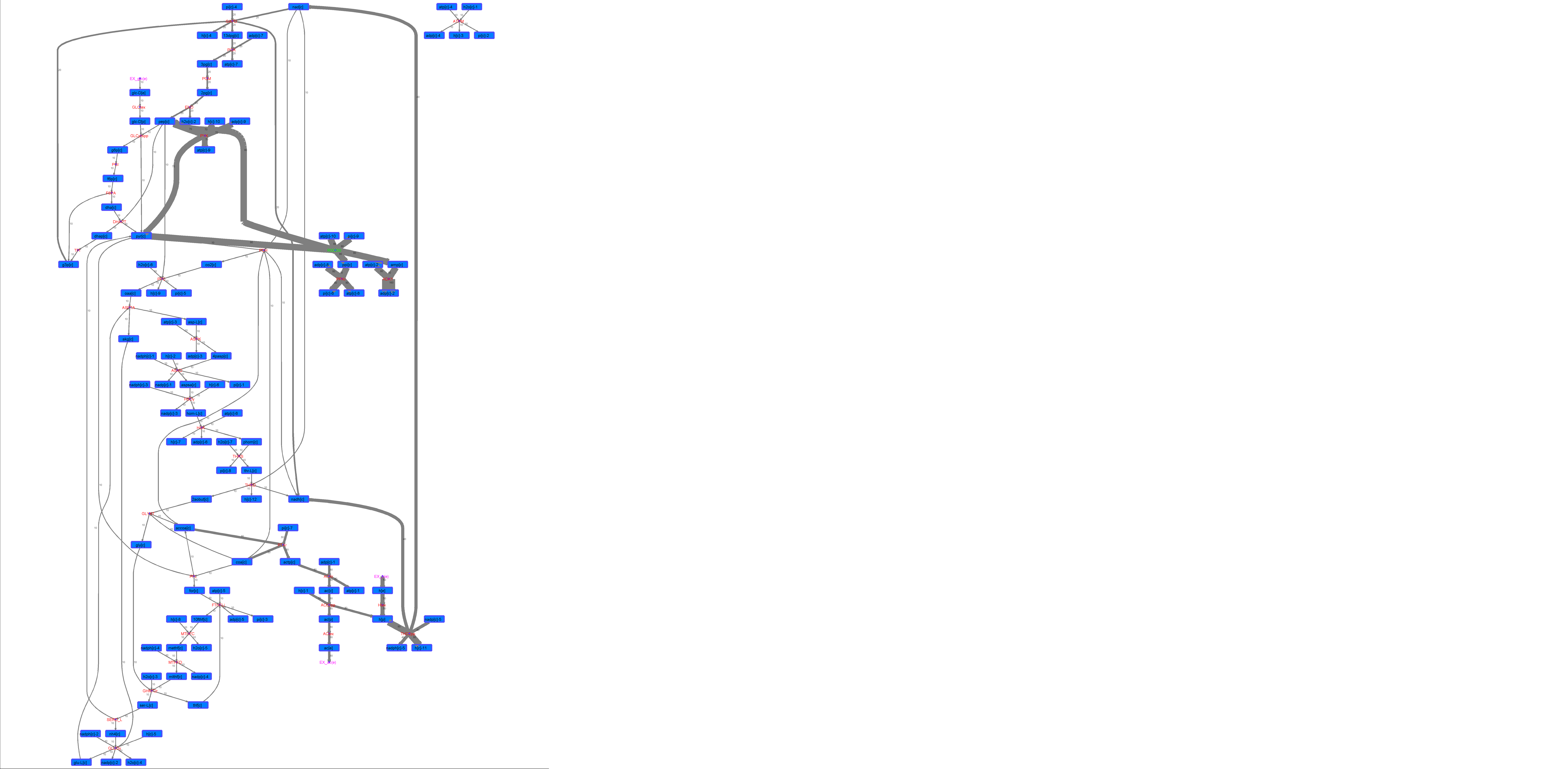

### R00206_For.png

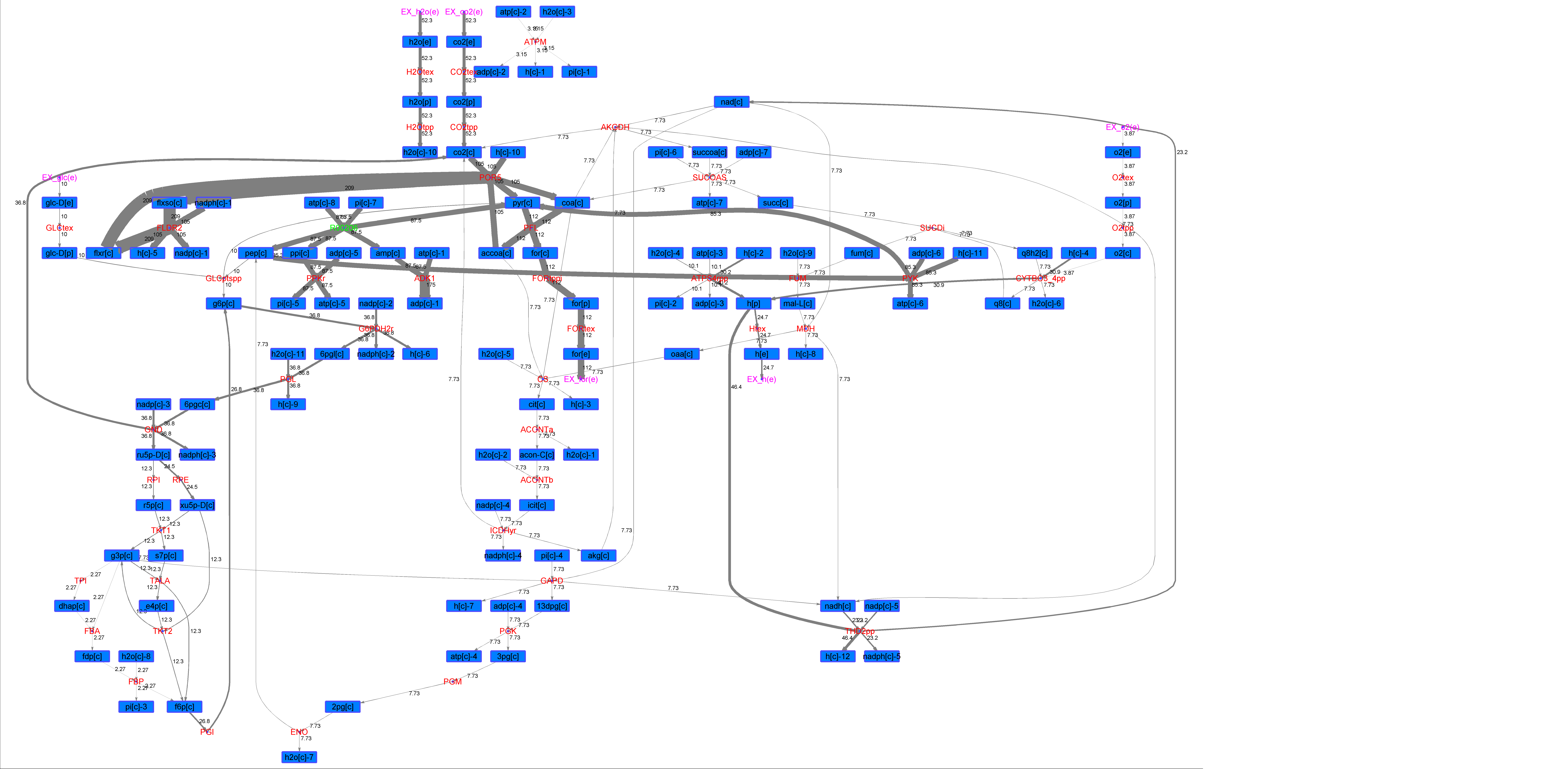

### R00206_Glu.png

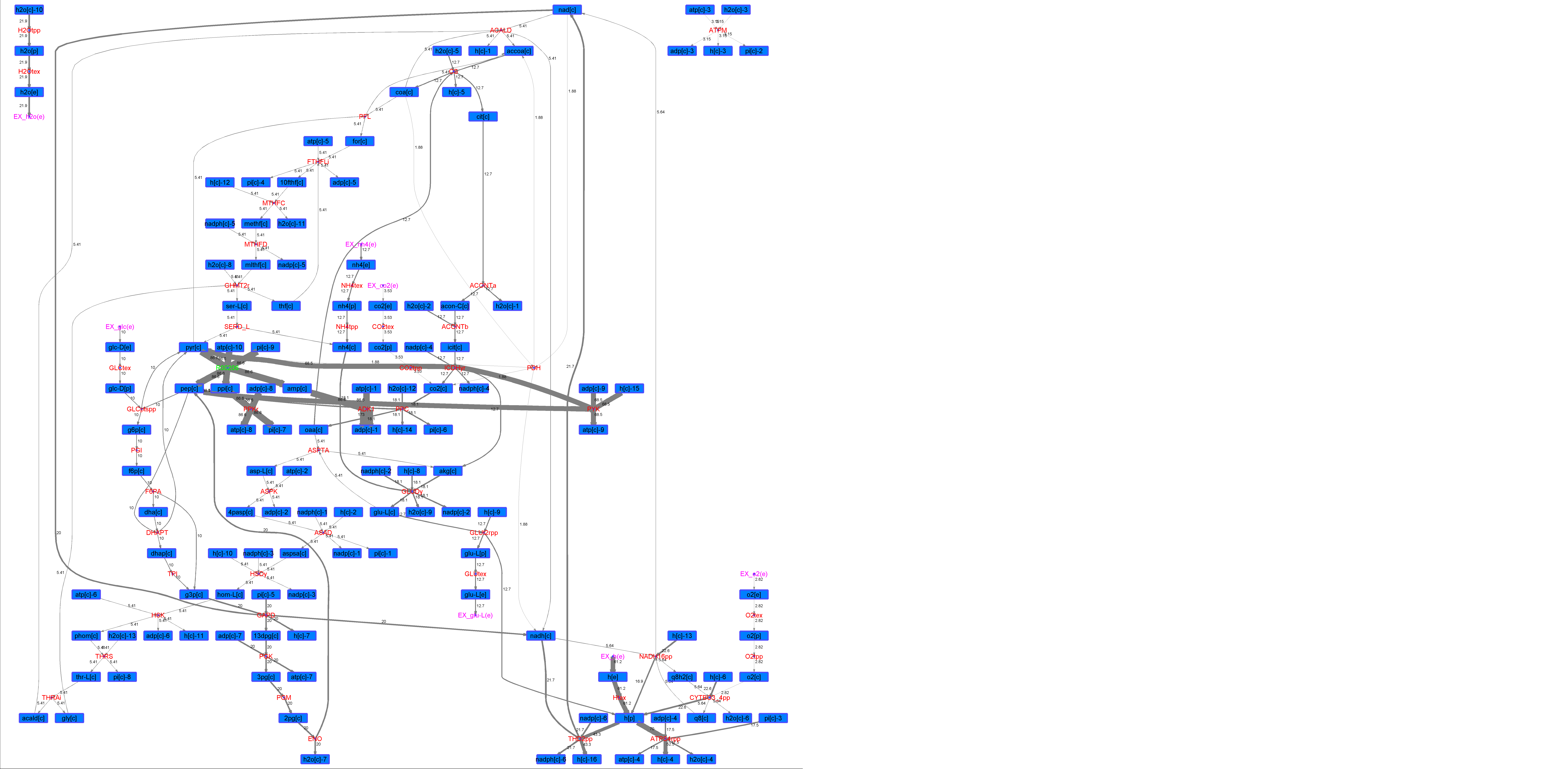

### R00206_L-ala.png

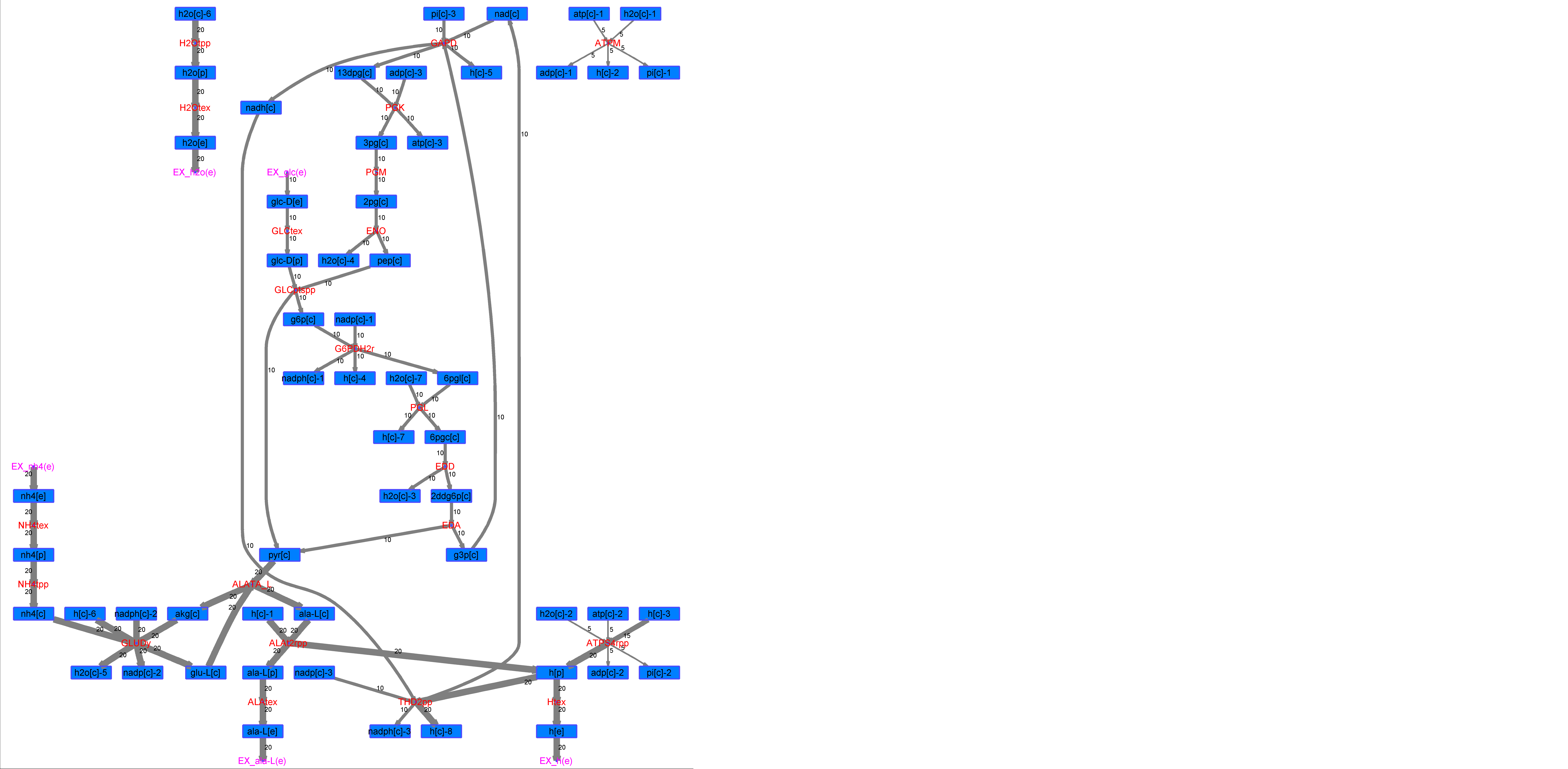

### R00206_L-Mal.png

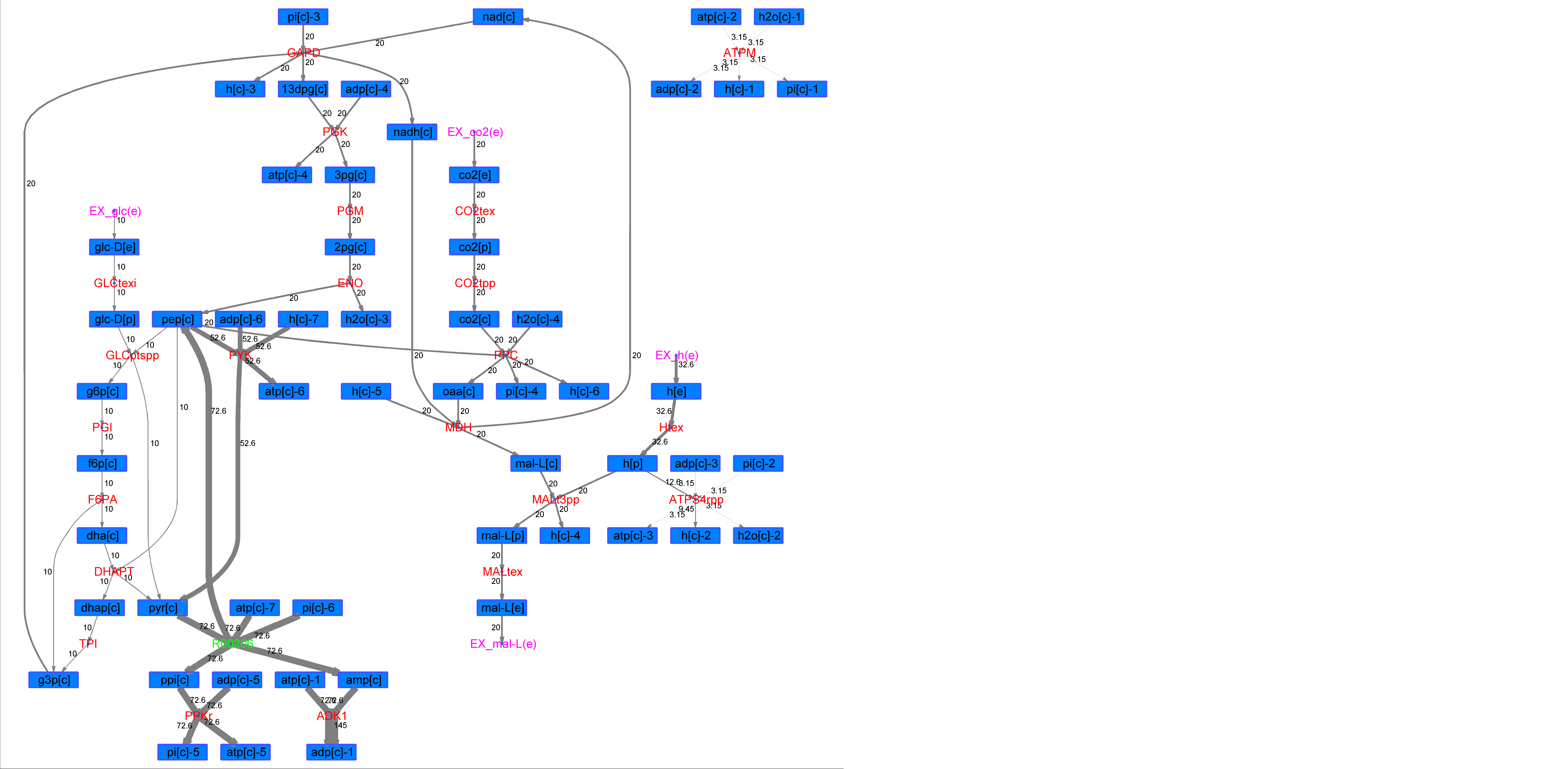

### R00206_L-Phe.png

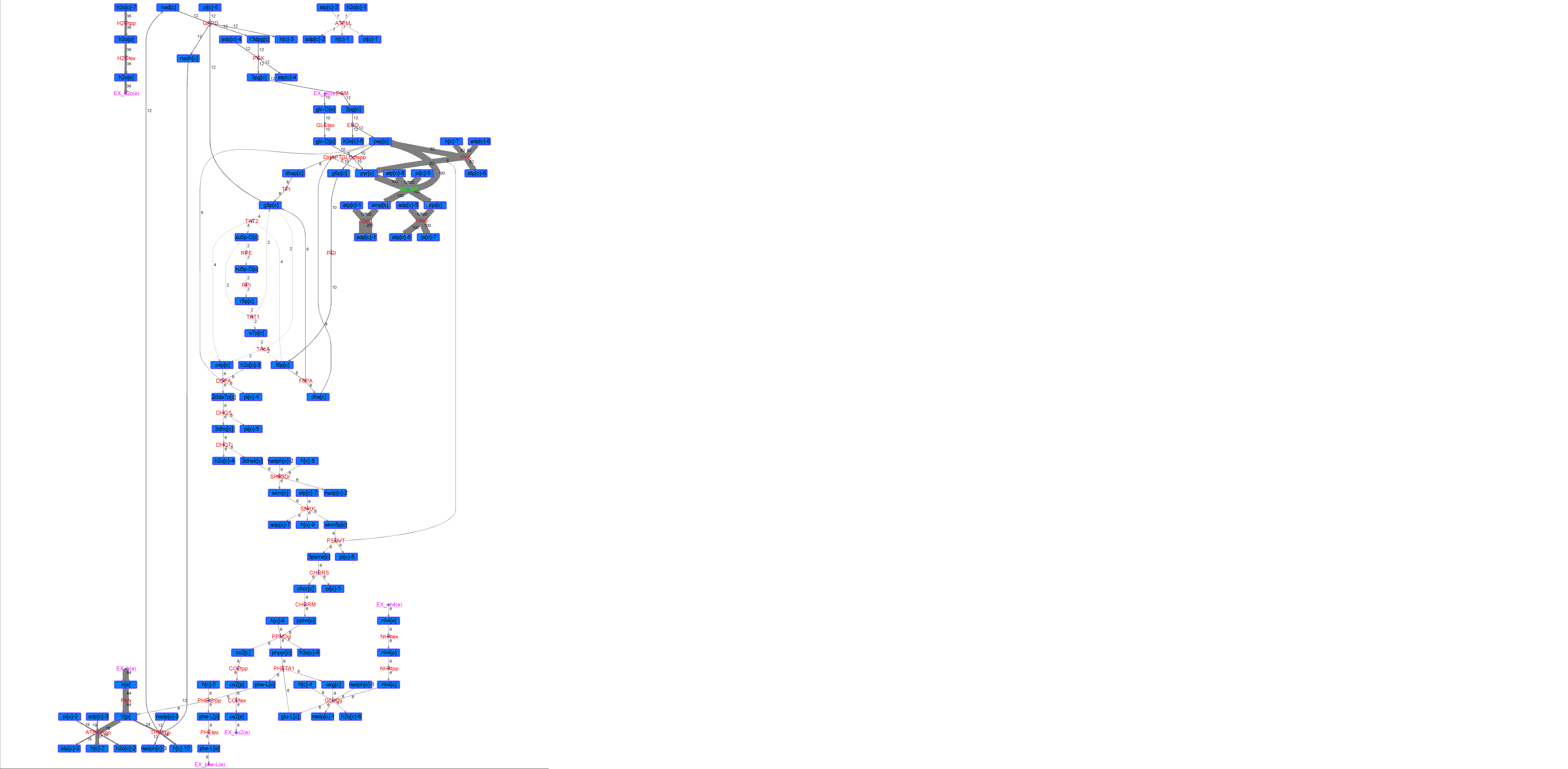

### R00206_L-Try.png

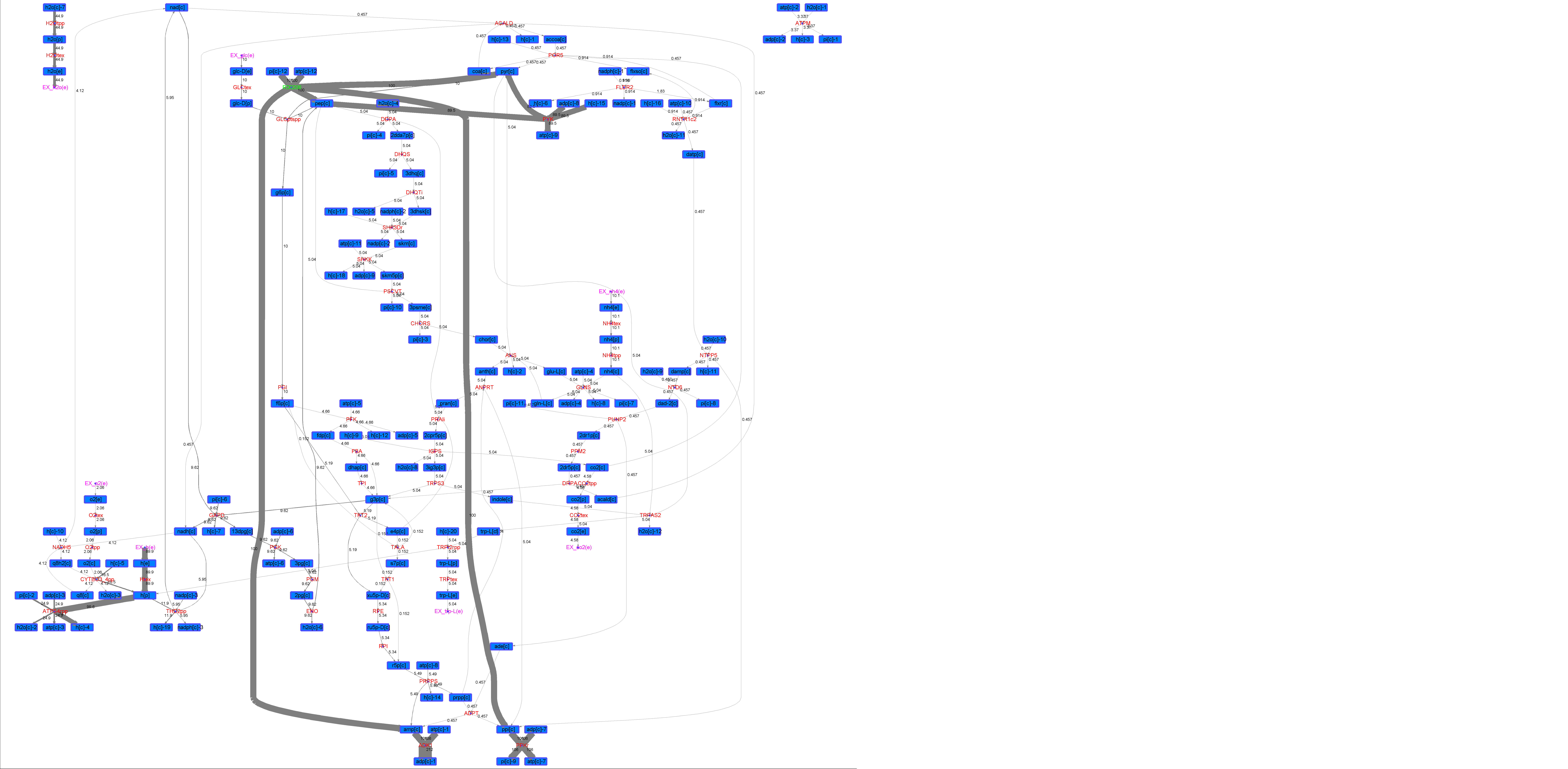

### R00206_Lys.png

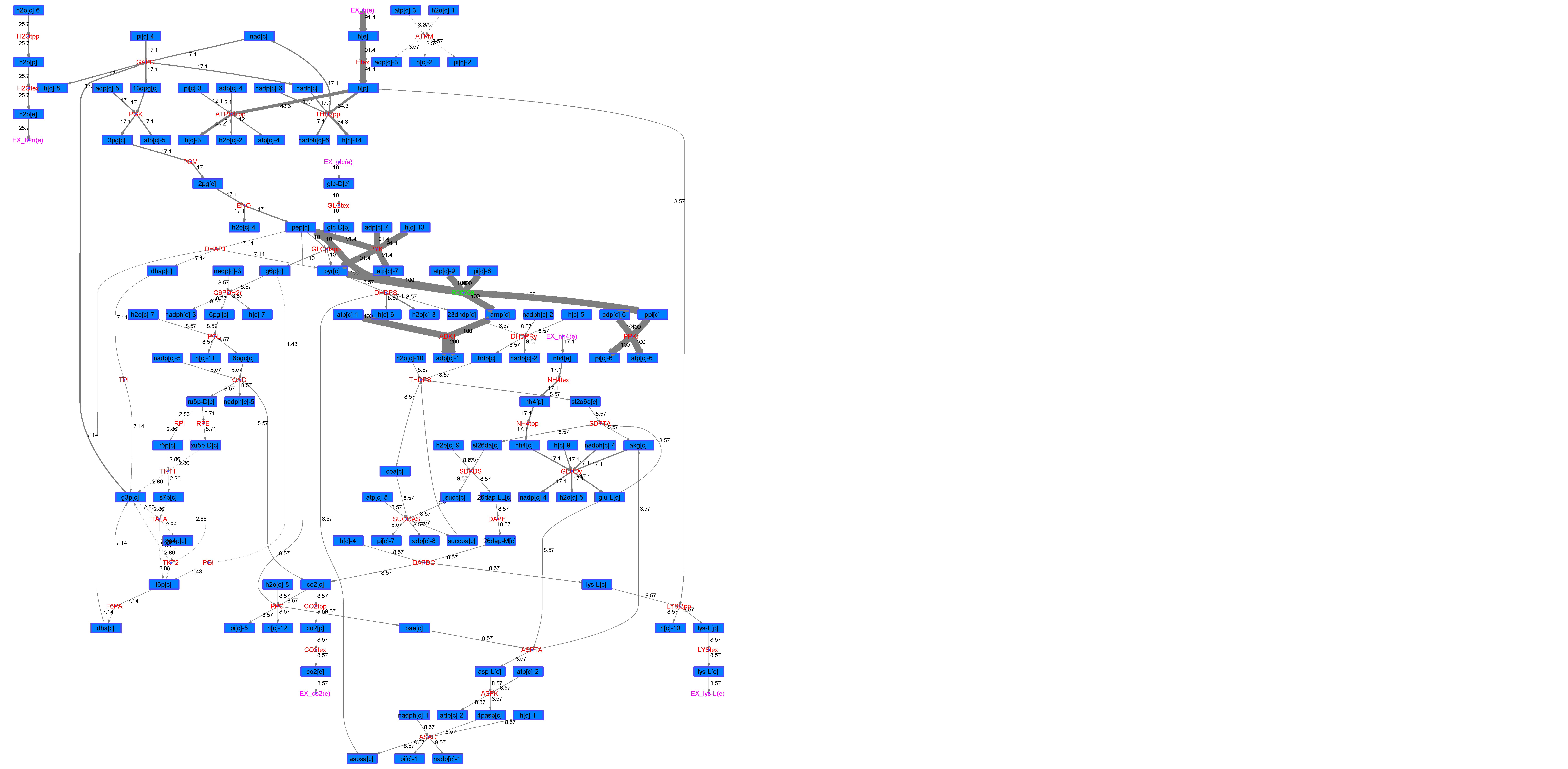

### R00206_Suc.png

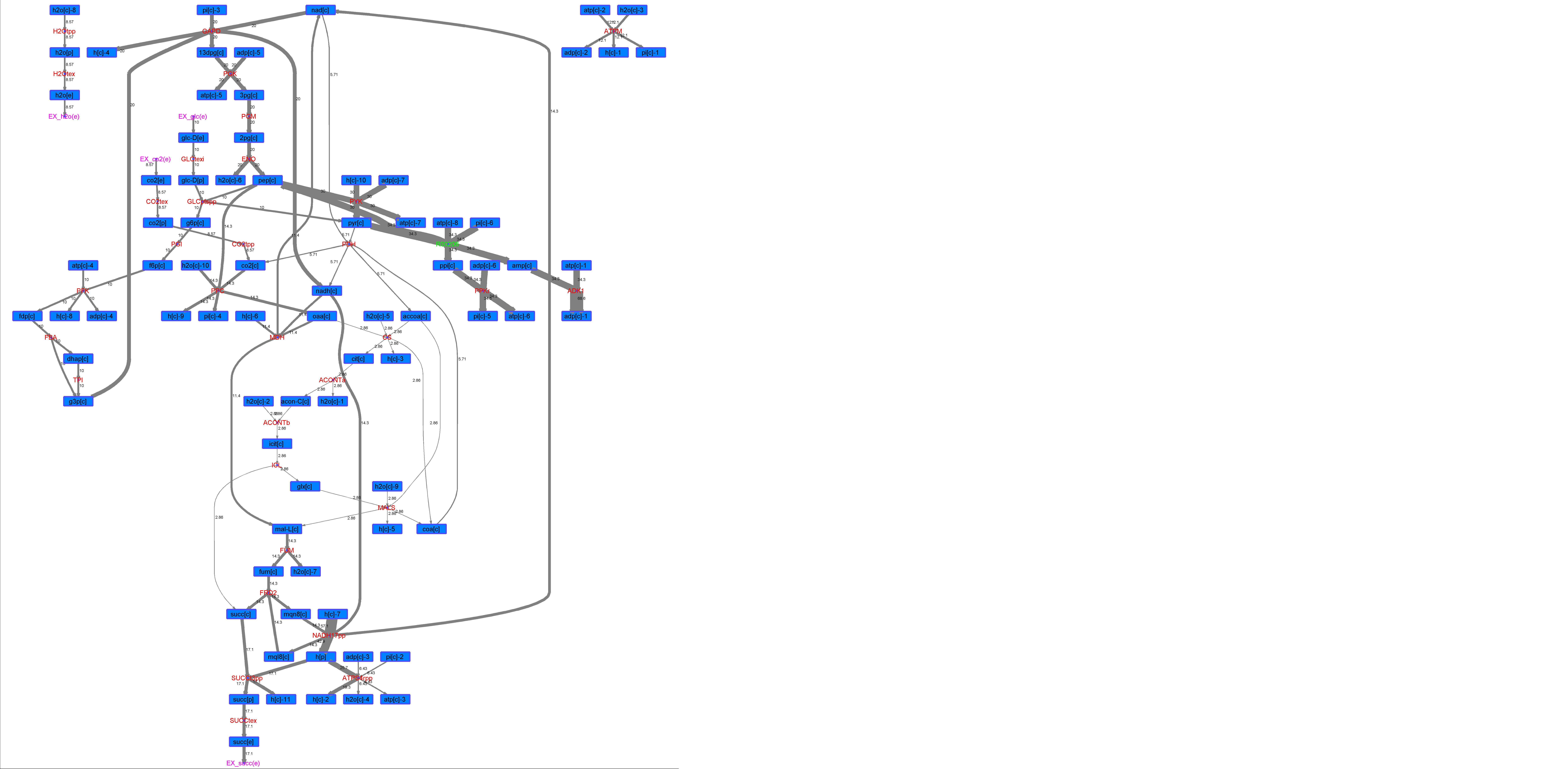

### R00206_Thr.png

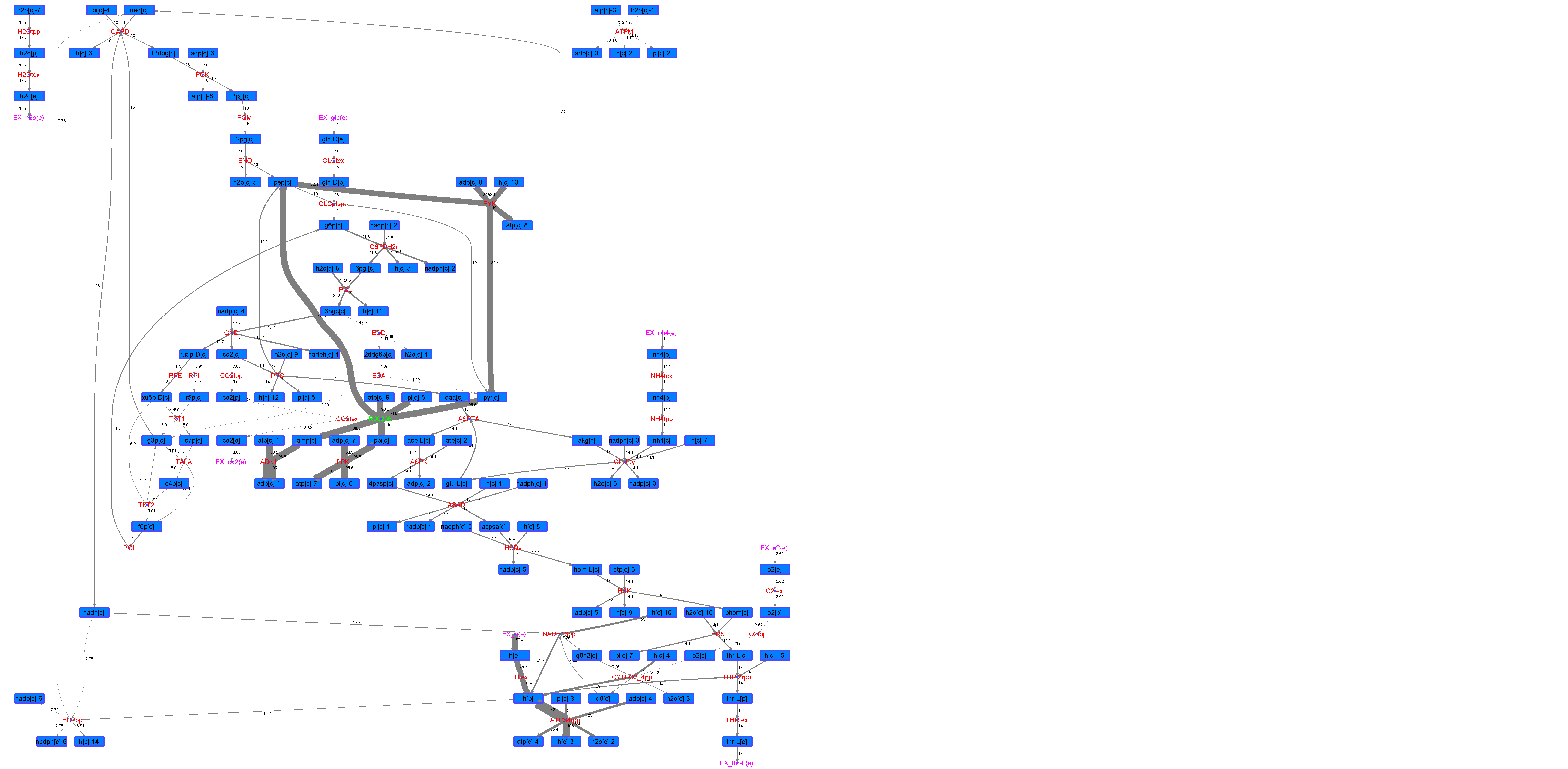

### R00210_Ace.png

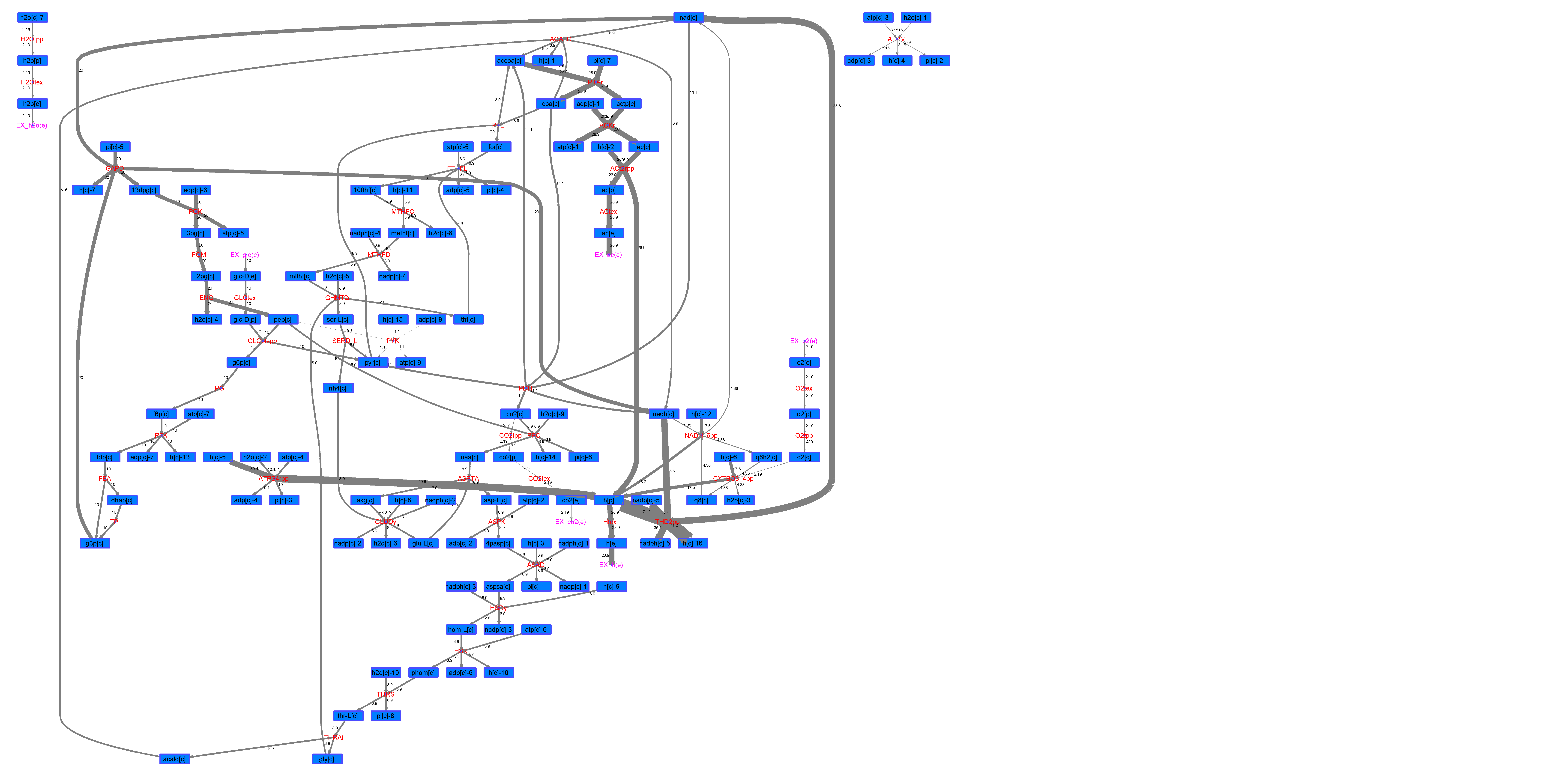

### R00210_For.png

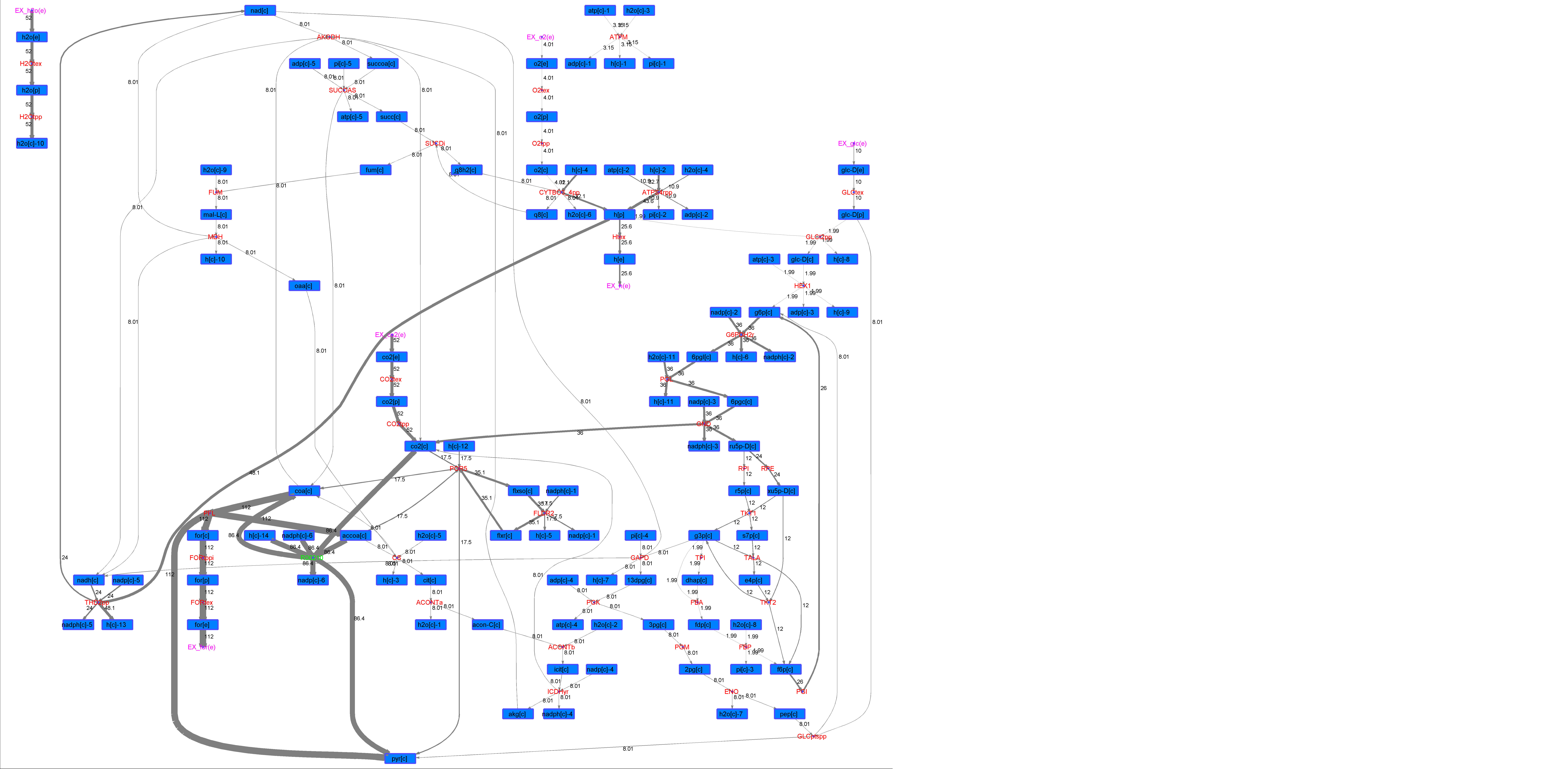

### R00210_Glu.png

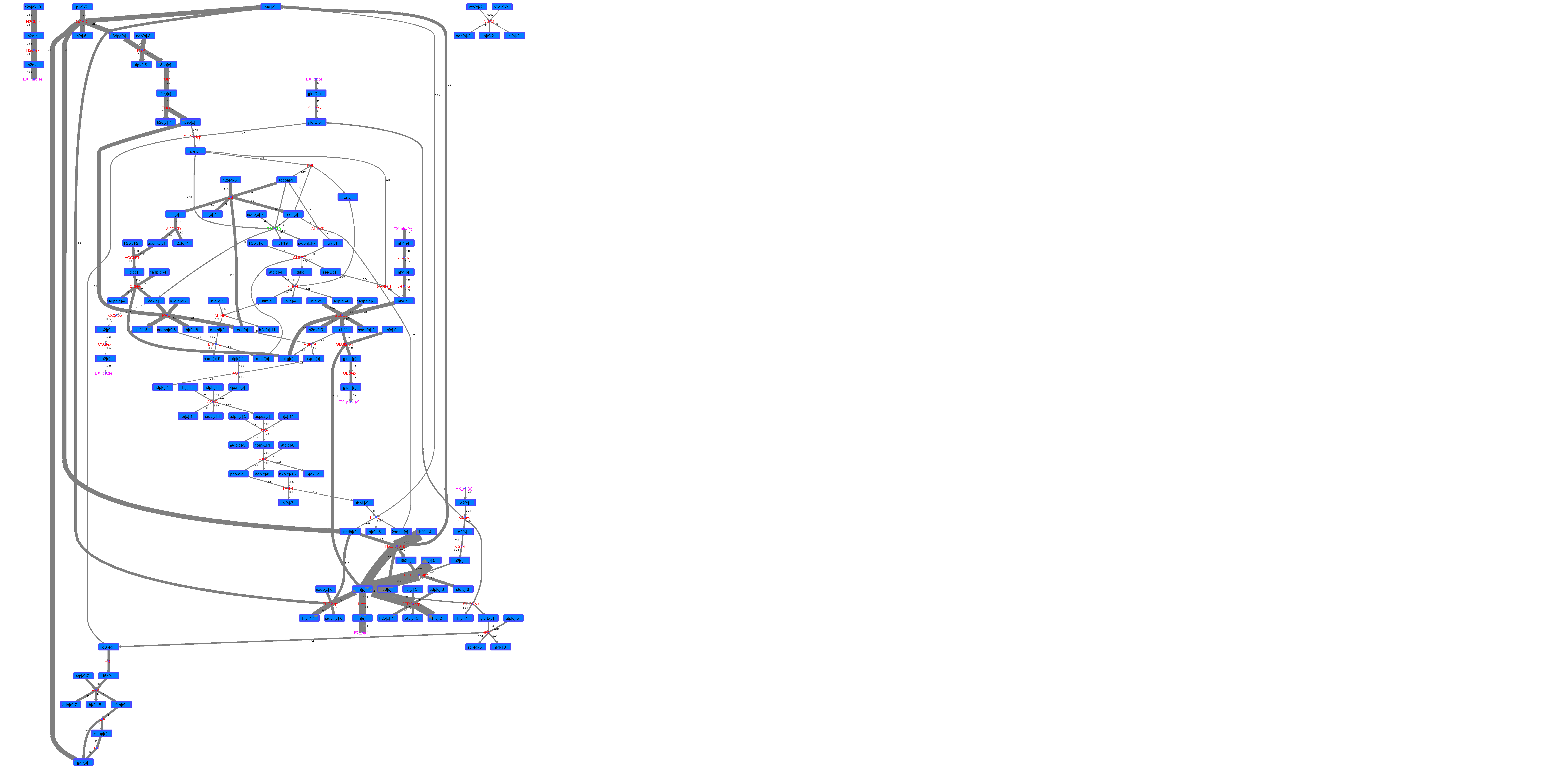

### R00210_L-ala.png

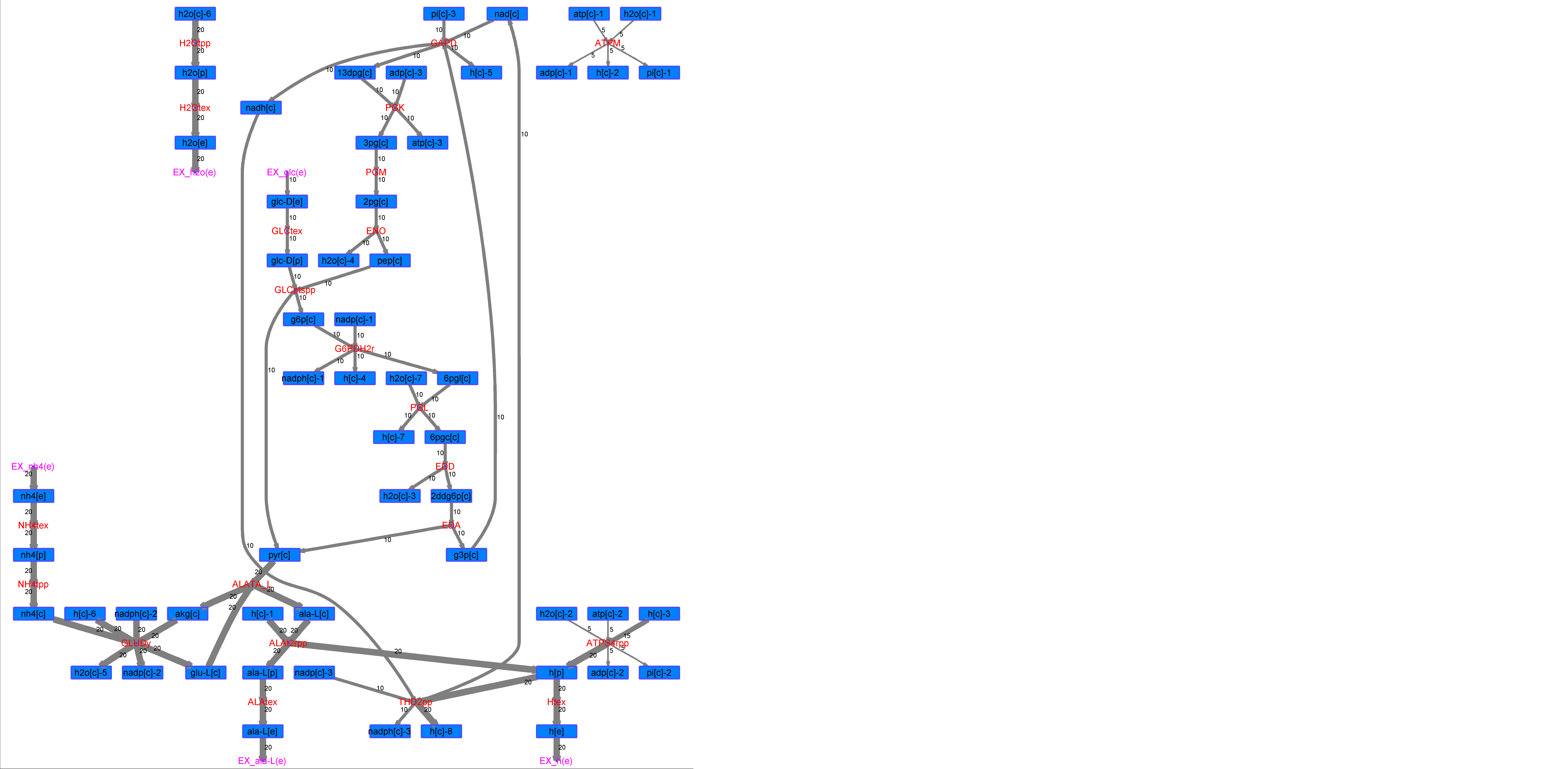

### R00210_L-Mal.png

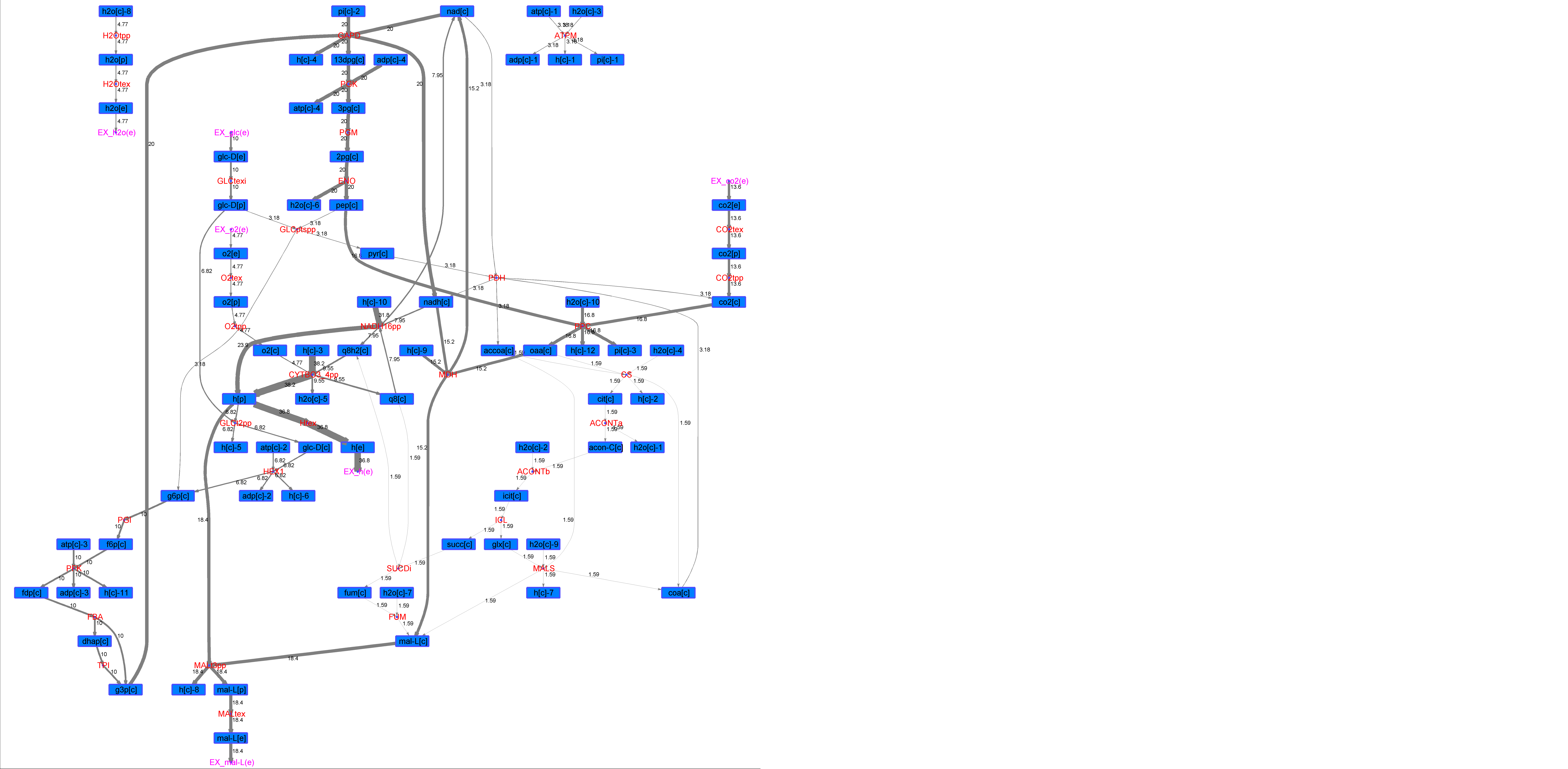

### R00210_L-Phe.png

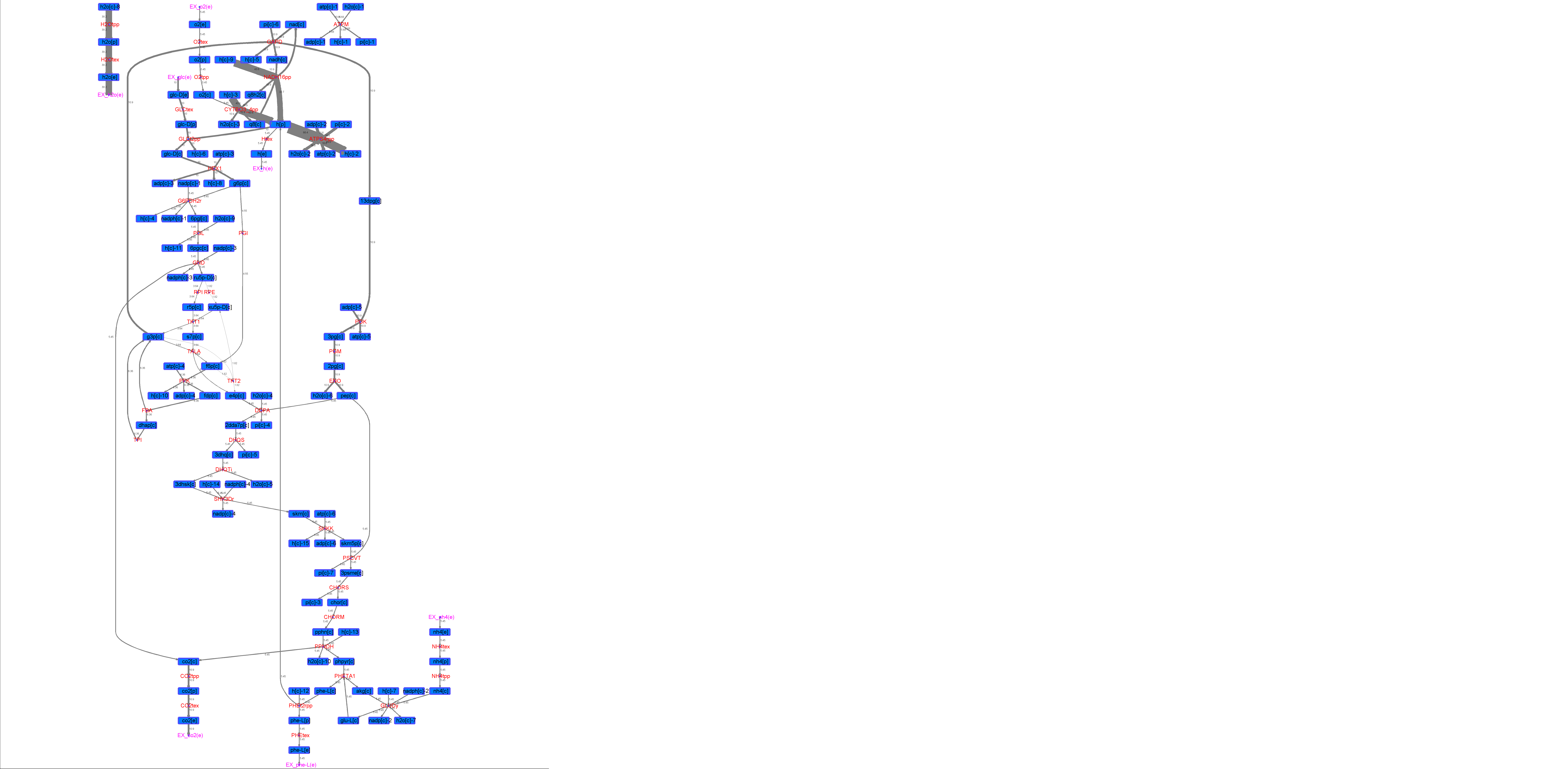

### R00210_L-Try.png

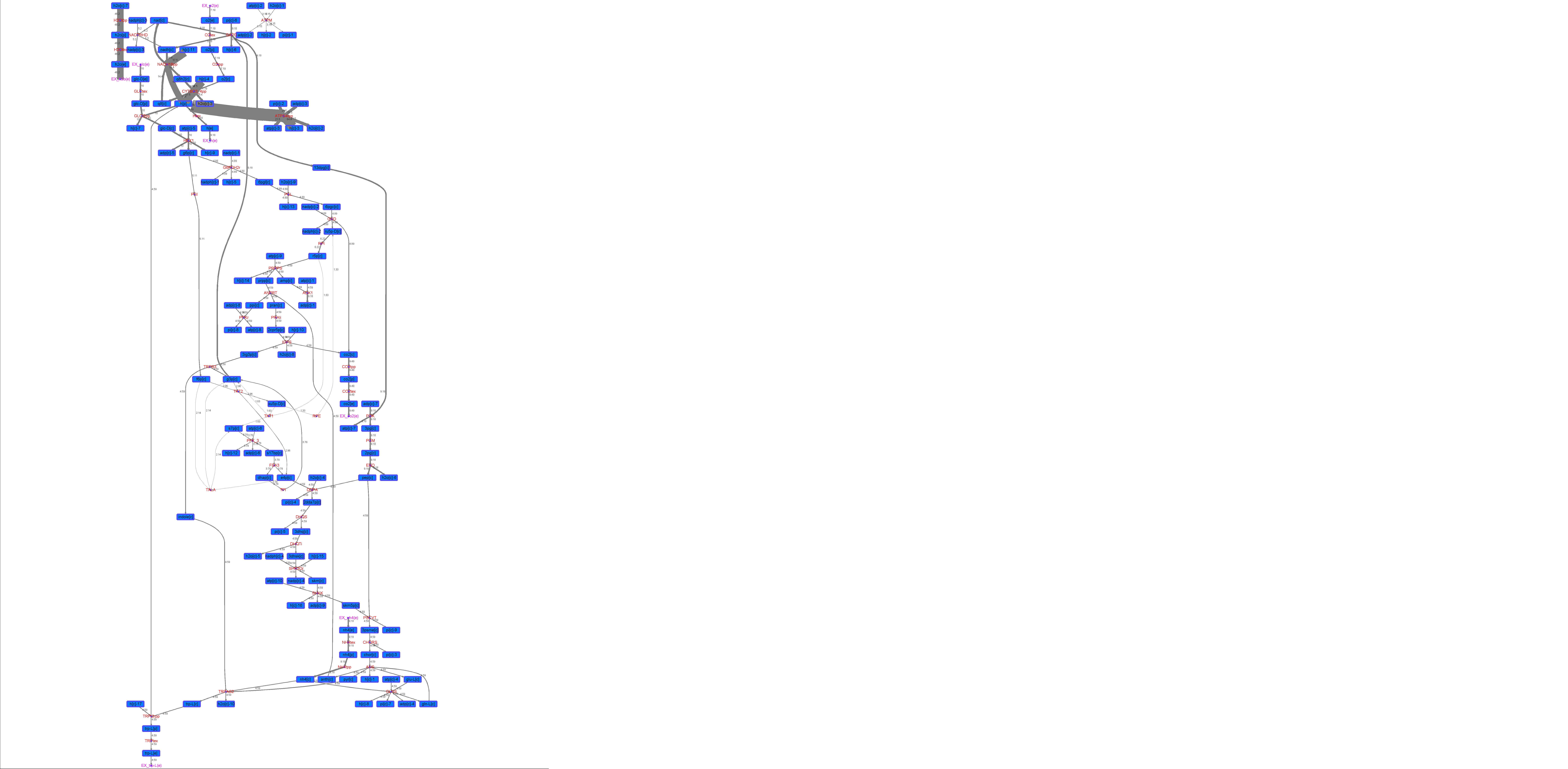

### R00210_Lys.png

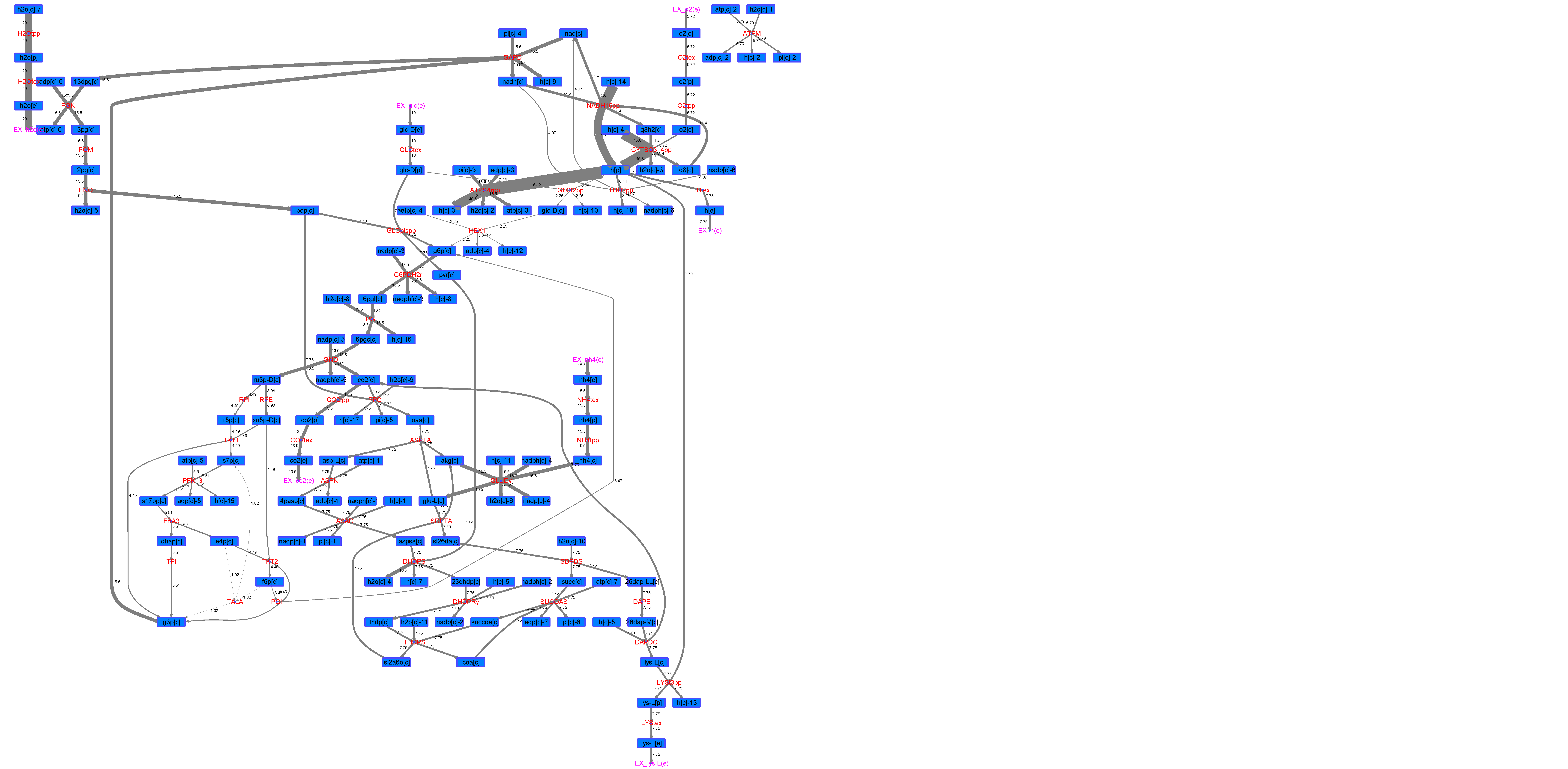

### R00210_Suc.png

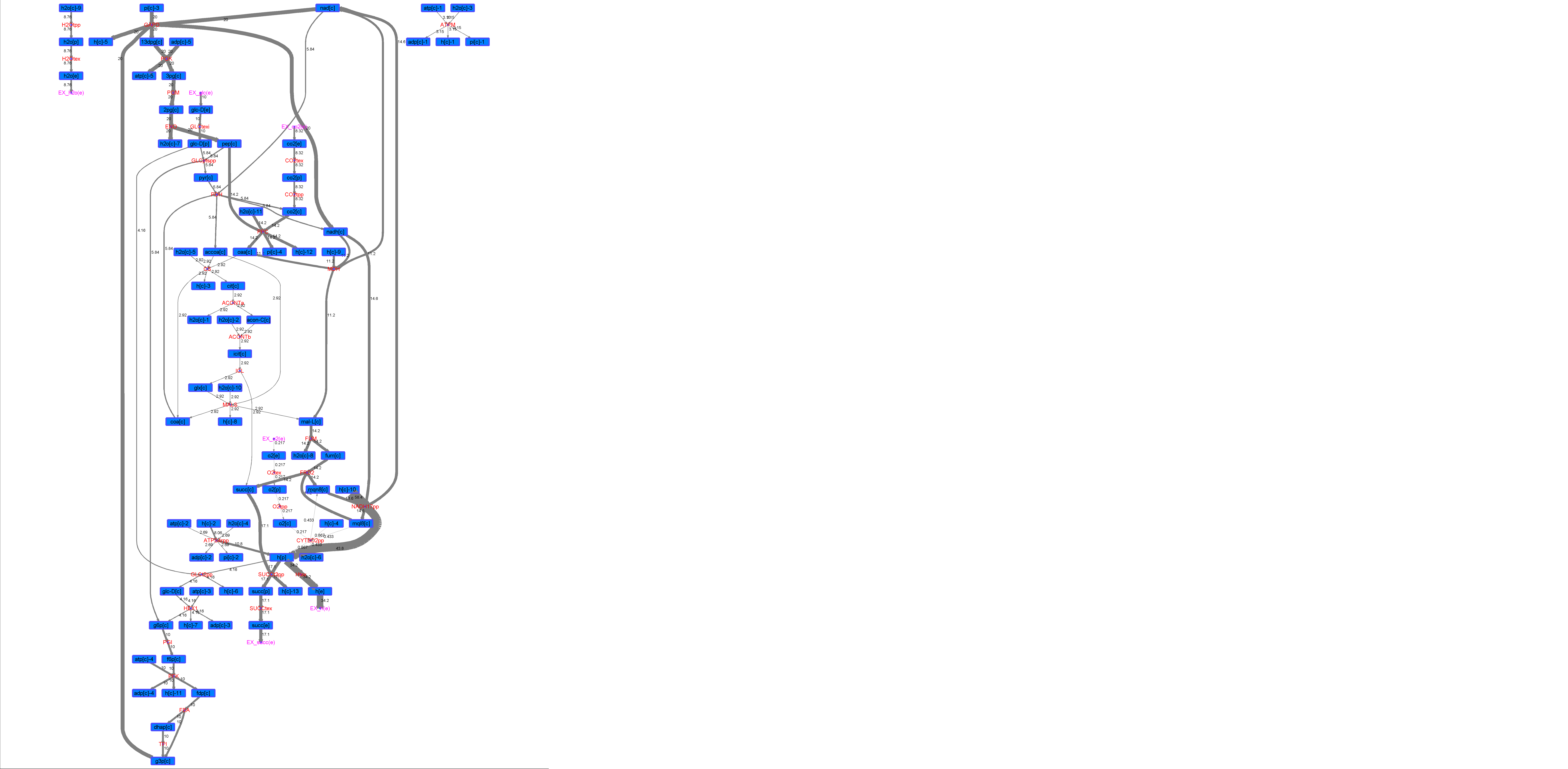

### R00210_Thr.png

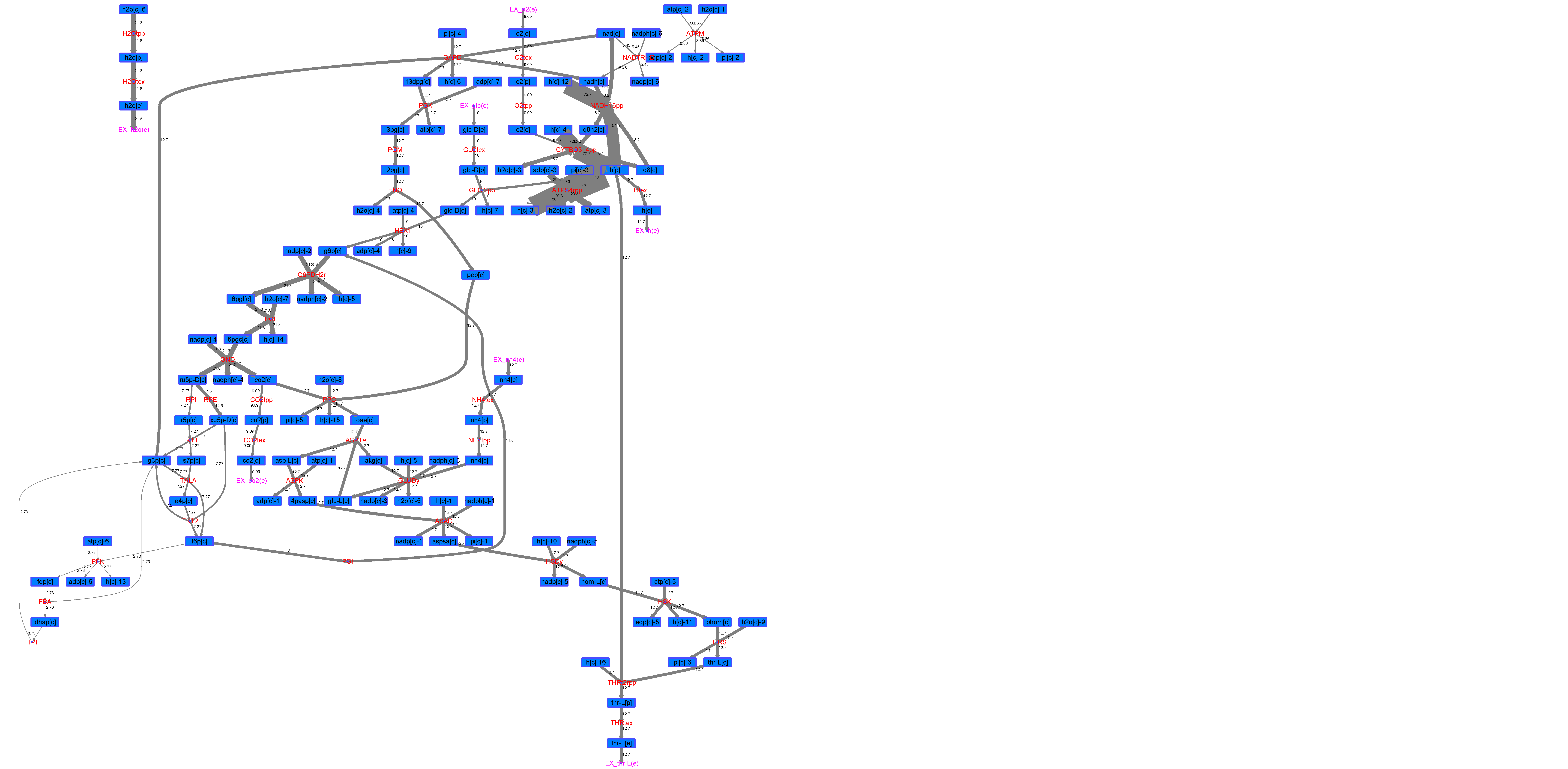

### R00224_Ace.png

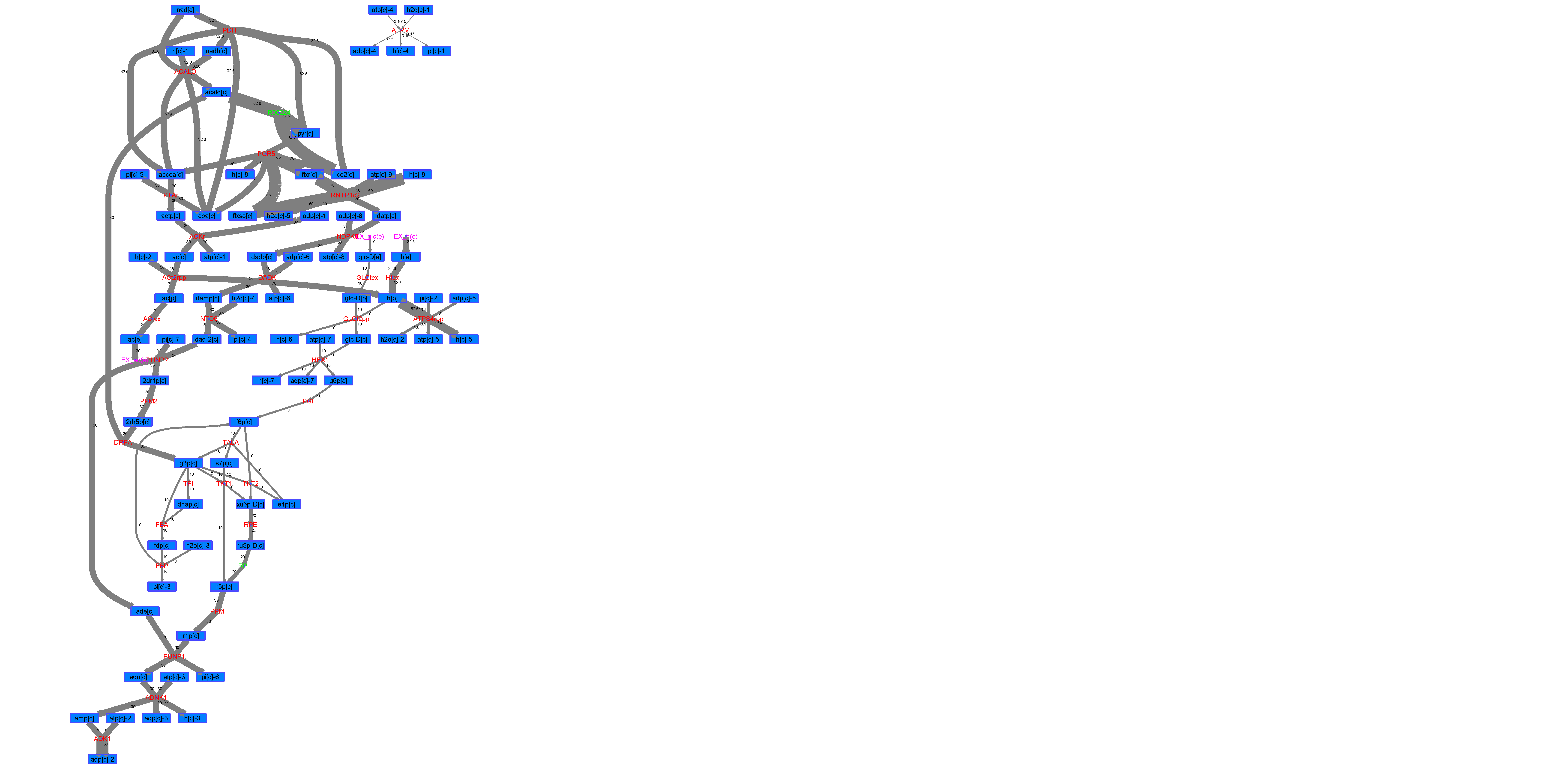

### R00224_For.png

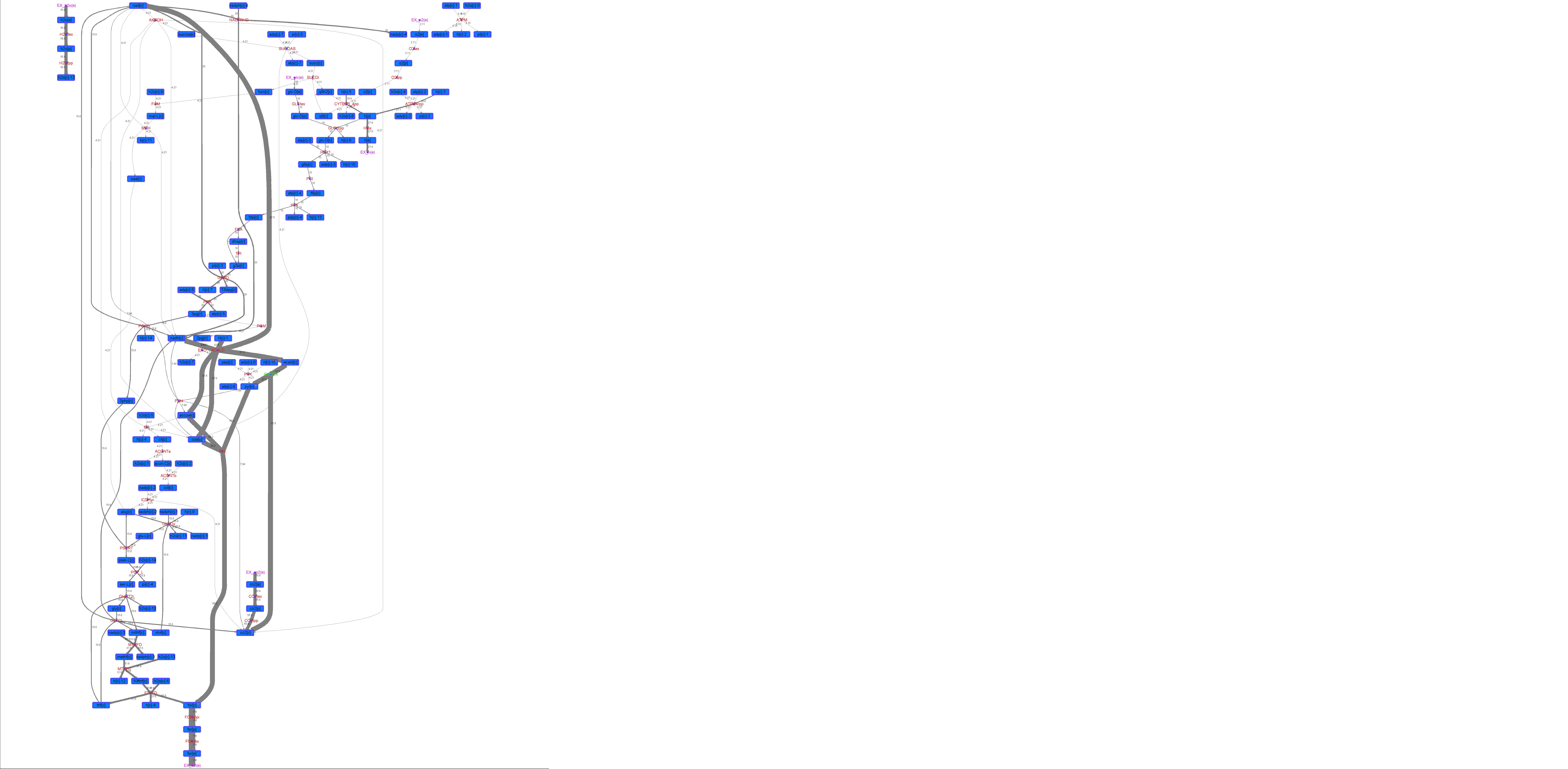

### R00224_Glu.png

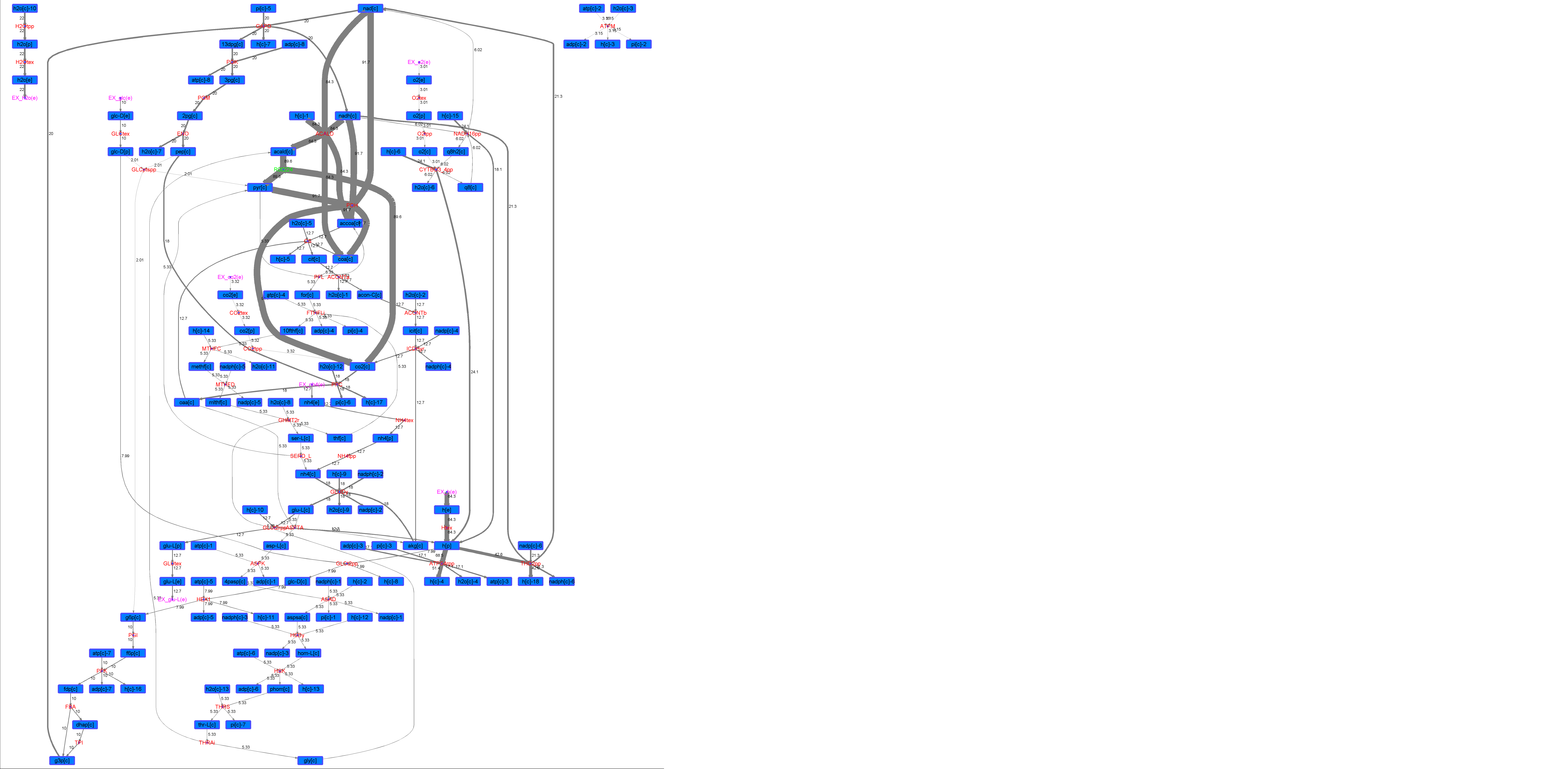

### R00224_L-ala.png

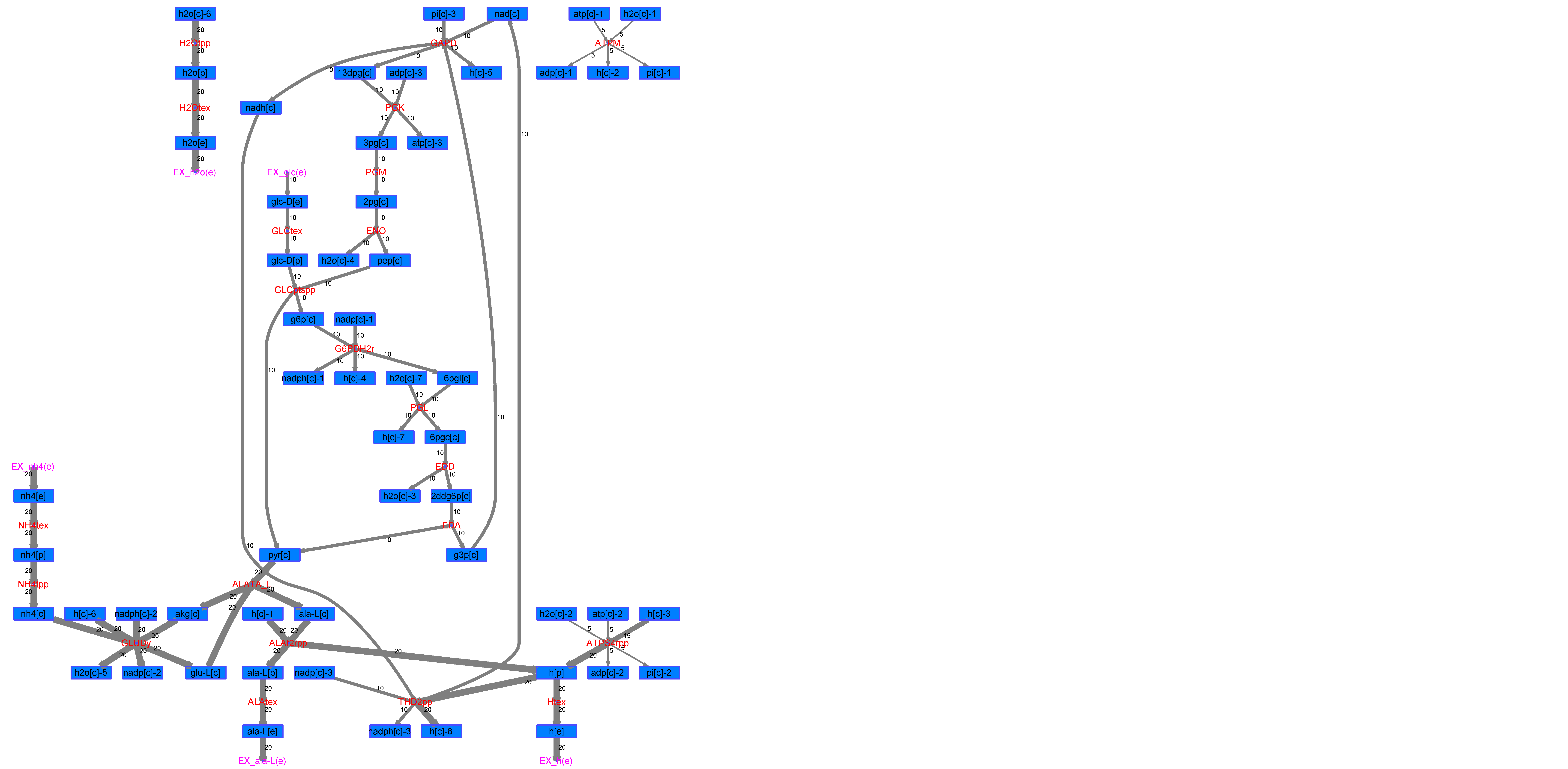

### R00224_L-Mal.png

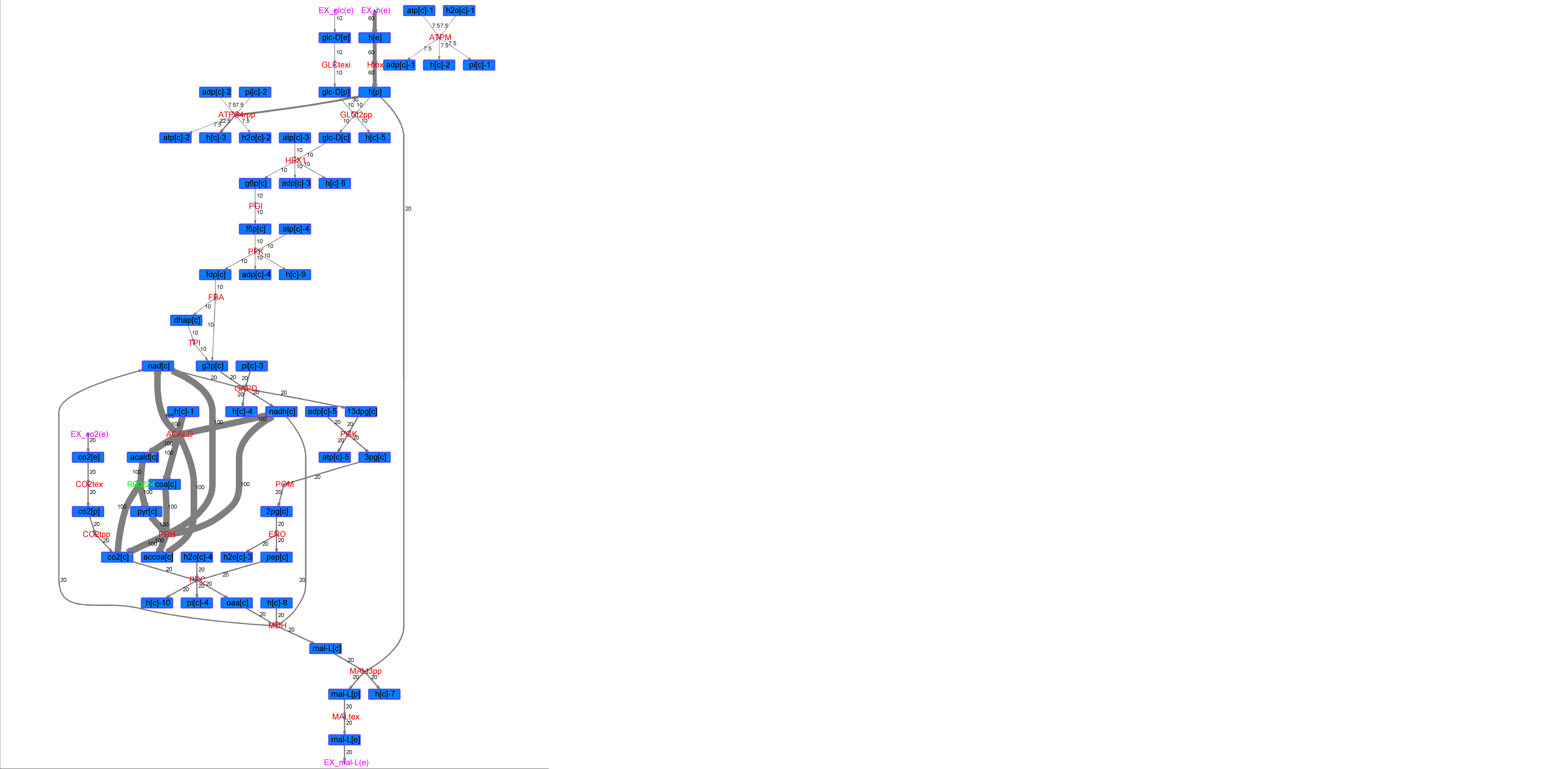

### R00224_L-Phe.png

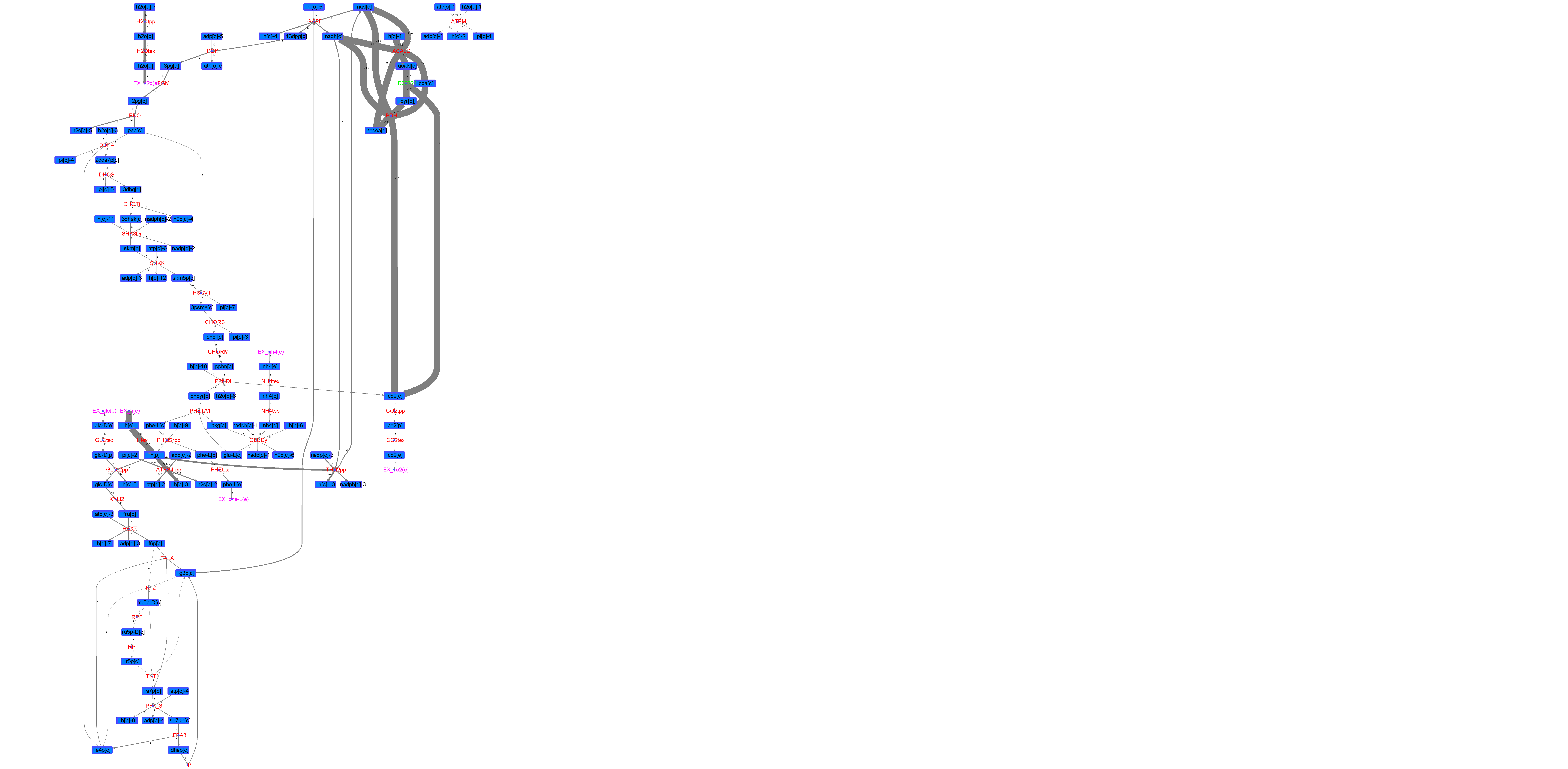

### R00224_L-Try.png

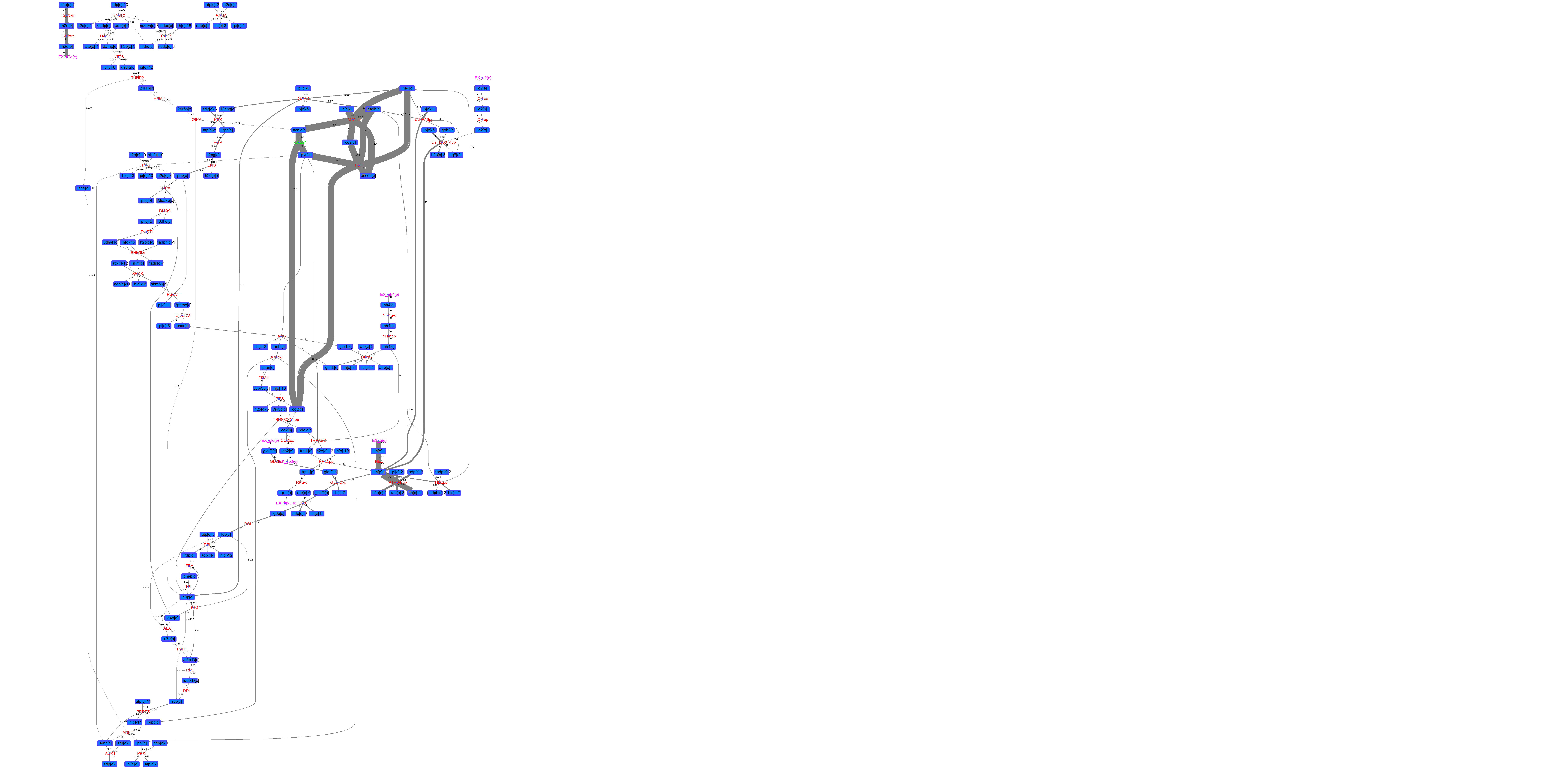

### R00224_Lys.png

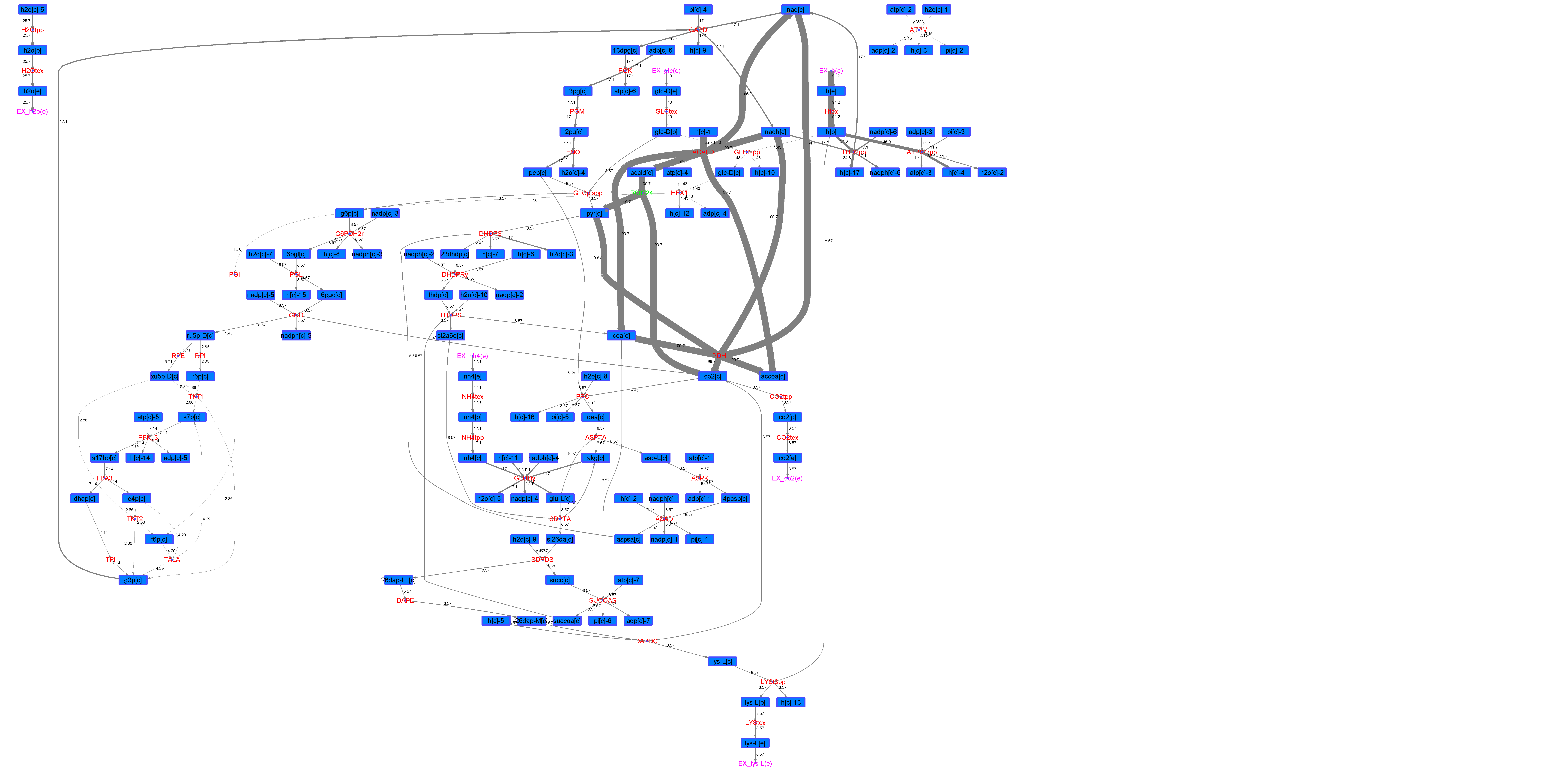

### R00224_Suc.png

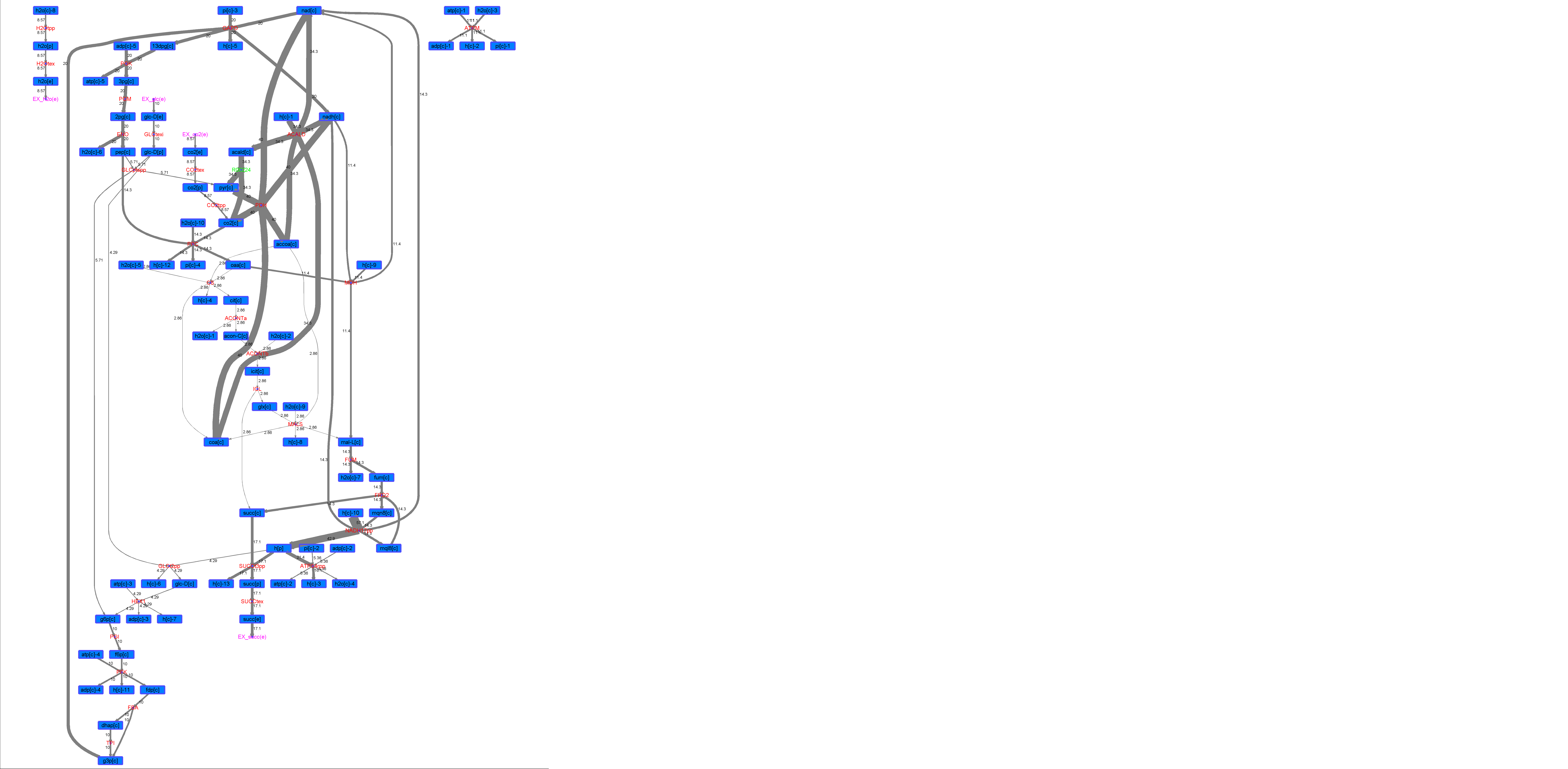

### R00224_Thr.png

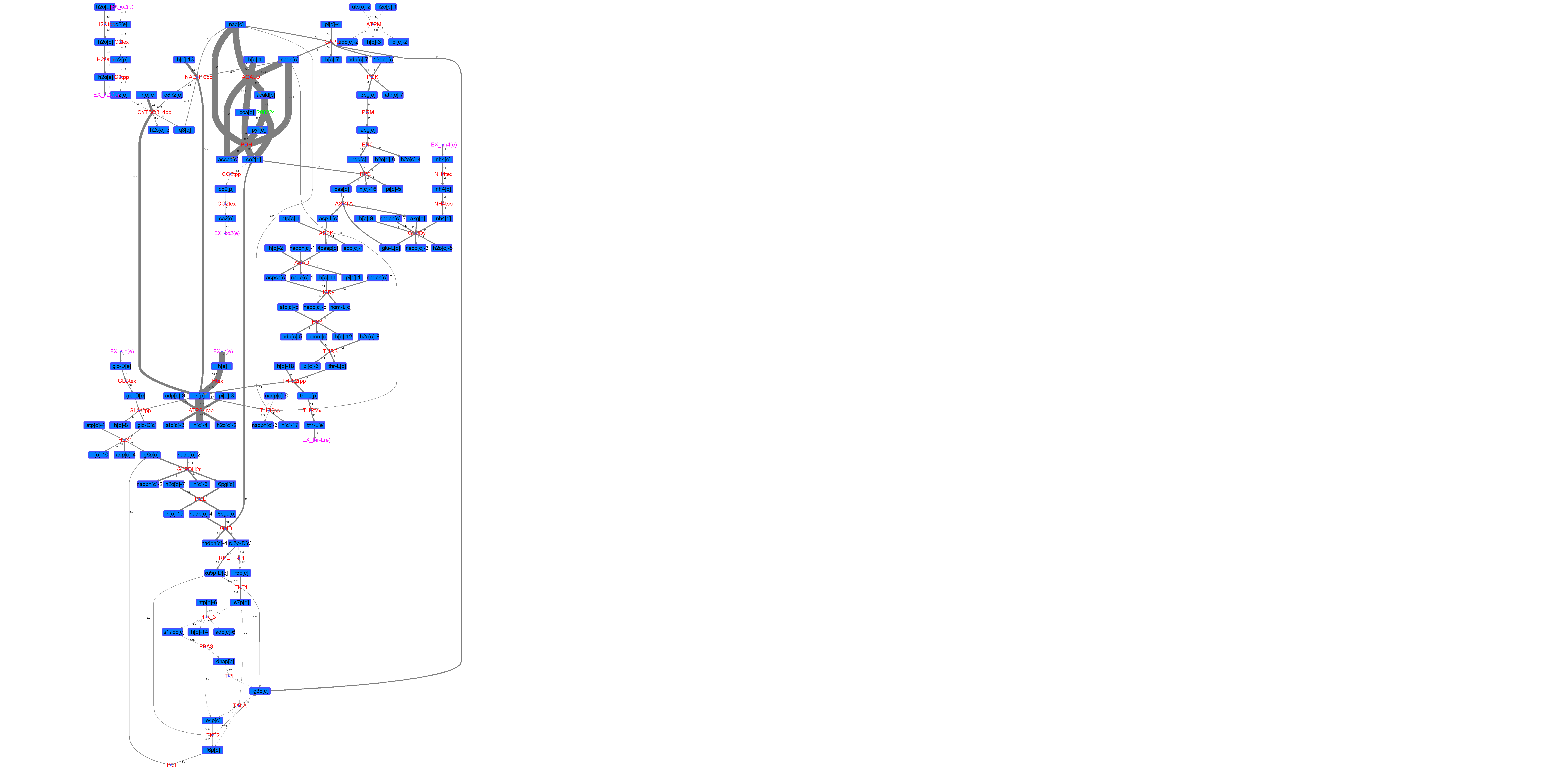

### R00243_Ace.png

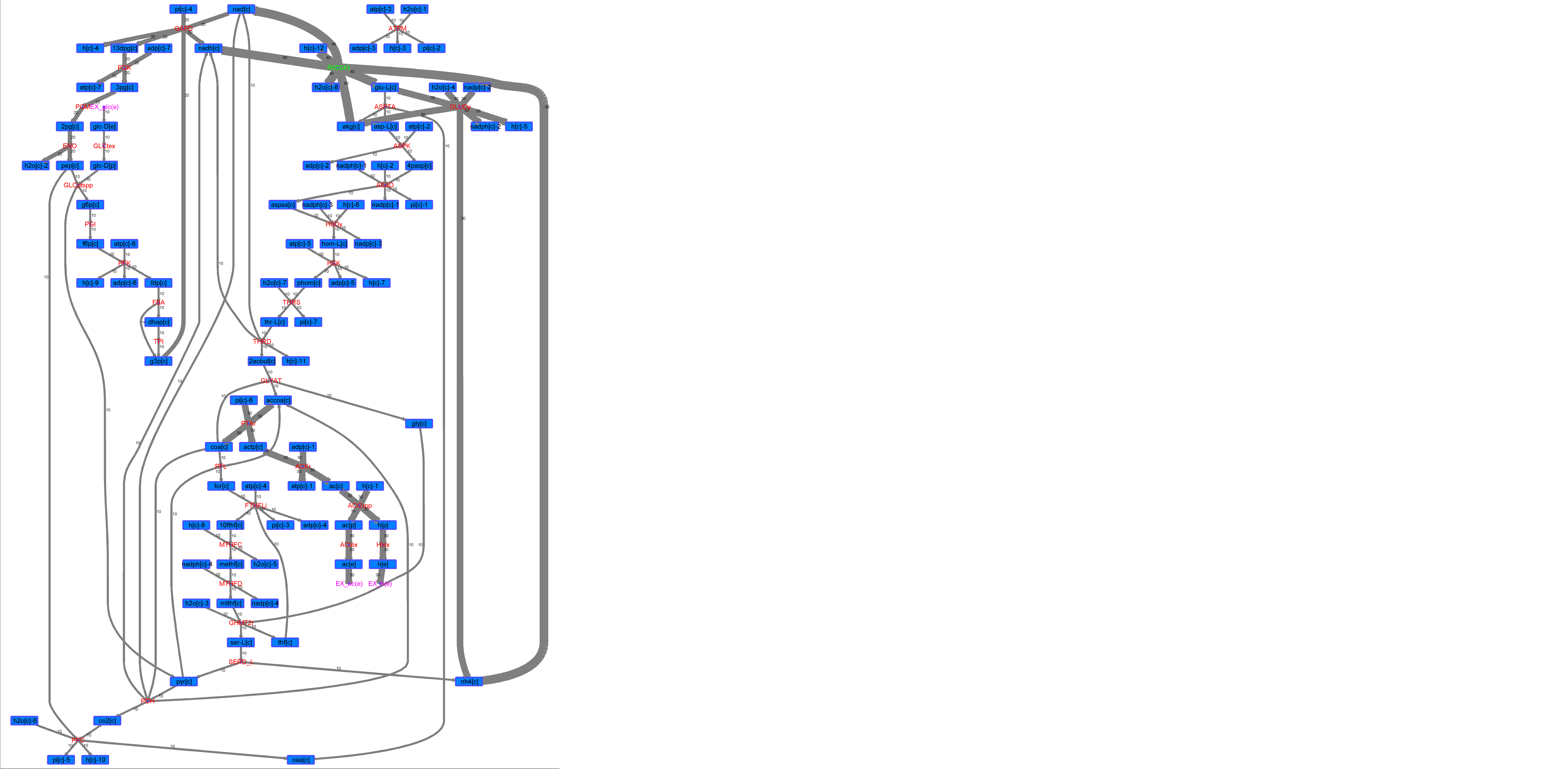

### R00243_For.png

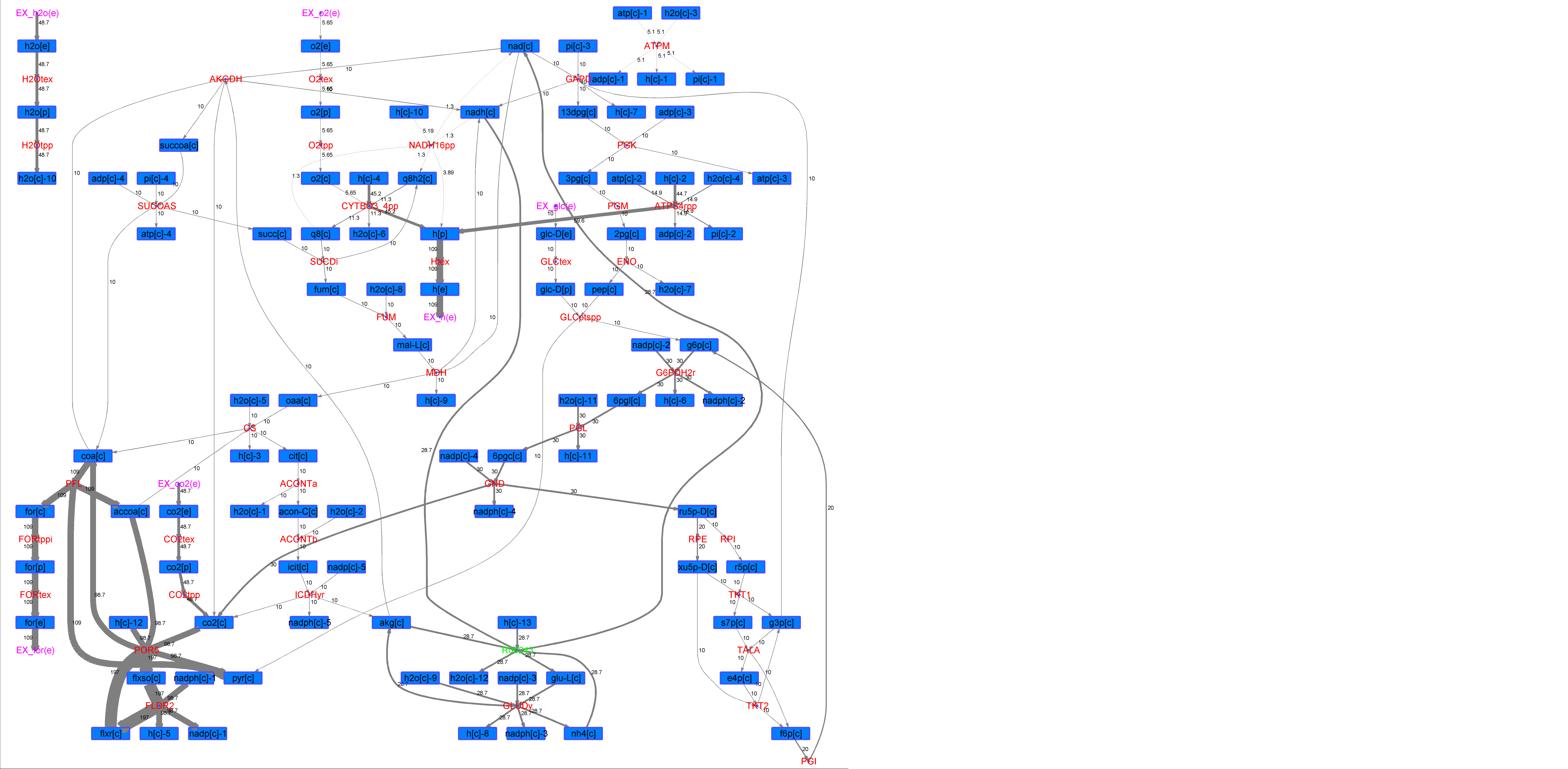

### R00243_Glu.png

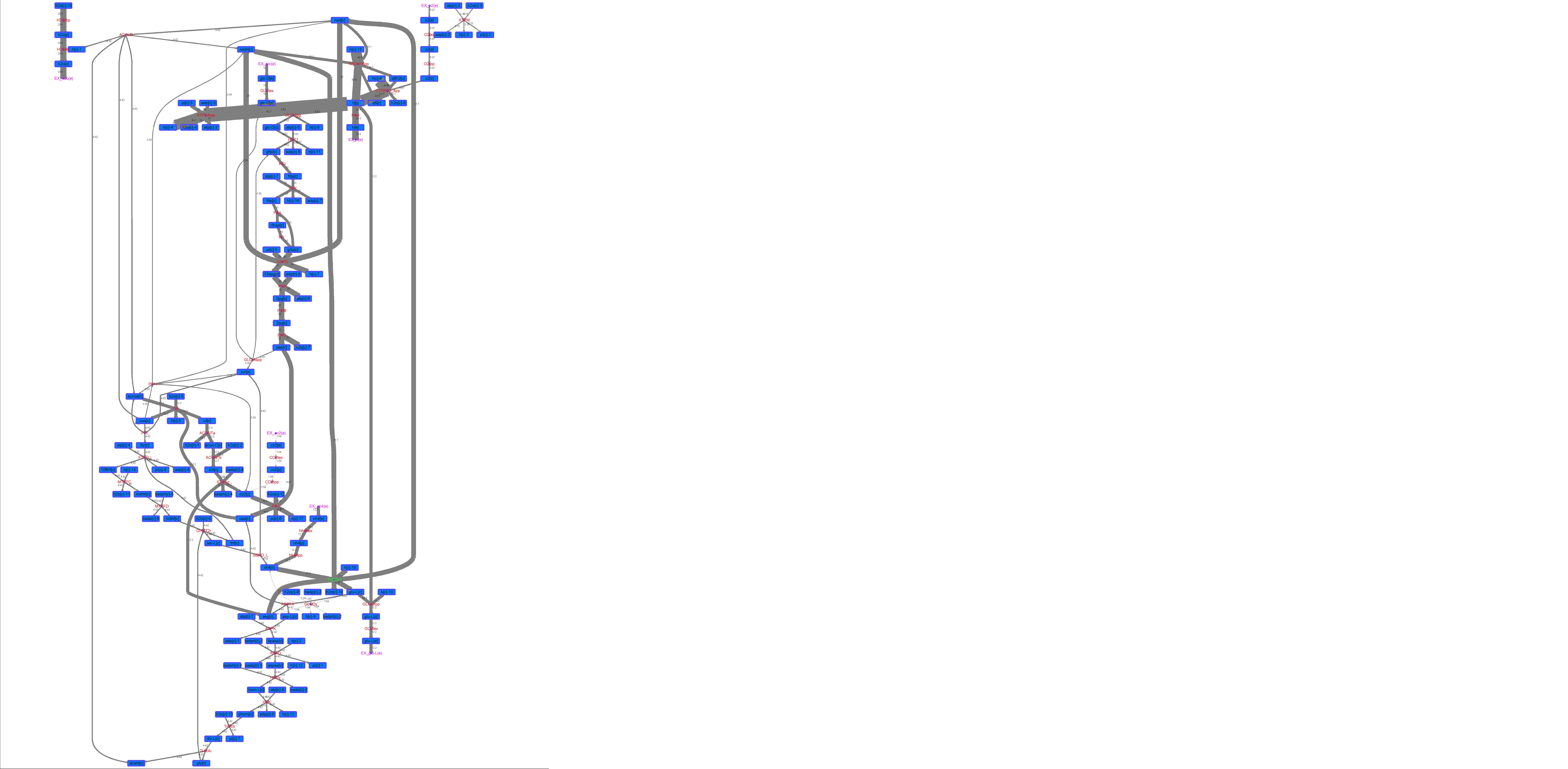

### R00243_L-ala.png

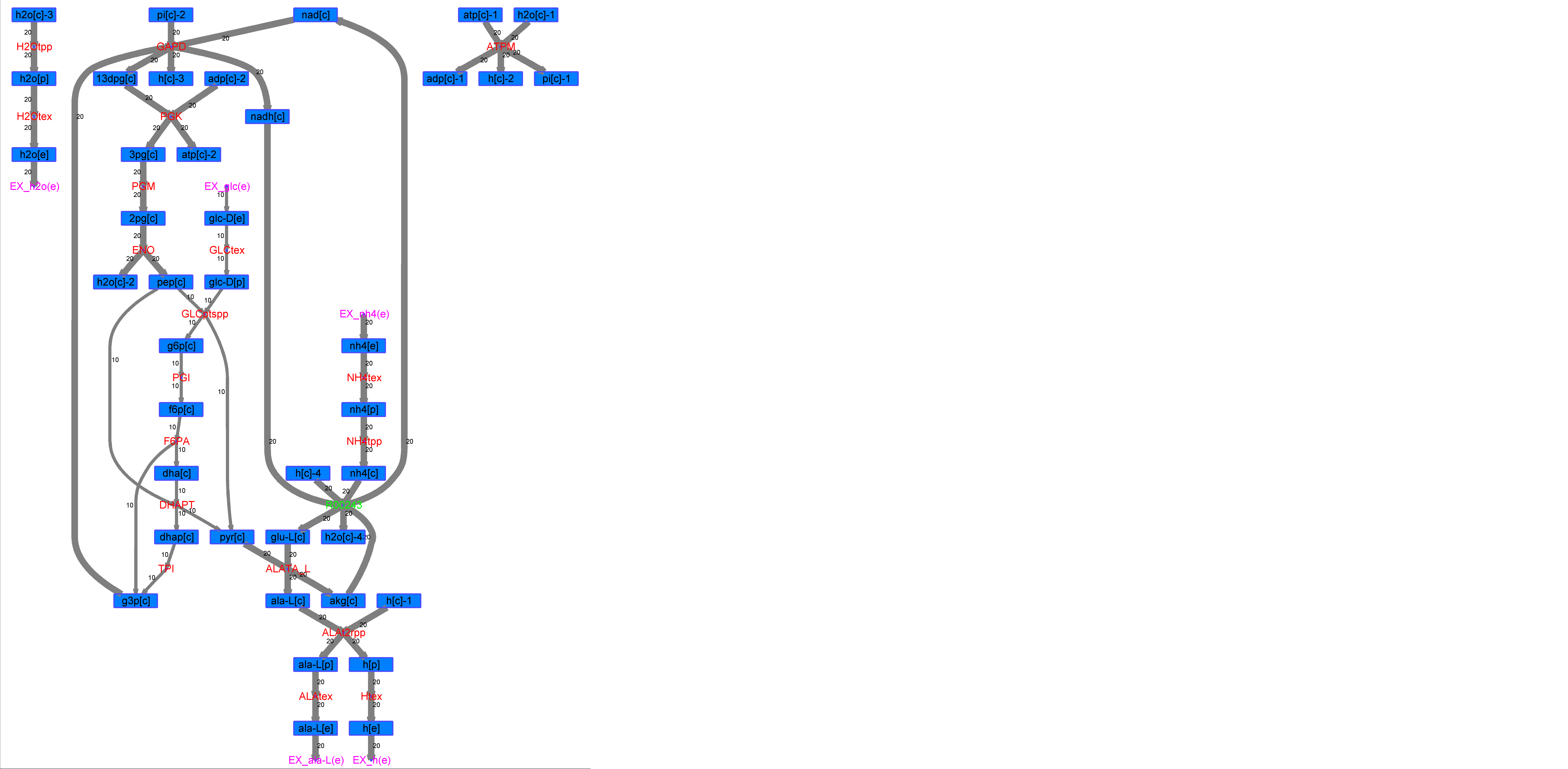

### R00243_L-Mal.png

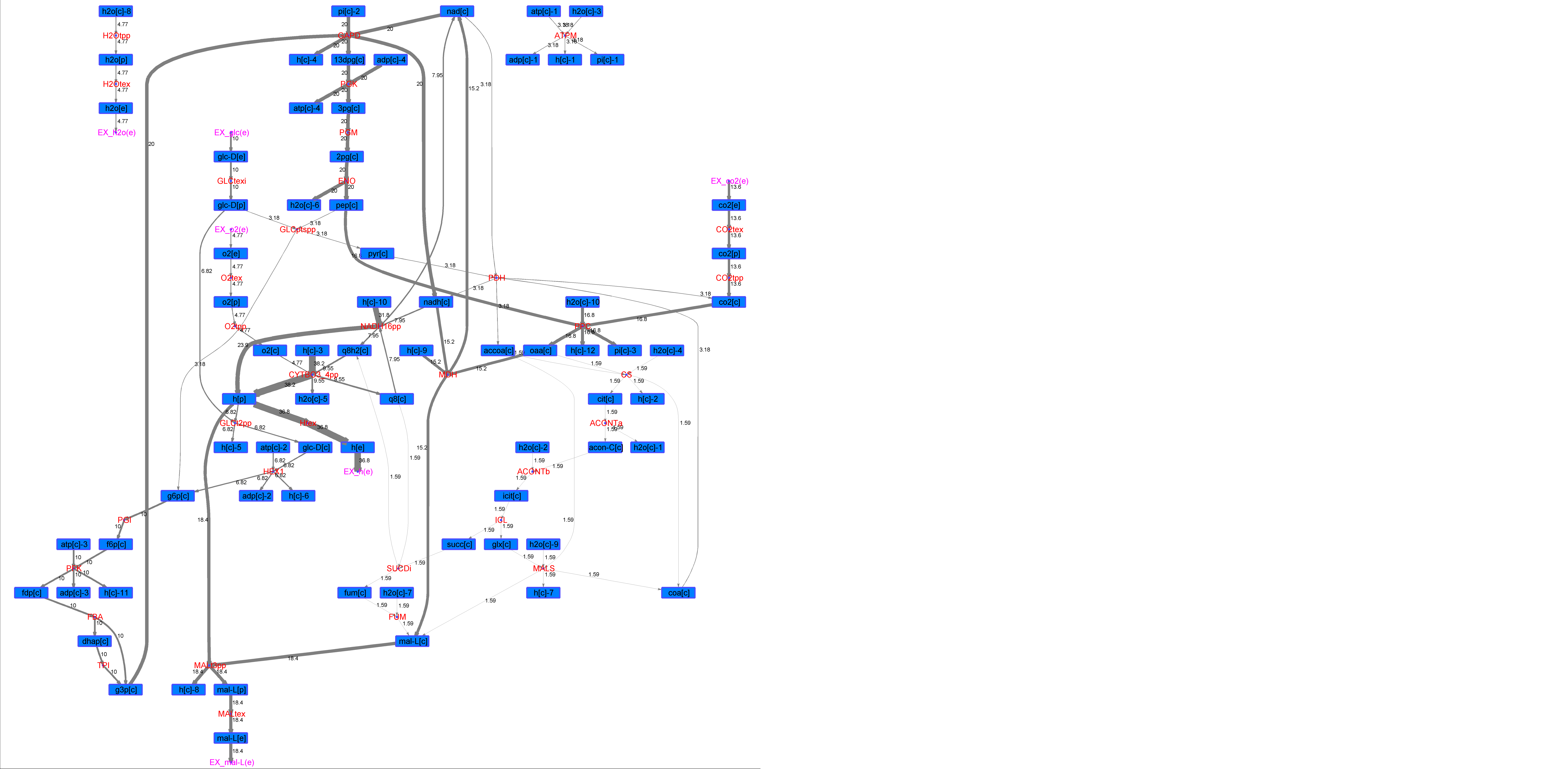

### R00243_L-Phe.png

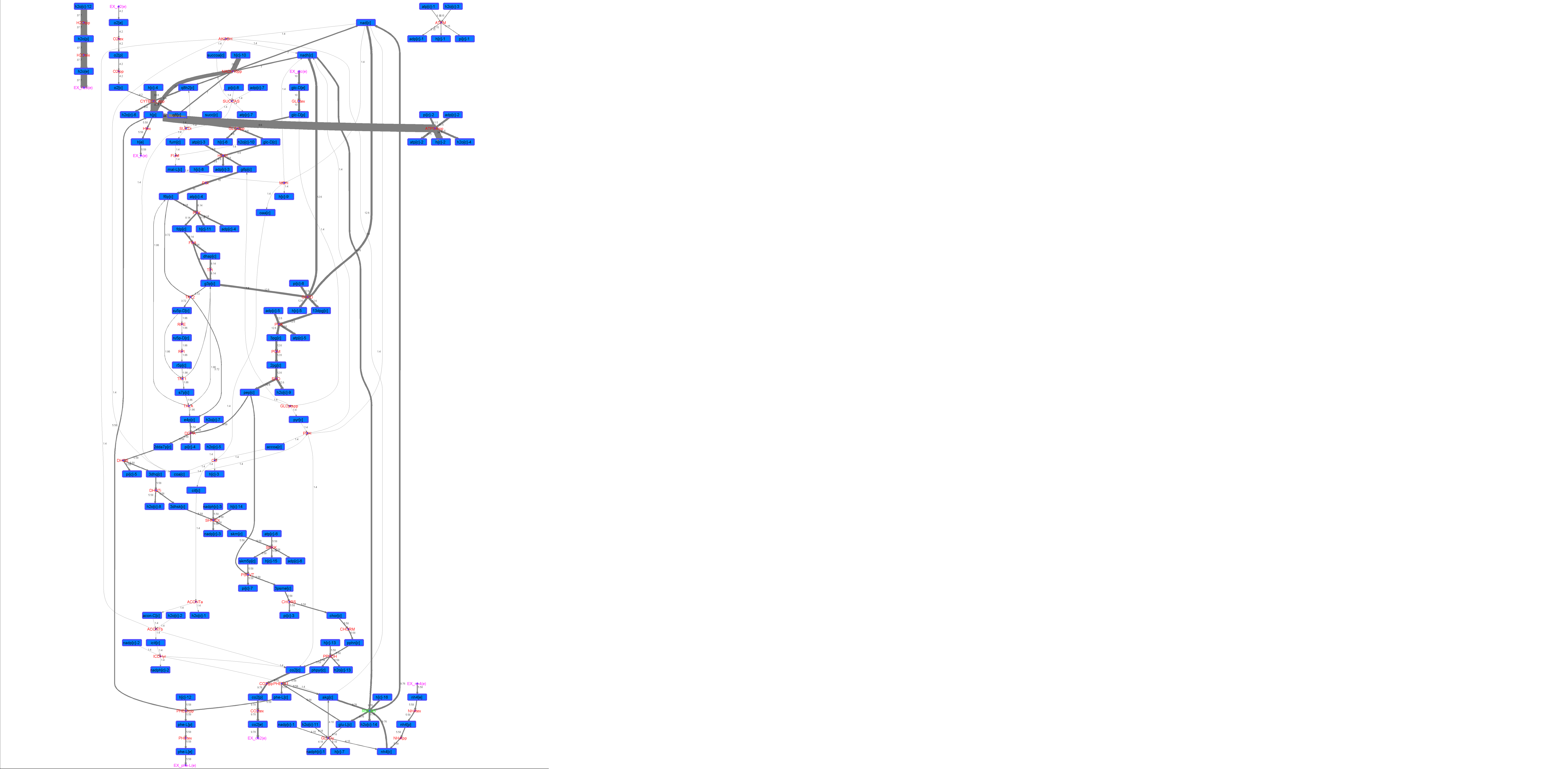

### R00243_L-Try.png

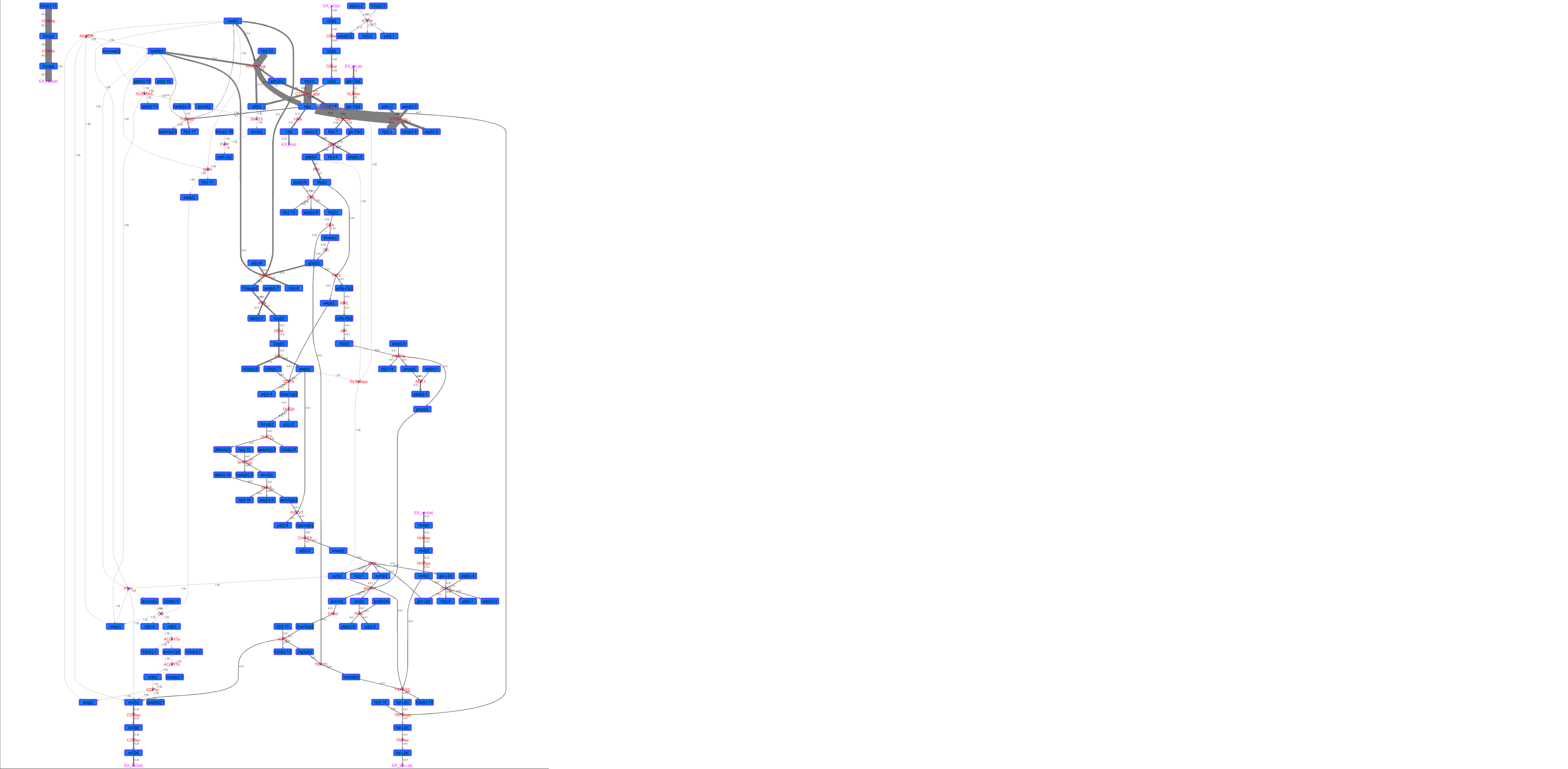

### R00243_Lys.png

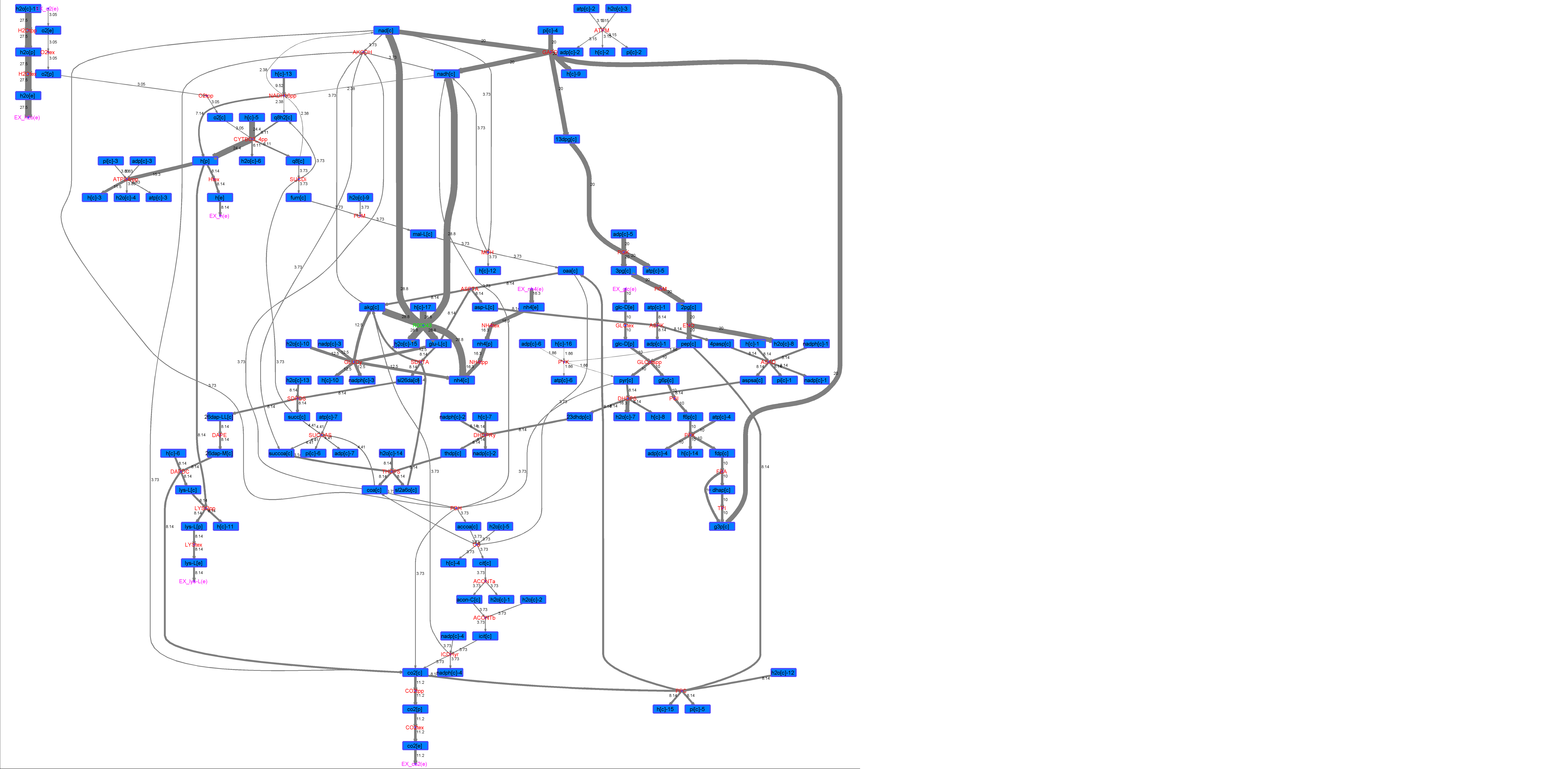

### R00243_Suc.png

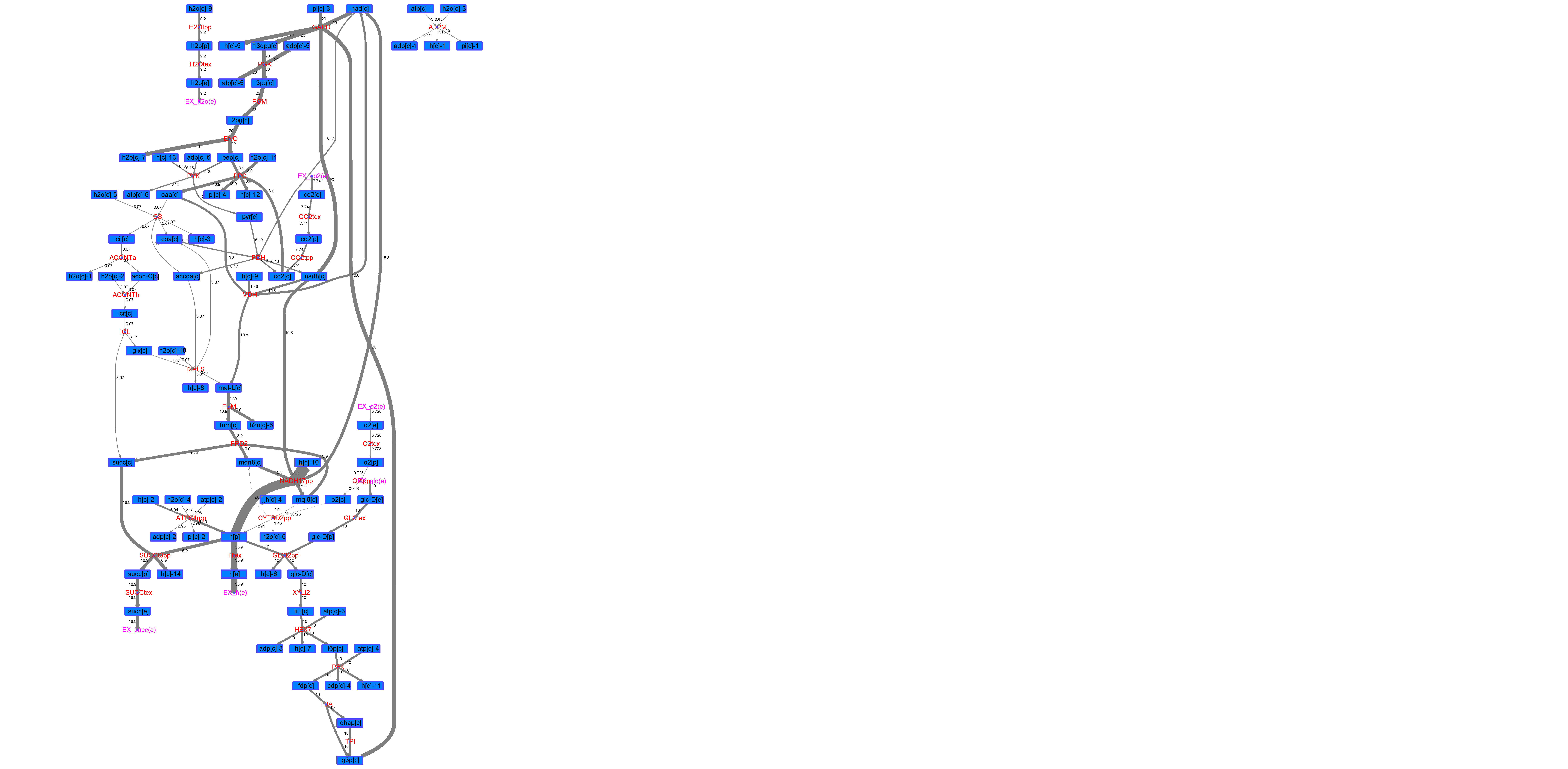

### R00243_Thr.png

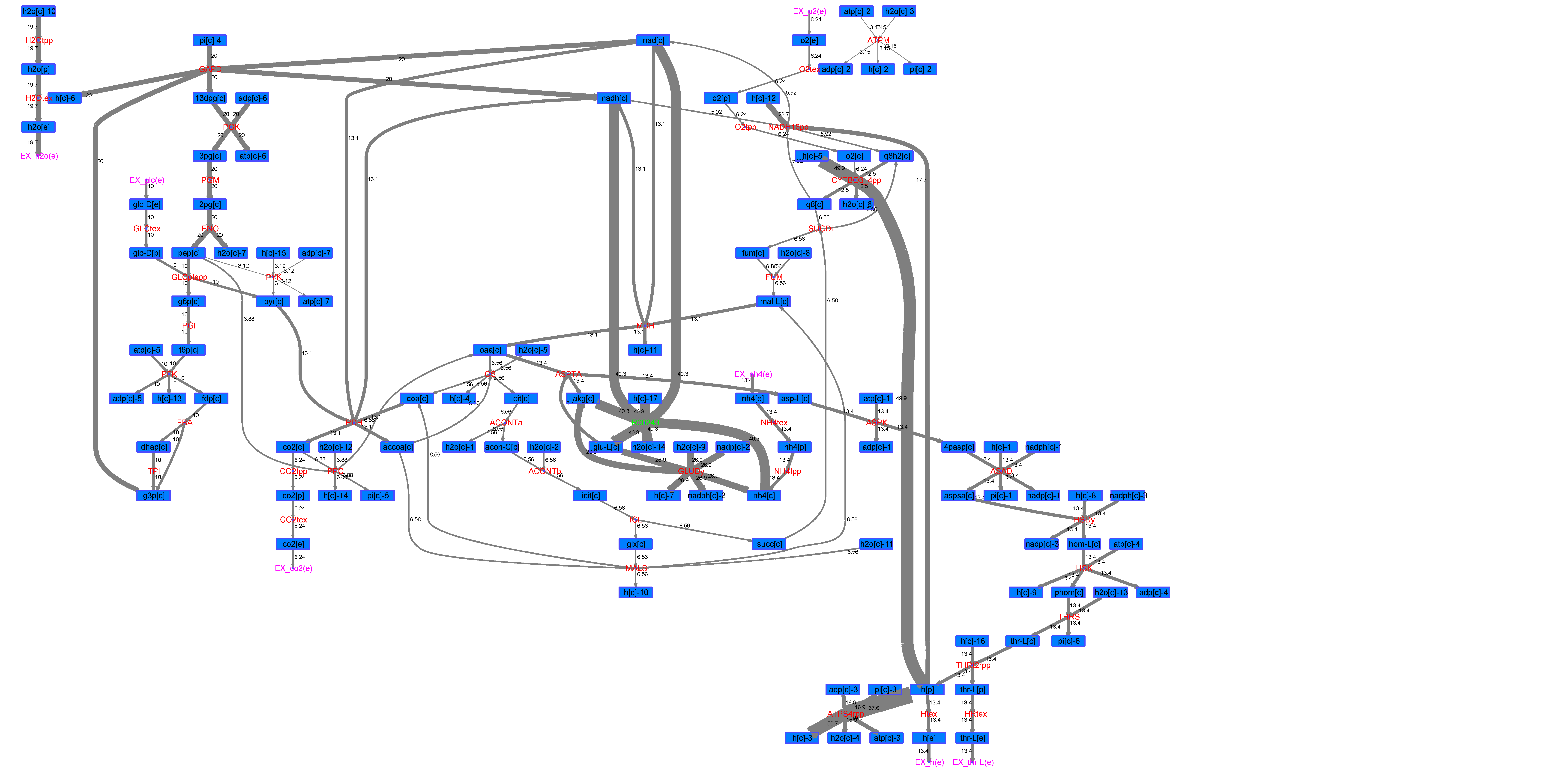

### R00265_Ace.png

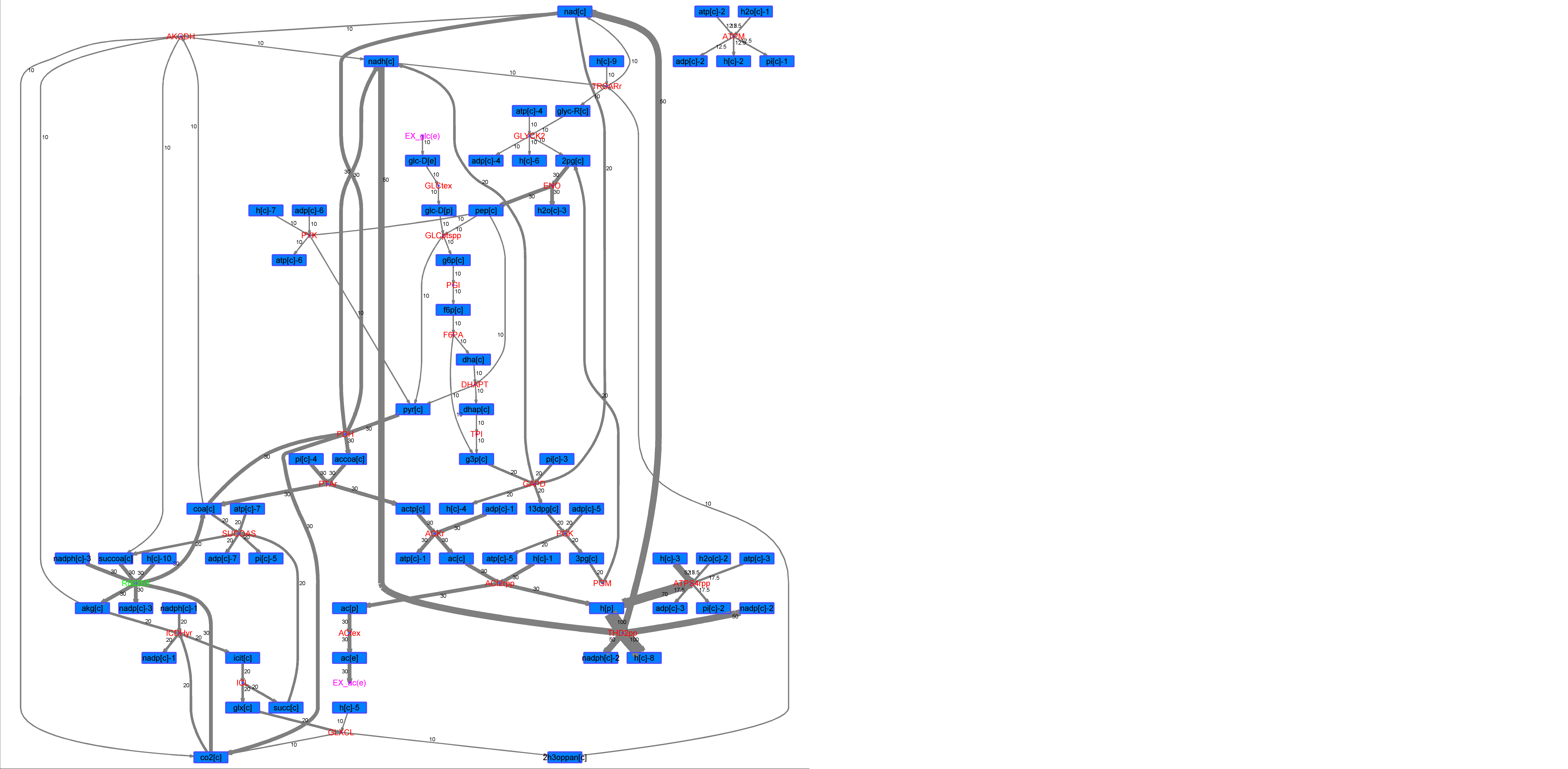

### R00265_For.png

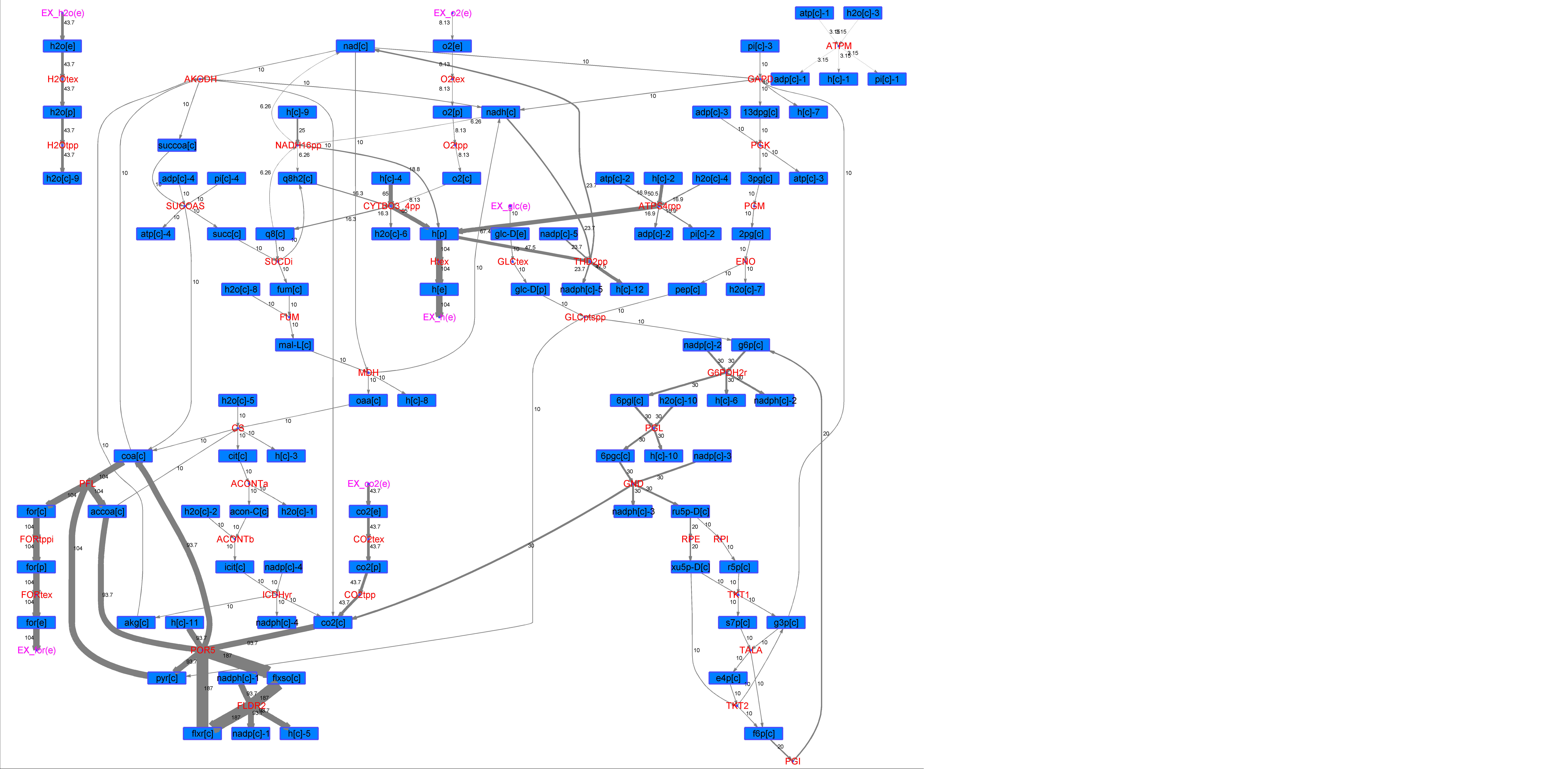

### R00265_Glu.png

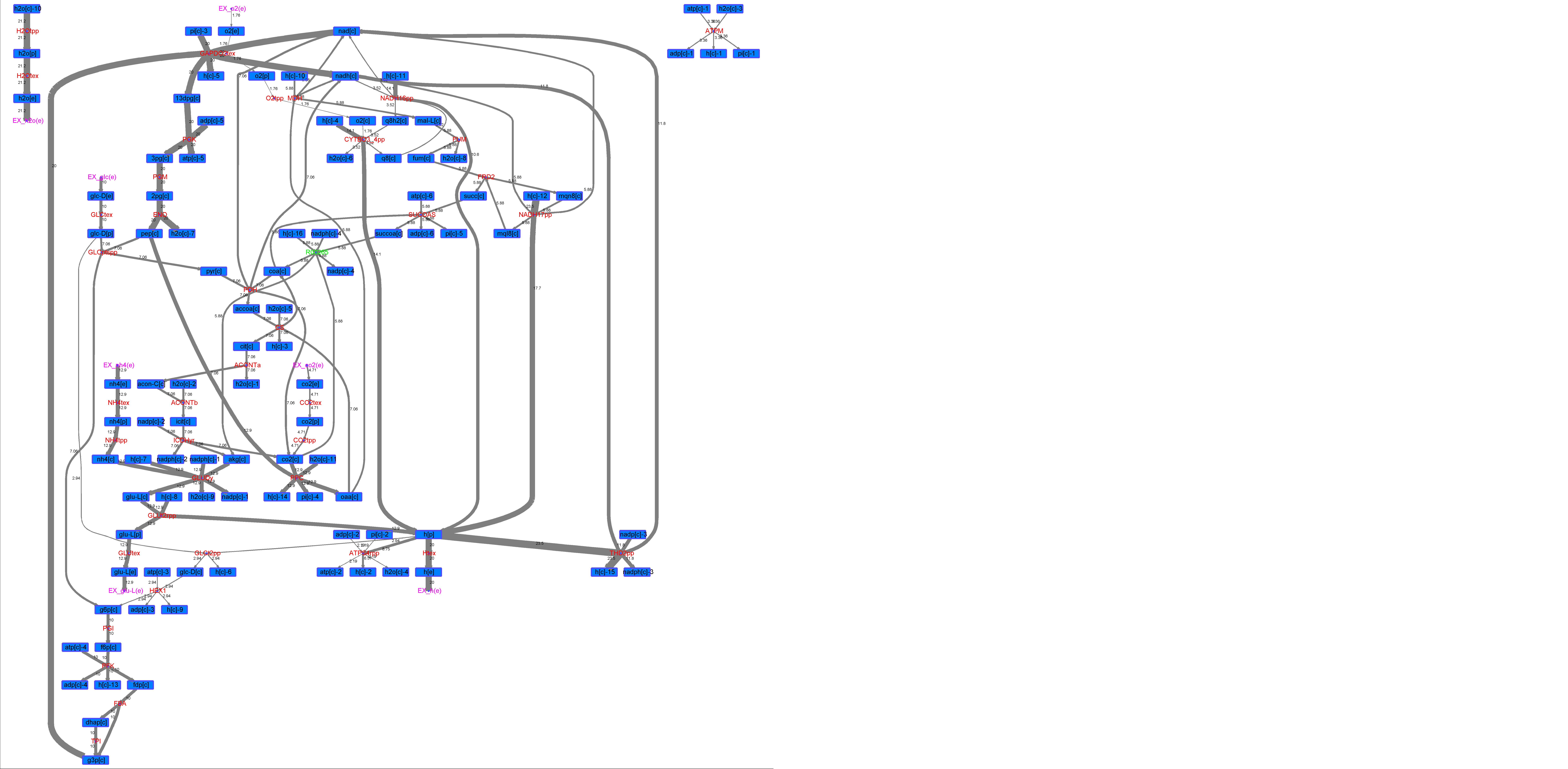

### R00265_L-ala.png

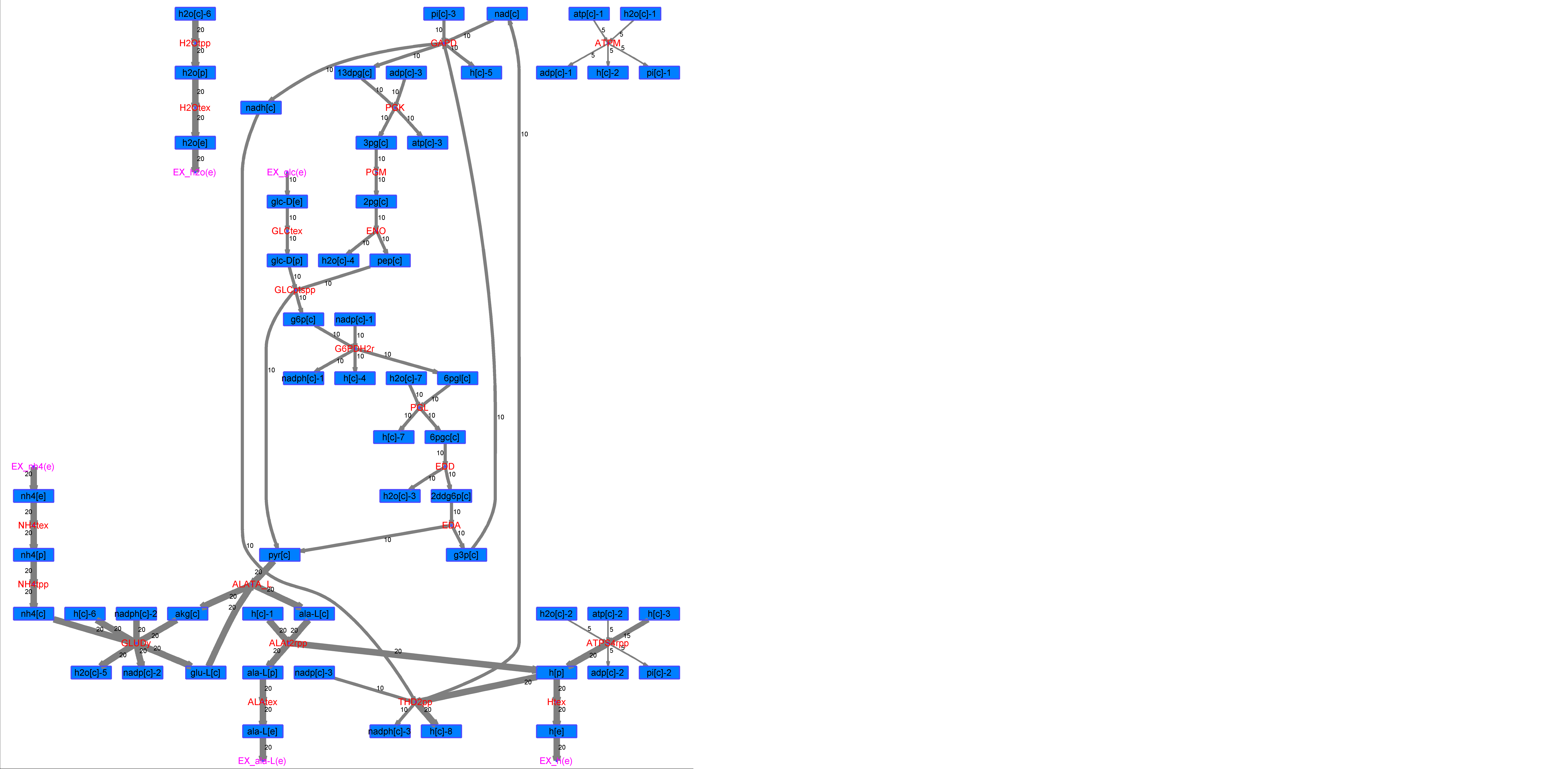

### R00265_L-Mal.png

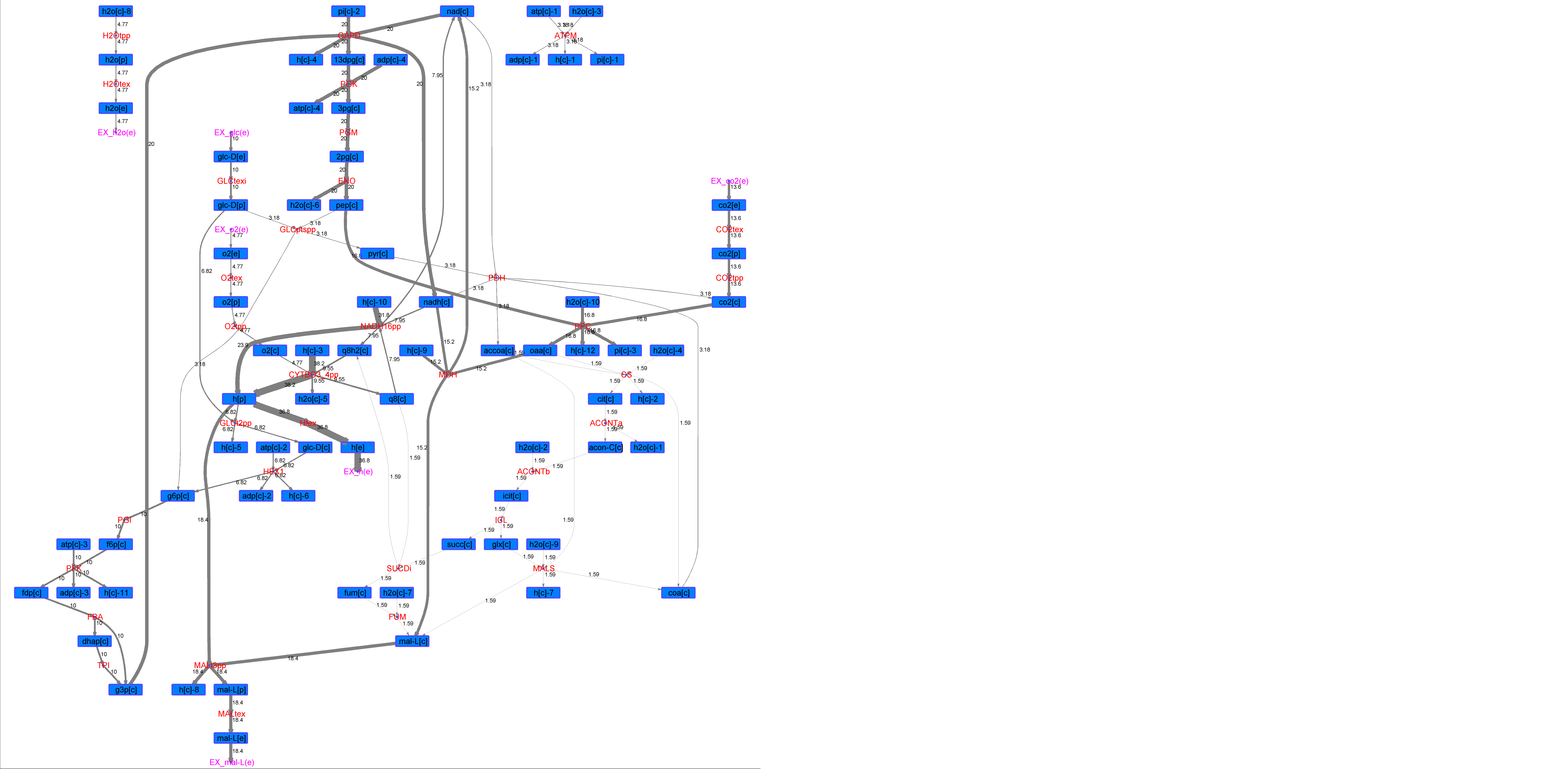

### R00265_L-Phe.png

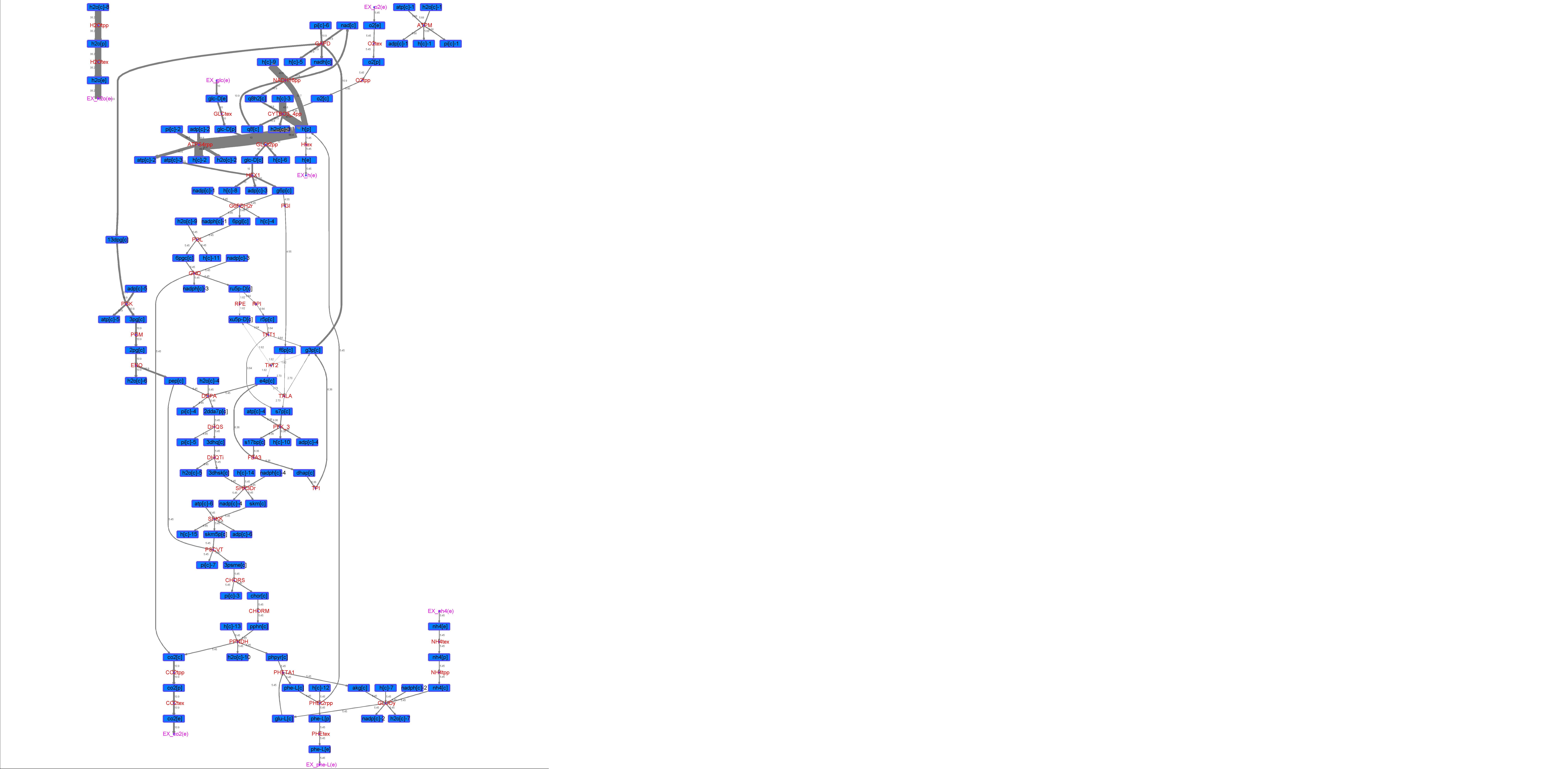

### R00265_L-Try.png

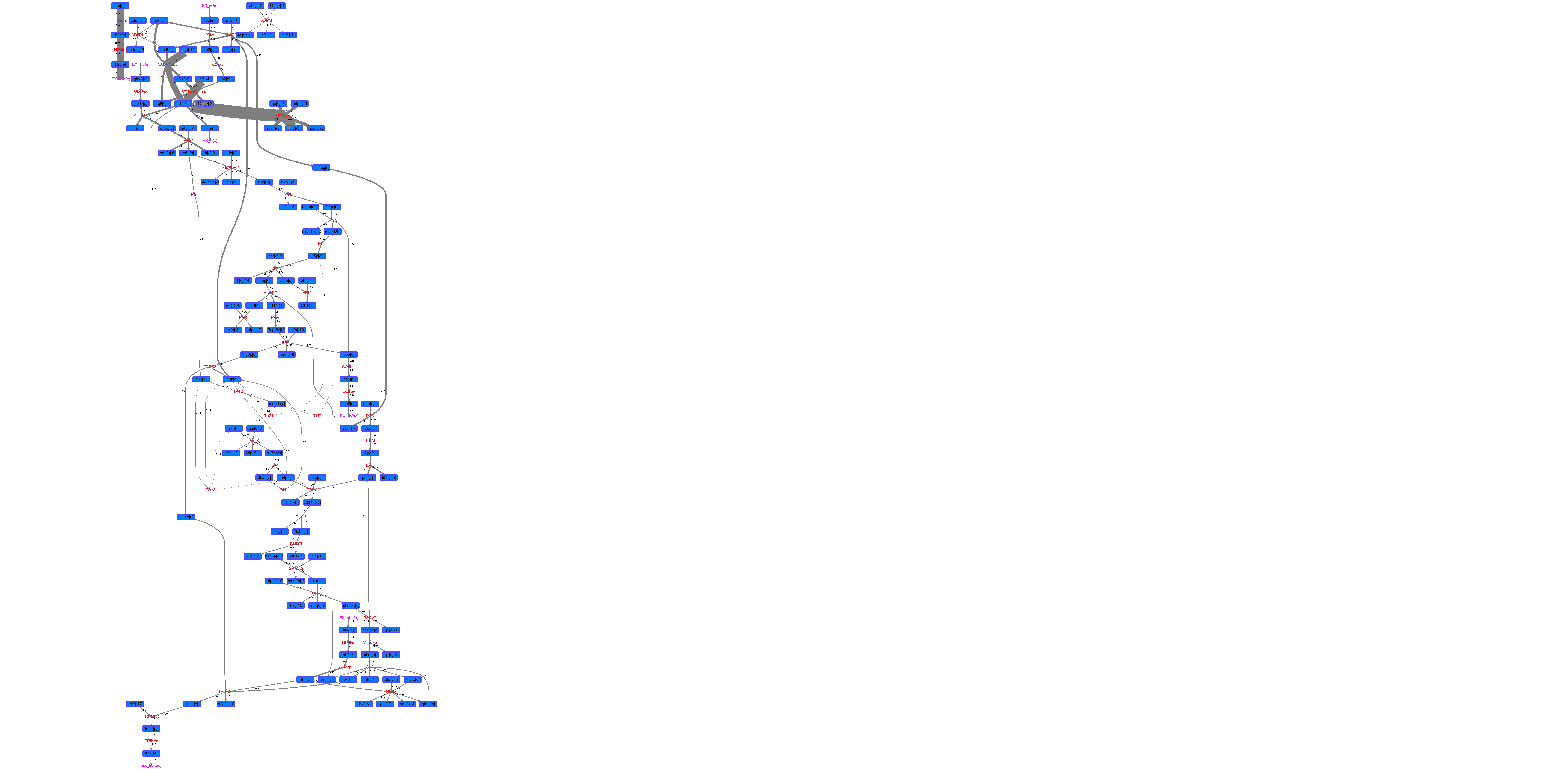

### R00265_Lys.png

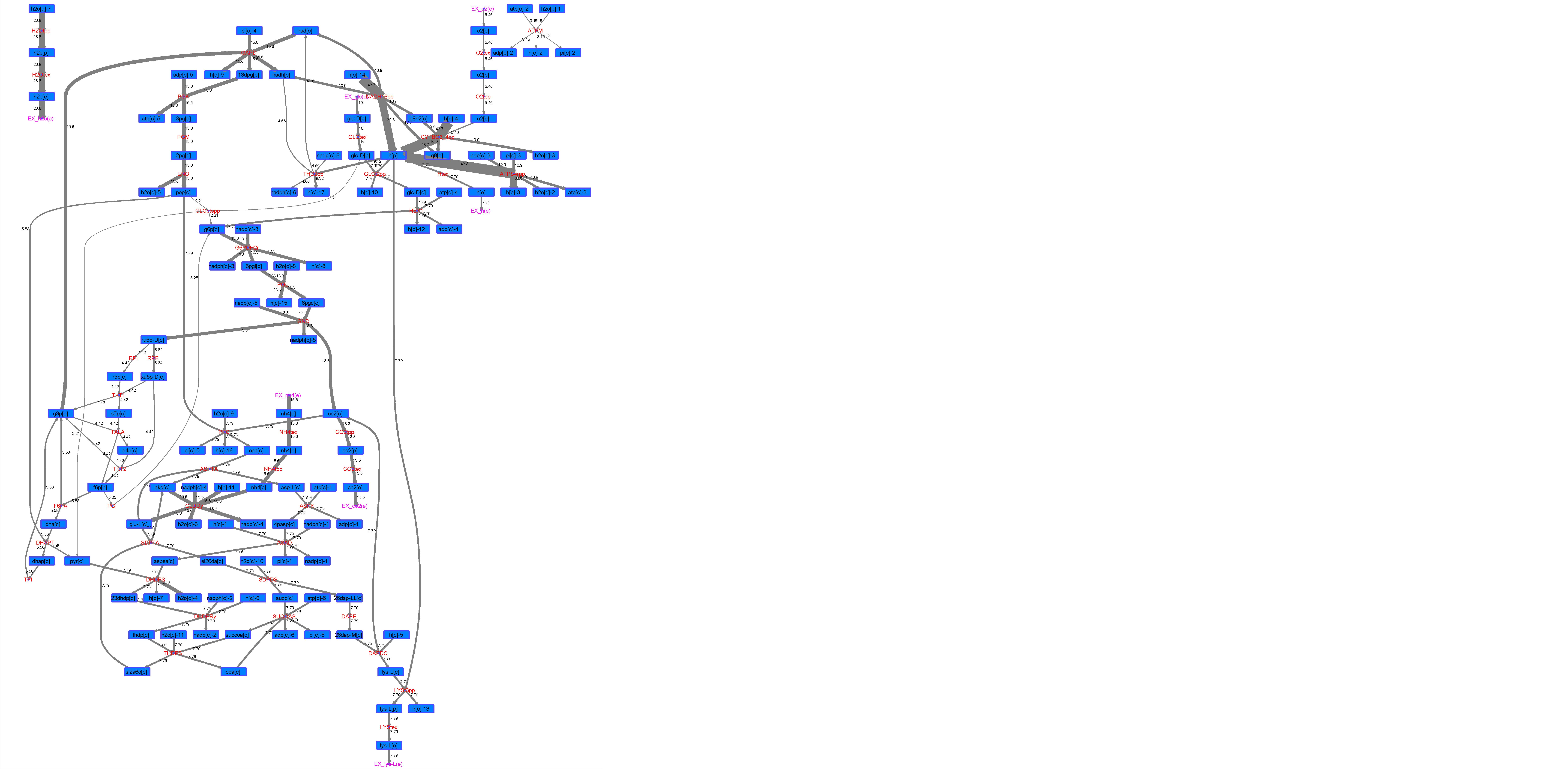

### R00265_Suc.png

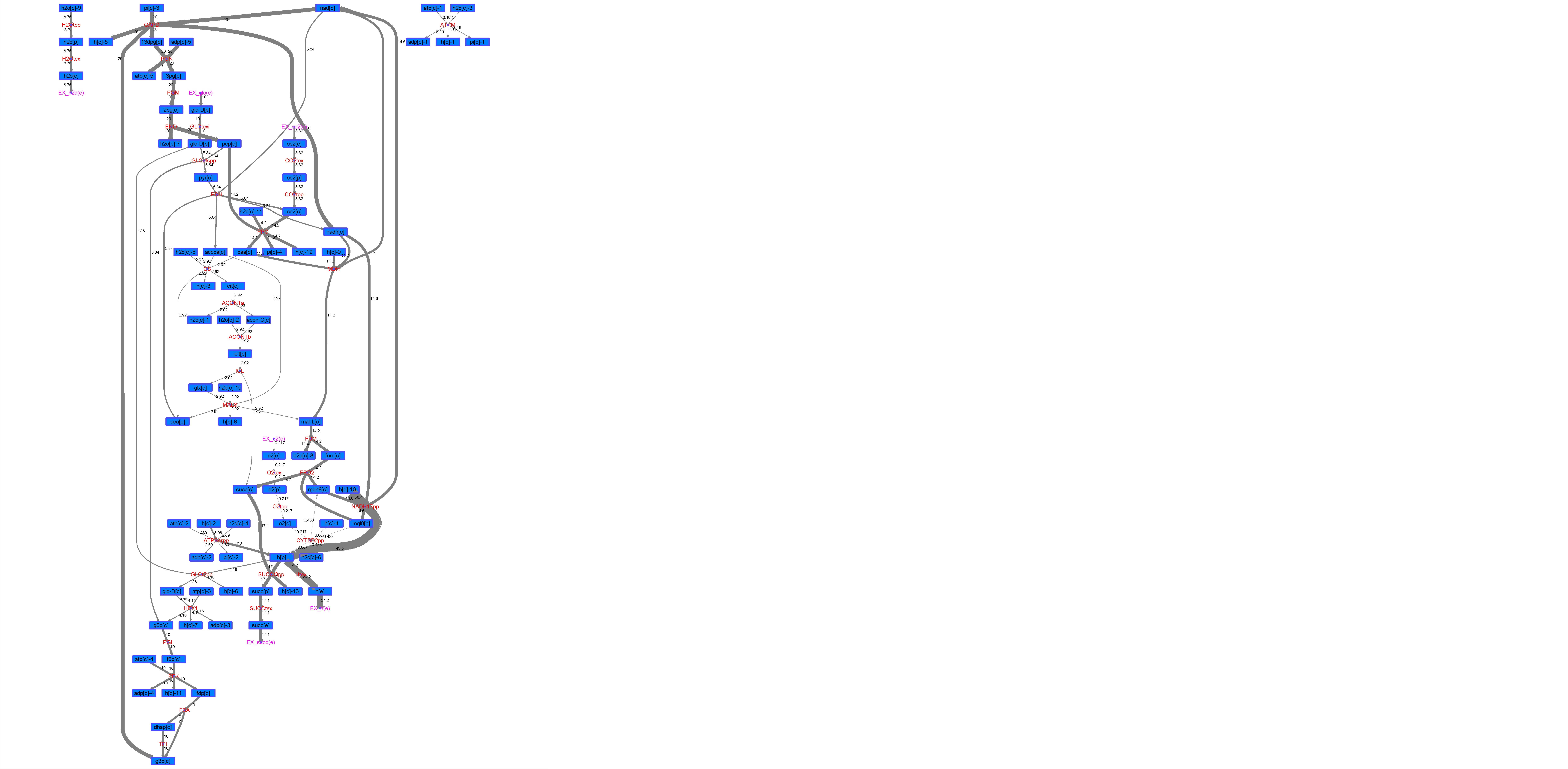

### R00265_Thr.png

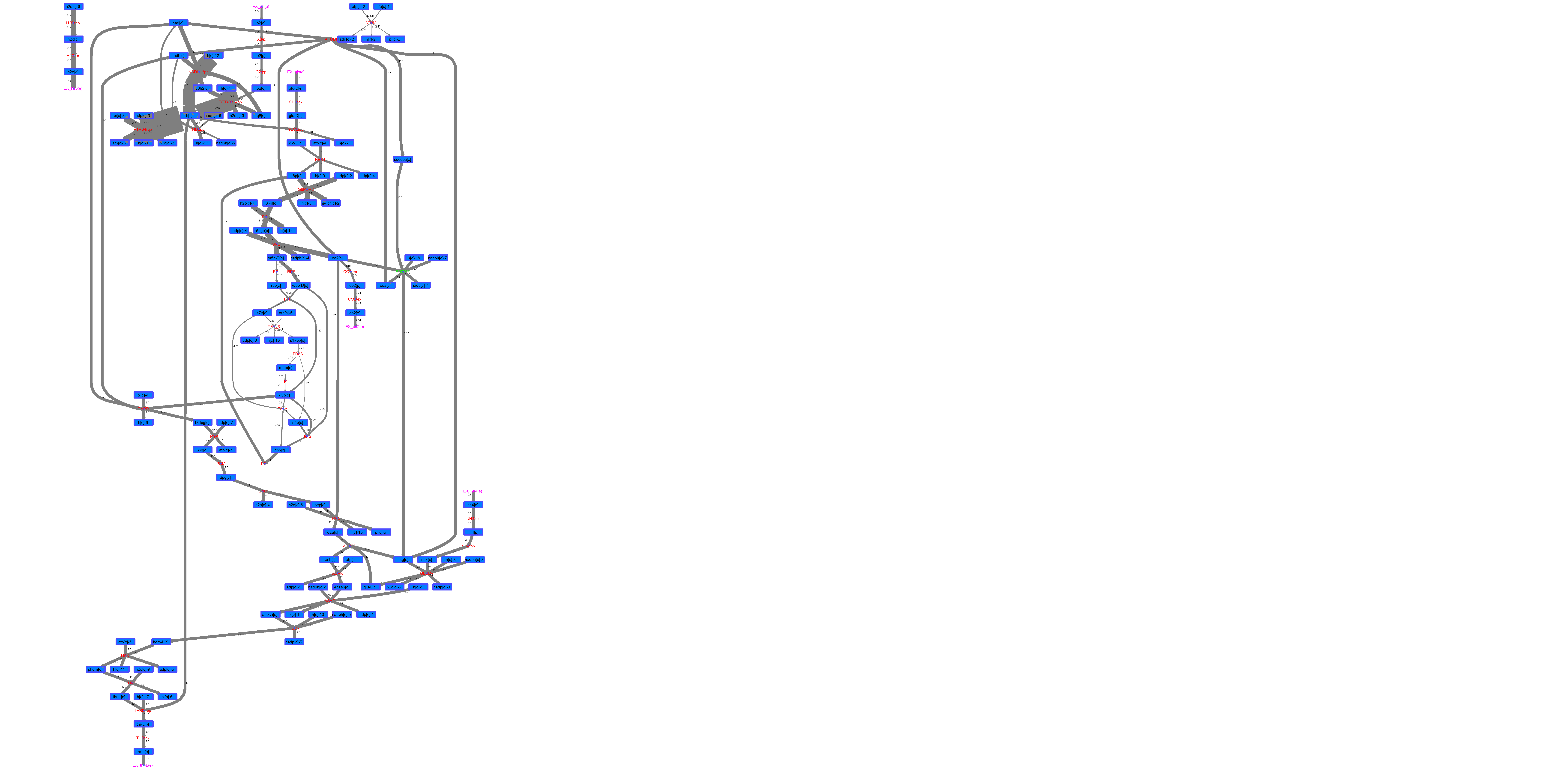

### R00343_Ace.png

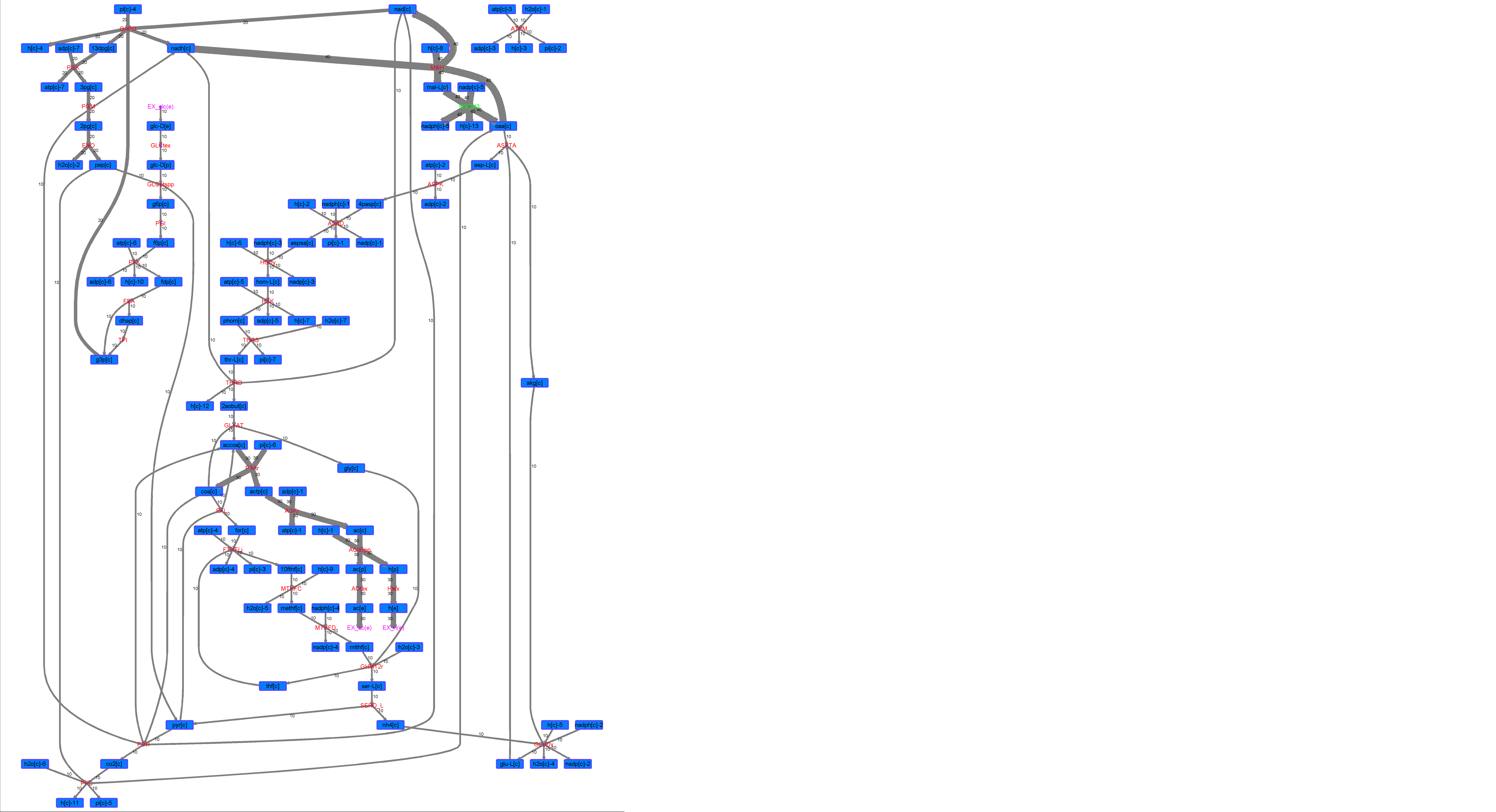

### R00343_For.png

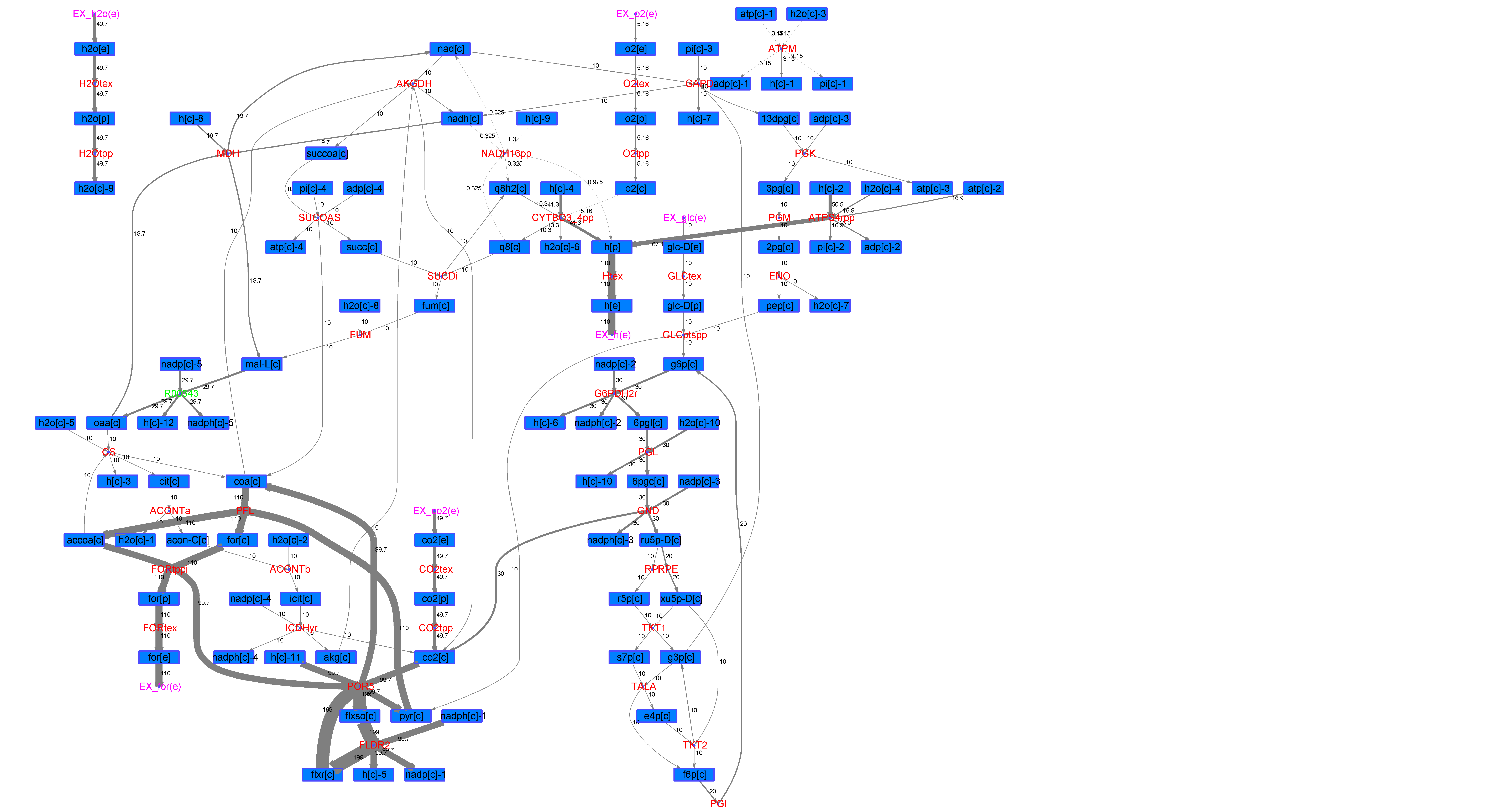

### R00343_Glu.png

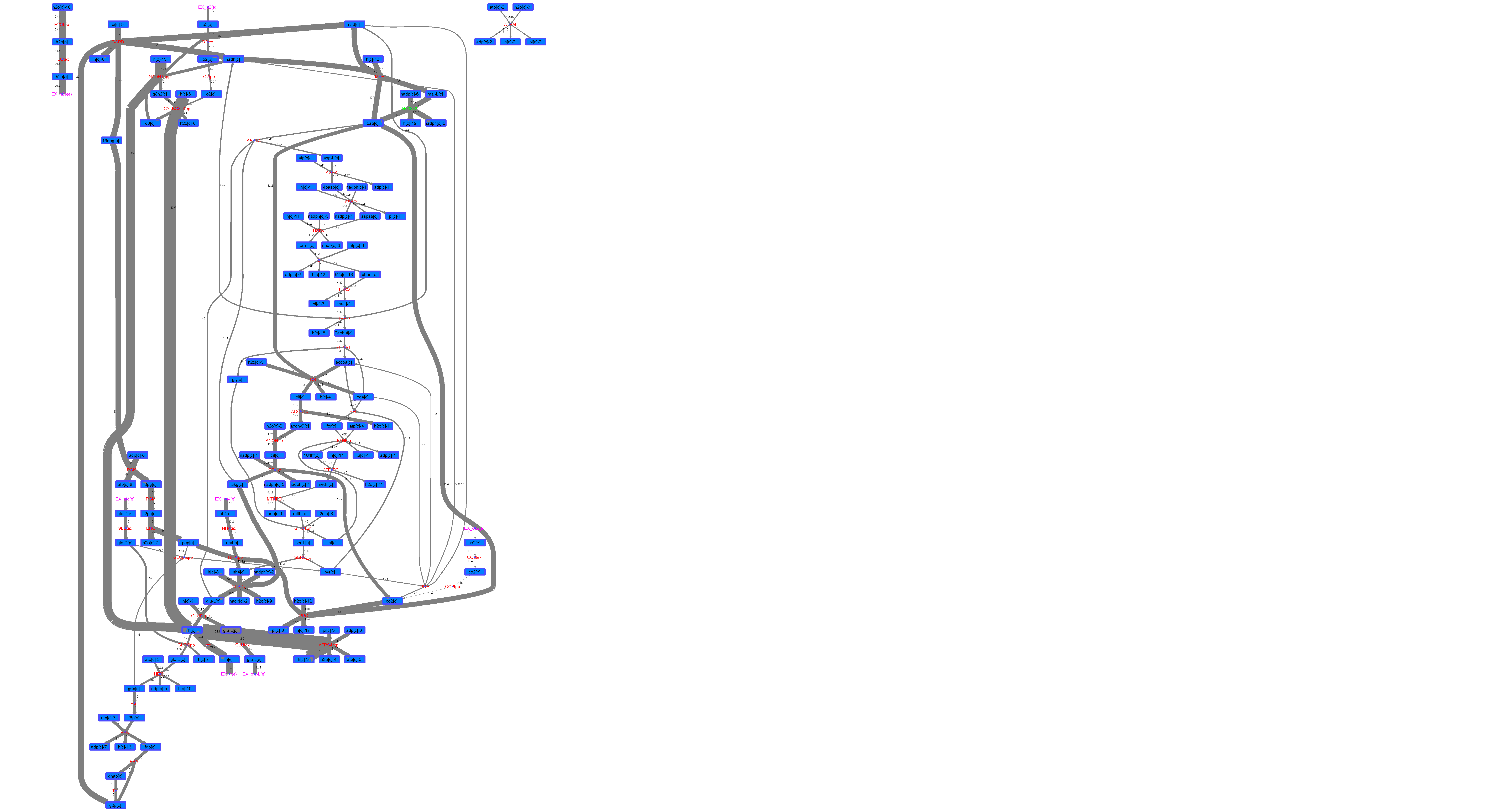

### R00343_L-ala.png

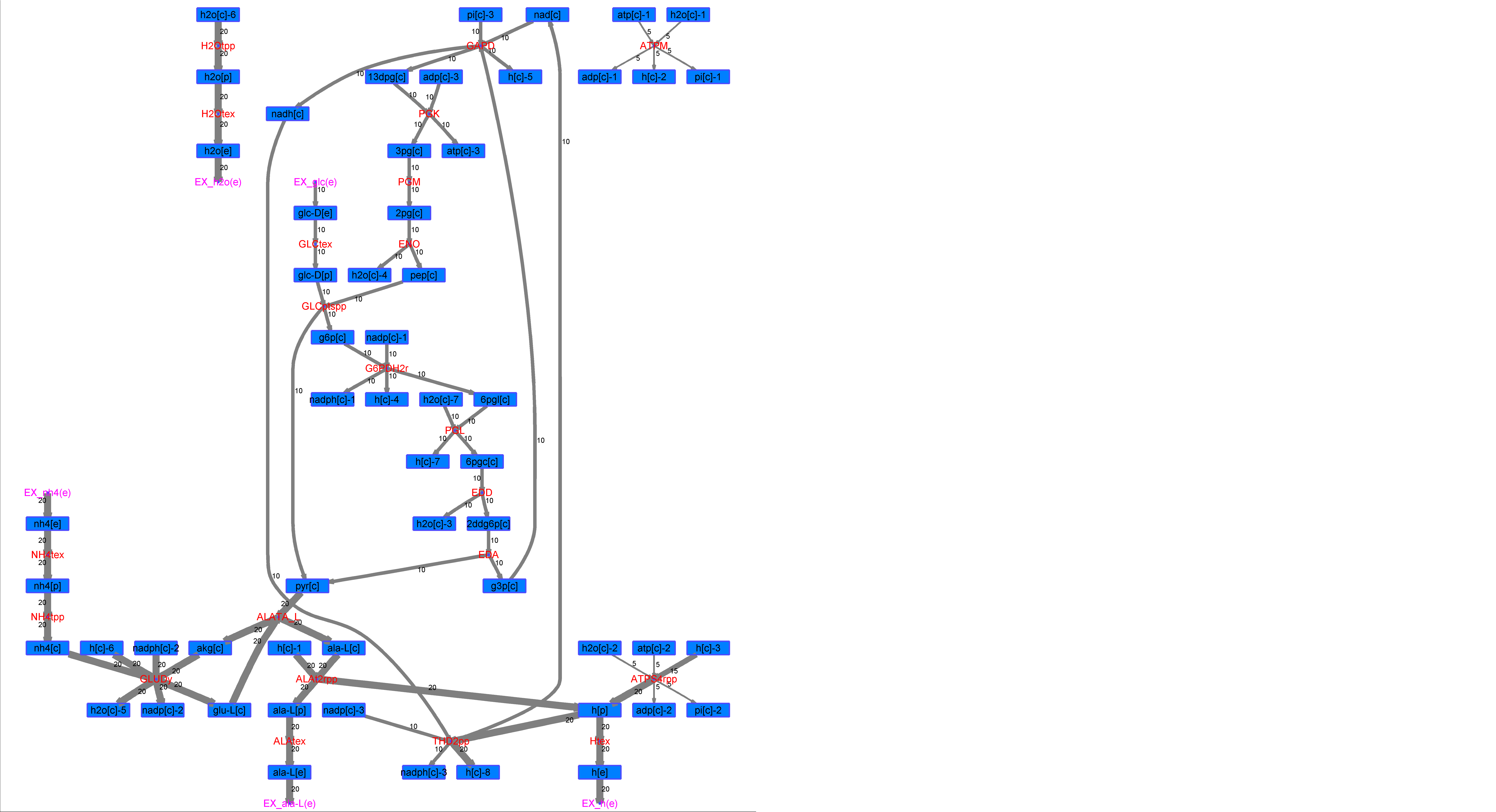

### R00343_L-Mal.png

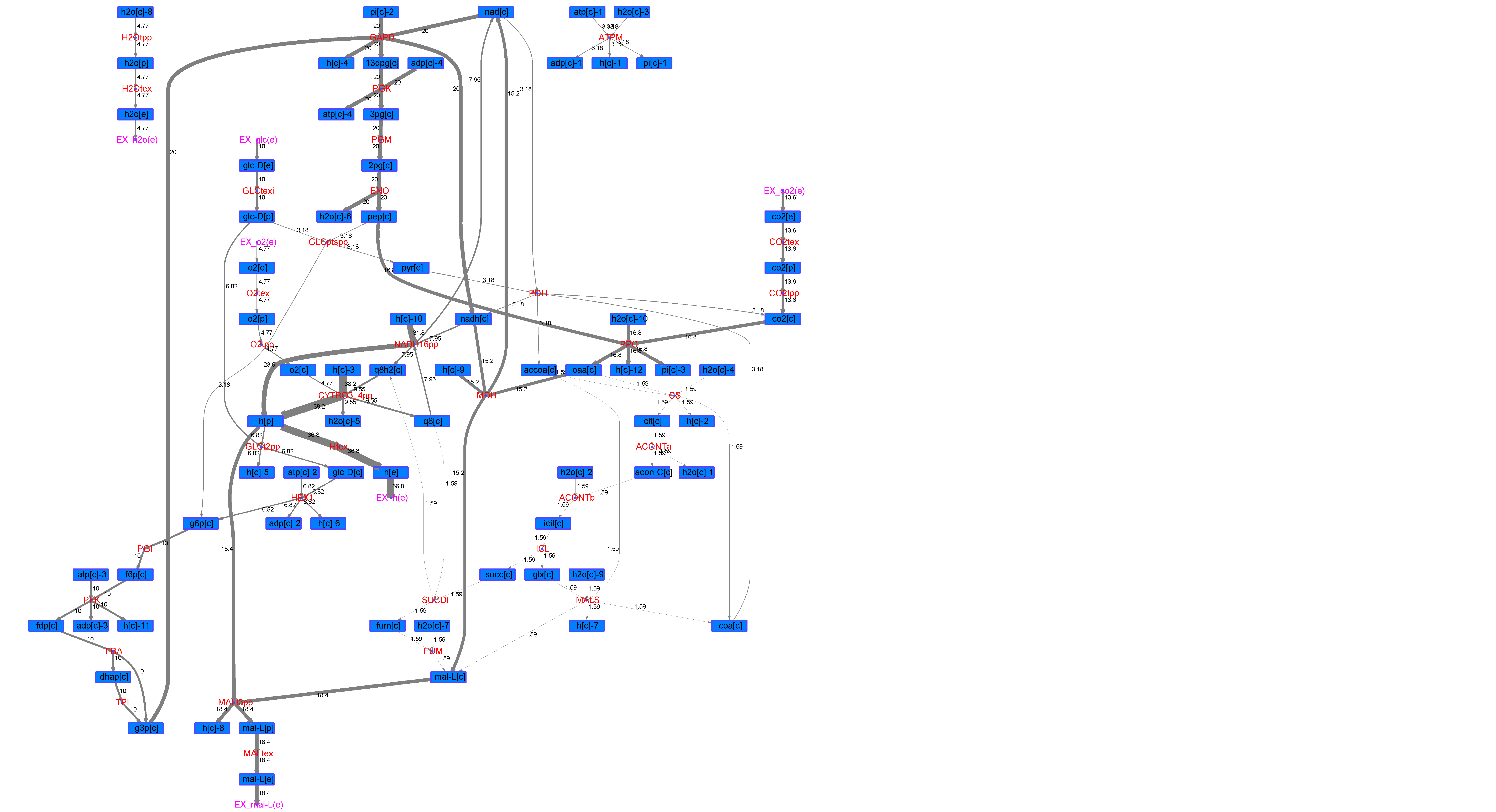

### R00343_L-Phe.png

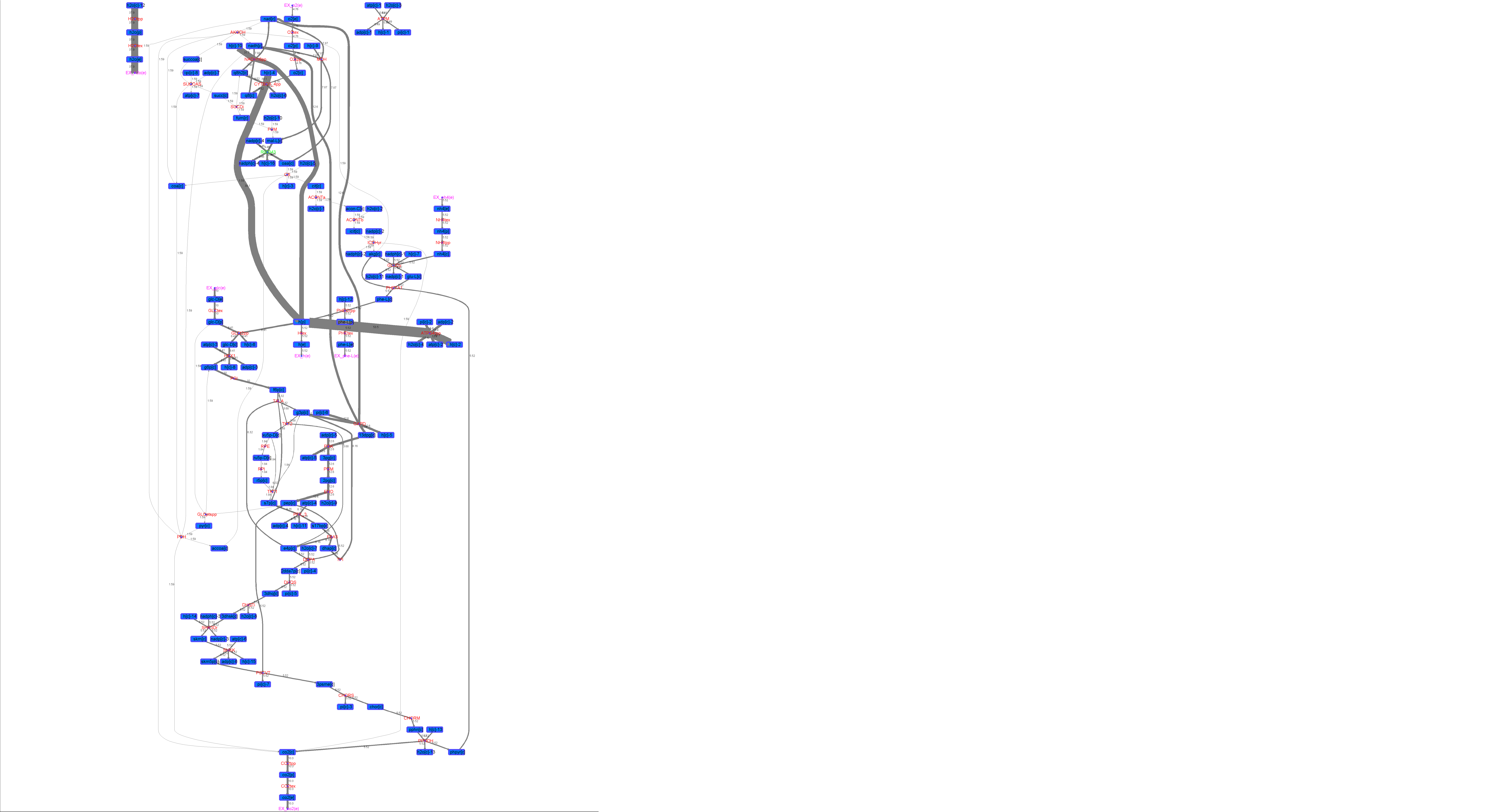

### R00343_L-Try.png

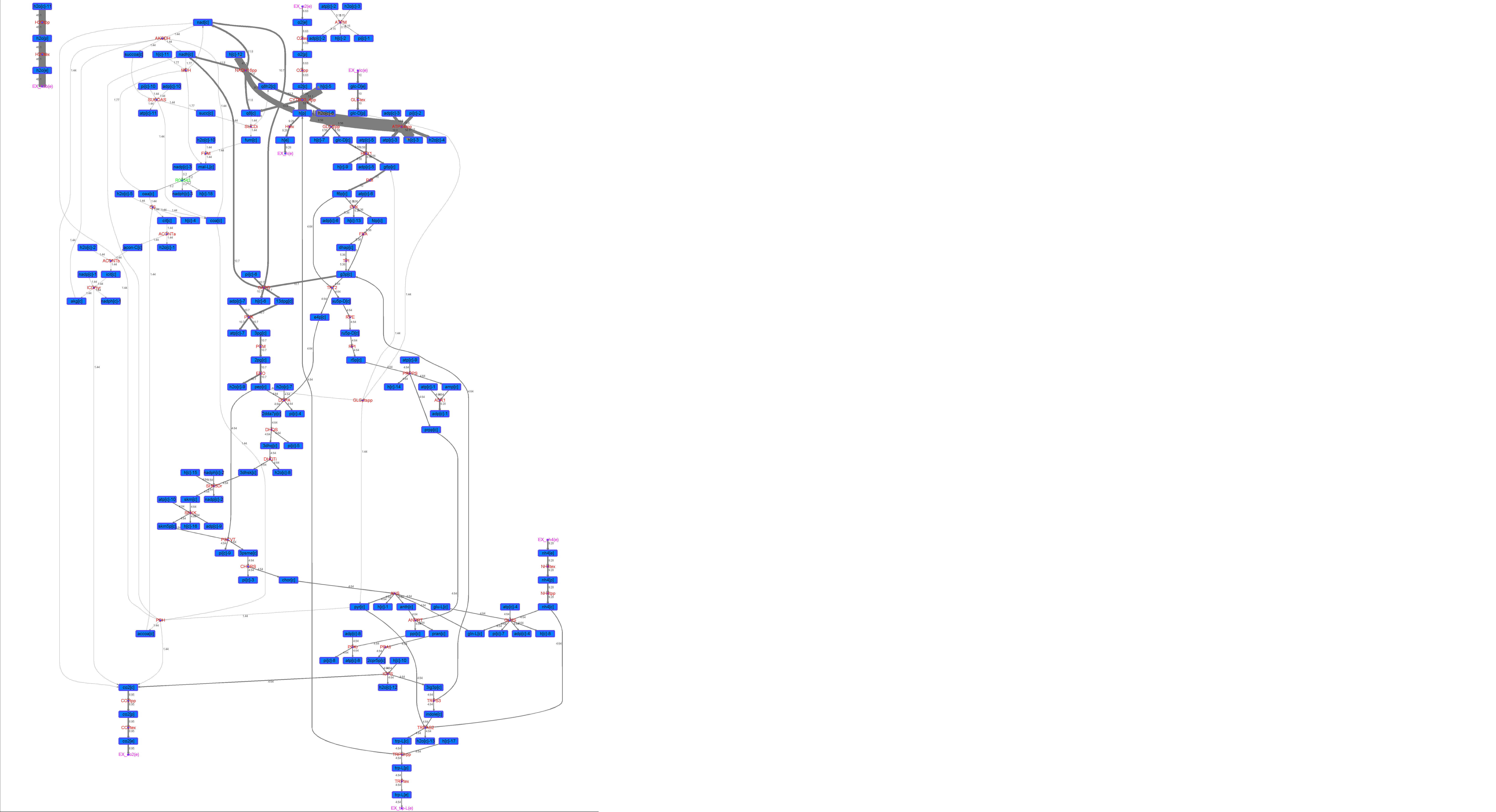

### R00343_Lys.png

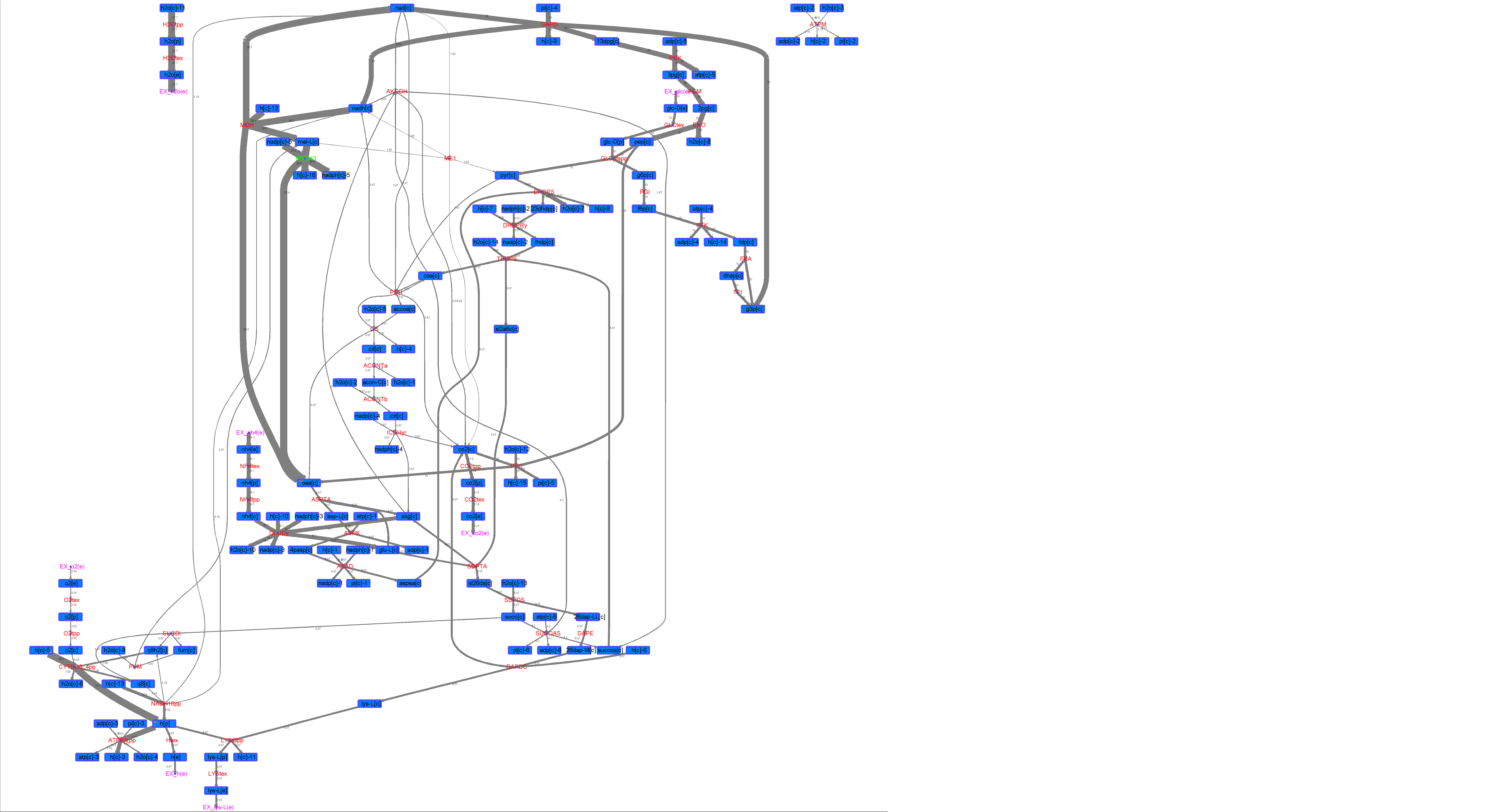

### R00343_Suc.png

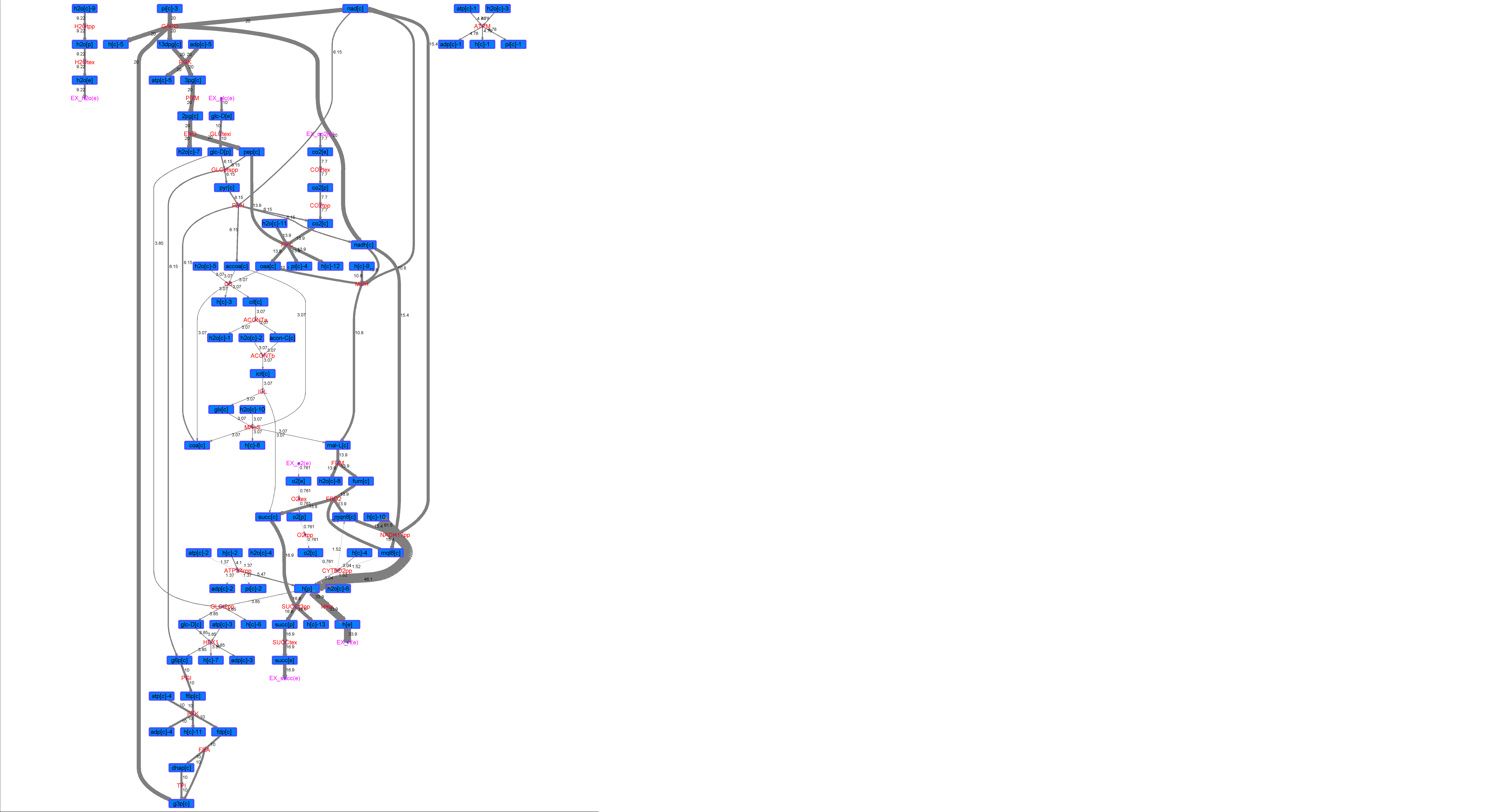

### R00343_Thr.png

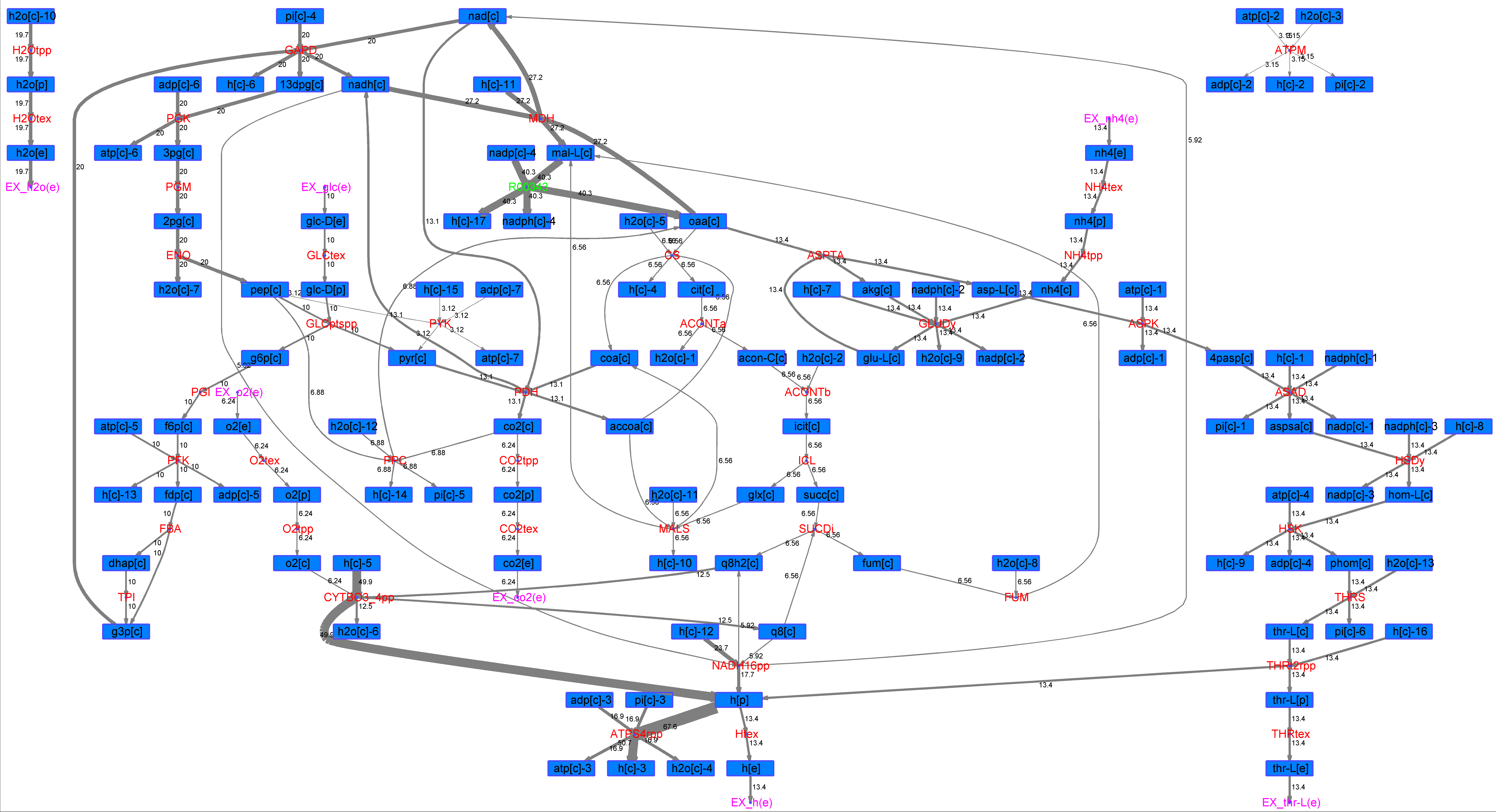

### R00346_Ace.png

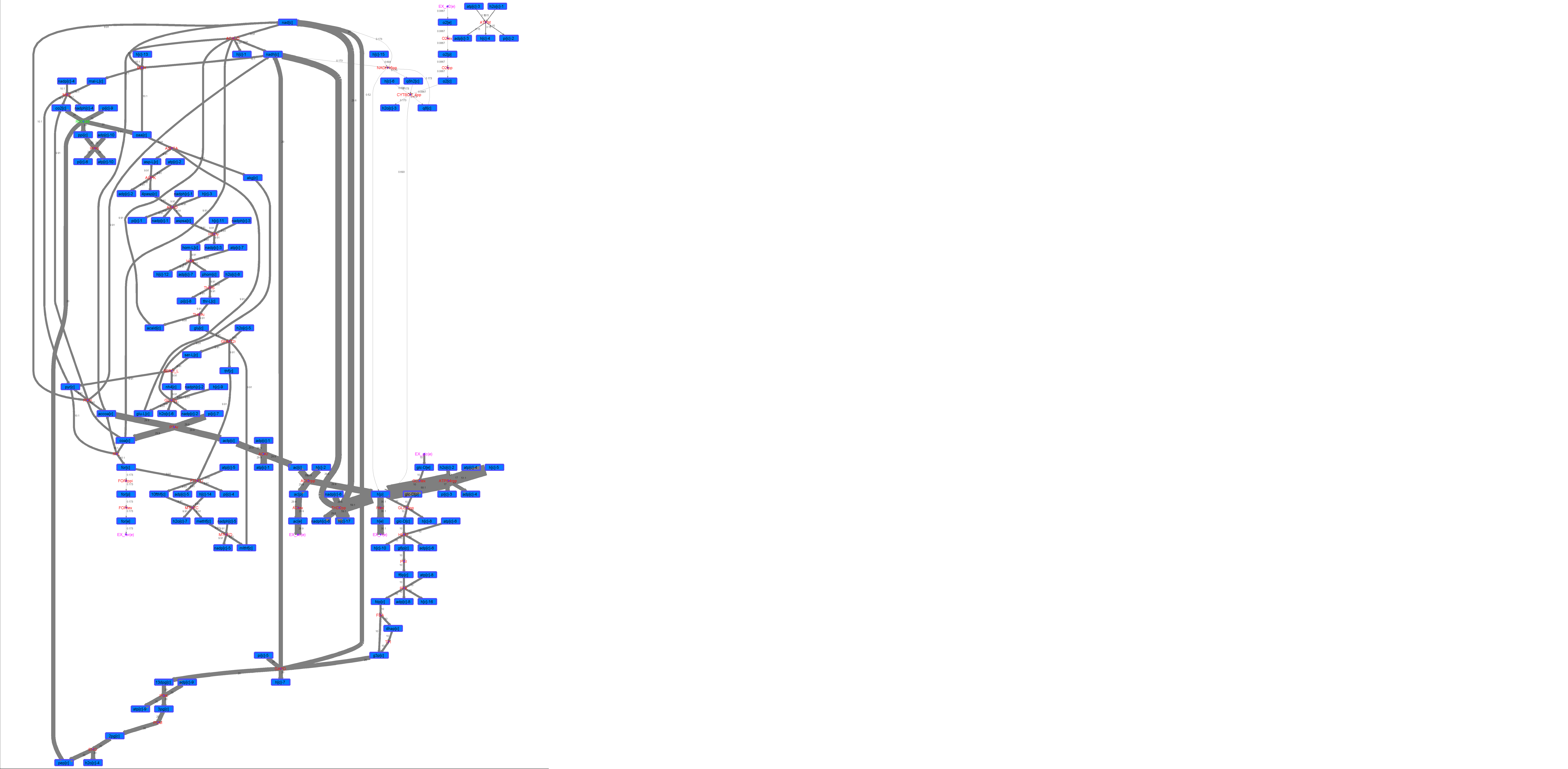

### R00346_For.png

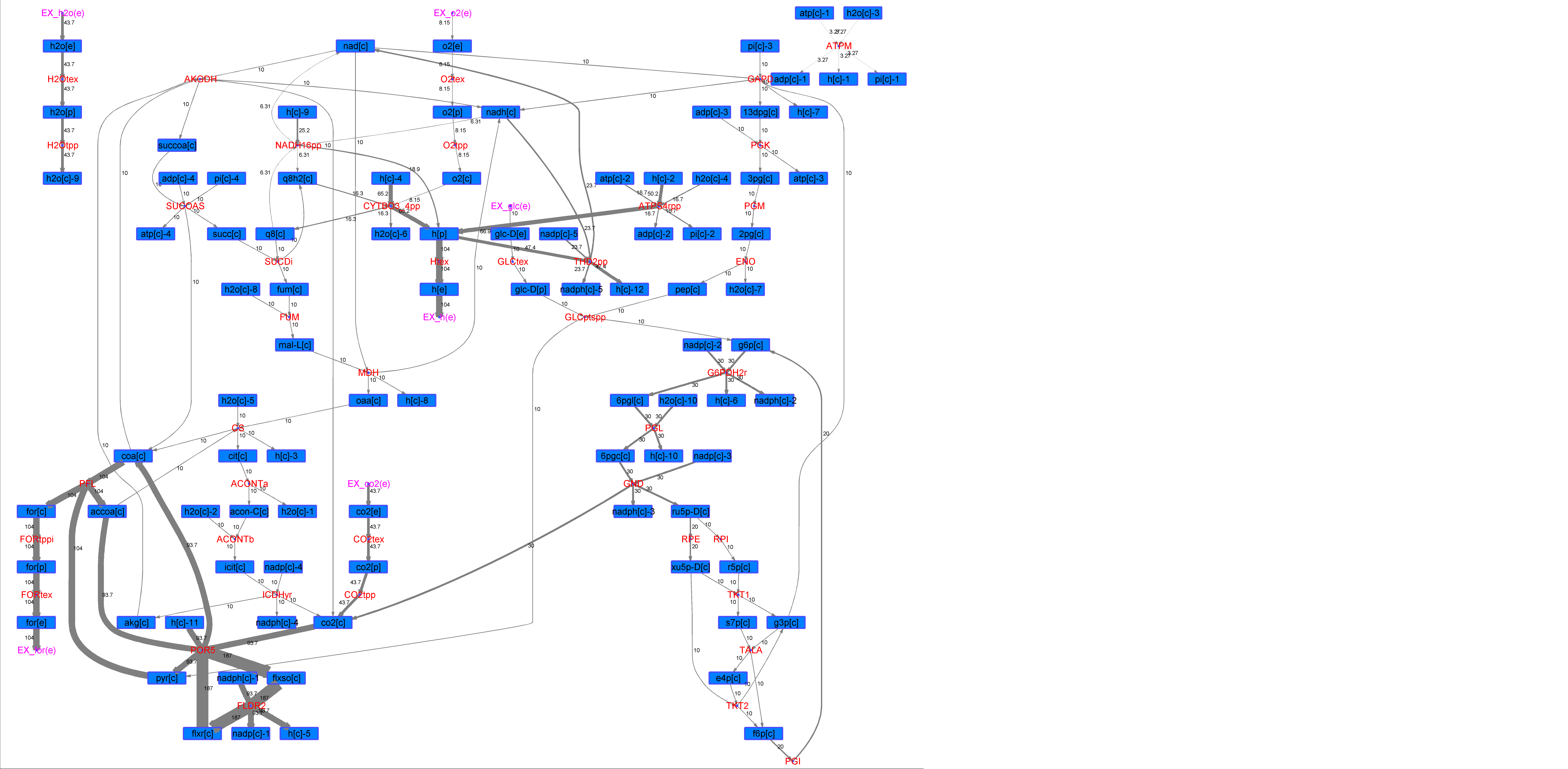

### R00346_Glu.png

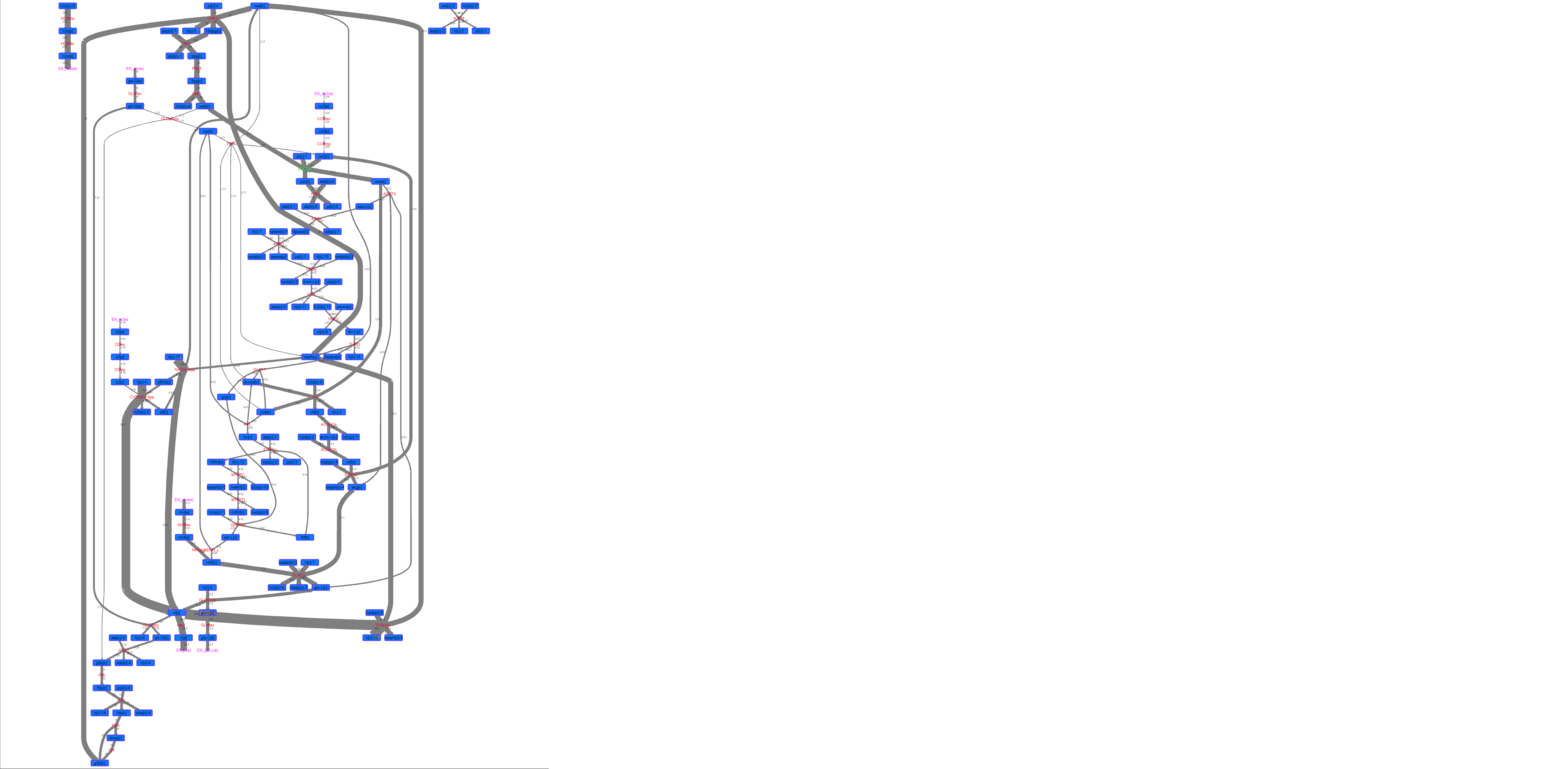

### R00346_L-ala.png

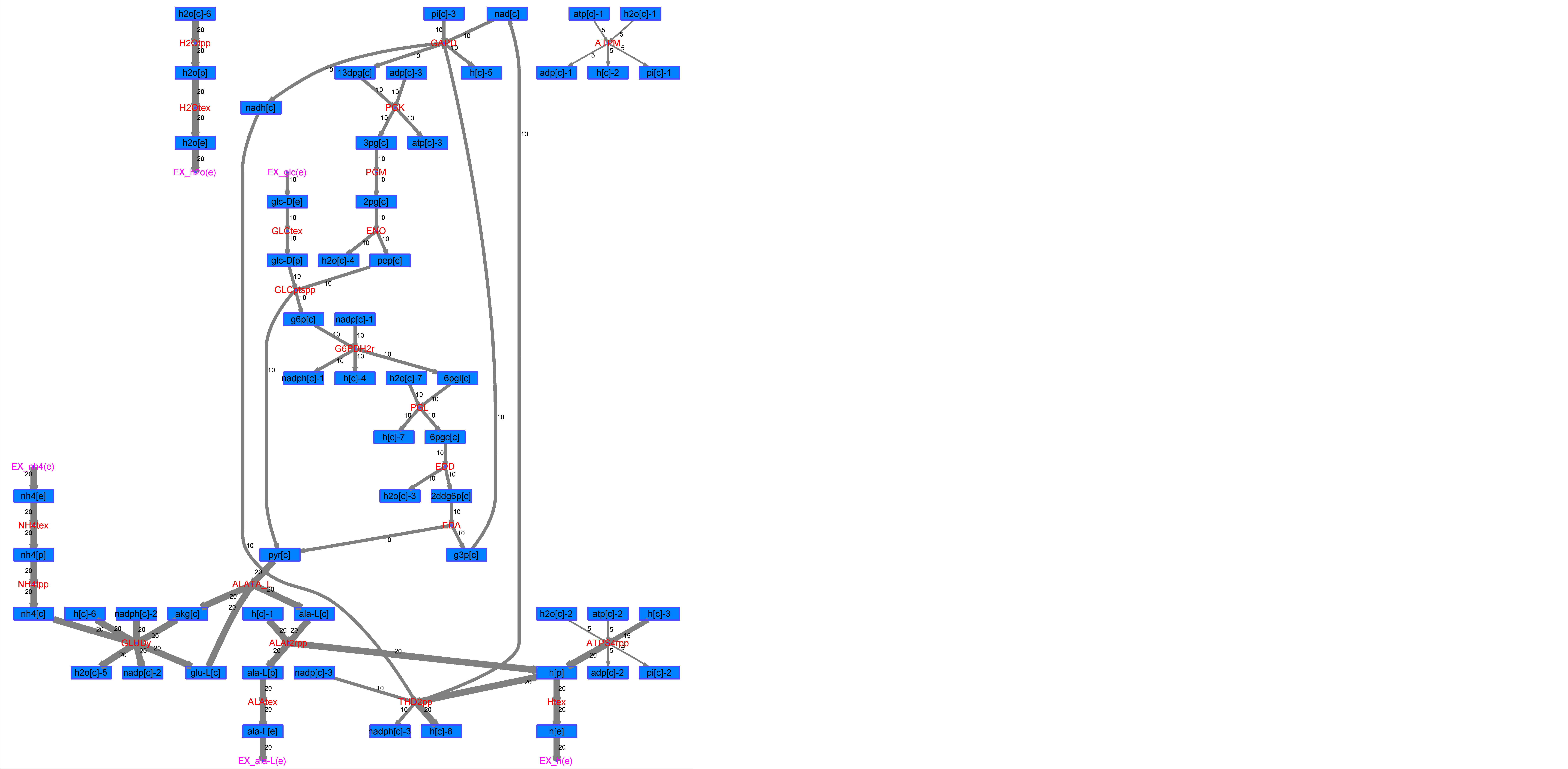

### R00346_L-Mal.png

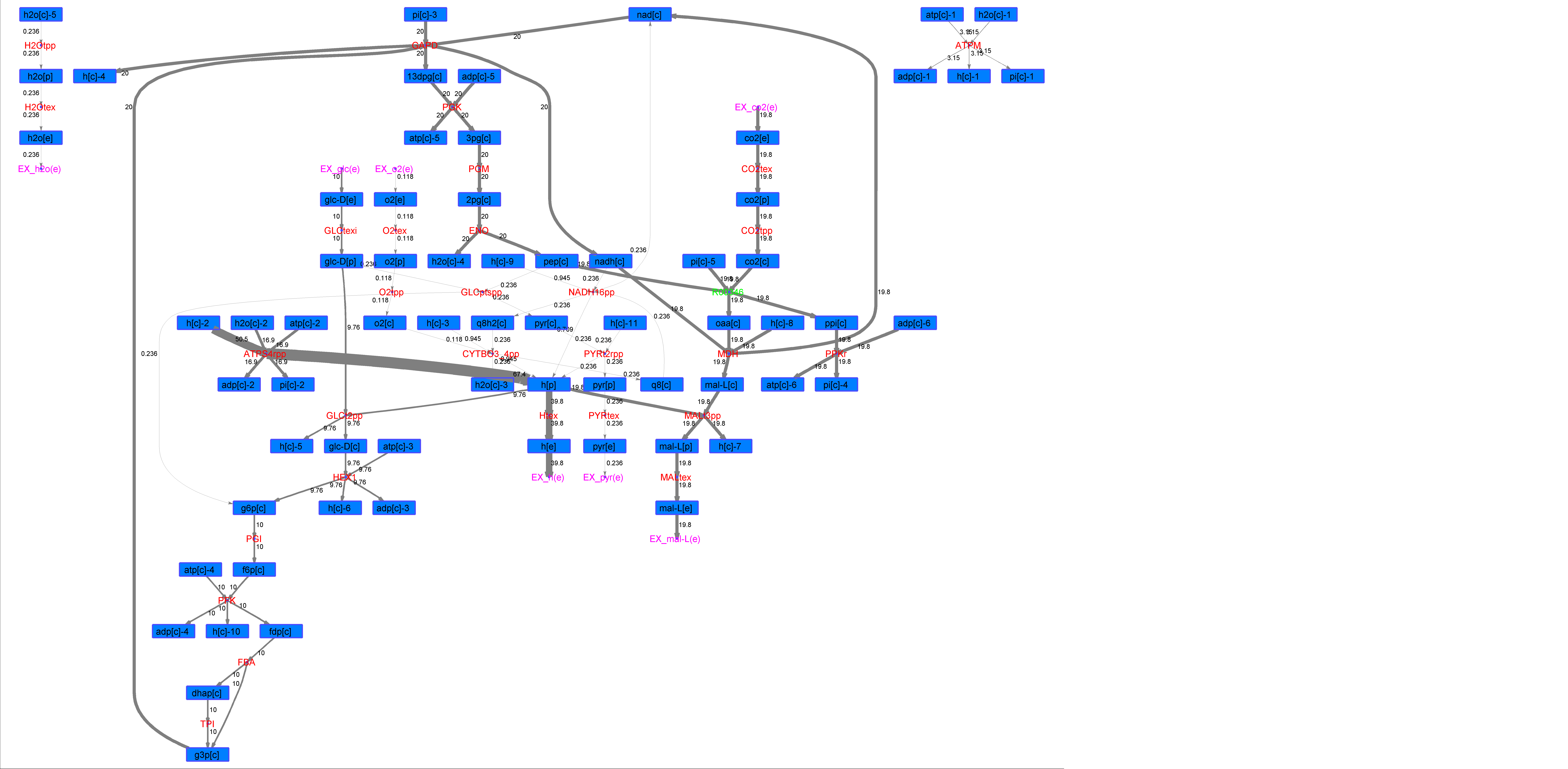

### R00346_L-Phe.png

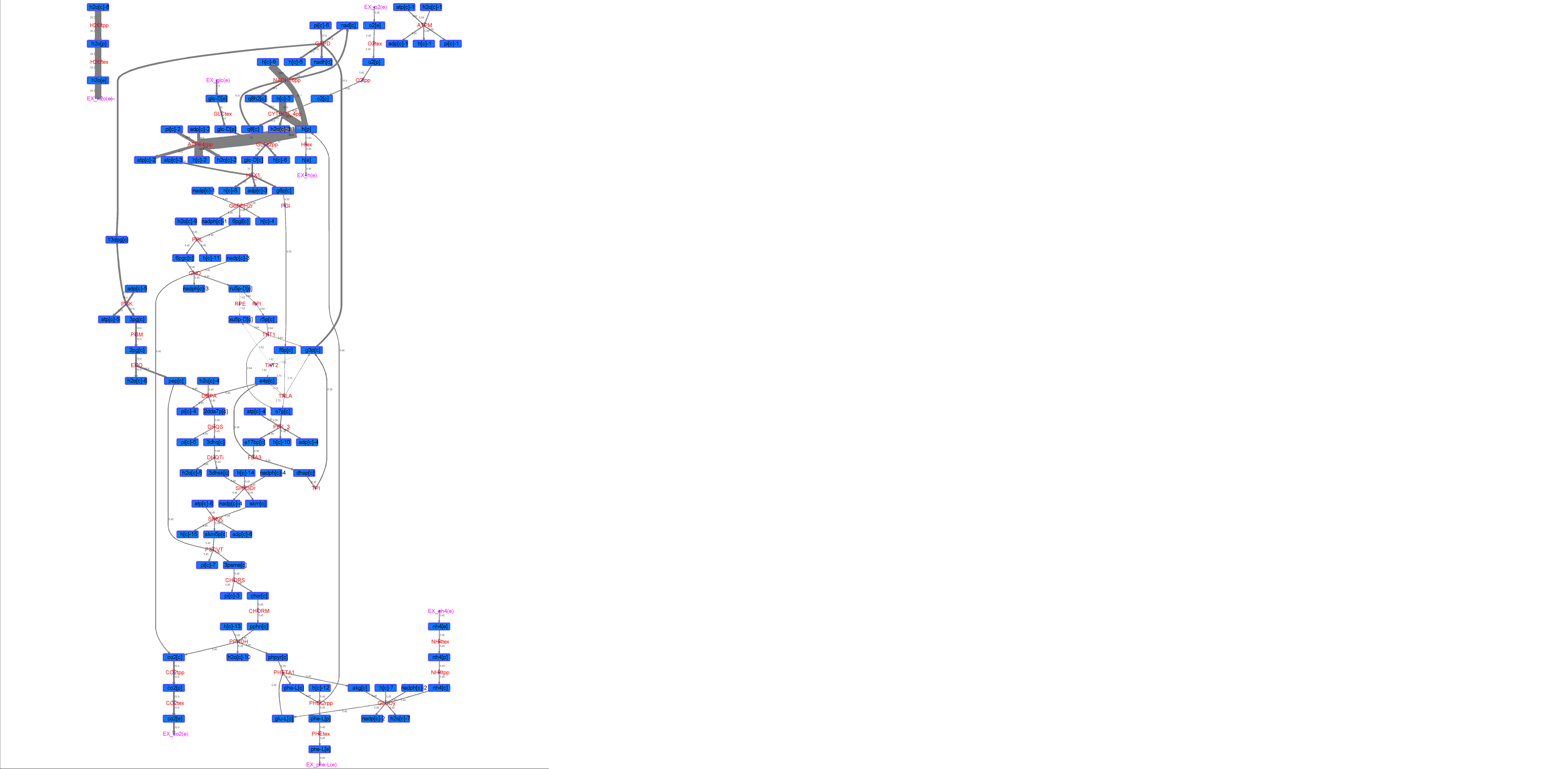

### R00346_L-Try.png

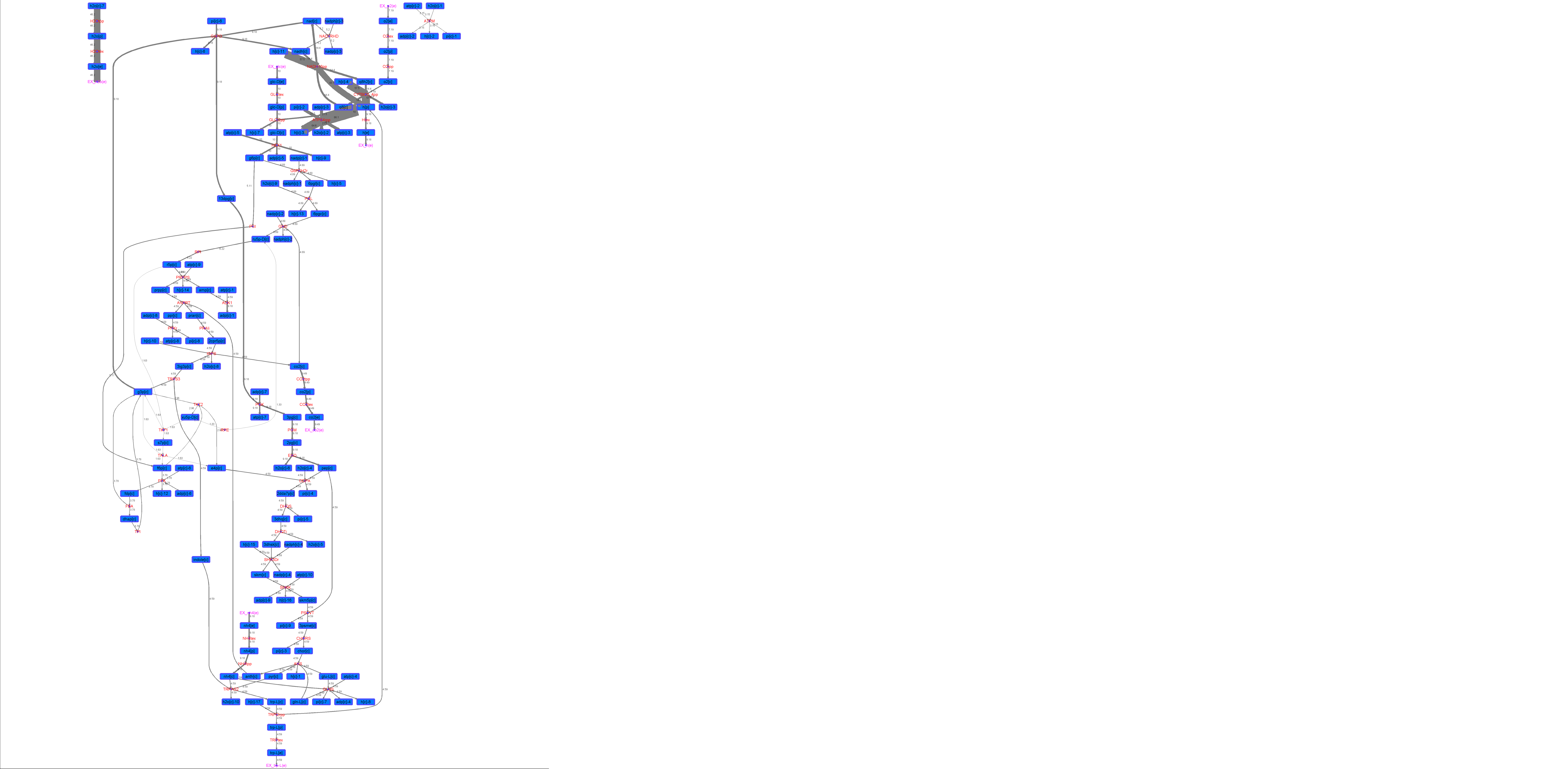

### R00346_Lys.png

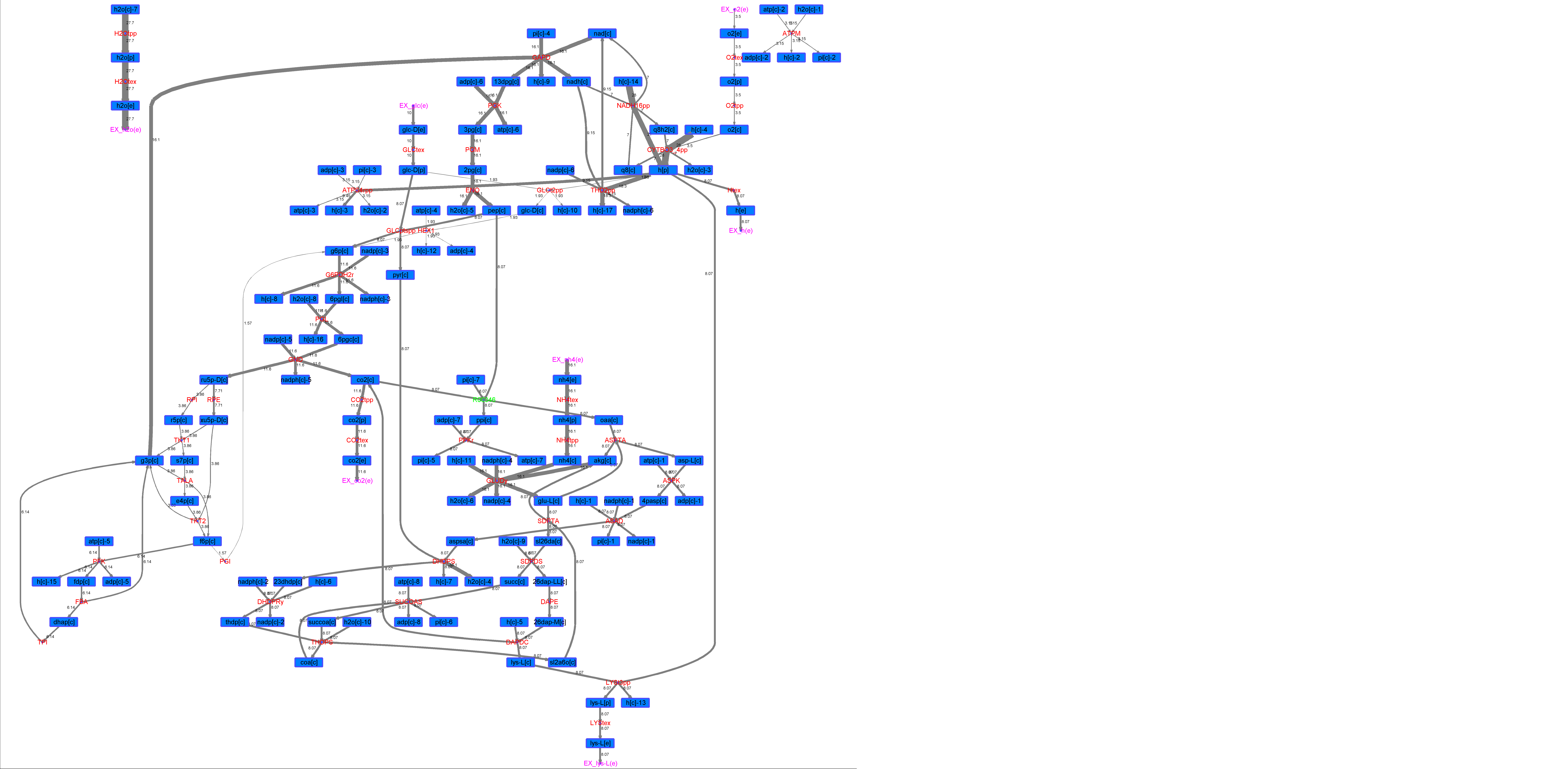

### R00346_Suc.png

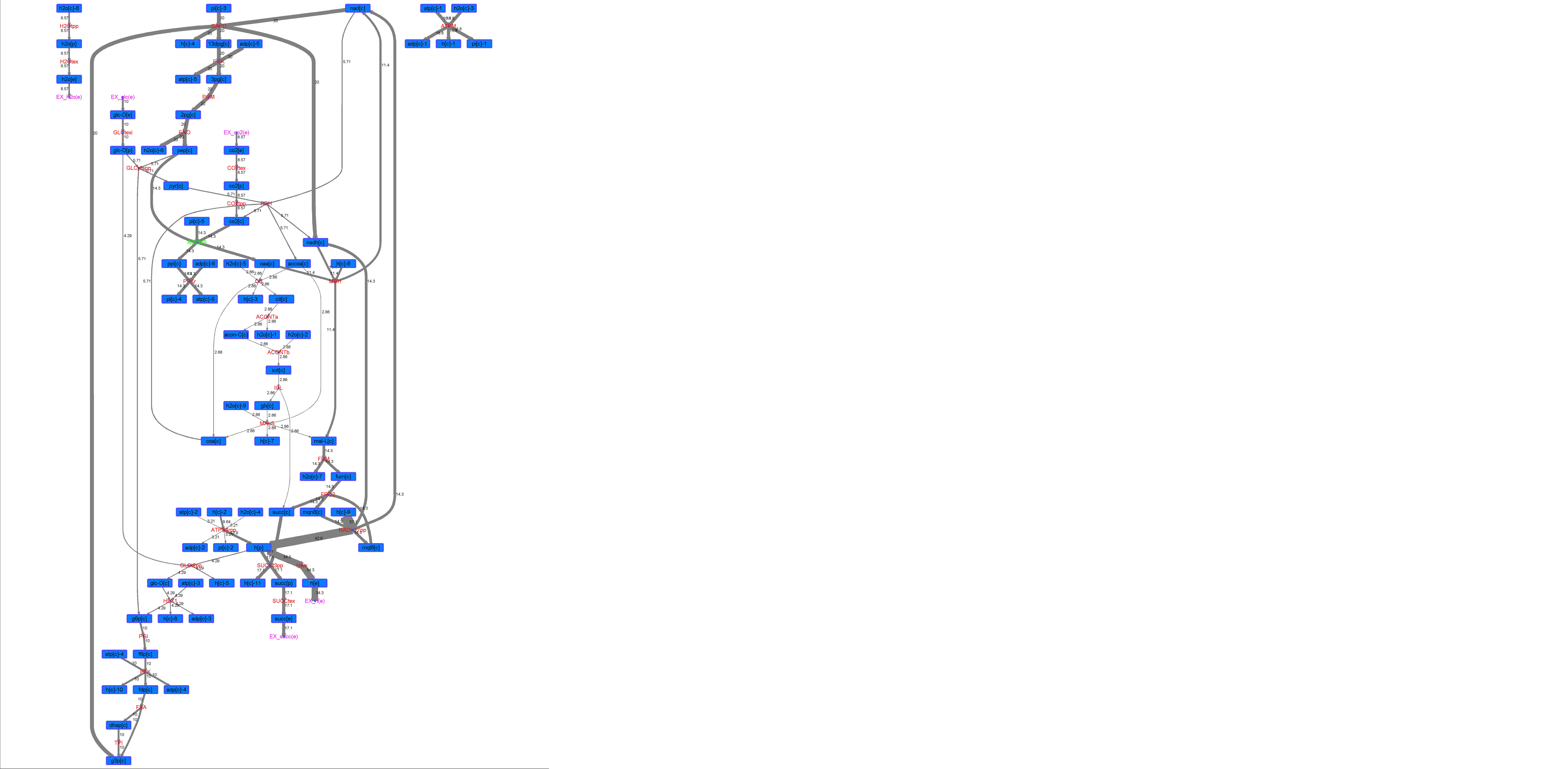

### R00346_Thr.png

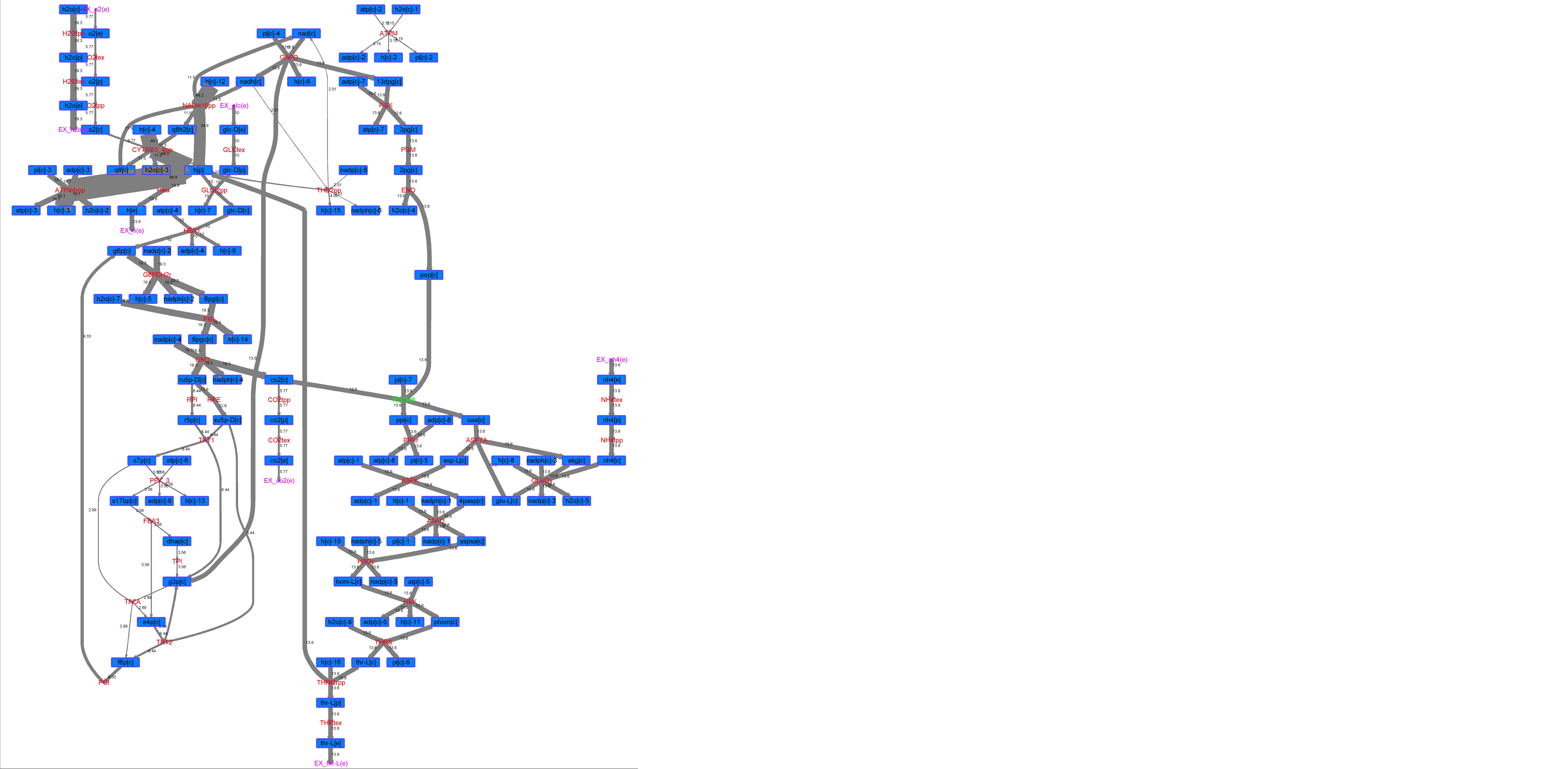
